# Generative Language Modeling for Antibody CDR Grafting and Alignment-driven De Novo Design

**DOI:** 10.64898/2026.09.06.749721

**Authors:** Ferran Gonzalez Hernandez, Oliver M Turnbull, Maliha Sultana, Lorena Roldán-Martín, Rohan J Kumar, Tom Diethe, Rebecca Croasdale-Wood, Charlotte Deane, Dino Oglic

## Abstract

Antibodies recognise their targets through hypervariable complementarity-determining regions (CDRs), which are interleaved with conserved frameworks in sequence space, making *de novo* CDR design an infilling problem. Autoregressive models generate residues left-to-right, which precludes full framework context during CDR generation and conflates framework and CDR likelihoods, leaving no natural prompt–response interface for feedback to steer generation. We present GenCDR, a family of LLaMa-based autoregressive language models that read all frameworks as a conditioning prompt and generate all CDRs jointly as a variable-length response, making CDR likelihoods a clean, separable target for reward attribution. The family comprises *IgGenCDR, p-IgGenCDR*, and *NanoGenCDR*, trained on unpaired, paired, and nanobody chains, respectively. GenCDR achieves the highest CDR recovery among autoregressive models and produces natural, diverse, human-like CDRs whose likelihoods correlate with fitness and developability assays. The prompt–response boundary also enables principled alignment: reward signals for binding affinity, expression, or developability can be composed to steer CDR generation. Over four rounds of alignment against antibody–antigen co-folding and developability objectives, we find that NanoGenCDR, which uses no explicit antigen encoding, can reach *in silico* structural interface metrics competitive with those of a structure-conditioned diffusion pipeline at roughly half the sampling budget, with more natural, developable designs. The same interface can be extended to integrate experimental feedback, opening a path to closed-loop antibody *de novo* design.

Code and pre-trained models for GenCDR are available at https://github.com/AstraZeneca/GenCDR.

## Introduction

Antibodies are fundamental to the immune response and represent a growing class of therapeutics owing to their ability to bind antigens with high specificity. The majority of antibody diversity, and thus antigen-binding capability, is concentrated within six hypervariable loops known as complementarity-determining regions (CDRs) of both heavy and light chains. A central challenge in therapeutic antibody development lies in the *de novo* design of CDRs that exhibit desired binding characteristics while maintaining favourable developability profiles, including low aggregation propensity, minimal polyspecificity, and acceptable expression and solubility (1, 2).

Protein language models have transformed protein engineering, enabling the generation of novel and diverse sequences with natural properties by learning evolutionary patterns from large sequence databases. Autoregressive (AR) protein language models, which predict the next amino acid given all preceding context, are particularly well-suited for sequence generation due to their scalability, natural support for variable-length outputs, well-established training paradigms, and compatibility with post-hoc alignment via reinforcement learning (3). Several AR models have been developed specifically for antibody generation, including p-IgGen (4) and ProGen2-OAS (3), which generate amino acids in canonical left-to-right order. However, therapeutic antibody *de novo* design is frequently framed as an infilling problem in which CDRs are designed conditioned on a given framework scaffold (5, 6). Standard left-to-right AR models are ill-suited to this task because when generating CDR1, the model has no context on subsequent frameworks or downstream CDRs, precisely the structural context constraining CDR composition and length.

IgLM partially addresses this limitation by masking a single variable-length span during training, appending it to the end of the sequence, and generating it conditioned on the full surrounding sequence (2). However, it handles only one span at a time and so cannot jointly design all CDR loops, since masked infilling of one loop presumes the others as context while they are themselves unknown in *de novo* design. The masked spans during training are also random windows of 10–20 residues rather than CDR-specific regions, and a single masked span yields no clean factorisation of sequence probability into framework and multi-CDR components, complicating CDR-specific scoring and reward attribution. Even combined with Proximal Policy Optimization (PPO), as demonstrated for single-loop CDR-H3 alignment (7), these properties prevent joint multi-CDR alignment. An alternative paradigm couples structure-based CDR generation via diffusion or hallucination with inverse folding, as in RFAntibody (8), BoltzGen (9), and Germinal (10). These pipelines directly incorporate antigen geometry and full framework context but require structural input at both training and inference times, limiting data scale relative to the hundreds of millions of sequences in repertoire databases. They also require pre-specifying the length of the designed region in sequence space and depart from the AR paradigm suitable for post-hoc alignment via reinforcement learning (7, 11, 12).

A key advantage of AR models is that their factorised log-probability provides a natural prompt–response interface for alignment methods such as Direct Preference Optimization (DPO) (13) or PPO (14). However, no existing AR antibody model cleanly exposes this interface for multi-CDR generation on a fixed framework scaffold. Left-to-right models interleave CDR and framework tokens, offering no natural boundary for reward attribution or variable-length CDR design. Hence, a pre-training paradigm that jointly generates all CDR loops at variable lengths in a single pass while providing a clean prompt–response interface for alignment is lacking.

In this work we introduce GenCDR, a family of framework-first autoregressive antibody language models trained explicitly for full-context CDR infilling. This ordering presents all four framework regions (FR1–FR4) as a conditioning prompt, after which the model generates CDR1, CDR2, and CDR3 sequentially in a single forward pass (Figure 1A). It can also generate a subset of CDR loops (e.g., CDR3 alone given CDR1 and CDR2) and naturally proposes variable CDR lengths compatible with the given frameworks. As the framework-first ordering separates framework and CDR loglikelihoods, it makes GenCDR the first AR antibody model family natively suited for joint multi-CDR generation and alignment.

**Fig. 1.**
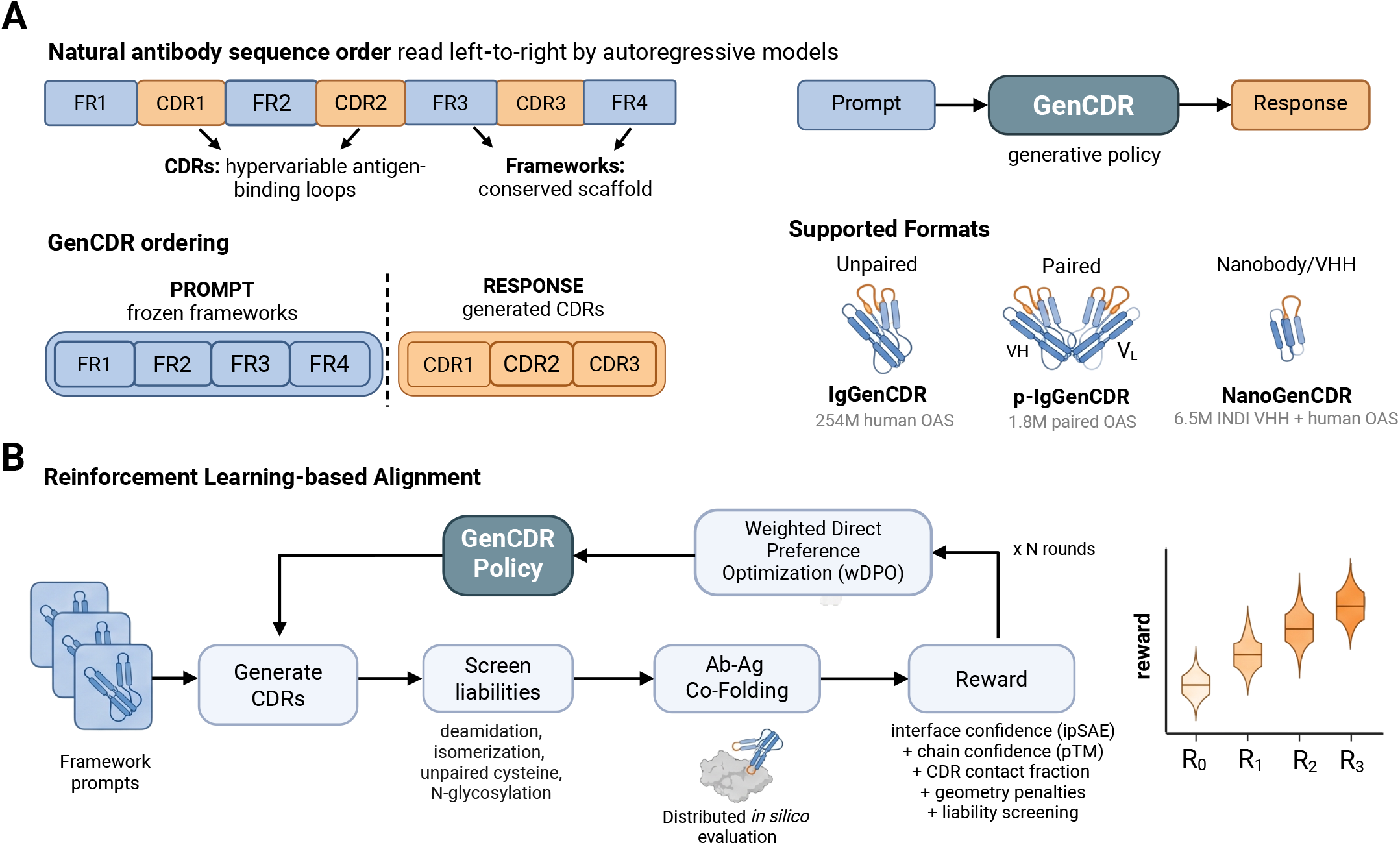
Framework-first antibody CDR generation and alignment. **(A)** GenCDR reads all four frameworks as a conditioning *prompt* and generates all CDRs jointly as a variable-length *response*, separating framework and CDR log-likelihoods. It supports three generative modalities trained on different data sources: unpaired chains (*IgGenCDR*), paired VH–VL (*p-IgGenCDR*), and nanobodies (*NanoGenCDR*). **(B)** The prompt–response boundary enables alignment. Each round generates CDRs on fixed prompts, screens liabilities, co-folds accepted designs against the antigen with Boltz-2, converts interface and geometry metrics into a composite reward, and updates the policy by scaffold-grouped weighted DPO (wDPO).

First, we show that framework-first pre-training achieves markedly better CDR amino acid recovery than prior AR baselines, including an identically sized left-to-right model p-IgGen (4), as well as ProGen2-OAS (3) and IgLM (2). Second, we show that framework-first conditioning can generate novel and high-quality CDRs as assessed by length distribution fidelity, humanness, naturalness, and diversity. Third, we train three model variants spanning the principal antibody design settings: *IgGenCDR*, pre-trained on ∼250M unpaired human sequences from the Observed Antibody Space (OAS) (15), *p-IgGenCDR*, fine-tuned on ∼1.8M natively paired VH–VL sequences from paired OAS to capture interchain dependencies, and *NanoGenCDR*, fine-tuned on a 1:1 mixture of camelid nanobody sequences from INDI (16) and unpaired human OAS sequences. Finally, we demonstrate how the framework-first paradigm can be used for alignment-driven *de novo* nanobody design by iteratively infilling CDRs prompted on four humanized nanobody frameworks against a target antigen. Starting from NanoGenCDR, we iteratively update the model weights through alignment with weighted DPO, where the model aims to maximize a composite structural reward combining nanobody–antigen co-folding metrics with non-differentiable developability liabilities, reaching *in silico* interface quality competitive with the structure-conditioned BoltzGen pipeline (9) at roughly half the sampling budget.

## System and Methods

### Data

*IgGenCDR* was pre-trained on unpaired human sequences from OAS (15), using the publicly released dataset of Turnbull et al. (4)^1^: ∼254M human antibody sequences (∼118M heavy, ∼136M light), quality-filtered, annotated with ANARCI (17), and clustered at 95% sequence identity. CDR and framework boundaries were then annotated under IMGT, Kabat, and Chothia numbering, with the scheme sampled uniformly per training example so that the model learns scheme-agnostic representations. We adopt the p-IgGen train and validation partitions as released. Because that partition clusters whole variable domains at 95% identity, two sequences can share a CDR yet differ in framework and fall into different splits, leaving it non-disjoint at the CDR level, so we rebuild the held-out test set and remove every sequence whose CDR3 lies within ≥ 95% length-matched identity of any training CDR3, retaining 202,138 heavy and 31,014 light sequences (Table 1, full construction and homology statistics in Supplementary Section S2). *p-IgGenCDR* was finetuned from the IgGenCDR checkpoint on ∼ 1.8M natively paired VH–VL sequences from paired OAS. *NanoGenCDR* was fine-tuned from the same checkpoint on a 1:1 per-batch mixture of ∼ 6.5M non-redundant camelid VHH sequences from INDI (16) and unpaired human heavy-chain OAS, the latter retained to preserve the humanness and diversity acquired during pre-training. The camelid corpus was processed through an eight-step filtering pipeline described in Supplementary Section S1. Every reported metric is computed on an independent, held-out test set.

**Table 1.** CDR amino acid recovery (next-token prediction accuracy, %, IMGT numbering) on the CDR-disjoint unpaired test set (Supplementary Section S2). p-IgGen is trained bidirectionally and is evaluated in both decoding directions. Subscripts are one bootstrap standard error (2000 sequence-level resamples). ^*†*^ Each CDR scored independently with full bidirectional context via IgLM’s single-span masked format (separate inference pass per CDR). Kabat and Chothia numbering in Supplementary Tables S6–S7. Stars mark the best-vs-second-best gap per column (paired Wilcoxon, Holm-corrected across loops): ^∗∗∗^*p* < 0.001, ^∗∗^*p* < 0.01, ^∗^*p* < 0.05.

| Model | Heavy chain (%) |  |  | Light chain (%) |  |  |
| --- | --- | --- | --- | --- | --- | --- |
|  | CDR1 | CDR2 | CDR3 | CDR1 | CDR2 | CDR3 |
| *Input-framing ablation* |  |  |  |  |  |  |
| IgGenCDR (framework-first) | **91.47 ± .03\*\*\*** | 89.81 ± .04 | **49.75 ± .04\*\*\*** | **85.63 ± .10\*\*\*** | **84.25 ± .13** | **58.54 ± .08\*\*\*** |
| IgGenCDR-LTR (left-to-right) | 88.56 ± .03 | **90.31 ± .04\*\*\*** | 48.76 ± .04 | 82.19 ± .10 | 84.15 ± .13 | 58.10 ± .08 |
| *Autoregressive baselines* |  |  |  |  |  |  |
| p-IgGen (left-to-right) (4) | 85.89 ± .04 | 85.39 ± .04 | 40.20 ± .03 | 77.61 ± .12 | 76.22 ± .14 | 50.38 ± .08 |
| p-IgGen (right-to-left) (4) | 88.20 ± .04 | 82.76 ± .05 | 40.33 ± .03 | 79.82 ± .11 | 75.85 ± .15 | 42.73 ± .08 |
| p-IgGen (developable, left-to-right) (4) | 86.02 ± .04 | 84.75 ± .04 | 39.93 ± .03 | 77.61 ± .12 | 75.83 ± .14 | 50.14 ± .08 |
| p-IgGen (developable, right-to-left) (4) | 87.99 ± .04 | 82.49 ± .05 | 39.74 ± .03 | 79.68 ± .11 | 75.60 ± .15 | 42.52 ± .08 |
| ProGen2-OAS | 85.64 ± .04 | 85.59 ± .04 | 41.38 ± .03 | 58.40 ± .20 | 43.88 ± .22 | 39.38 ± .12 |
| ProGen2-small | 68.44 ± .04 | 50.88 ± .05 | 21.17 ± .02 | 45.47 ± .11 | 39.03 ± .17 | 24.55 ± .08 |
| ProGen2-base | 80.92 ± .04 | 80.42 ± .05 | 33.16 ± .03 | 66.06 ± .15 | 63.55 ± .17 | 40.66 ± .09 |
| ProGen2-large | 81.73 ± .04 | 80.18 ± .05 | 32.81 ± .03 | 70.67 ± .14 | 67.00 ± .17 | 40.11 ± .10 |
| IgLM (CDR infill) | 88.52 ± .04<sup>†</sup> | 86.00 ± .04<sup>†</sup> | 39.09 ± .03<sup>†</sup> | 80.68 ± .11<sup>†</sup> | 77.81 ± .14<sup>†</sup> | 53.17 ± .08<sup>†</sup> |
| IgLM (full LTR) | 86.24 ± .04 | 85.63 ± .04 | 30.65 ± .03 | 79.77 ± .11 | 77.99 ± .14 | 41.08 ± .08 |
| IgLM-S (CDR infill) | 87.56 ± .04<sup>†</sup> | 84.85 ± .04<sup>†</sup> | 39.57 ± .03<sup>†</sup> | 78.73 ± .12<sup>†</sup> | 76.10 ± .15<sup>†</sup> | 49.34 ± .08<sup>†</sup> |
| IgLM-S (full LTR) | 85.94 ± .04 | 85.10 ± .04 | 32.40 ± .03 | 79.14 ± .11 | 76.88 ± .14 | 40.85 ± .09 |

### Architecture and Training

All GenCDR variants use a decoder-only transformer following the LLaMA architecture (18), with RoPE positional embeddings, RMSNorm, SwiGLU activations, and grouped-query attention, updating the GPT-2-based backbone of p-IgGen (4) and IgLM (2). Sequences are tokenised at the residue level. The models used throughout this work have 4 layers (∼25M parameters). Full architecture and tokenisation details are given in Supplementary Section S3.

All models were trained with cross-entropy loss over all input tokens, framework and CDRs. Computing the loss over framework tokens as well comes at no additional computational cost and keeps framework representations well-calibrated, which benefits full-sequence log-likelihood scoring. IgGenCDR was pre-trained for 10 epochs, with each batch comprising 80% unpaired single-chain inputs (heavy and light) and 20% randomly paired heavy–light inputs so that the model is exposed to both single-chain and cross-chain context. Both p-IgGenCDR and NanoGenCDR were then fine-tuned from the IgGenCDR checkpoint for 5 epochs, on natively paired VH–VL sequences and the camelid– human mixture respectively. Optimiser, schedule, precision, and batch-size settings are given in Supplementary Section S3.

### Framework-first input representation

The defining feature of IgGenCDR is a framework-first input ordering in which all four framework regions are presented as a conditioning prompt before any CDR token is generated. For a single chain (shown for heavy, with light symmetric using `<L>` and `<LPRED>`):

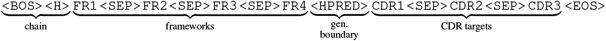

The numbering scheme (IMGT, Kabat, or Chothia) is sampled uniformly per training example so that the model learns scheme-agnostic representations. Paired heavy–light inputs use the same framework-first layout extended across both chains: all framework regions of both chains form a single joint conditioning prompt, followed by the two CDR blocks, each introduced by its own generation-boundary token, so that every CDR is generated with both chains’ frameworks in context and the model can capture inter-chain dependencies (Supplementary Section S9, Figure S9). During IgGenCDR pre-training this paired rendering is applied to 20% of batches by randomly pairing heavy and light chains to expose the model to the paired input formulation, whereas p-IgGenCDR is then fine-tuned on natively paired VH–VL sequences.

### Antigen-aware alignment for *de novo* design

Pre-training teaches the model the natural distribution of antibody sequences and yields a framework-conditioned CDR generator. Antibody engineering, however, is usually driven by specific objectives, ranging from developability properties such as expression, stability, or reduced immunogenicity to binding a chosen antigen. A growing body of work has shown that pre-trained protein and antibody language models can be steered toward such biophysical and functional objectives with reinforcement learning, mirroring how language models are aligned by human feedback toward preferred responses (7, 11, 19). Antigen-conditioned *de novo* design is a particularly demanding instance, requiring all CDR loops to be generated jointly to meet target-specific structural and biophysical constraints that are absent from the natural distribution. GenCDR models make the alignment step especially direct for this task: framework tokens form a fixed prompt *x* and CDR tokens a variable-length response *y*, giving offline RL an unambiguous boundary for reward attribution. Here, we test whether such models can be steered effectively toward realistic nanobody design objectives against a target of interest. To that end, we define a reward that resembles the objectives of real structure-based design pipelines (10): nanobody–antigen interface confidence, nanobody chain foldability, CDR shape geometry, and the absence of developability liabilities. This last term makes our setting distinct: liabilities are discrete, non-differentiable sequence-level rules that cannot be expressed as a differentiable objective, and are therefore harder to control for pipelines based on inverse folding or hallucination. Alignment instead lets the policy learn to avoid them directly in sequence space by learning motifs from liable sequences. We align NanoGenCDR using weighted DPO (11, 19), which operates directly on scalar rewards without constructing explicit preference pairs.

### Weighted DPO

Standard DPO (13) aligns a policy *π*_*θ*_ toward preferred outputs by contrasting winner–loser pairs, regularised by KL divergence from a frozen reference *π*_ref_. When continuous scalar rewards are available, weighted DPO (11, 19) replaces the pairwise sigmoid with a listwise cross-entropy over responses to a *shared* prompt. For a fixed framework scaffold *x* and *K* responses with rewards 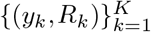 sampled from it, the loss is:

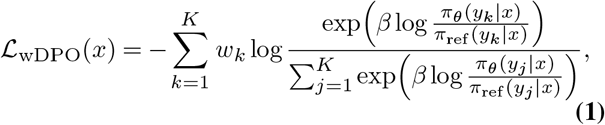

i.e. a cross-entropy between the soft targets *w*_*k*_ = softmax(**R**)_*k*_ derived from the scalar rewards and the logits 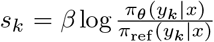, the *β*-scaled sequence log-likelihood ratios. We adapt this objective to CDR infilling in two ways. First, the shared prompt *x* is a fixed framework scaffold, and each log-ratio is a mean over CDR completion tokens only rather than the full sequence (Supplementary Section C). Second, since the listwise softmax is meaningful only among responses to the same prompt and a training batch spans all four scaffolds, we apply Eq. 1 separately per scaffold and average the per-scaffold losses weighted by their sample count (Eq. S4). Each framework is generated in equal numbers per round, so no scaffold dominates the training pool, and the sample-count weighting keeps each batch update proportional to the scaffolds it contains (Supplementary Figure S3). We set a conservative *β*=0.15 (relative to the 0.01–0.1 used for single-property protein alignment (11)) to limit drift from the reference and mitigate reward hacking, and freeze *π*_ref_ throughout.

### Iterative alignment loop

Each round proceeds in five steps (Figure 1B, detailed in Figure S1): (1) *generate* CDR completions from the current policy, (2) *screen* against CDR-length and liability rules, labelling sequences ACCEPT, REJECT, or HARD REJECT, (3) *score* ACCEPT sequences via Boltz-2 co-folding, (4) *assign rewards* (composite for ACCEPT, surrogate bands for rejected sequences), and (5) *align* with the grouped wDPO loss (Eq. S4). The updated model becomes the policy for the next round.

### Composite reward

Following recent structure-conditioned design pipelines that optimise co-folding confidence and interface geometry (9, 10), we define a composite reward that the purely sequence-based model must learn to satisfy: high-confidence nanobody–antigen interfaces, CDR-localised binding, and realistic loop geometry:

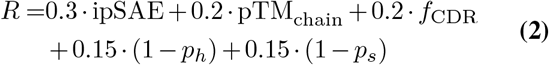

where ipSAE (20) is an interface-confidence proxy for binding quality and pTM_chain_ the predicted TM-score of the nanobody chain. The remaining three terms are adapted from the Germinal pipeline (10) and evaluated as post-hoc metrics on the Boltz-2 co-folded structures. *f*_CDR_ is the fraction of paratope contacts on CDR rather than framework residues, a counterbalance to ipSAE ensuring that interface confidence is driven by CDR-mediated binding. The geometry penalties (1 V *p*_*h*_) and (1 − *p*_*s*_) discourage regular *α*-helical or *β*-strand structure in the predicted CDR loops, which natural CDRs (especially CDR-H3) rarely adopt and which could otherwise inflate co-folding confidence. Full metric definitions are given in Supplementary Section B.

## Results

### A. IgGenCDR

We first measured CDR recovery, the next-token prediction (NTP) accuracy over CDR tokens in teacher-forcing mode (3, 4), to evaluate how faithfully each model reconstructs held-out CDRs from their surrounding framework context. Table 1 compares IgGenCDR with published AR baselines on the disjoint held-out test set (IMGT numbering, Kabat/Chothia in Supplementary Tables S6–S7). Additionally, we trained an identically sized left-to-right model (IgGenCDR-LTR) on the same dataset to ablate the effect of framework-first formulation under identical settings. For IgLM (2) we report both full left-to-right and CDR-infill modes, where we run three separate inference runs per sequence masking CDR1, CDR2, and CDR3 respectively. Framework-first ordering yields gains over the LTR ablation on five of the six loops, with the largest improvement on CDR1, the loop that benefits most from downstream framework context absent in LTR decoding (21). IgGenCDR also surpasses all published baselines on every loop. IgLM’s CDR-infill mode narrows the gap by providing bidirectional context for each loop individually, underscoring the value of surrounding-sequence conditioning, but cannot supply simultaneous context for all three CDRs in a single pass. The gains over published baselines likely reflect the combination of the framework-first factorisation itself, the updated LLaMA-style decoder departing from the GPT-2 backbone used by previous autoregressive models, and training exclusively on unpaired human sequences (in contrast to IgLM and ProGen which train across species, or p-IgGen which fine-tunes on paired OAS). These patterns are consistent across numbering schemes (Supplementary Tables S6–S7). Because p-IgGen is trained bidirectionally, we report it in both decoding directions: right-to-left, where decoding modestly improves CDR1 recovery but lowers L-CDR3 recovery, and left-to-right, but neither direction approaches the framework-first model. A more detailed per-baseline discussion is provided in Supplementary Section S6.

### Generated CDRs are natural, diverse, and human-like

Having established recovery, we next assessed the quality of *de novo* generated CDRs, to verify that framework-first conditioning produces realistic and diverse loops rather than merely reconstructing held-out ones. We prompted IgGenCDR with 2,000 held-out framework scaffolds from the test set and generated all CDR loops simultaneously across multiple sampling temperatures (under fixed top-*p* = 0.95), computing all downstream metrics under IMGT numbering. Figure 2 compares the CDR length distributions of generated (*T* = 1.0) and natural test-set sequences. For all six CDR loops the generated and natural distributions overlap almost perfectly, confirming that IgGenCDR faithfully captures both the germline-determined length preferences of CDR1 and CDR2 and the broader V(D)J-driven CDR3 distribution. This overlap is maintained for sampling temperatures up to 1.1 and broadens only beyond it, mirroring the smooth degradation of the other quality metrics at higher temperatures (Supplementary Figures S5–S8). The paired (p-IgGenCDR) and nanobody (NanoGenCDR) variants reproduce their respective reference distributions equally faithfully (Supplementary Figures S10 and S11).

**Fig. 2.**
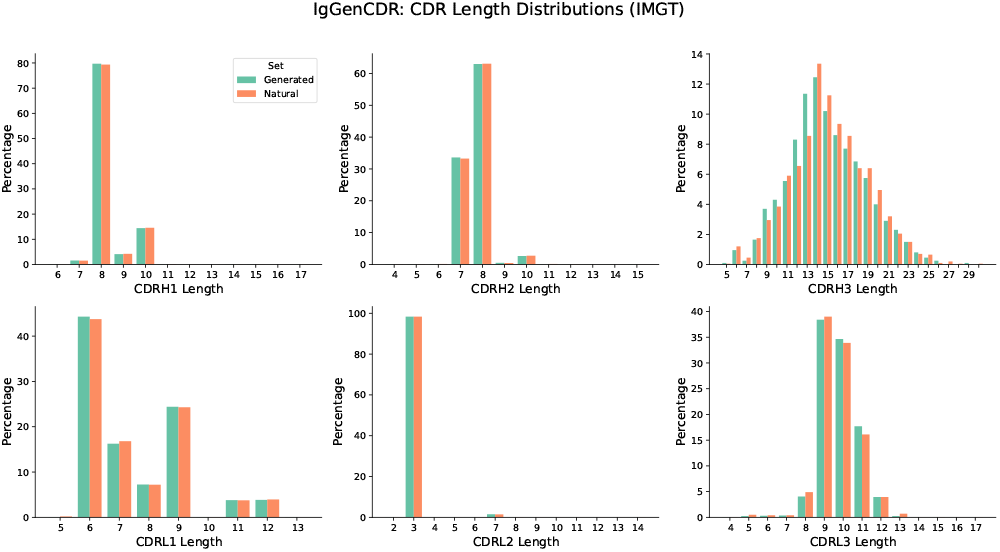
CDR length distributions (IMGT) for IgGenCDR-generated sequences at *T* = 1.0 (green) versus natural OAS test-set sequences (coral). Heavy chain (top) and light chain (bottom); CDR1, CDR2, CDR3 from left to right. Bars show the per-length proportion within each set.

We additionally evaluated humanness (OASis 9-mer scores (22)), naturalness (ESM-2 pseudo-log-likelihood per residue (23)), and sequence novelty (minimum Hamming distance to the training set) across the full temperature sweep. All metrics degrade smoothly at higher temperatures while novelty increases monotonically (Supplementary Figure S8). Together, these results suggest that generating three CDRs simultaneously at temperatures in the range *T* ∈ [0.90, 1.15] yields loops that are novel, human-like, and natural. Full temperature-sweep figures are provided in Supplementary Figures S5–S8.

### Likelihoods retain fitness-predictive signal

Beyond generation, a model’s likelihoods are useful only if they track antibody fitness, so we next tested whether the framework-first pre-training objective preserves fitness-predictive signal. We measured zero-shot correlation with antibody fitness assays from the FLAb benchmark (24, 25), scoring each sequence by its mean per-token log-likelihood (Supplementary Section S8), restricted either to all variable-domain tokens (full-sequence scoring) or to CDR tokens only (CDR scoring). The CDR score isolates the loops the model must design, whereas the framework and full-sequence scores probe the surrounding context. For IgGenCDR the CDR score uses the framework-conditioned factorisation that matches its training objective. We compared against the left-to-right ablation (IgGenCDR-LTR) and IgLM (2), the most architecturally similar infilling baseline. Results averaged across IMGT, Kabat, and Chothia numbering schemes are shown in Table 2, with per-scheme results in Supplementary Table S8. Among the unpaired models, IgGenCDR maintains comparable or superior correlation to the left-to-right baseline and substantially outperforms IgLM across both datasets and all sequence regions, indicating that the framework-first factorisation preserves fitness-predictive signal in its loglikelihoods. The two assays also localise to different antibody regions, reflecting that distinct regions govern distinct properties: the immunogenicity assay of Marks et al. (24), which derives from humanization mutations, is the more CDR-driven, with CDR scoring reaching *ρ*=0.227, whereas the expression assay of Koenig et al. (25) is dominated mostly by framework signal and shows near-zero CDR correlation. The paired models (p-IgGenCDR, p-IgGenCDR-LTR), discussed in Section B, score each H+L pair jointly and are reported in the same table for comparison. These patterns are robust across numbering schemes (Supplementary Table S8).

**Table 2.** Log-likelihood–fitness correlation (Spearman *ρ*) on FLAb for paired heavy+light chain sequences, averaged across numbering schemes (IMGT, Kabat, Chothia). Unpaired models score each chain independently and average, whereas paired models score the H+L pair as a single joint sequence. **Bold**: highest *ρ* in each column within each model group.

| Model | Marks et al. (24) (n=217) |  |  | Koenig et al. (25) (n=4276) |  |  |
| --- | --- | --- | --- | --- | --- | --- |
|  | Full | Framework | CDR | Full | Framework | CDR |
| *Independent-chain scoring (pair-mean)* |  |  |  |  |  |  |
| IgGenCDR | **0.351** | **0.317** | **0.227** | **0.238** | 0.203 | **0.089** |
| IgGenCDR-LTR | 0.310 | 0.302 | 0.226 | 0.237 | **0.214** | 0.044 |
| IgLM | 0.244 | 0.298 | 0.165 | 0.211 | 0.199 | 0.044 |
| *Joint paired scoring* |  |  |  |  |  |  |
| p-IgGenCDR | **0.374** | **0.328** | 0.227 | 0.296 | 0.265 | 0.020 |
| p-IgGenCDR-LTR | 0.373 | 0.317 | **0.275** | **0.327** | **0.283** | **0.041** |

### B. p-IgGenCDR

p-IgGenCDR is fine-tuned from the IgGenCDR checkpoint on ∼ 1.8M natively paired VH–VL sequences from paired OAS. Presenting both heavy- and light-chain frameworks as a joint conditioning prompt (Supplementary Section S9) lets the model learn the inter-chain co-evolutionary dependencies inaccessible to models trained on unpaired sequences. We also fine-tune a left-to-right ablation, p-IgGenCDR-LTR, from the IgGenCDR-LTR checkpoint under the same data and schedule.

#### Paired fine-tuning improves recovery on every loop

To test whether joint conditioning on both chains improves reconstruction, we measured CDR recovery on the held-out paired OAS test set (∼69.8K pairs), scoring all six loops jointly under the paired framework-first input and evaluating the public p-IgGen baseline (4) in both decoding directions it was trained with (Supplementary Table S9).

Paired fine-tuning improves recovery on every loop over the unpaired base checkpoint, with the largest gains on the inter-chain-constrained H-CDR3 (+1.80%) and L-CDR3 (+3.20%), and the framework-first variant again leads its LTR ablation consistently (mirroring the unpaired pattern, Table 1). The public p-IgGen baseline behaves as expected for a bidirectional model: in each direction the late-generated loops recover well (right-to-left even edges p-IgGenCDR on H-CDR1) while the early-generated ones collapse (L-CDR3 drops to 73.5% right-to-left, versus 84.4% for p-IgGenCDR). Because p-IgGen must commit to one direction at inference, it cannot avoid this trade-off, whereas p-IgGenCDR conditions every CDR on all frameworks in a single pass and stays uniformly strong across all six loops. This pattern holds across numbering schemes (Supplementary Table S9).

#### Joint paired scoring improves fitness correlation

Because both FLAb assays (24, 25) comprise paired heavy+light sequences, p-IgGenCDR and p-IgGenCDR-LTR can score each pair as a single joint render, complementing the independent-chain scoring of the unpaired models. Both scoring modes are reported together in Table 2. Joint paired scoring significantly improves full- and framework-region correlation across both datasets indicating that both immunogenicity and expression benefit from joint consideration of heavy and light chains. Interestingly, no paired model (framework-first or LTR) showed a consistently higher correlation than its counterpart, indicating that full paired sequence scoring might be similarly correlated in framework-first than LTR input formulations. Per-scheme breakdowns are reported in Supplementary Table S10.

### C. NanoGenCDR

NanoGenCDR is initialised from the IgGenCDR checkpoint and fine-tuned on a 1:1 mixture (50% each per batch) of ∼6.5M camelid VHH sequences from INDI (16), and unpaired human OAS sequences, the latter retained to preserve the humanness and diversity learned during pre-training (relevant because nanobody design often starts from humanised camelid frameworks). The framework-first input representation applies without modification to nanobodies, which share the same single-domain architecture as an antibody heavy chain.

#### Fine-tuning closes the camelid recovery gap

To assess whether framework-first conditioning transfers to the camelid single-domain setting, we measured CDR recovery on the INDI test set (Supplementary Table S11). The base IgGenCDR checkpoint, trained only on human OAS, falls below 50% recovery on all three loops, reflecting the distributional shift between human and camelid VHH repertoires. IgLM remains similarly low even when prompted with its camelid species token, despite its multi-species OAS pretraining, and fine-tuning on INDI closes this gap entirely. NanoGenCDR then outperforms all baselines on every loop and leads the LTR ablation consistently, with the largest margin on CDR1 where full framework conditioning helps most (per-scheme results in Supplementary Table S12).

#### Generated nanobodies are plausible and developable

As for the human models, we assessed the quality of *de novo* generated nanobody CDR loops, generating 2,000 candidates prompted on test-set frameworks and comparing them against the INDI training distribution (reference set) under IMGT numbering. Their CDR length distributions overlap the reference closely across all three loops, reproducing the characteristic per-loop length preferences (Supplementary Figure S11).

The designs are also structurally plausible and developable. NanoBodyBuilder2 (26) fold-confidence scores match the reference across all three CDRs (Supplementary Figure S13), and the proportion passing the Therapeutic Nanobody Profiler developability flags (27) (hydrophobicity, charge, and CDR length) is likewise maintained relative to the reference cohort (Supplementary Figure S14).

#### The OAS mixture preserves humanness on humanised scaffolds

A key motivation for the 1:1 camelid–human OAS training mixture is to preserve humanness for design on humanised camelid frameworks, the practical substrate for therapeutic nanobodies. To isolate this effect we compare NanoGenCDR (mix) against an otherwise-identical *INDI-only* checkpoint (same initialisation, schedule and seed) on five humanised VHH scaffolds: four from Stark et al. (9) and the framework of ozoralizumab, an approved anti-TNF*α* humanised VHH therapeutic. Both models complete CDRs (*T* =1.0, IMGT, 500 sequences per framework per model on identical prompts), scored with OASis (22) restricted to the CDR-overlapping 9-mers so the shared framework does not dilute the signal (Figure 3). The OAS mixture yields significantly more human CDRs on every humanised framework (mean OASis 0.471 vs 0.410 across all scaffolds, *p<*0.001), while on non-humanised camelid INDI frameworks the two models are indistinguishable (median OASis 0.31 for both), confirming that the benefit appears precisely where it matters. This humanness gain comes at little cost to camelid sequence recovery: on held-out CDR recovery the mixed model trails the INDI-only model by only 1.35% on camelid INDI (72.02 vs 73.37) yet retains +36% on human OAS (80.31 vs 44.19), where the camelid-only model collapses (Table S14). Full per-scheme analyses are in Supplementary Section A.

**Fig. 3.**
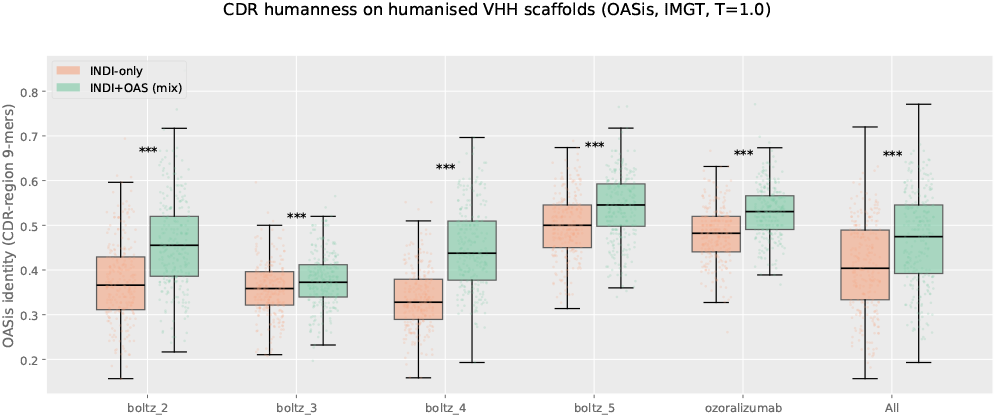
CDR-region OASis humanness (*T* =1.0, IMGT) of NanoGenCDR (mix, green) versus the INDI-only checkpoint (coral) on five humanised VHH scaffolds and pooled (All). Boxes show IQR with median; *n*=500 generated sequences per framework per model. The OAS mixture yields more human CDRs on every framework (*** *p<*0.001, two-sided Mann–Whitney *U*).

#### Likelihoods predict nanobody thermostability

Beyond CDR recovery, we tested whether NanoGenCDR’s loglikelihood preserves correlation with experimentally measured developability properties. We measured zero-shot correlation on the NbBench thermostability benchmarks (28), thermo-seq (*n*=764) and thermo-tm (*n*=567), using mean per-token log-likelihood as the fitness predictor. NanoGenCDR achieves the highest framework correlation on both datasets (*ρ*=0.331 and 0.325, averaged across schemes), substantially outperforming the base IgGenCDR checkpoint and IgLM. This confirms that domain-specific fine-tuning on camelid sequences not only improves CDR generation quality but also strengthens the model’s ability to discriminate thermostable nanobodies in a zero-shot setting, without any task-specific supervision. Full results are reported in Supplementary Section C (Table S15). Per-scheme breakdowns are given in Table S16.

### D. Alignment-driven *de novo* nanobody design

Finally, we tested whether the framework-first formulation lets a purely sequence-based GenCDR model be steered toward developable CDRs specific to an antigen through reinforcement learning. Starting from NanoGenCDR, we iteratively aligned the model against the PD-L1 target with weighted DPO (Section) over four rounds of generation, co-folding, and preference optimisation. To benchmark the resulting designs, we compared against BoltzGen (9), a structure-conditioned diffusion pipeline that generates CDR backbones on the fixed frameworks, inverse-folds each backbone into sequences with BoltzIF, and re-folds the sequences with Boltz-2 for scoring. Because our reward is built from the same *in silico* interface metrics that we then report, our aim is not to outscore BoltzGen on those metrics, but to test whether GenCDR models that explicitly optimise for them can reach the interface quality of a structure-conditioned pipeline that does not, at a lower sampling cost and with more natural designs.

#### Design conditions

Both pipelines were held to the same design problem: grafting three CDRs into the four nanobody frameworks from Stark et al. (9) under identical length constraints, liability rules (Table S2), and Boltz-2 co-folding settings (Table S3), with the same co-folding budget of 20,000 complexes each (4 rounds of 5,000 accepted designs for our model, 20,000 for BoltzGen). Structural metrics and the composite reward (Eq. 2) are computed identically for both.

#### Alignment pipeline

Each round applies the five-step loop of Section (Figure S1): the current policy generates without length constraints, sequences are screened and labelled, the 5,000 ACCEPT designs are co-folded and scored, and the model is updated with scaffold-grouped weighted DPO (Eq. S4) before seeding the next round. Full pre-screening and reward-assignment details are given in Supplementary Section A.

#### Alignment effectively removes sequence liabilities

The results in Table 3 show that after a single alignment round the model internalises the length window and the six liability rules almost completely: the ACCEPT rate rises from 23.8% before alignment to 99.8% at round 1 and 100% thereafter. Because each round trains on both the 5,000 co-folded ACCEPT designs and the flagged REJECT generations, the model learns to avoid the liability motifs directly in sequence space rather than treating them as an external screen, and after alignment it produces developable, length-compatible CDRs almost without any explicit constraint at sampling time. This matters because the Boltz-2 co-folding oracle dominates the compute cost, so the ACCEPT rate sets how many raw generations must be drawn to fill each round’s pool of ∼5,000 cofolded designs. At 23.8% acceptance NanoGenCDR needs ∼21,000 generations at round 0, whereas from round 1 onward almost every generation is usable from the aligned checkpoint. Filling the full ∼20,000-design co-folded pool therefore takes ∼36,000 generations for the aligned model versus ∼63,000 for BoltzGen, whose inverse-folding step had a ∼31.8% pass rate, a ∼43% reduction in the sampling needed to saturate the same co-folding budget.

**Table 3.**
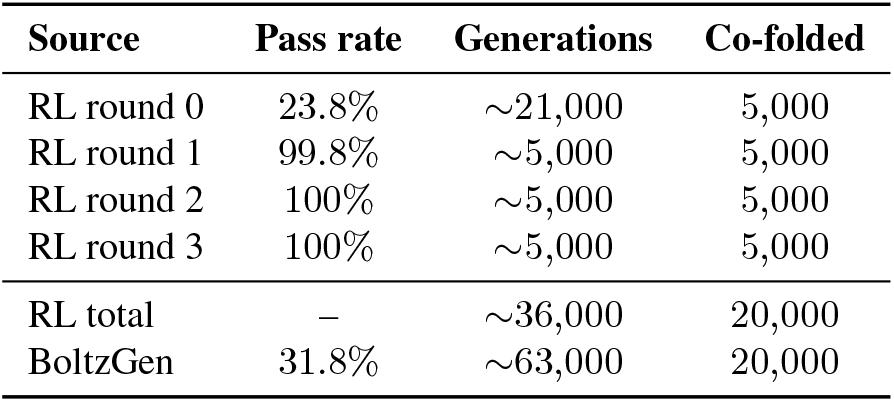
Generations required to fill the co-folding pool. Alignment satisfies the length and liability rules within a single round, so from round 1 onward almost every generated sequence is accepted and little of the generation budget is wasted. Generation counts are estimated from the per-round acceptance rate for a target of 5,000 accepted designs per round.

| Source | Pass rate | Generations | Co-folded |
| --- | --- | --- | --- |
| RL round 0 | 23.8% | ~21,000 | 5,000 |
| RL round 1 | 99.8% | ~5,000 | 5,000 |
| RL round 2 | 100% | ~5,000 | 5,000 |
| RL round 3 | 100% | ~5,000 | 5,000 |
| RL total | — | ~36,000 | 20,000 |
| BoltzGen | 31.8% | ~63,000 | 20,000 |

#### Alignment reaches BoltzGen-competitive interface metrics

Alignment steadily improves the composite reward across rounds (Figure 4, full medians and IQRs in Supplementary Table S4), from a median of 0.715 at round 0 (pre-alignment) to 0.779 at round 2, after which it plateaus (round 3, 0.775) as the KL penalty imposed by *β* increasingly resists further drift from the reference (Supplementary Figure S2). The gain is driven almost entirely by the two terms, interface confidence (ipSAE, median 0.404 → 0.530) and the CDR paratope fraction (*f*_CDR_, 0.828 → 0.943), so the aligned model designs CDRs with more confident interfaces to the target antigen and more CDR-localised contacts while continuing to satisfy the liability rules. A high *f*_CDR_ is the biologically desirable regime, since natural nanobody– antigen paratopes are concentrated on the CDRs and approved therapeutic VHHs sit at the upper end of the distribution across experimental complexes from SAbDab (29) (Supplementary Figure S12). The remaining reward terms have little room to move and stay flat. Nanobody chain foldability is already saturated from the outset (pTM_chain_ 0.973 → 0.975), and the CDR geometry terms (1 − *p*_*h*_, 1 − *p*_*s*_) barely move from their round-0 values. This stability is informative, showing that the higher interface confidence is achieved through loop-like, CDR-localised contacts rather than by introducing regular *α*-helical or *β*-strand structure into the CDRs.

**Fig. 4.**
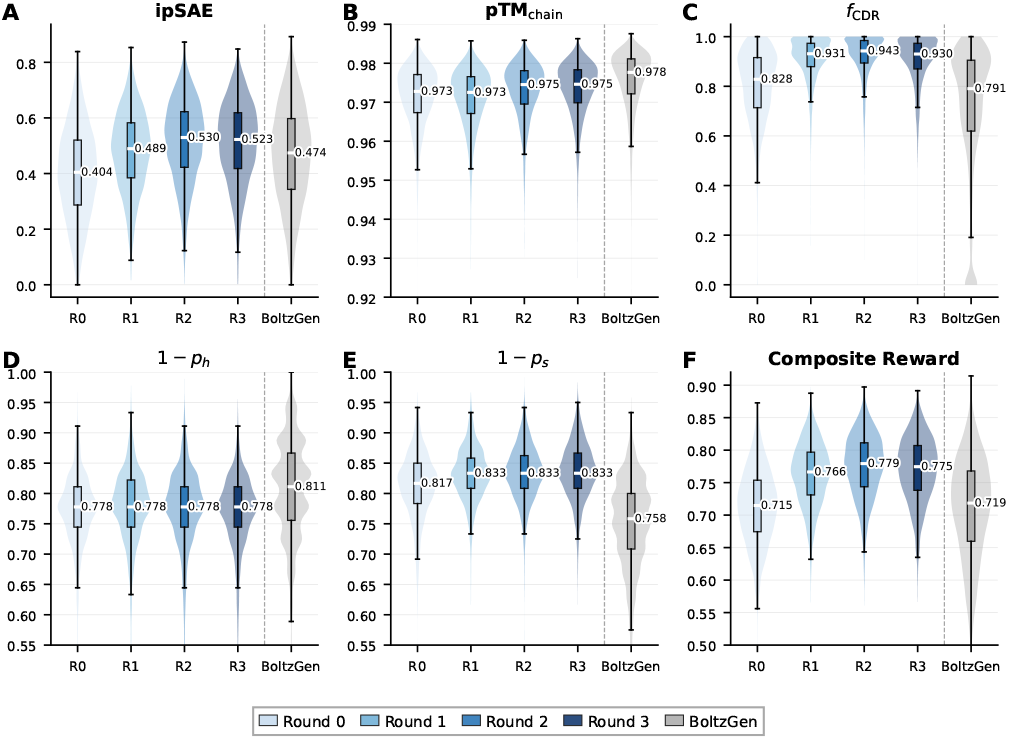
Distribution of reward components and composite reward across alignment rounds (R0–R3) and BoltzGen baseline. Violin plots show full distributions; boxes indicate IQR with median (white line). Annotated values are medians. Dashed vertical line separates alignment rounds from the BoltzGen reference.

In comparison to BoltzGen, GenCDR models never see the antigen sequence or geometry explicitly as an input, but rather learn it through the reward during alignment. At round 0 its ipSAE (0.404) sits below BoltzGen’s structure-conditioned designs (0.474), yet alignment lifts it above BoltzGen by round 2 (0.530), while localising far more of the paratope on the CDRs (*f*_CDR_ 0.943 vs 0.791). The two pipelines leave different residual secondary structure in the loops. Our aligned model more effectively suppresses *β*-strand character (1 − *p*_*s*_ 0.833 vs 0.758), whereas BoltzGen more effectively suppresses *α*-helix (1 − *p*_*h*_ 0.811 vs 0.778) and retains a marginally higher chain foldability (pTM_chain_ 0.978 vs 0.975) from its inverse-folding step.

Beyond the interface and geometry terms, the aligned designs are also markedly more natural than BoltzGen’s. Across all four rounds they score higher on both the frozen NanoGenCDR reference policy (median per-token log-likelihood −0.84 to −0.88 vs −1.02) and an independent AbLang (30) pseudo-log-likelihood (0.96 to 1.01 vs 1.01), staying closer to the natural antibody distribution despite the optimisation pressure (Supplementary Figure S2). Naturalness is not a reward term. It is preserved instead by the KL anchor to the pre-trained reference.

Taken together, these results show that a purely sequencebased generative model, aligned over four rounds with cofolding feedback, reaches *in silico* interface quality competitive with a structure-conditioned diffusion pipeline on the same scaffolds and co-folding budget, while co-adapting all three CDR loops jointly within each round, a capability architecturally inaccessible to left-to-right or single-span masked AR models.

Because a real campaign advances only its best designs, we also asked whether alignment improves the top-tail of the distribution, the cohort that would enter experimental screening. Ranking by composite reward and taking the top 5% clones, alignment improves this tail (reward 0.826 to 0.859, ipSAE 0.683 to 0.745 from round 0 to round 2) and matches BoltzGen at equal selection (Supplementary Section G), a genuine effect of alignment rather than oversampling, as recovering this cohort from the pre-alignment checkpoint would require roughly *×*170 more generation and co-folding. The tail gain is nonetheless smaller than the median gain. Because the composite reward combines bounded components without variance normalisation, and several of them (*f*_CDR_, pTM_chain_) approach their ceilings among the best designs, its spread within the top cohort is narrow, so the reward softmax yields near-uniform preference targets there. Temperature-scaling the reward softmax, or standardising rewards within each scaffold group, is a natural lever to sharpen these targets and push the tail further, which we leave to future work.

#### Alignment improves every scaffold in step

Disaggregating the composite reward by scaffold shows that reward improves across the four frameworks, from round 0 to round 2 and plateauing together, with no scaffold left behind (Supplementary Figure S3). This is a direct effect of the scaffold-grouped weighted DPO loss (Eq. S4), which balances optimisation pressure across scaffolds rather than letting a dominant one bias the update, and confirms that the framework-first formulation steers each nanobody framework toward the target in parallel.

#### Alignment adapts CDR length to the antigen

CDR length is not fixed by the reward, only bounded by the permitted window (CDR3 14–20 residues, IMGT), leaving the model free to choose where within it to place its mass. Alignment shifts that choice: from the broad round-0 distribution inherited from the reference policy, the aligned model concentrates CDR3 on the longer, higher-reward loops favoured by the PD-L1 interface by round 2, then tightens further as the reward plateaus (Supplementary Figure S4). Loop length is therefore one of the degrees of freedom the reward actively shapes, letting the model adapt CDR geometry to the target antigen rather than merely reproducing the reference length preferences. This makes aligned GenCDR models unique compared to generative pipelines relying on inverse-folding, where designed CDR length needs to be specified beforehand rather than letting the model learn the most suitable lengths to a given epitope.

## Conclusions and future work

We have presented GenCDR, a family of framework-first autoregressive antibody language models that read all framework regions as a conditioning prompt and generate the CDR loops as a variable-length response. This single change to the input ordering yields the highest CDR recovery among autoregressive antibody models across heavy, light, paired, and nanobody settings, produces generated CDRs that remain natural, diverse, and human-like, and preserves likelihoods that correlate with fitness and developability assays. Crucially, separating framework and CDR likelihoods exposes a clean prompt–response interface, making GenCDR the first autoregressive antibody model family that natively supports joint multi-CDR alignment. Exploiting this, we aligned NanoGenCDR against a PD-L1 co-folding oracle with weighted DPO and reached *in silico* interface quality competitive with BoltzGen, a structure-conditioned diffusion pipeline that requires antigen geometry at both training and inference time, while using roughly half the generation budget and producing more natural, developable designs.

Several directions remain open, particularly for the alignment setting. We aligned with a single offline preference method (weighted DPO), and other reinforcement-learning strategies, whether online, on-policy, or alternative offline objectives, may steer the policy more effectively or efficiently, and a systematic comparison would clarify which are best suited to multi-CDR design. Our alignment study also targets a single antigen (PD-L1), and extending it to a broader panel of antigens and epitopes is a natural next step. Finally, all design quality reported here is measured *in silico*. Experimental characterisation of expression, stability, and binding for aligned designs will be essential to establish whether the gains in co-folding metrics and naturalness translate into therapeutic candidates.

## Supporting information

TeX files

Supplementary Split

## Acknowledgements

We extend our gratitude to everyone in AI in Biologics Engineering and Discovery Science at AstraZeneca, with special thanks to Leon Gerard, Cedric Malherbe, and Peizhen Bai for their useful feedback and technical discussions.

## Data availability

The training corpora are derived from publicly available resources: the human OAS-derived p-IgGen dataset (https://zenodo.org/records/13880874) and the INDI nanobody databases^2^. Inference code and trained models are available at https://github.com/AstraZeneca/GenCDR.

## Supplementary Material

### Supplementary Note S1: Nanobody fine-tuning data processing

NanoGenCDR was fine-tuned on nanobody sequences from the INDI (2,697,100 sequences) and INDI2 (11,900,684 sequences) databases^3^ (16), a curated collection of camelid heavy-chain variable domains. We combine both releases and refer to the resulting corpus simply as INDI throughout the main text. Both datasets were processed through an eight-step filtering pipeline, similar to the processing performed in p-IgGen (4). Sequences were first deduplicated on full sequence identity, then filtered to remove entries exceeding 150 residues, containing unknown residues (X, *) or an odd number of cysteine residues (indicative of unpaired cysteines). Remaining sequences were annotated using ANARCI (17) under three numbering schemes (IMGT, Kabat, and Chothia) via the abnumber library. Sequences for which at least one scheme succeeded were retained. Post-annotation, sequences with IMGT framework regions shorter than expected minimums (FR1 ≥ 20, FR2 ≥ 14, FR3 ≥ 30, FR4 ≥ 9 residues) or missing the conserved cysteine residues at IMGT positions 23 and 104 were removed. Finally, the remaining sequences were clustered at 95% sequence identity using MMseqs2 linclust (coverage mode 1, coverage threshold 0.8), retaining only cluster representatives.

For INDI v1, this pipeline reduced the dataset from 2,697,100 to 1,852,588 sequences (31.3% removed). For INDI v2, the pipeline yielded 8,762,295 sequences after annotation and quality filtering, which were further reduced to 5,492,727 unique cluster representatives (37.3% redundancy removal by linclust).

The processed sequences from both datasets were combined into a unified corpus of 6,486,354 non-redundant sequences, which was then partitioned into training (94.0%), validation (1.0%), and test (5.0%) sets. To prevent data leakage, splitting was performed at the CDR cluster level rather than the individual sequence level: for each CDR (IMGT numbering), sequences were grouped by CDR length and clustered at 95% sequence identity using CD-HIT (31). A union-find algorithm was then used to merge sequences sharing any CDR cluster into super-clusters, ensuring that no two sequences in different splits share a CDR with ≥ 95% sequence identity at any CDR position. Super-clusters were assigned to splits stratified by source dataset to preserve the original dataset proportions across all partitions.

During fine-tuning this camelid corpus was combined with unpaired human OAS sequences in a 1:1 ratio within each batch (50% camelid VHH, 50% human heavy chains from OAS). The human OAS sequences are drawn from the same pre-training corpus and serve to preserve the humanness and sequence diversity learned by IgGenCDR, which is beneficial because nanobody design frequently targets humanised camelid frameworks (see Figure 3).

### Supplementary Note S2: CDR-disjoint unpaired test set

Our unpaired human models reuse the train/validation/test partitions released with p-IgGen (4). That partition reduces redundancy by clustering whole variable domains at 95% identity, which is appropriate for the whole-sequence generation it was built to benchmark but not for the per-CDR recovery we report. Because clustering operates on the entire domain, two sequences that share a CDR but differ in their frameworks can be assigned to different splits, so the partition is not disjoint at the level of any individual CDR. This inflates recovery relative to genuine generalisation, and relative to the camelid splits, which enforce per-CDR disjointness by construction (Supplementary Section S1). On the released heavy test set, 55.3% of test CDR3s appear verbatim in training and the germline-templated CDR1 and CDR2 overlap almost completely.

We therefore rebuild a CDR-disjoint test set by applying the same per-CDR criterion used for the camelid splits, after first removing test sequences that overlap the IgLM training set (517,774 heavy, 471,267 light). We gate on CDR3 alone, since CDR1 and CDR2 are largely germline-determined and would remove nearly the entire set without testing generalisation, whereas CDR3 is the discriminative, non-germline loop. A test sequence is removed if its CDR3 lies within ≥ 95% length-matched identity of any training CDR3.

This retains 202,138 of 517,774 heavy sequences (39.0%) and 31,014 of 471,267 light sequences (6.6%). The far higher lightchain removal rate reflects its smaller, germline-dominated CDR3 space, in which most test loops recur in training, and both sets remain large enough for stable recovery estimates. Relative to the raw p-IgGen partition, the disjoint set lowers heavy CDR3 recovery by ∼ 13 percentage points while leaving CDR1 and CDR2 essentially unchanged, confirming that recovery on the raw partition was inflated by CDR3 leakage. All main-text recovery is reported on the disjoint set (Table 1, with Kabat and Chothia numbering in Supplementary Tables S6–S7).

### Supplementary Note S3: Architecture and training hyperparameters

Table S1 lists the full architecture and training hyperparameters, complementing the summary in the main text. All variants share the same architecture, differing only in the fine-tuning corpus. AdamW used *β*_1_=0.9, *β*_2_=0.999 and a weight decay of 0.1, with gradient norms clipped to 1.0. The effective batch size of 1,024 was reached on 8*×*A100 (40 GB) GPUs under Distributed Data Parallel with a per-GPU batch size of 32 and 4 gradient accumulation steps.

**Table S1.**
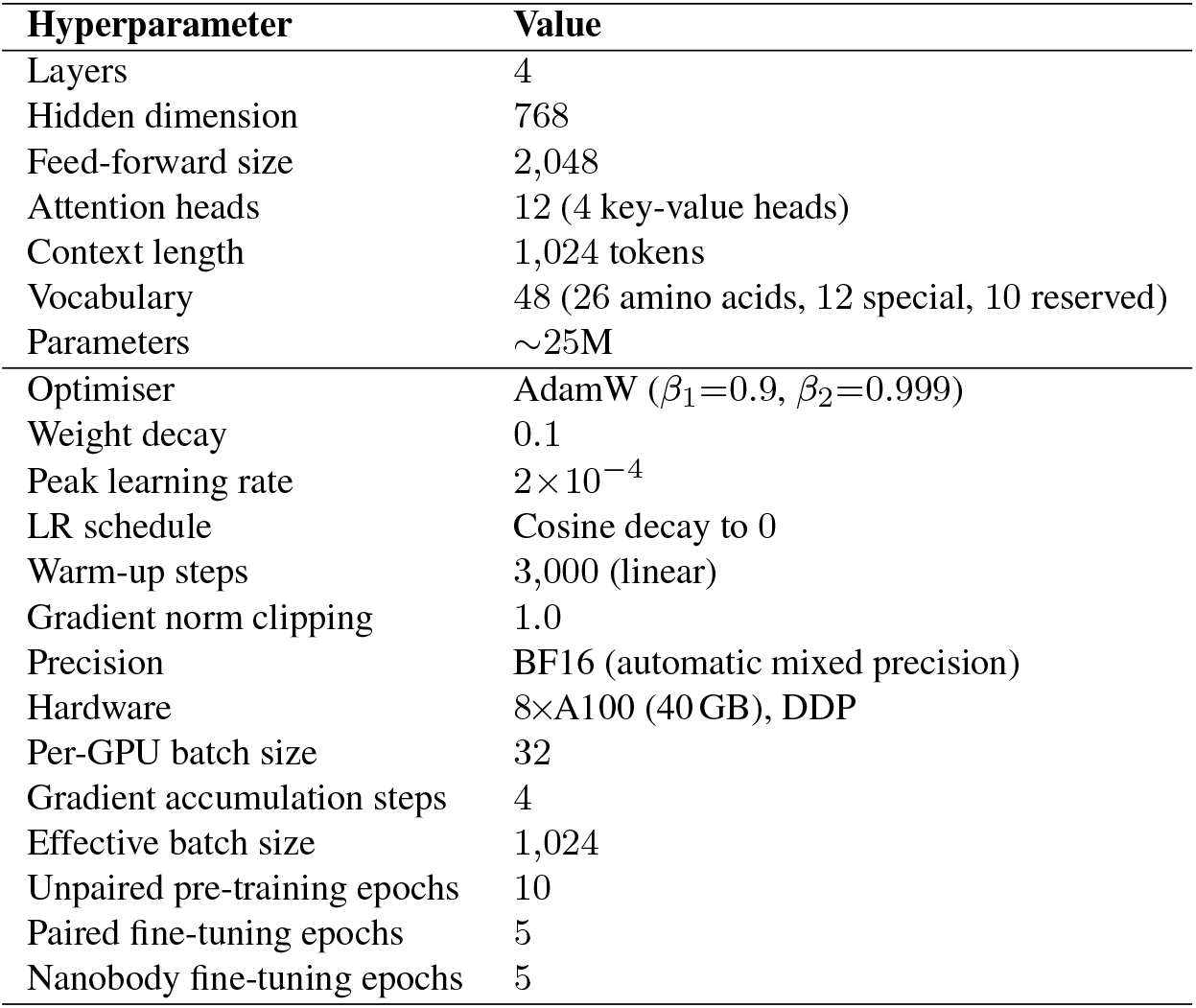
Architecture and training hyperparameters for all IgGenCDR variants. All variants share the LLaMA-style architecture above. Both p-IgGenCDR and NanoGenCDR were initialised from the IgGenCDR (unpaired) checkpoint and fine-tuned for 5 epochs on paired VH–VL sequences and the camelid–human mixture respectively.

**Fig. S1.**
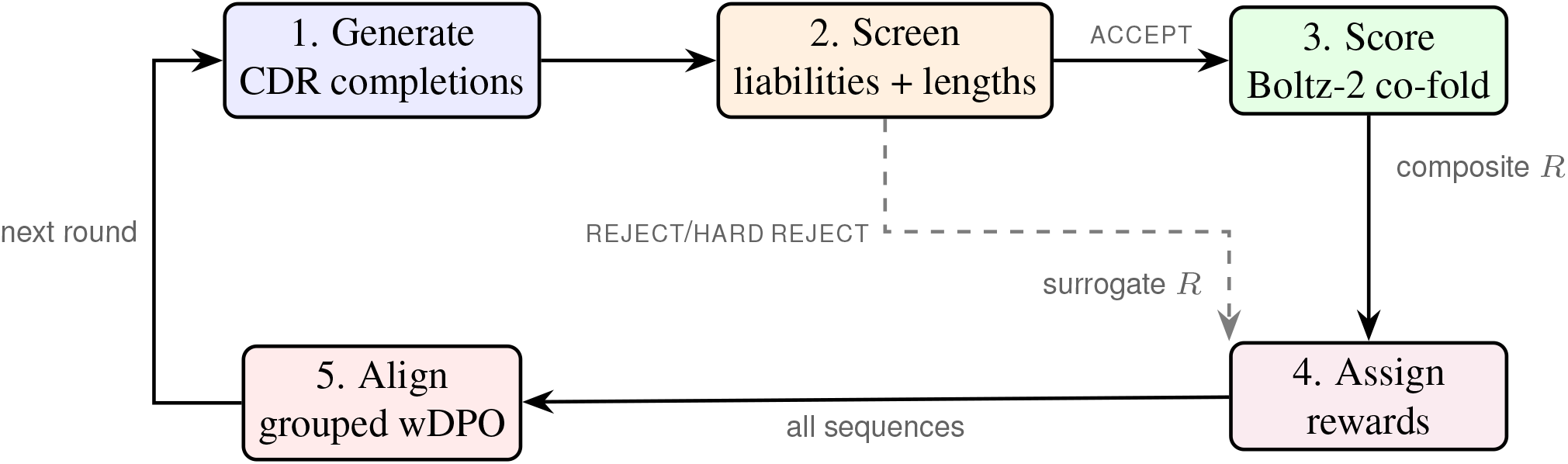
Iterative alignment loop. ACCEPT sequences are co-folded and receive the composite reward; REJECT/HARD REJECT sequences bypass co-folding and receive surrogate rewards from calibrated bands. The updated policy generates CDR sequences for the next round.

### Supplementary Note S4: Antigen-driven alignment: methods and details

Full methodological detail for the antigen-driven alignment experiment (Section D).

#### A. Pre-screening and reward calibration

Figure S1 illustrates the complete iterative alignment loop.

Before any structural evaluation, generated CDR sequences are gated to scaffold-compatible length ranges (CDR1: 6–10, CDR2: 6–10, CDR3: 14–20 residues, IMGT) and screened against six sequence-level developability liability rules, adapted from the motif set of the Blaze antibody sequence-analysis platform and consistent with established developability rules (32) (Table S2), both shared with the BoltzGen reference (Section E). Sequences passing all checks are labelled ACCEPT and proceed to Boltz-2 co-folding, the only sequences to incur that cost. Those failing one or more rules are labelled REJECT or HARD REJECT according to the number of violations.

Rejected sequences are still included in the DPO signal so the model learns that liability-prone regions yield low reward, but they bypass co-folding and instead receive surrogate rewards sampled uniformly from bands placed below the ACCEPT distribution. The REJECT band spans the two robust standard deviations immediately below the 1st percentile of the initial ACCEPT rewards, and the HARD REJECT band the two below that, where the robust standard deviation is 1.4826 times the median absolute deviation of the ACCEPT rewards. These bounds are calibrated once on the first round and fixed thereafter, keeping the surrogate rewards close enough to provide a gradient signal yet separated enough not to confound the reward landscape.

**Table S2.**
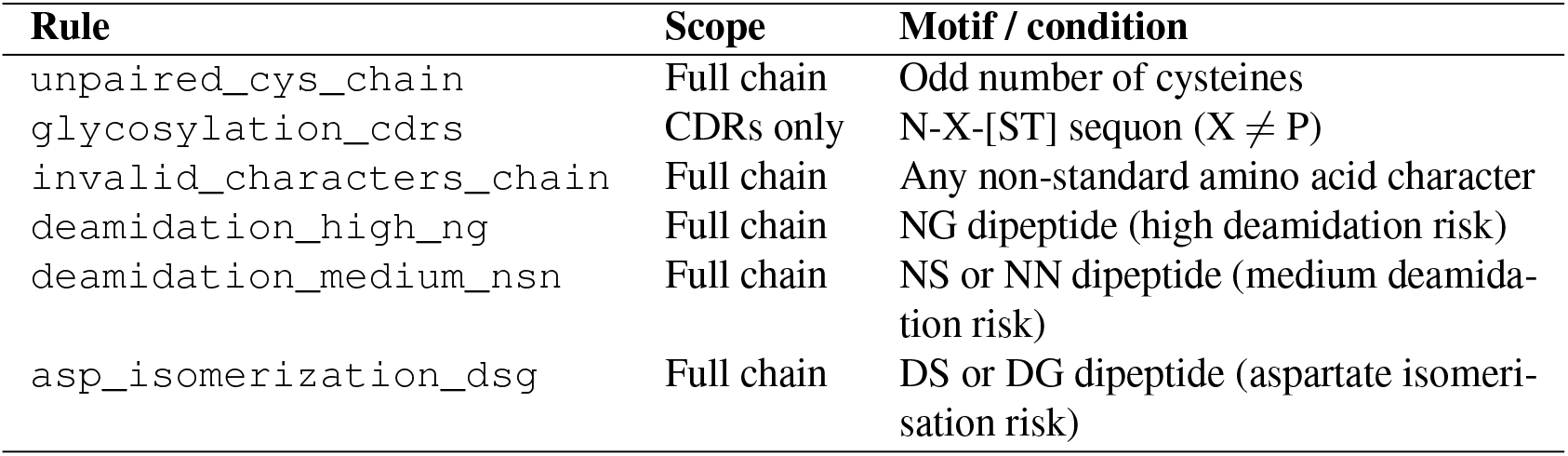
Liability rules applied during pre-screening, adapted from the Blaze platform and consistent with established developability motifs (32). A sequence failing any single rule is rejected.

#### B. Composite reward: metric definitions. Formal definitions of the five terms in Eq. 2, all evaluated on Boltz-2 (6) cofolded nanobody–antigen predictions

**Interface confidence** (ipSAE): the interaction prediction Score from Aligned Errors (20), computed from the PAE matrix restricted to inter-chain residue pairs. Higher values indicate greater confidence in the predicted binding interface.

**Chain TM-score** (pTM_chain_): predicted TM-score restricted to the nanobody chain, measuring internal fold quality independently of the interface.

**Paratope CDR fraction** (*f*_CDR_): with paratope residues defined as nanobody residues having any heavy atom within 4 Å of any antigen heavy atom and classified as CDR or framework by IMGT boundaries, *f*_CDR_ = *C/*(*C* + *F*) where *C* and *F* are CDR and framework contact counts.

**CDR geometry penalties** (*p*_*h*_, *p*_*s*_): adapted from the *α*-helical and *β*-strand distogram losses in Germinal (10), converted to post-hoc metrics on predicted structures. The *α*-helix propensity *p*_*h*_ is the fraction of CDR-internal (*i, i*+3) C*α* pairs within the helix window:

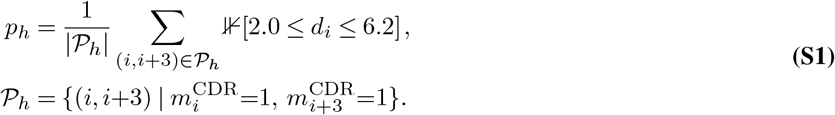

*β-strand propensity* (*p*_*s*_): the same calculation with a one-residue-expanded CDR mask (to capture CDR–framework junctions) and the strand window:

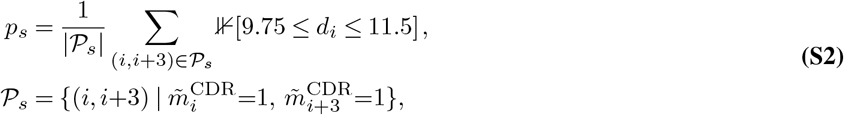

where *d*_*i*_ = ‖*x*_*i*+3_ − *x*_*i*_ ‖_2_ and 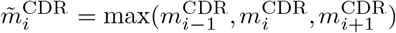. The distance windows and mask expansion follow Mille-Fragoso et al. (10).

#### C. Scaffold-grouped weighted DPO: derivation and implementation details. Single-trajectory alignment

Standard DPO (13) operates on preference pairs (*x, y*_*w*_, *y*_*l*_). When continuous scalar rewards are available, constructing preference pairs introduces information loss. The single-trajectory setting (33) instead works with triplets (*x, y, R*) and derives the optimal KL-regularised policy directly. Weighted DPO (11, 19) implements this by replacing the pairwise sigmoid with a listwise cross-entropy over softmax-normalised rewards (Eq. 1), avoiding the need for an auxiliary value function. Alternative single-trajectory formulations exist (e.g. DRO (33)) and a comparison is left to future work.

##### Completion-only log-ratio

The sequence-level log-ratio in Eq. 1 is computed as a mean over CDR completion tokens only, those after the `<HPRED>` (or `<LPRED>`) boundary through `<EOS>`, including inter-CDR `<SEP>` tokens. Framework tokens are excluded since they form the fixed prompt:

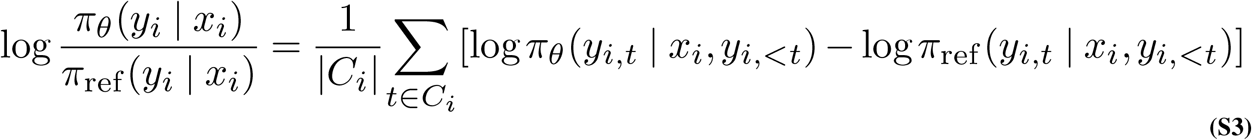

where *C*_*i*_ is the set of CDR completion token positions for sequence *i*. This decomposition follows directly from the framework-first input ordering.

##### Scaffold grouping

Each batch contains CDR completions from all four framework scaffolds. The listwise softmax of Eq. 1 is meaningful only among completions of a shared prompt: the *β*-scaled log-ratios *s*_*k*_ are comparable across responses to the same scaffold, but not across different scaffolds, whose frameworks impose different baseline likelihoods. We therefore compute the softmax targets *w*_*k*_ and logits *s*_*k*_ within each scaffold group, never across the full batch. This grouping is what makes Eq. 1 well defined, rather than a post-hoc variance-reduction step.

The per-scaffold losses are then combined by group fraction |*g*|*/B*, each group *g* collecting the |*g*| responses to one scaffold prompt:

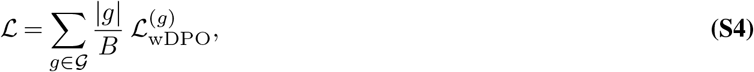

with *B* = ∑_*g* |_*g*| the batch size. Because each round generates comparable numbers of accepted completions per scaffold, group sizes are near-balanced (|*g*| ≈ *B/*|*G*|), so the group-fraction weighting reduces to a uniform average over scaffolds (|*g*| */B* ≈ 1*/*|*G*|) and no scaffold is systematically up-weighted. We adopt |*g*|*/B* as it requires no assumption of exact balance while reducing to the uniform weighting whenever balance holds. Figure S3 confirms that all four scaffolds improve in lockstep, with none dominating the update. A systematic comparison of the two aggregations under strong scaffold imbalance is left to future work.

##### Choice of *β*

We set *β* = 0.15, more conservative than the 0.01–0.1 used for single-property protein alignment (11), for three reasons: (i) the multi-CDR action space amplifies the risk of degenerate drift, (ii) the composite reward is derived from stochastic Boltz-2 predictions and is therefore noisy, (iii) we aim to preserve pre-trained CDR length and composition distributions while steering only interface-relevant properties.

##### Optimisation

Each round performs 10 epochs of wDPO on that round’s generated set with AdamW (learning rate 5*×*10^−5^, weight decay 0.01), a batch size of 512, gradient-norm clipping at 1.0, and 16-bit mixed precision, on a single A100 GPU. A stratified 20% of each round’s data, by reward, is held out for validation. The reference policy *π*_ref_ is the frozen pre-aligned NanoGenCDR checkpoint and is kept fixed across all four rounds.

#### D. Boltz-2 co-folding inference settings

All structural evaluations in this work (both alignment rounds and BoltzGen reference designs) use the same Boltz-2 (6) co-folding configuration to ensure directly comparable scores (Table S3). No template structure is provided for the antigen, only an MSA is supplied for the antigen sequence (empty for nanobody inputs).

**Table S3.**
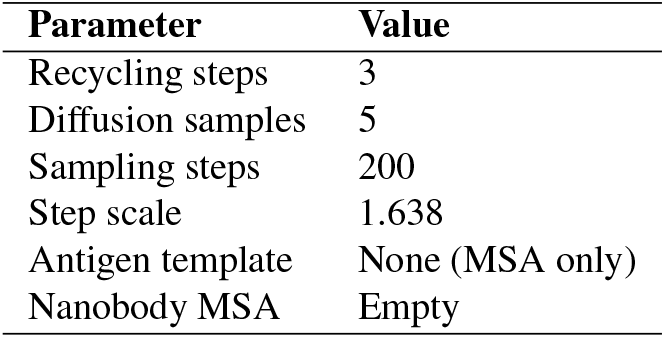
Boltz-2 co-folding inference settings used for all structural evaluations. Scores are aggregated by taking the average across the 5 diffusion samples.

#### E. BoltzGen reference design methodology

BoltzGen designs were generated against PD-L1 using the four nanobody scaffolds of Section D, following the protocol of Stark et al. (9): for each scaffold the framework structure and sequence are fixed while the three CDR regions are replaced with loops of random lengths within the shared constraints (Section A). We ran BoltzGen with `num_design = 20,000`, `diffusion_batch_size = 10`, `protocol = nanobody_anything`, `step_scale = 1.8`, and `noise_scale = 0.9`, inverse-folding each generated backbone into 4 sequences with the BoltzIF inverse-folding model (9), a soluble-protein inverse folder analogous to SolubleMPNN. Of the 80,000 candidates, 31.8% passed the shared liability and length filters from which we sampled 20,000 ACCEPT sequences for co-folding. These were co-folded with Boltz-2 under the shared settings (Table S3, ≈48 h on 8*×*A100), with structural and interface metrics computed identically to the alignment pipeline.

#### F. Alignment naturalness monitoring

Table S4 reports the full per-round distributions (median and IQR) of every reward component, the composite reward, and the reference policy log-likelihood summarised in the main text (Section D). To monitor policy drift during alignment, we track the reference policy log-likelihood (mean per-token log-probability under the frozen NanoGenCDR checkpoint) and AbLang pseudo-log-likelihood (30) across rounds (Figure S2).

The round-by-round trajectory resolves the drift underlying the reward plateau. The reference log-likelihood decreases modestly from round 0 (−0.837) to round 2 (− 0.883) as the policy diverges to push reward, then recovers in round 3 (−0.845), tracking the reward plateau (Table S4). The conservative *β*=0.15 therefore bounds drift, with the policy exploring early and being pulled back toward the reference as the KL penalty grows. For reference, BoltzGen designs score −1.020 (NanoGenCDR log-likelihood) and −1.010 (AbLang PLL), below every alignment round.

**Table S4.**
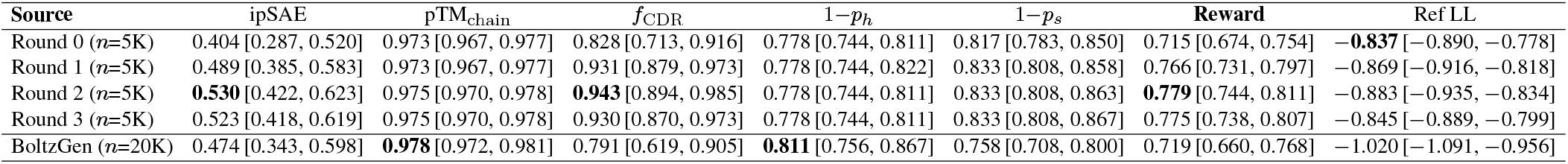
Median [IQR] of reward components, composite reward (Eq. 2), and reference policy log-likelihood across alignment rounds (ACCEPT sequences only). Both pipelines use identical Boltz-2 co-folding settings and scoring (Table S3). *f*_CDR_: paratope CDR fraction; 1−*p*_*h*_/1−*p*_*s*_: CDR geometry penalties (higher=less regular secondary structure). Ref LL: mean per-token log-probability under the frozen NanoGenCDR reference policy (not a reward component; monitors policy drift). **Bold**: best value in each column.

**Fig. S2.**
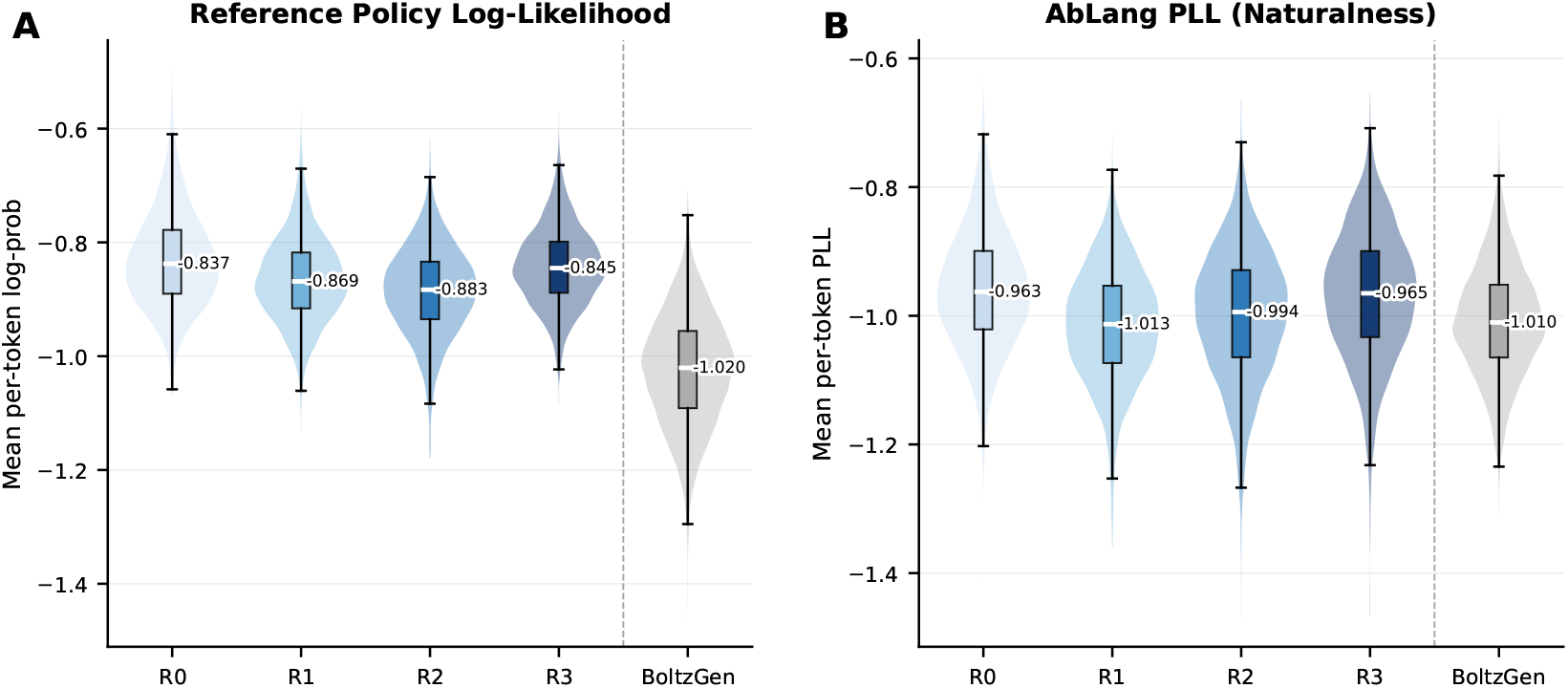
Naturalness monitoring across alignment rounds. **(A)** Reference policy (NanoGenCDR) log-likelihood of generated CDR sequences. **(B)** AbLang pseudo-log-likelihood (30), an independent antibody-specific naturalness metric. Violins show full distributions; boxes indicate IQR with median (white line). Annotated values are medians. Dashed vertical line separates alignment rounds from BoltzGen.

#### G. Tail analysis: top-5% designs

This section details the tail analysis summarised in the main text (Section D). To mirror how a campaign triages designs, we rank all ACCEPT designs of each pipeline by the composite reward, take the top 5%, and report the mean of every metric over that single selected cohort (Table S5). Ranking by a single criterion, rather than per metric, means the reported values are jointly realised by the same set of designs. The round-2 threshold defining this cohort is a composite reward of 0.847, which only 0.58% of round-0 designs meet.

**Table S5.**
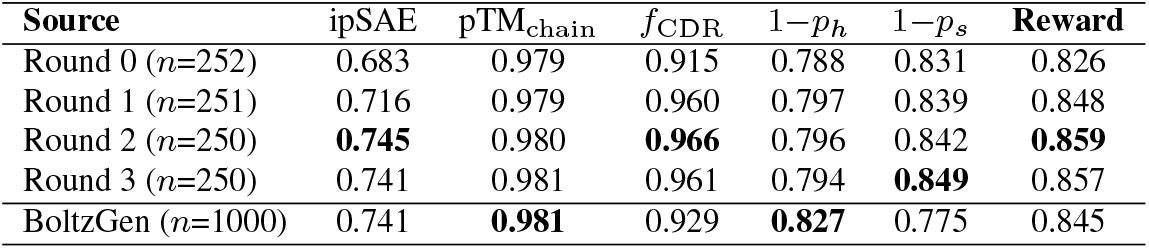
Tail analysis: designs ranked by composite reward, top 5% selected, and the mean of every metric reported over that single selected cohort (ACCEPT sequences only), complementing the median comparison of Table S4. Alignment improves the selected cohort across rounds, and at equal 5% selection the aligned round-2 cohort matches BoltzGen on ipSAE and leads on composite reward and *f*_CDR_. **Bold**: best value in each column.

#### H. Per-scaffold reward and length steering

The main-text progression (Section D, Figure 4) pools the four scaffolds. Figure S3 disaggregates the composite reward by scaffold, with all four rising from a median of ∼0.71 at round 0 to ∼0.78 by round 2 and then plateauing.

Figure S4 disaggregates the CDR3 length steering described in the main text (Section D) by scaffold, showing the broad round 0 distributions concentrating toward the longer loops at round 2 and tightening again at round 3 as the reward plateaus, consistently across all four scaffolds.

**Fig. S3.**
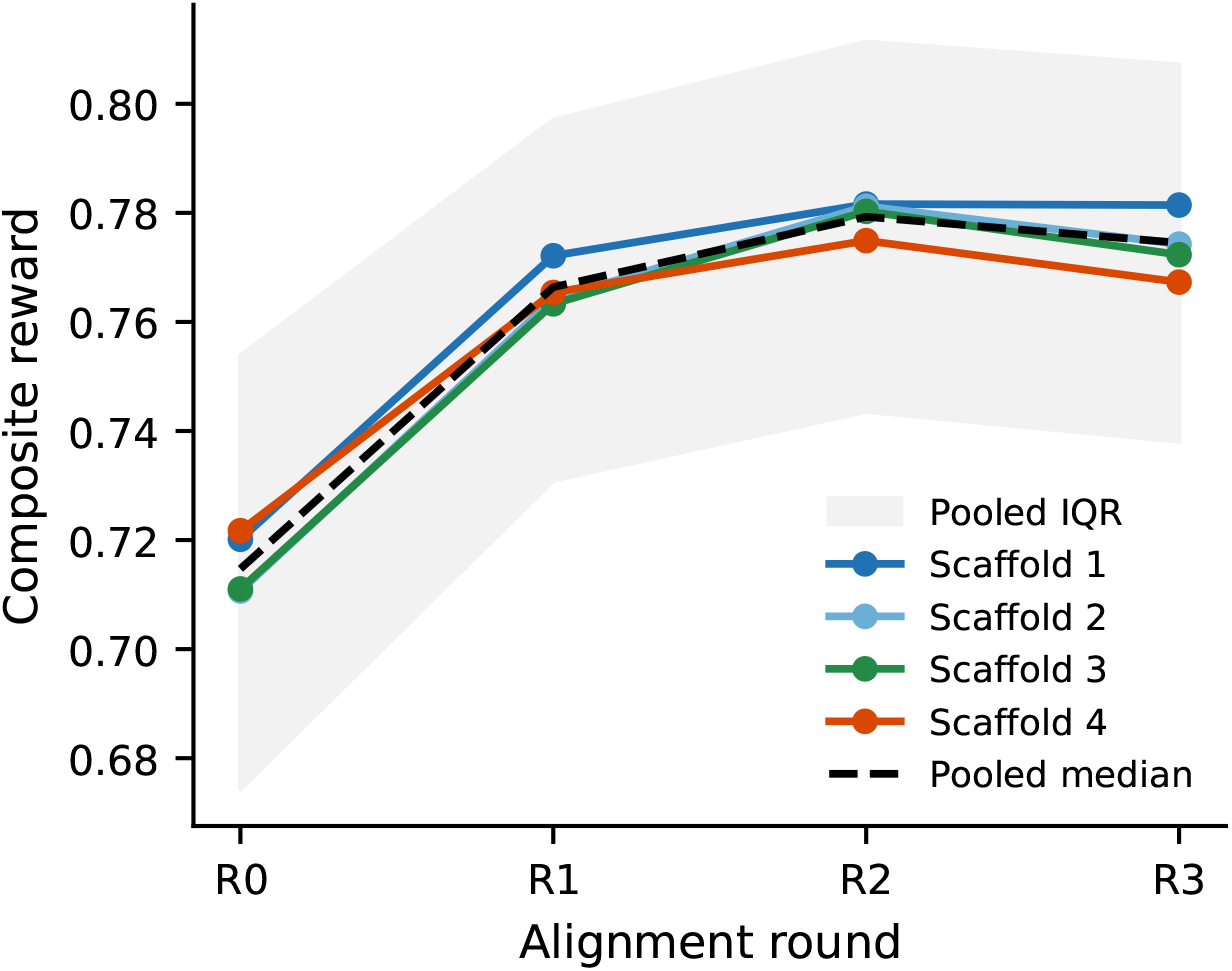
Composite reward per scaffold across alignment rounds (median over accepted designs, shading shows the pooled interquartile range). All four scaffolds improve together and plateau after round 2, mirroring the pooled trend (dashed).

**Fig. S4.**
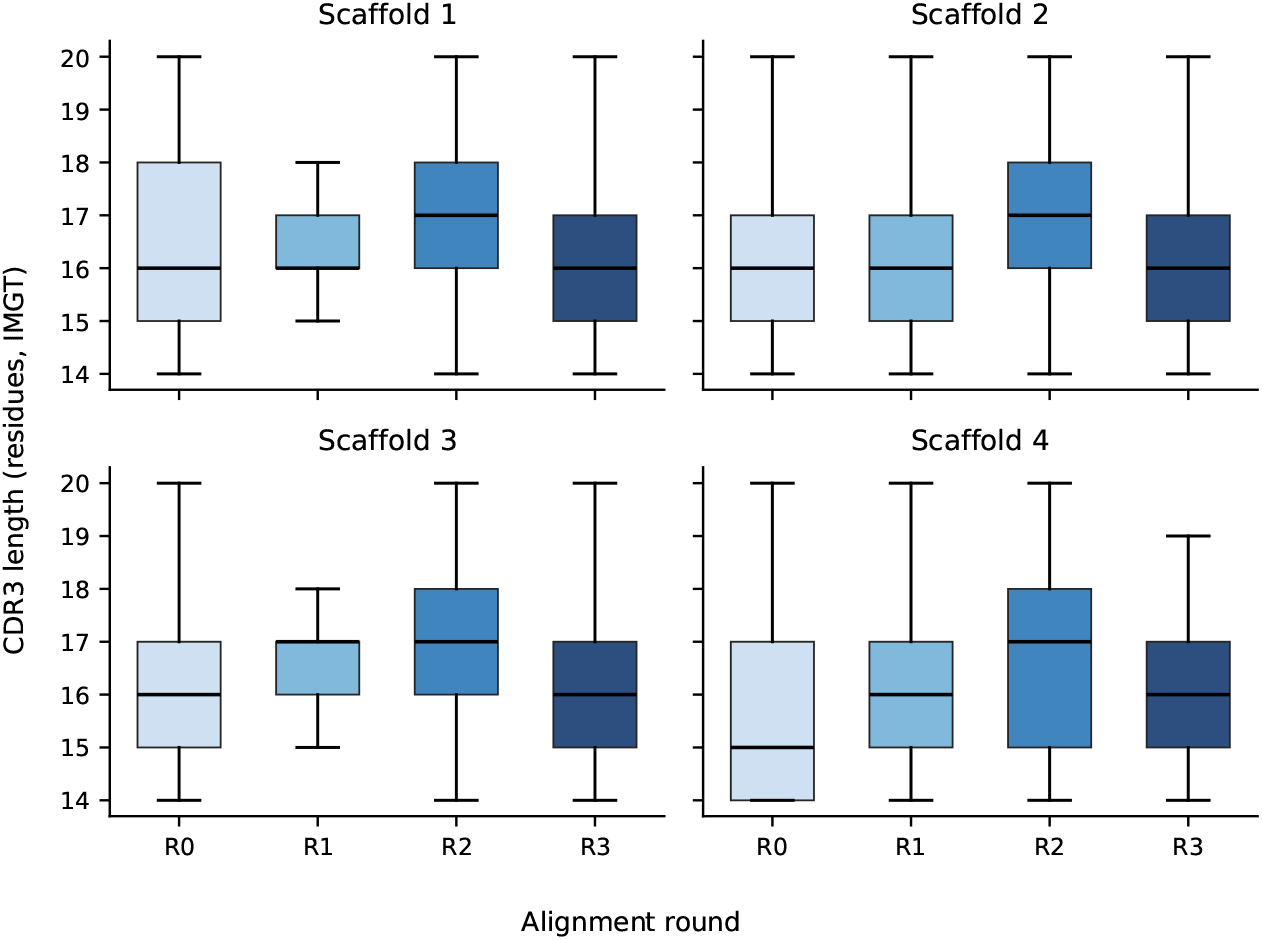
CDR3 length distribution per scaffold across alignment rounds (boxes show IQR with median; whiskers to 1.5*×*IQR). Distributions shift toward longer loops at round 2, coincident with the reward peak, then partly relax at round 3.

### Supplementary Note S5: CDR amino acid recovery under alternative numbering schemes

Tables S6 and S7 reproduce the main-text CDR-disjoint recovery comparison (Table 1) under Kabat and Chothia CDR boundaries. The model ordering is preserved across schemes.

### Supplementary Note S6: CDR recovery: detailed baseline analysis

#### Input-framing ablation

With an identical 25 M-parameter architecture and training set, switching from left-to-right to framework-first ordering improves recovery on five of the six CDR loops. CDR1 benefits the most (+2.91% heavy, +3.44% light) and CDR3 improves moderately (+0.99% heavy, +0.44% light), whereas CDR2 barely moves (−0.50% heavy, +0.10% light). CDR1 appears before FR2–FR4 in the canonical left-to-right order and thus never sees the downstream frameworks that constrain it. The framework-first layout resolves this by conditioning CDR1 on all four frameworks. CDR2, by contrast, already receives most of its relevant context from FR1 and FR2 even under left-to-right decoding.

**Table S6.**
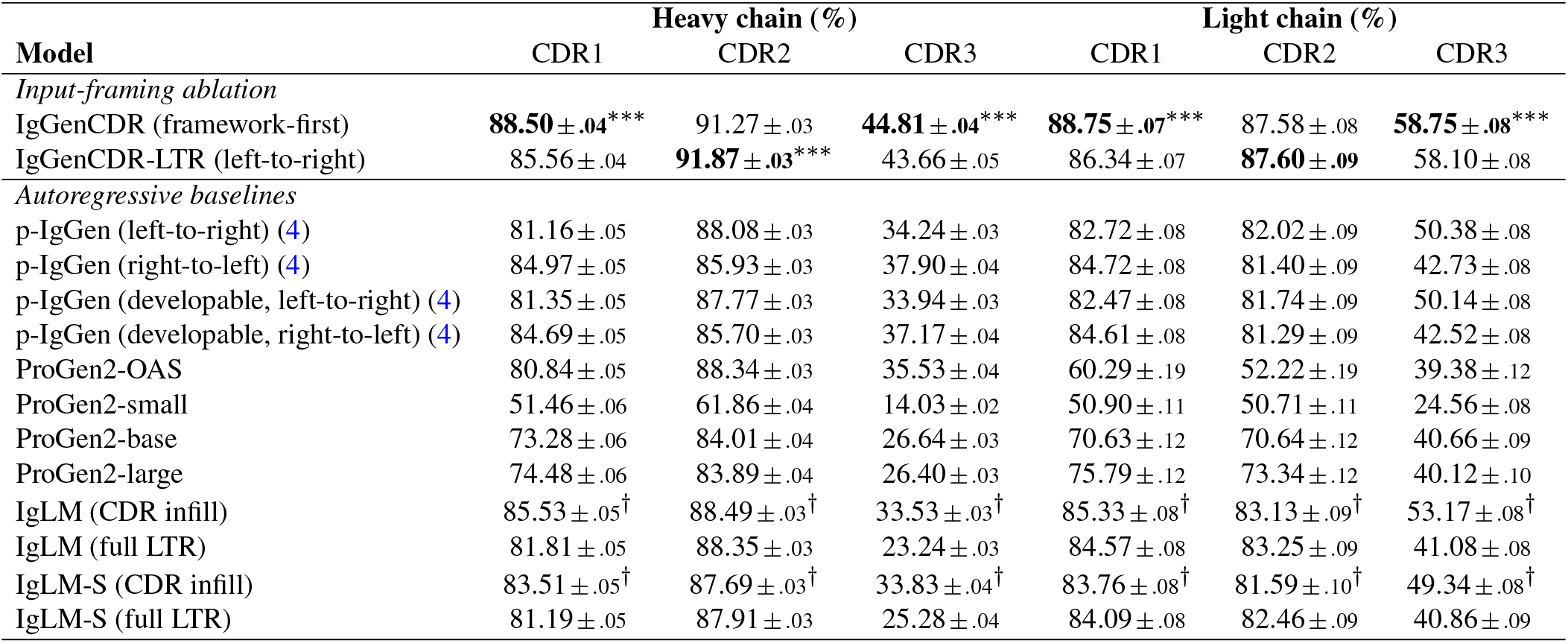
CDR amino acid recovery (NTP accuracy, %, Kabat numbering) on the CDR-disjoint unpaired test set, following the protocol of the main-text Table 1. Subscripts are one bootstrap standard error (2000 sequence-level resamples). ^*†*^Each CDR scored independently with full bidirectional context via IgLM’s single-span masked format. Stars mark the best-vs-second-best gap per column (paired Wilcoxon, Holm-corrected across loops): ^∗∗∗^*p* < 0.001, ^∗∗^*p* < 0.01, ^∗^*p* < 0.05.

**Table S7.**
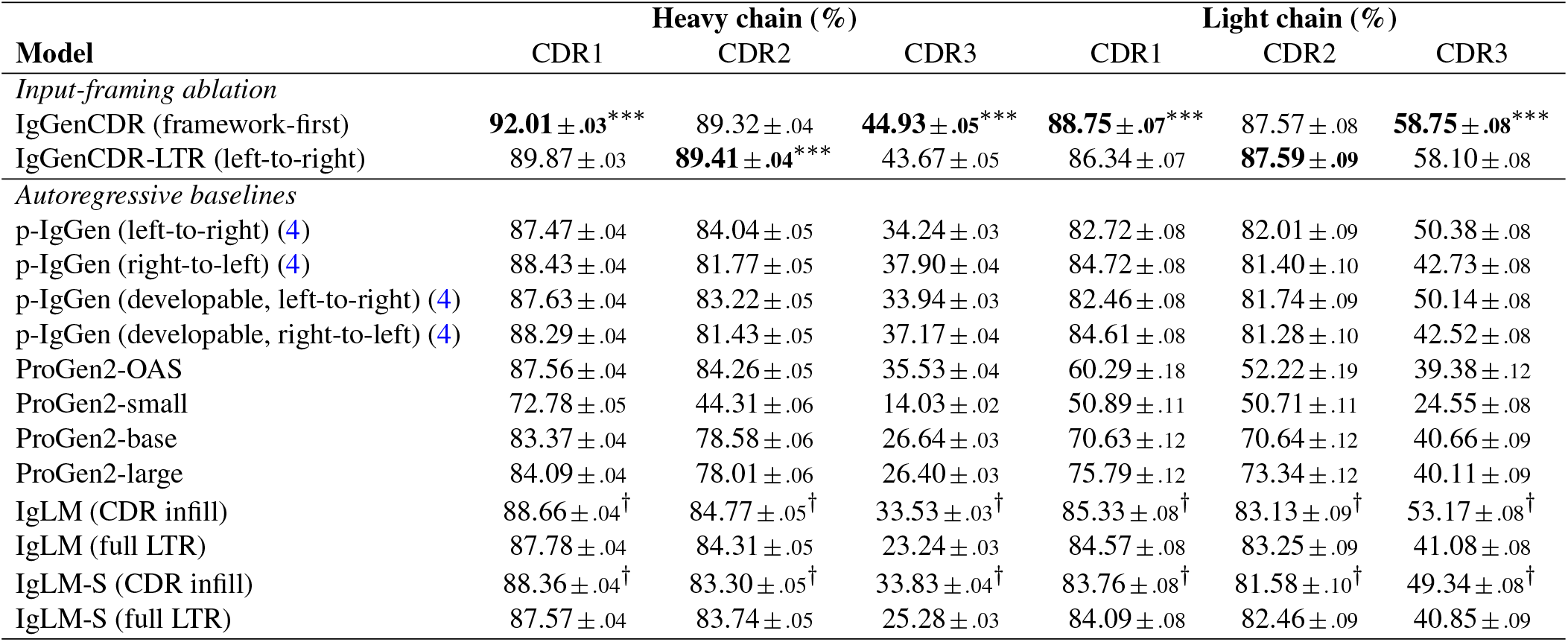
CDR amino acid recovery (NTP accuracy, %, Chothia numbering) on the CDR-disjoint unpaired test set, following the protocol of the main-text Table 1. Subscripts are one bootstrap standard error (2000 sequence-level resamples). ^*†*^Each CDR scored independently with full bidirectional context via IgLM’s single-span masked format. Stars mark the best-vs-second-best gap per column (paired Wilcoxon, Holm-corrected across loops): ^∗∗∗^*p* < 0.001, ^∗∗^*p* < 0.01, ^∗^*p* < 0.05.

#### Published baselines

p-IgGen (17 M parameters) was fine-tuned for paired heavy–light generation on ∼1.8 M paired sequences (4). ProGen2-OAS (151 M parameters) achieves competitive CDR1 and CDR2 accuracy on the heavy chain but collapses on light-chain CDR2 (43.88%), likely because its OAS training mixture is skewed towards heavy chains and lacks chain-type conditioning (3). IgLM (13 M parameters) was trained on 558 M multi-species sequences from OAS with a GPT-2 backbone and species-conditioning tokens (2).

#### Bidirectional CDR context

IgLM’s CDR-infill mode provides bidirectional context for a single loop at a time. In this mode, heavy-chain CDR3 accuracy rises from 30.65% to 39.09% and light-chain CDR3 from 41.08% to 53.17%, confirming the value of surrounding-sequence conditioning. However, IgLM’s masked-span format cannot provide simultaneous context for all three CDRs.

**Fig. S5.**
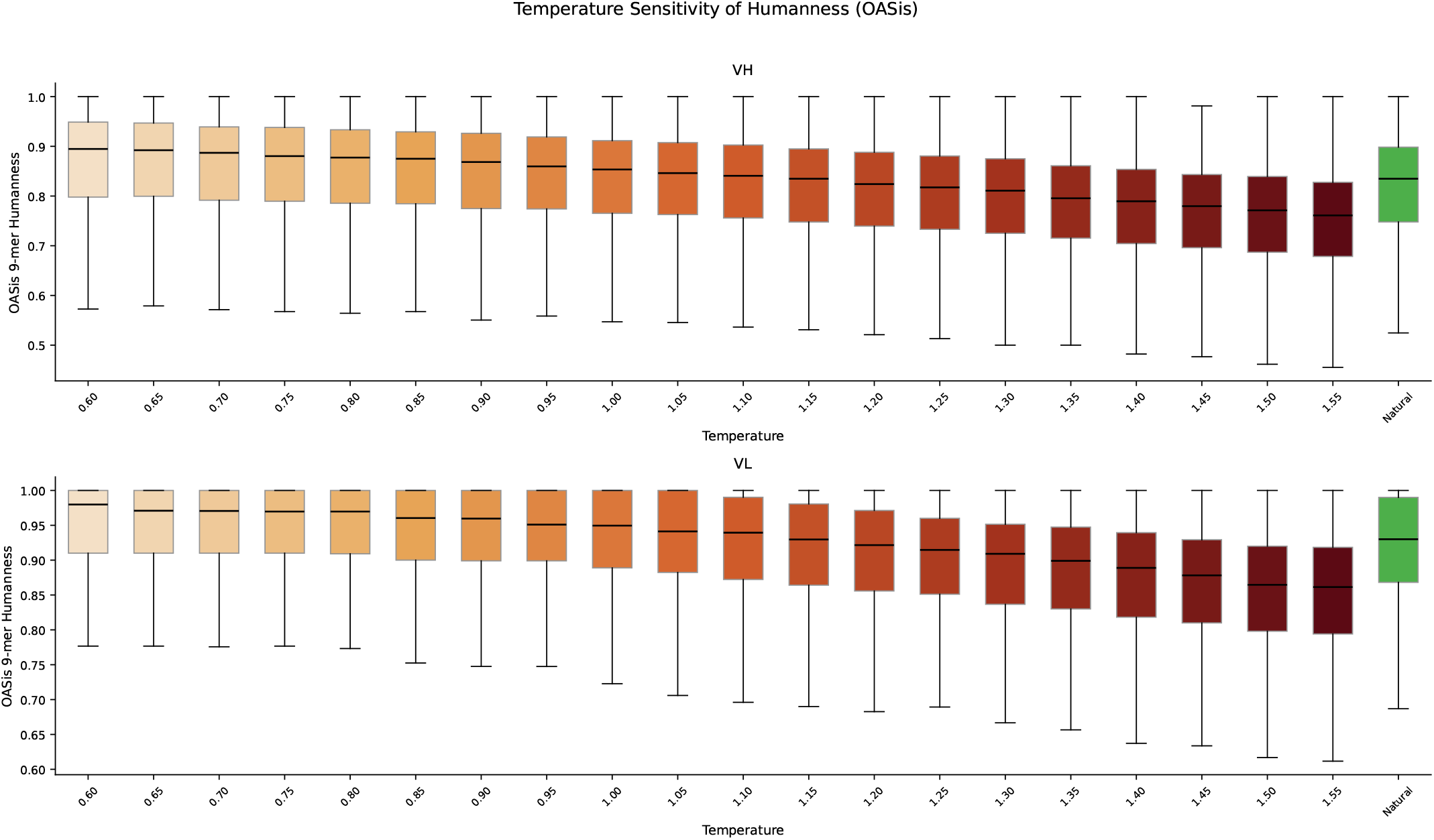
OASis 9-mer humanness across the temperature sweep (*T* = 0.60–1.55) for heavy (top) and light (bottom) chains. The green box (far right) shows the natural test-set distribution. At *T* ≤ 1.0 the generated distribution exceeds the natural reference; humanness degrades smoothly at higher temperatures.

### Supplementary Note S7: Generated CDR quality: full temperature sweep

This section collects the full temperature-sweep quality figures summarised in the IgGenCDR generation-quality results (main text). Across the sweep (*T* = 0.60–1.55), OASis 9-mer humanness (Figure S5) and ESM-2 pseudo-log-likelihood (Figure S7) degrade smoothly while sequence novelty (Figure S8) rises monotonically, and at *T* = 1.0 the generated ESM-2 distribution overlaps closely with the natural test set (Figure S6). Together these confirm that *T* ∈ [0.90, 1.15] offers the best trade-off between naturalness and diversity.

### Supplementary Note S8: Zero-shot fitness correlation

We evaluate zero-shot fitness correlation on two assays from the FLAb benchmark collection (24, 25). Marks et al. (24) (*n*=217) measures the percentage of anti-drug antibody (%ADA) response of clinical-stage therapeutics to humanisation mutations, providing a proxy for immunogenicity, lower %ADA is preferable. Koenig et al. (25) (*n*=4,276) reports the expression ratio (ER) of single-point mutants of the G6.31 anti-VEGF antibody, where higher expression is preferable. Both datasets contain paired heavy and light chain sequences with a scalar fitness label.

Each sequence is scored by its mean per-token log-likelihood under the model:

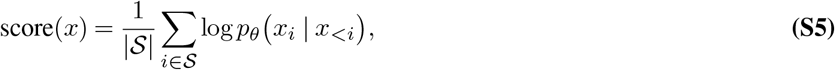

where *S* indexes either all variable-domain tokens (full-sequence scoring), the framework tokens only, or the CDR tokens only, giving the Full, Framework, and CDR scores reported throughout. For the framework-first models the CDR score uses the framework-conditioned factorization 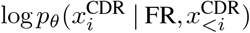, the quantity directly optimised during training. Log-likelihoods are computed separately for the heavy and light chains, and the pair score is their arithmetic mean. CDR boundaries follow the indicated numbering scheme, and for each dataset the sign is adjusted so that a positive Spearman *ρ* indicates correct model behaviour: for Marks et al. (24) immunogenicity, lower is better and for Koenig et al. (25) expression, higher is better. Reported values are averaged across IMGT, Kabat, and Chothia to marginalise over the arbitrary choice of scheme, with per-scheme results in Table S8.

IgGenCDR uses the framework-first factorisation: framework tokens condition CDR log-likelihoods, matching the training objective. IgGenCDR-LTR is the left-to-right ablation trained on identical data. Its full-sequence log-likelihood is schemeindependent (same token order regardless of numbering), but framework and CDR decompositions differ across schemes as FR/CDR boundaries change. IgLM (2) is a GPT-2 language model conditioned on chain type and species. It is the most architecturally similar infilling baseline and is evaluated in its standard full-sequence left-to-right mode (HEAVY/LIGHT + HUMAN prefix).

**Fig. S6.**
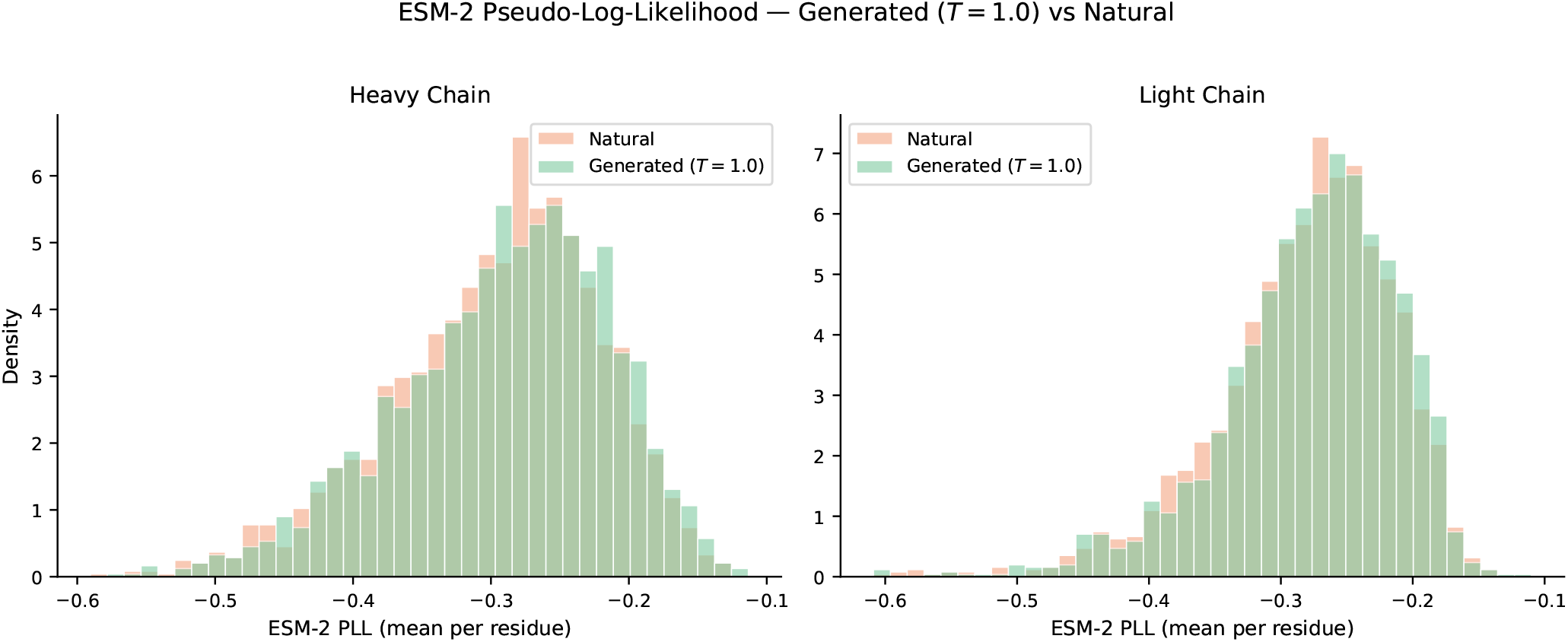
ESM-2 pseudo-log-likelihood histograms for IgGenCDR-generated sequences at *T* = 1.0 (green) overlaid on natural OAS test-set sequences (coral). Heavy chain (left) and light chain (right). The two distributions overlap extensively, indicating comparable naturalness.

**Fig. S7.**
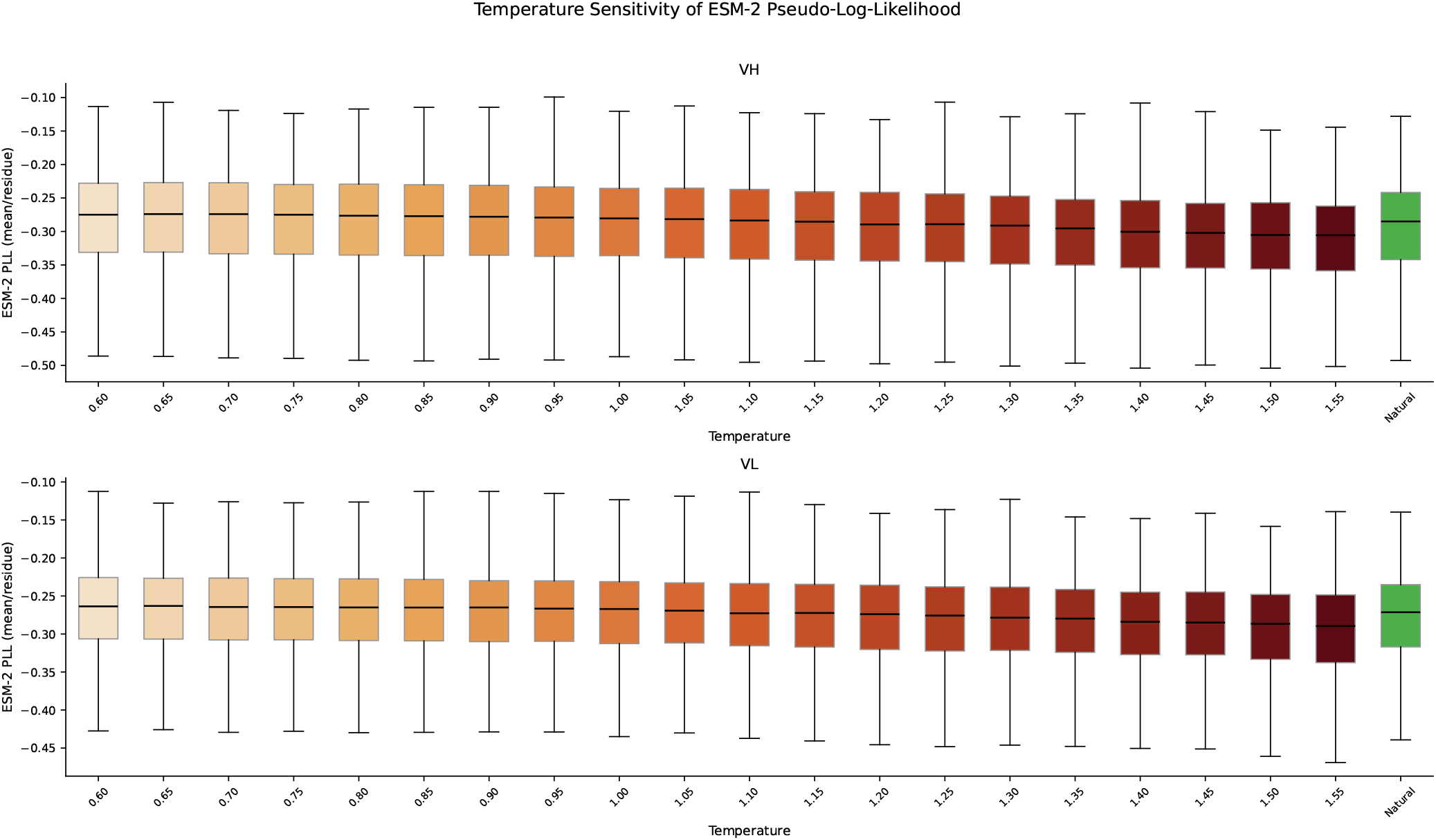
ESM-2 pseudo-log-likelihood (PLL, mean per residue) across the temperature sweep for heavy (top) and light (bottom) chains. Natural test-set reference in green. PLL is stable up to *T* ≈ 1.0 and declines smoothly at higher temperatures.

**Fig. S8.**
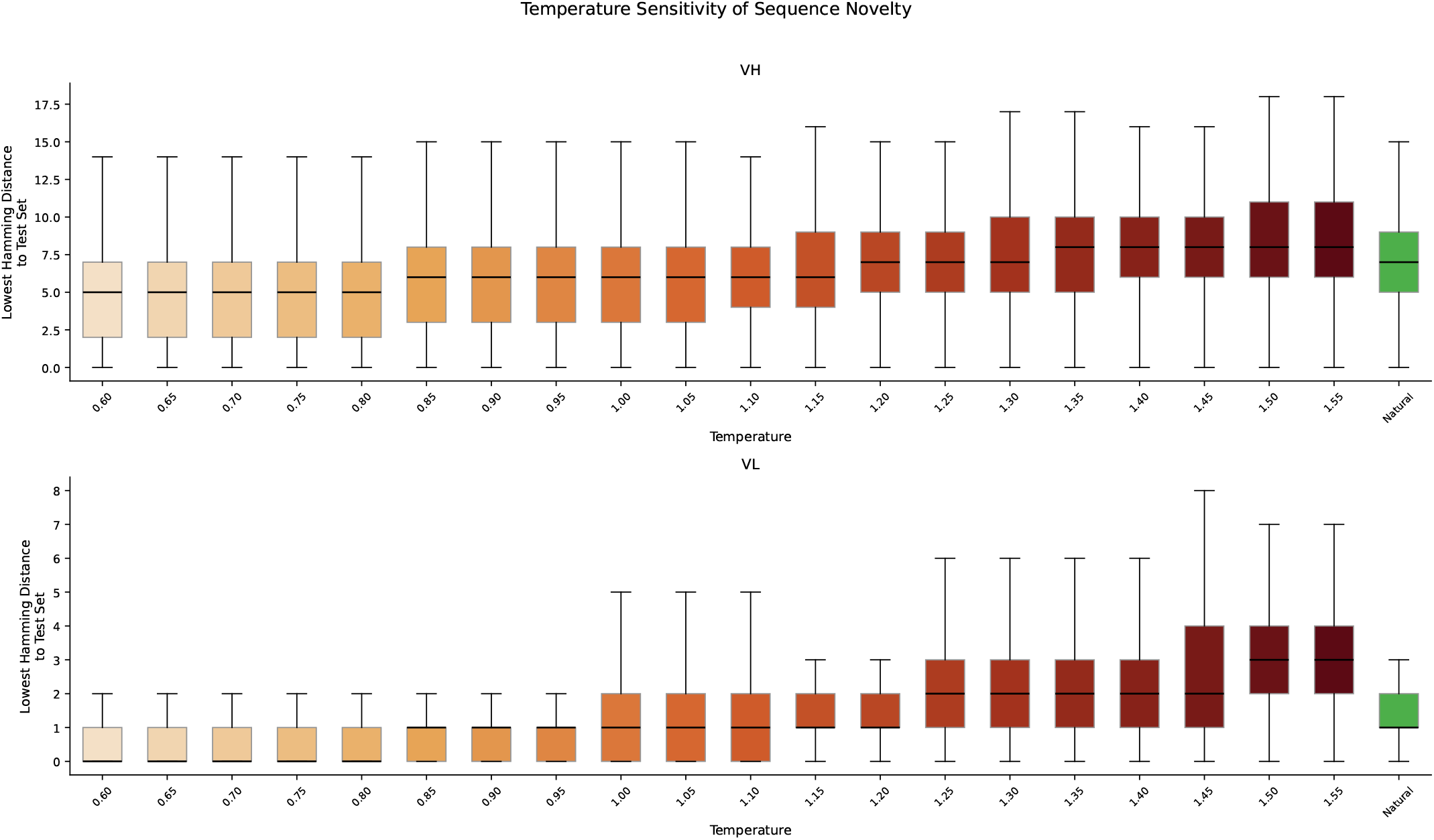
Sequence novelty across the temperature sweep, measured as the minimum Hamming distance to the nearest training-set sequence. Heavy chain (top) and light chain (bottom). Natural test-set reference in green. Novelty increases smoothly with temperature.

The scheme-averaged correlations are reported in the main text (Table 2), and the per-scheme breakdown is given in Table S8.

**Table S8.**
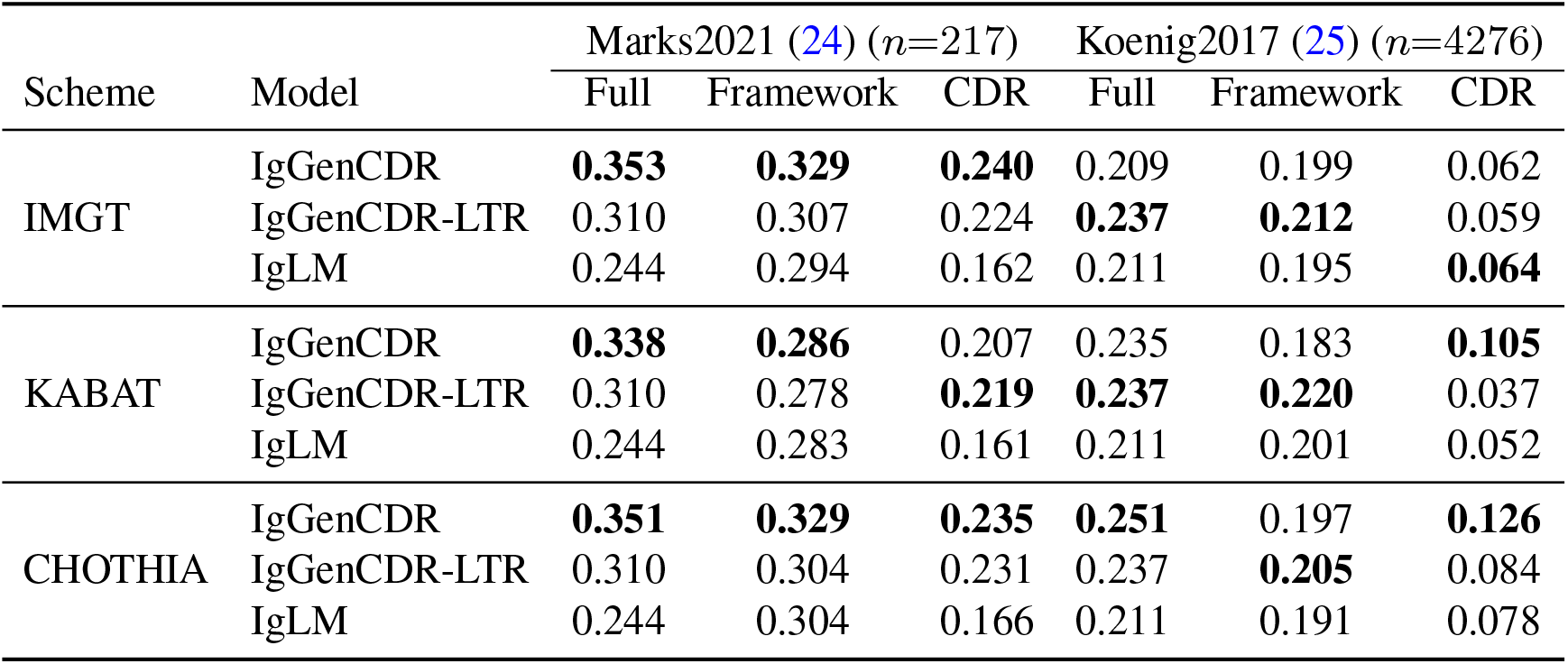
Per-scheme log-likelihood–fitness correlation (Spearman *ρ*) for paired heavy+light chain sequences. Region boundaries (FR/CDR) are defined by each numbering scheme. **Bold**: highest *ρ* in each column within each numbering scheme.

### Supplementary Note S9: Paired IgGenCDR (p-IgGenCDR)

This section provides additional detail and per-scheme breakdowns for the paired model introduced in Section B. p-IgGenCDR is fine-tuned from the IgGenCDR checkpoint on ~1.8M natively paired VH–VL sequences from paired OAS only. Both chains are rendered as a single paired sequence in the framework-first layout (Figure S9), presenting all framework regions of both chains first as a joint conditioning prompt, followed by the two CDR blocks, each introduced by its own generationboundary token (`<HPRED>`, `<LPRED>`). This lets the model learn inter-chain co-evolutionary dependencies inaccessible to models trained on unpaired sequences. We also fine-tune a left-to-right ablation, p-IgGenCDR-LTR, from the IgGenCDR-LTR checkpoint under the same data and schedule, and include the unpaired IgGenCDR (base) checkpoint as an ablation. The full per-scheme recovery breakdown for all models follows in Table S9.

**Fig. S9.**
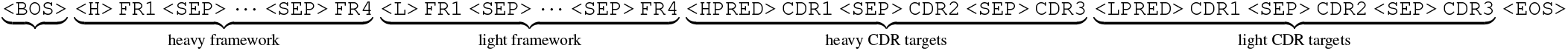
Paired framework-first input representation (heavy-first shown). Both chains’ framework regions form a single joint conditioning prompt, followed by the heavy and light CDR blocks, each preceded by its own generation-boundary token (`<HPRED>`, `<LPRED>`). The chain order (heavy-first or light-first) is sampled uniformly per training example. This extends the single-chain layout of the main text to two chains without architectural change.

#### A. CDR recovery: per-scheme breakdown

Table S9 reports CDR recovery for all models under each numbering scheme (IMGT, Kabat, Chothia).

**Table S9.**
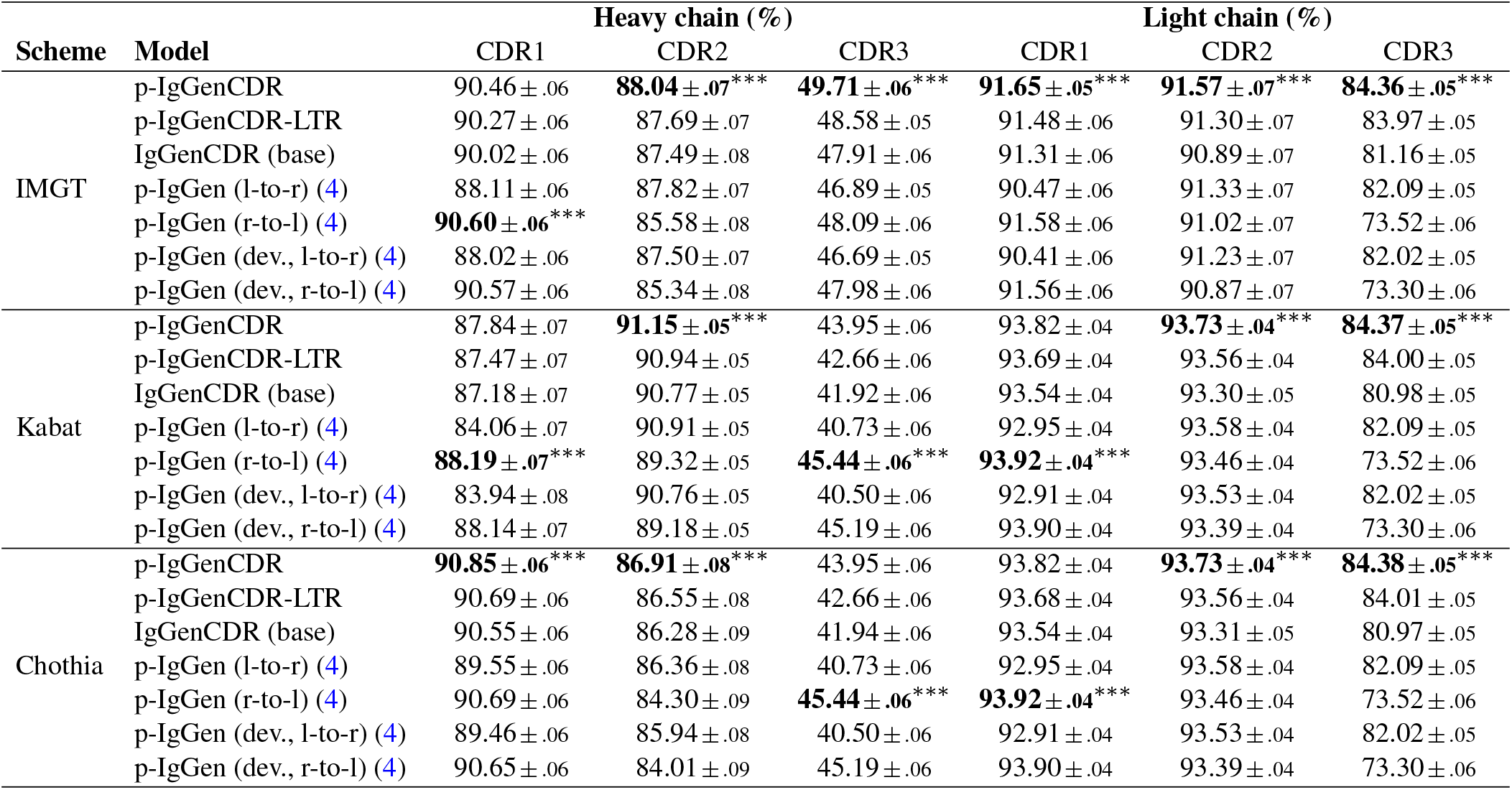
Per-scheme CDR amino acid recovery (NTP accuracy, %) on the paired OAS test set (∼69.8K pairs), for all models under each numbering scheme. p-IgGen and p-IgGen (developable) are the public baselines (4), evaluated on the same pairs in both decoding directions (l-to-r: `1+VH+VL+2`, r-to-l: `2+rev+1`). Subscripts are one bootstrap standard error (2000 sequence-level resamples). Stars mark the best-vs-second-best gap per column (paired Wilcoxon, Holm-corrected across the six loops) and sit on the column-best (bold) cell. Framework-first also exceeds its LTR ablation on all six loops under every scheme (paired Wilcoxon, Holm-corrected). ^∗∗∗^*p* < 0.001, ^∗∗^*p* < 0.01, ^∗^*p* < 0.05.

### B. Generated CDR length distributions

We assess *de novo* generation quality for p-IgGenCDR by prompting the model with 2,000 held-out paired framework scaffolds from the paired OAS test set and generating all six CDR loops at *T* = 1.0 (top-*p* = 0.95, IMGT numbering). Figure S10 compares the resulting CDR length distributions against the natural paired test-set sequences. As for the unpaired model (main-text Figure 2), the generated and natural distributions overlap closely across all six loops of both chains, confirming that paired fine-tuning preserves the length-distribution fidelity of the base model.

### C. Zero-shot fitness correlation on paired FLAb

The scheme-averaged paired FLAb correlations are reported in the main text (Table 2), where they appear alongside the independent-chain (pair-mean) scores of the unpaired models. For p-IgGenCDR and p-IgGenCDR-LTR, the H+L pair is rendered as a single paired sequence (`render_paired`) and per-region log-likelihoods are computed jointly: *Full* sums over all amino-acid tokens of both chains, *Framework* over the H-and L-framework blocks, and *CDR* over the post-`<HPRED>` and post-`<LPRED>` blocks. The unpaired IgGenCDR baseline cannot be evaluated under this rendering because it has not been trained on the paired layout, so its independent-chain pair-mean values (from Table S8) are reproduced here for reference. Sign convention follows Section S8: positive Spearman *ρ* indicates correct model behaviour. Table S10 gives the per-scheme breakdown.

**Table S10.**
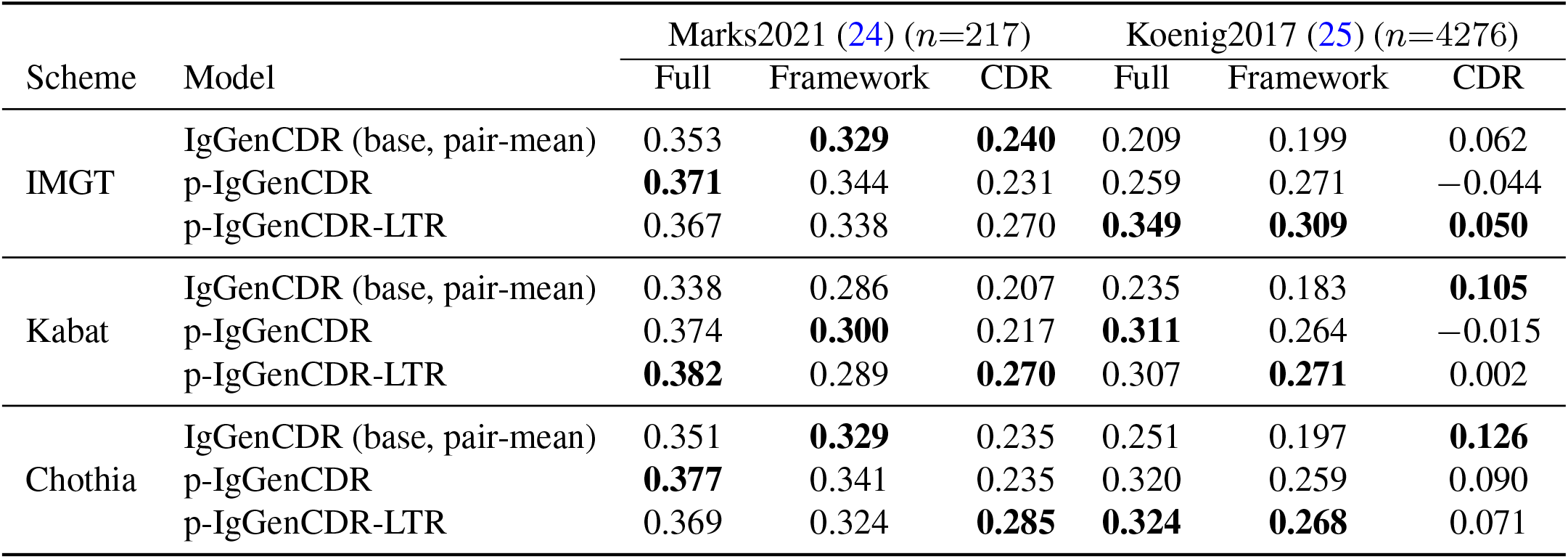
Per-scheme log-likelihood–fitness correlation (Spearman *ρ*) on FLAb. p-IgGenCDR and p-IgGenCDR-LTR use joint paired scoring, whereas the IgGenCDR (base) row reproduces the per-scheme pair-mean values from Table S8. Sign convention: positive *ρ* indicates correct model behaviour. **Bold**: highest *ρ* in each column within each numbering scheme.

**Fig. S10.**
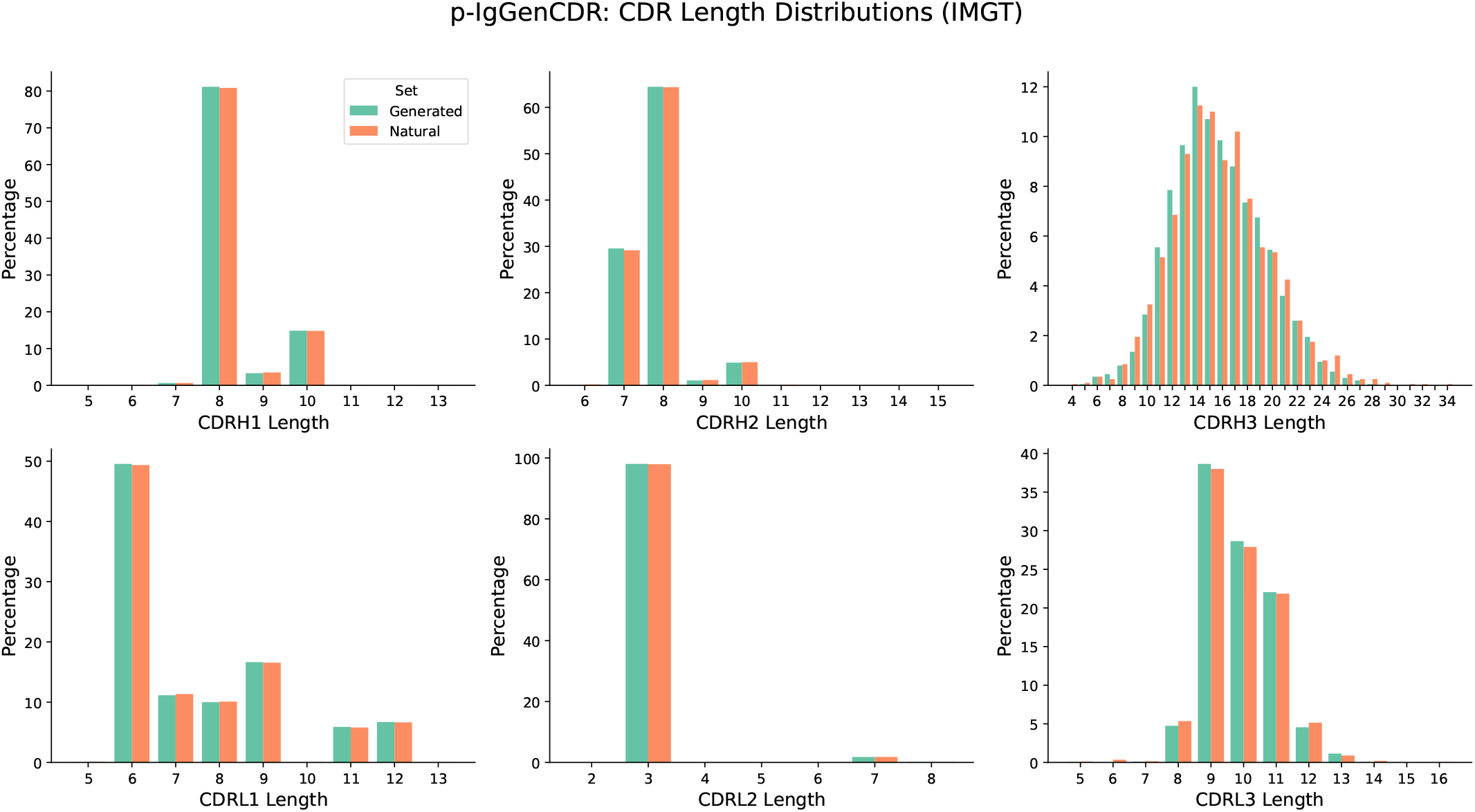
CDR length distributions (IMGT) for p-IgGenCDR-generated sequences at *T* = 1.0 (green) versus natural paired OAS testset sequences (coral). Heavy chain (top) and light chain (bottom); CDR1, CDR2, CDR3 from left to right. Bars show the per-length proportion within each set.

### Supplementary Note S10: NanoGenCDR additional results

Table S11 reports the scheme-averaged CDR recovery on the INDI test set summarised in the main text (Section C), and Table S12 gives the per-scheme breakdown.

#### A. Effect of the OAS mixture on CDR humanness

This section expands the humanness comparison summarised in the main text (Section C, Figure 3, Table S13). NanoGenCDR is fine-tuned on a 1:1 mixture of camelid (INDI) and human OAS sequences (Section S1). To test whether the human OAS fraction improves the humanness of generated CDRs, we compare it against an otherwise-identical checkpoint trained on camelid sequences only (*INDI-only*), sharing the same initialisation, schedule and seed and differing solely in the OAS mixture. Both models complete CDRs (*T* =1.0, IMGT) on five humanised VHH scaffolds: four humanised camelid frameworks from Stark et al. (9) and the framework of ozoralizumab, an approved anti-TNF*α* humanised VHH therapeutic. These scaffolds are hybrids that combine a human VH3-derived framework with retained camelid hallmark and loop-supporting residues, and are the practical substrate for therapeutic nanobody design. The ozoralizumab framework was taken from its sequence (deposited in SAbDab (29)) and is representative of the humanised synthetic VHH frameworks now used to build therapeutic libraries (34), and we segment it into FR1–FR4 with ANARCI (17) under IMGT. We generate 500 sequences per framework per model on identical prompts, and score humanness with OASis (22), restricted to the 9-mers overlapping the generated CDRs so that the shared framework does not dilute the signal. Per-framework means are given in Table S13.

**Fig. S11.**
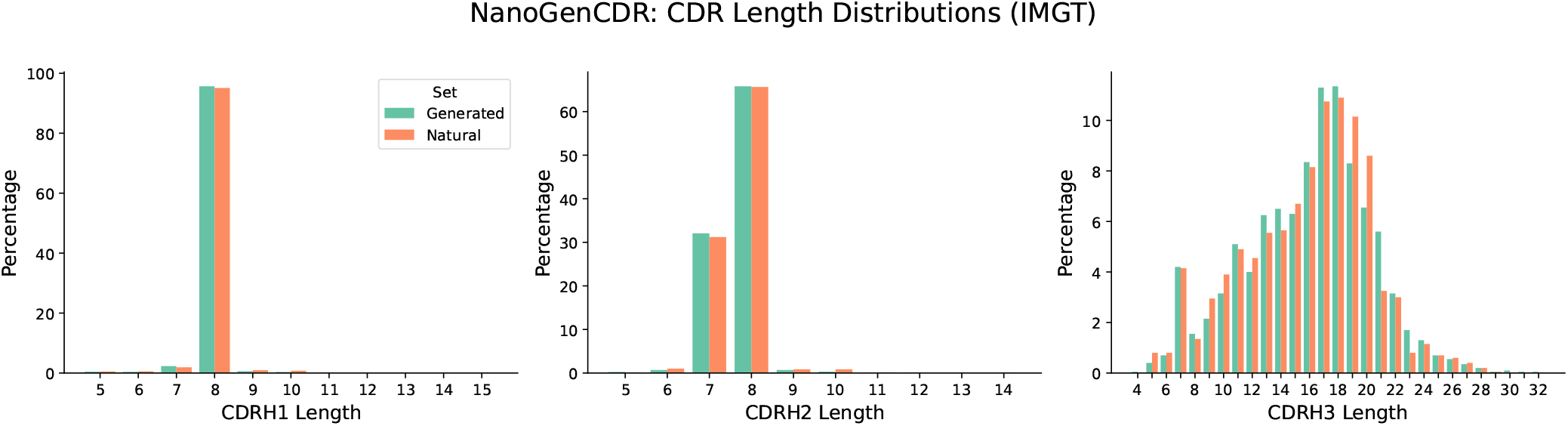
CDR length distributions (IMGT) for NanoGenCDR generations from 2,000 INDI framework prompts at *T* = 1.0 (green) versus natural INDI test-set heavy-chain sequences (coral). CDR1, CDR2, CDR3 from left to right; bars show the per-length proportion within each set.

**Fig. S12.**
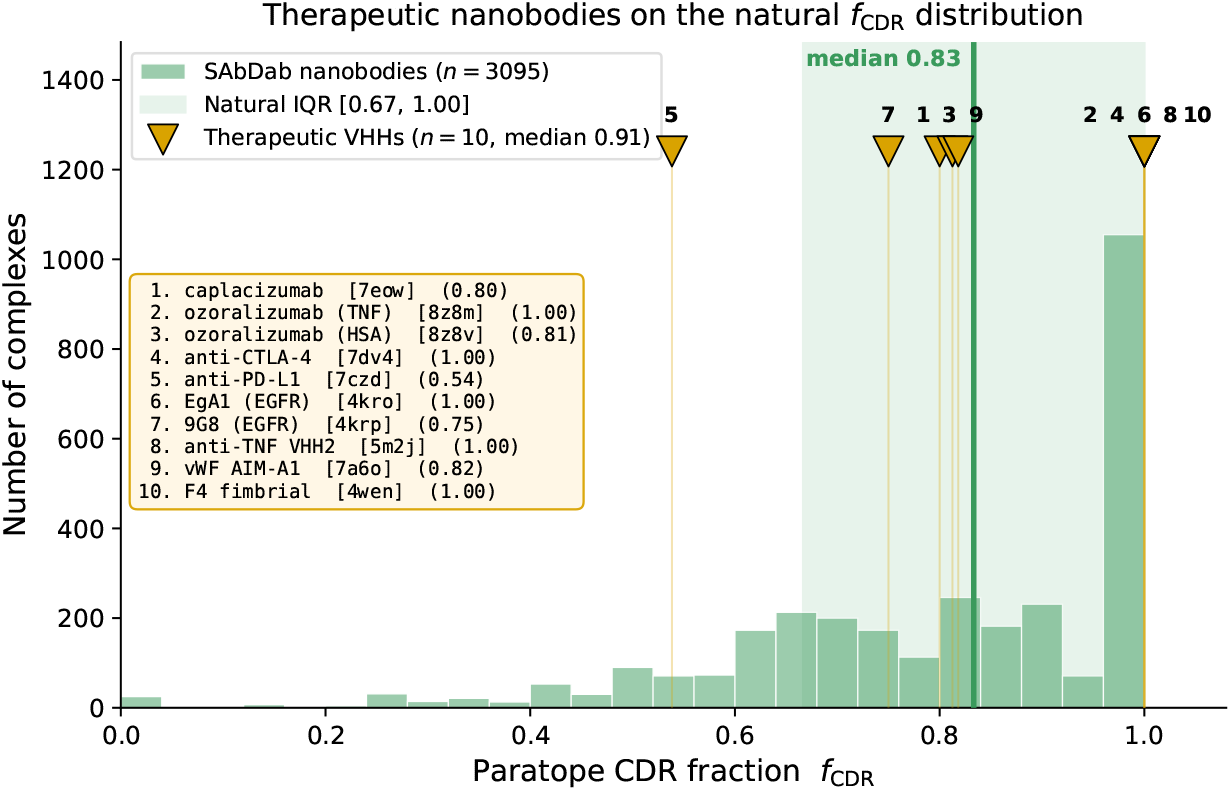
Reference paratope CDR fraction (*f*_CDR_) over 3,095 experimental nanobody–antigen complexes from SAbDab (29) (C*β* contacts within 6 Å, IMGT). Gold triangles mark 10 clinical/approved therapeutic VHHs (key inset, with PDB identifiers).

**Table S11.**
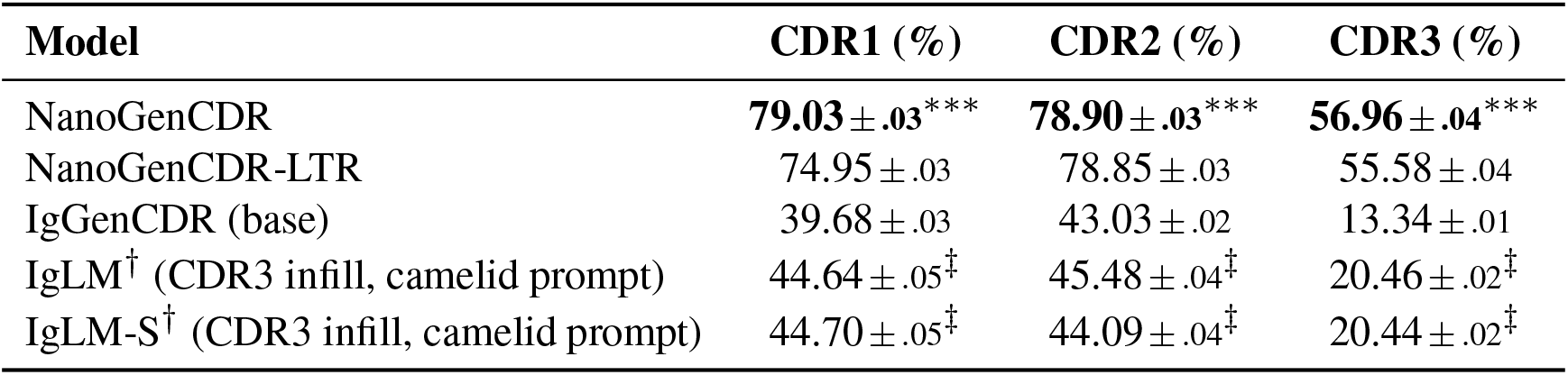
CDR amino acid recovery (NTP accuracy, %) on the INDI nanobody test set, averaged across numbering schemes (IMGT, Kabat, Chothia). Subscripts are one bootstrap standard error (2000 sequence-level resamples, resampling unit is the test sequence aligned across schemes). Stars mark the best-vs-second-best gap per column on the scheme-averaged per-sequence recovery (paired Wilcoxon, Holm-corrected across the three loops) and sit on the column-best (bold) cell: ^∗∗∗^*p* < 0.001, ^∗∗^*p* < 0.01, ^∗^*p* < 0.05. ^*†*^IgLM is prompted with its camelid species token. ^*‡*^IgLM evaluated under IMGT only with CDR3 masked, leveraging bidirectional context.

| Model | CDR1 (%) | CDR2 (%) | CDR3 (%) |
| --- | --- | --- | --- |
| NanoGenCDR | **79.03** ± .03*** | **78.90** ± .03*** | **56.96** ± .04*** |
| NanoGenCDR-LTR | 74.95 ± .03 | 78.85 ± .03 | 55.58 ± .04 |
| IgGenCDR (base) | 39.68 ± .03 | 43.03 ± .02 | 13.34 ± .01 |
| IgLM<sup>†</sup> (CDR3 infill, camelid prompt) | 44.64 ± .05<sup>‡</sup> | 45.48 ± .04<sup>‡</sup> | 20.46 ± .02<sup>‡</sup> |
| IgLM-S<sup>†</sup> (CDR3 infill, camelid prompt) | 44.70 ± .05<sup>‡</sup> | 44.09 ± .04<sup>‡</sup> | 20.44 ± .02<sup>‡</sup> |

**Table S12.** Per-scheme CDR amino acid recovery (NTP accuracy, %) on the INDI nanobody test set. IgLM and IgLM-S are evaluated under IMGT numbering only (CDR3-masked, bidirectional context) and are therefore reported only in the scheme-averaged Table S11. Subscripts are one bootstrap standard error (2000 sequence-level resamples). Stars mark the best-vs-second-best gap per column (paired Wilcoxon, Holm-corrected across the three loops) and sit on the column-best (bold) cell: ^∗∗∗^*p* < 0.001, ^∗∗^*p* < 0.01, ^∗^*p* < 0.05.

| Scheme | Model | CDR1 (%) | CDR2 (%) | CDR3 (%) |
| --- | --- | --- | --- | --- |
| IMGT | NanoGenCDR | **78.96** ± .05*** | **78.61** ± .05*** | **58.49** ± .06*** |
| | NanoGenCDR-LTR | 73.29 ± .05 | 78.16 ± .05 | 58.23 ± .06 |
| | IgGenCDR (base) | 44.58 ± .05 | 41.06 ± .04 | 15.12 ± .02 |
| Kabat | NanoGenCDR | **78.19** ± .05 | 82.57 ± .03 | **56.18** ± .07*** |
| | NanoGenCDR-LTR | 78.17 ± .05 | **83.46** ± .03*** | 55.75 ± .07 |
| | IgGenCDR (base) | 26.30 ± .04 | 55.63 ± .03 | 12.47 ± .02 |
| Chothia | NanoGenCDR | **79.94** ± .05*** | **75.51** ± .06*** | **56.20** ± .07*** |
| | NanoGenCDR-LTR | 73.38 ± .05 | 74.94 ± .06 | 55.75 ± .07 |
| | IgGenCDR (base) | 48.17 ± .05 | 32.41 ± .04 | 12.44 ± .02 |

**Table S13.**
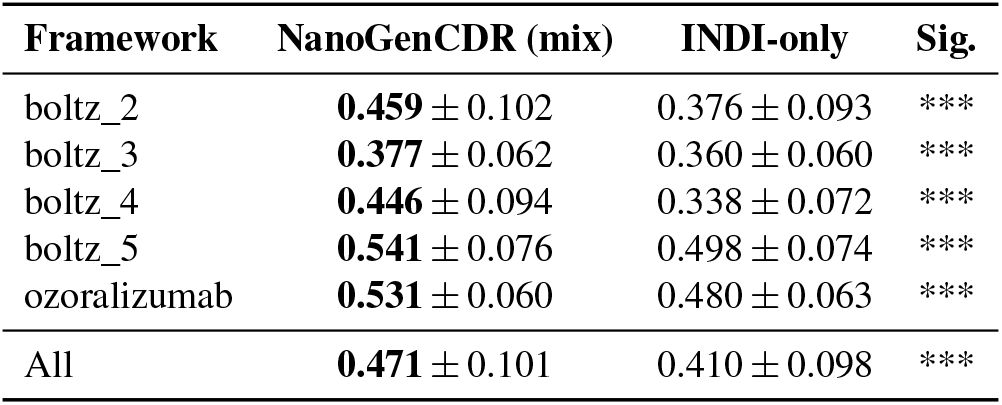
CDR-region OASis humanness (mean *±* s.d., *n*=500 per framework per model) of CDRs generated on humanised VHH scaffolds. The OAS mixture yields significantly more human CDRs on every framework. Significance from a two-sided Mann–Whitney *U* test: * *p<*0.05, ** *p<*0.01, *** *p<*0.001. **Bold**: higher mean.

| Framework | NanoGenCDR (mix) | INDI-only | Sig. |
| --- | --- | --- | --- |
| boltz_2 | **0.459** ± 0.102 | 0.376 ± 0.093 | *** |
| boltz_3 | **0.377** ± 0.062 | 0.360 ± 0.060 | *** |
| boltz_4 | **0.446** ± 0.094 | 0.338 ± 0.072 | *** |
| boltz_5 | **0.541** ± 0.076 | 0.498 ± 0.074 | *** |
| ozoralizumab | **0.531** ± 0.060 | 0.480 ± 0.063 | *** |
| All | **0.471** ± 0.101 | 0.410 ± 0.098 | *** |

The per-framework means (Table S13) show the OAS-mixture gain holds on every individual humanised scaffold, and on camelid INDI frameworks the two models are indistinguishable (median OASis 0.31 for both), as expected when the scaffold itself is non-human.

The effect is even more pronounced on a fully humanised acceptor framework. Repeating the experiment on the human IGHV3-23 (DP-47) germline framework, the canonical human acceptor toward which camelid VHHs are humanised but which retains no camelid hallmark residues and is therefore not included in the humanised-VHH average above, the OAS mixture reaches an OASis of 0.864 *±* 0.061 versus 0.691 *±* 0.071 for the INDI-only model (*n*=500 each, *p<*10^−100^, two-sided Mann–Whitney *U*). As the scaffold approaches a fully human framework, the human-distribution signal retained by the mixture becomes increasingly decisive.

#### B. Effect of the OAS mixture on recovery: a favourable trade-off

Table S14 details the recovery trade-off summarised in the main text (Section C), comparing NanoGenCDR (mix) and the INDI-only checkpoint on next-token CDR recovery (IMGT, heavy) on the camelid INDI and unpaired human OAS test sets. The collapse on human OAS is sharpest on the discriminative CDR3 loop, where the INDI-only model falls to 24.34% against the mixture’s 60.53%, indicating that camelid-only fine-tuning largely forgets the human CDR3 distribution while the mixture preserves it.

**Table S14.**
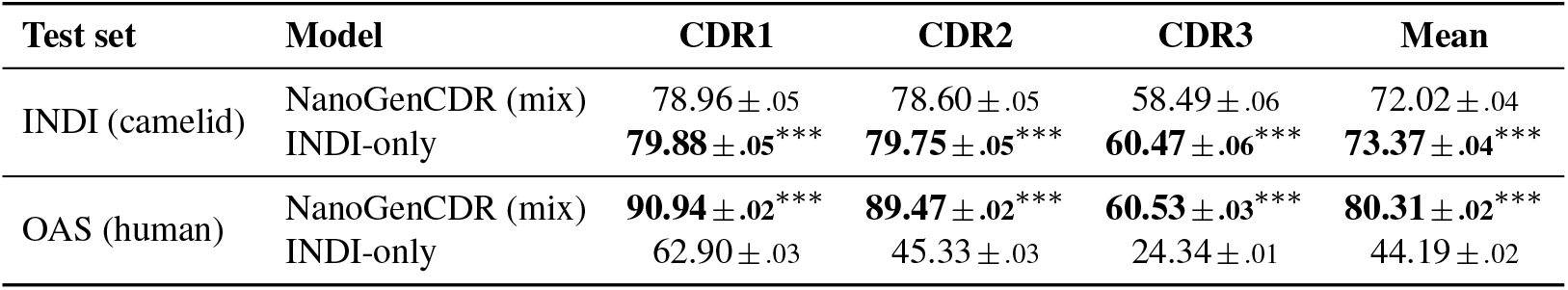
CDR amino acid recovery (NTP accuracy, %, IMGT, heavy) for the OAS-mixed vs INDI-only NanoGenCDR checkpoints on camelid (INDI) and human (OAS) test sets. The mixture costs 1.35 mean points on camelid but retains +36 mean points on human relative to the camelid-only model. Subscripts are one bootstrap standard error (2000 sequence-level resamples, resampling unit is the test sequence). Stars mark the mix vs INDI-only gap per loop (paired Wilcoxon on per-sequence recovery, Holm-corrected across the three loops and the mean) and sit on the better model: ^∗∗∗^*p* < 0.001, ^∗∗^*p* < 0.01, ^∗^*p* < 0.05.

#### C. Zero-shot fitness correlation on NbBench

We evaluate zero-shot fitness correlation on two nanobody thermostability assays from the NbBench collection (28): thermo-seq (*n*=764), measuring sequencing-based enrichment labels, and thermo-tm (*n*=567), measuring melting temperature (*T*_*m*_), with higher values preferable for both. All datasets contain single heavy-chain (VHH) sequences with a scalar fitness label.

Sequences are scored by mean per-token log-likelihood (Eq. S5) computed over the single heavy chain. Region-specific scores (Full, Framework, CDR) use the same token partitioning as in Section S8, with CDR boundaries defined by the indicated numbering scheme. For framework-first models (NanoGenCDR, IgGenCDR), the CDR log-likelihood is conditioned on all framework tokens, matching the training objective. IgLM (2) is evaluated in full-sequence left-to-right mode with its camelid species token.

**Table S15.**
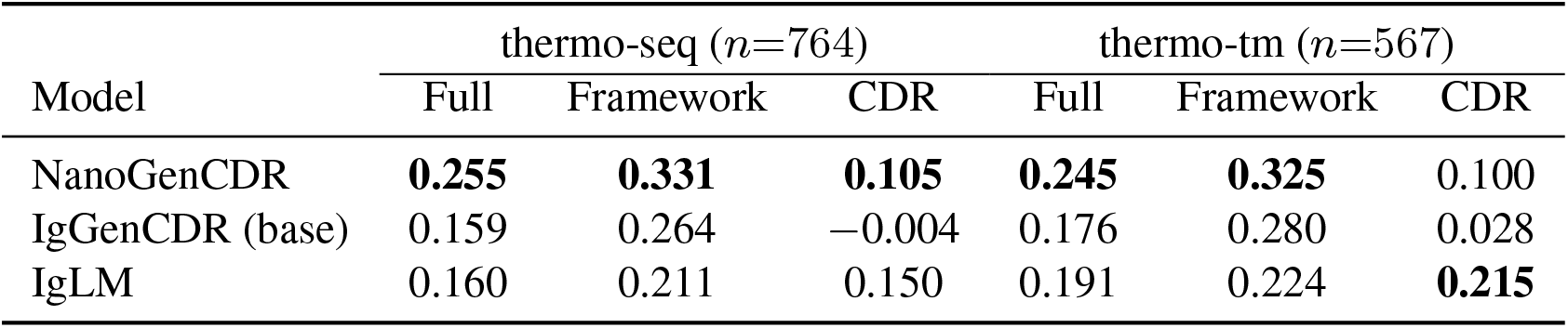
Log-likelihood–fitness correlation (Spearman *ρ*) on NbBench nanobody benchmarks, averaged across numbering schemes (IMGT, Kabat, Chothia). **Bold**: highest *ρ* in each column.

**Table S16.**
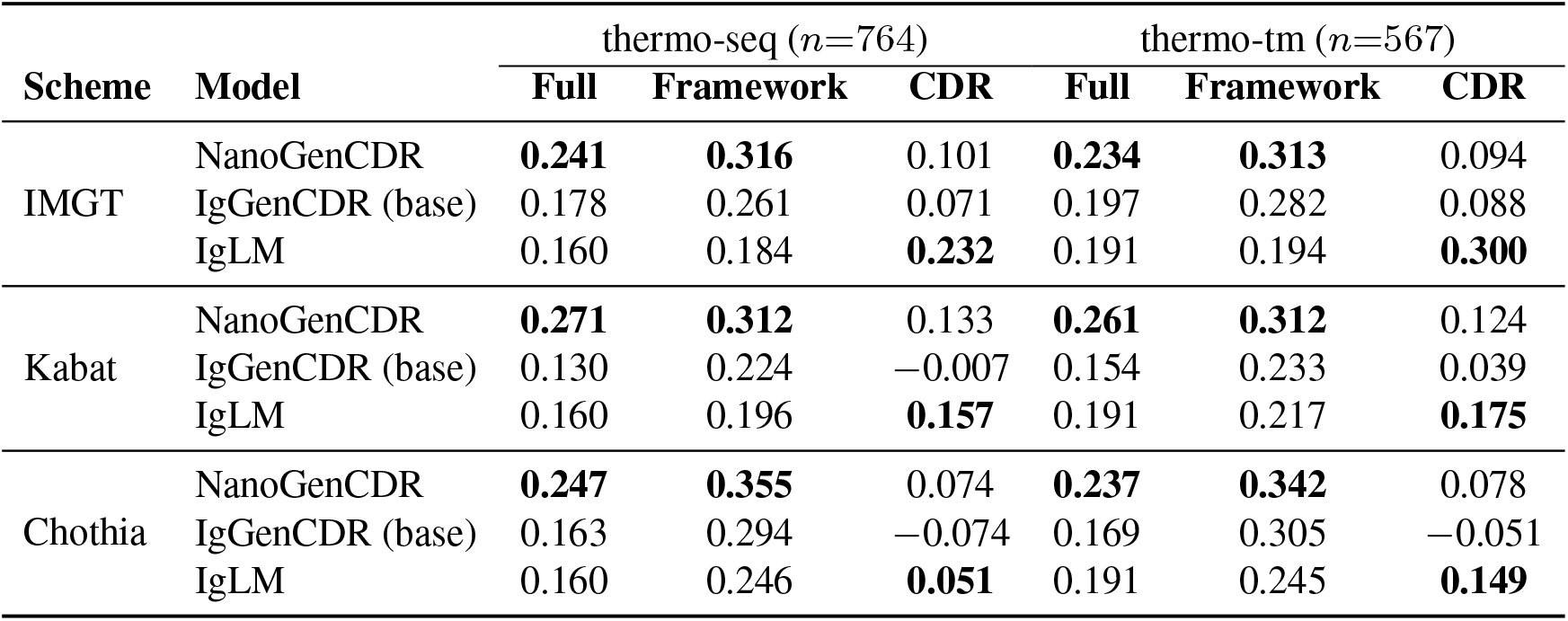
Per-scheme Spearman *ρ* between log-likelihood and label on NbBench thermostability benchmarks. **Bold**: highest *ρ* in each column within each numbering scheme.

**Fig. S13.**
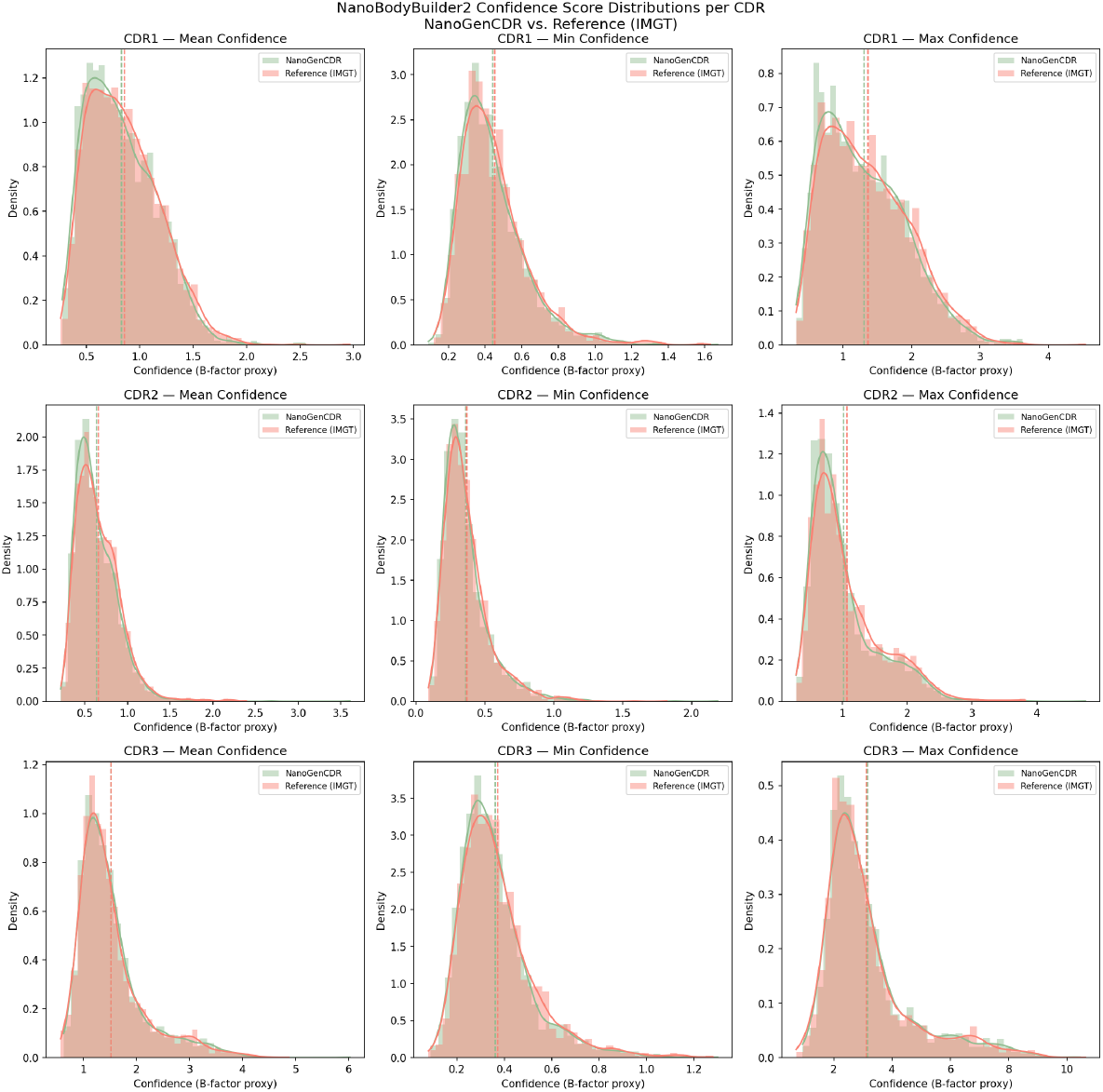
NanoBodyBuilder2 (26) mean confidence score per each CDR (IMGT), with minimum and maximum confidence values, based on b-factor proxy. NanoGenCDR results are overlaid against Reference set.

**Fig. S14.**
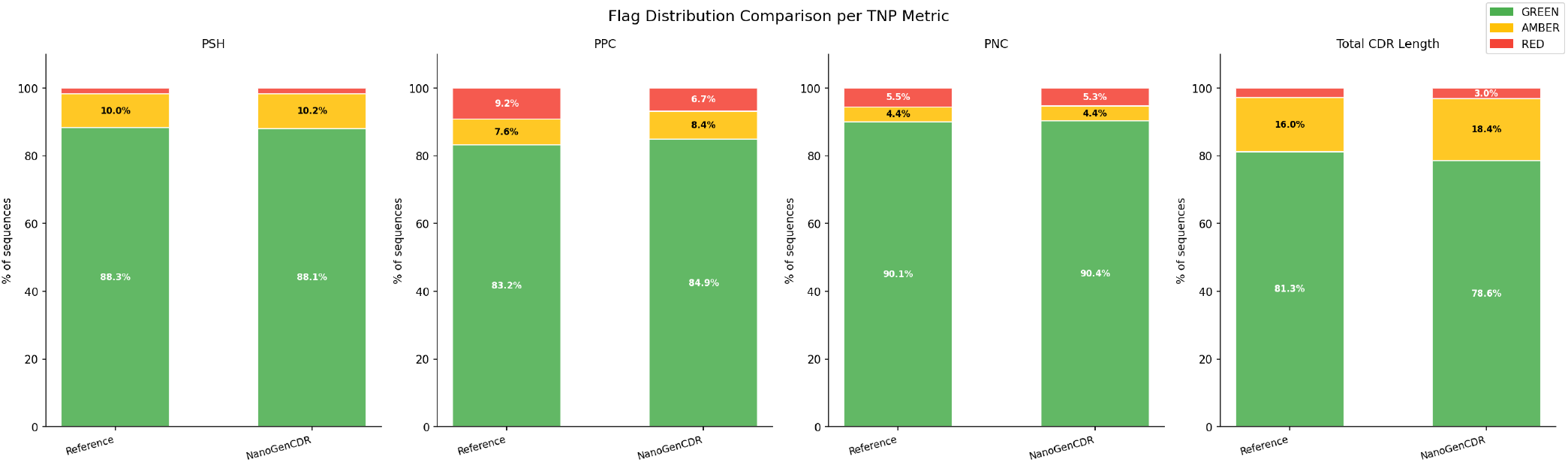
Developability flag distributions for NanoGenCDR versus the reference set, computed with the Therapeutic Nanobody Profiler (TNP) (27). PSH (Patch CDR Surface Hydrophobicity) flags hydrophobic surface patches on the CDRs that predispose to aggregation. PPC and PNC (Patch Positive and Negative Charge) flag charged CDR patches associated with poor specificity and clearance. Total IMGT CDR length flags unusually long combined loops, which tend to be harder to express and develop. Each metric is binned into TNP green/amber/red risk bands, and the plot compares the proportion of designs in each band against the reference cohort.

## Footnotes

1 https://zenodo.org/records/13880874

2 https://naturalantibody.com/indi2/

3 https://naturalantibody.com/indi2/

