## Supplementary material for "Generative Language Modeling for Antibody CDR Grafting and Alignment-driven De Novo Design": TeX files: bibliography.bib

@article{jain2017biophysical,
  title={Biophysical properties of the clinical-stage antibody landscape},
  author={Jain, Tushar and Sun, Tingwan and Durand, St{\'e}phanie and Hall, Amy and Houston, Nga Rewa and Nett, Juergen H and Sharkey, Beth and Bobrowicz, Beata and Caffry, Isabelle and Yu, Yao and others},
  journal={Proceedings of the National Academy of Sciences},
  volume={114},
  number={5},
  pages={944--949},
  year={2017},
  publisher={National Academy of Sciences}
}

@inproceedings{rafailov2023direct,
  title={Direct Preference Optimization: Your Language Model is Secretly a Reward Model},
  author={Rafailov, Rafael and Sharma, Archit and Mitchell, Eric and Manning, Christopher D and Ermon, Stefano and Finn, Chelsea},
  booktitle={Advances in Neural Information Processing Systems},
  volume={36},
  year={2023}
}

@article{richemond2024offline,
  title={Offline Regularised Reinforcement Learning for Large Language Models Alignment},
  author={Richemond, Pierre Harvey and Tang, Yunhao and Guo, Daniel and Calandriello, Daniele and Azar, Mohammad Gheshlaghi and Rafailov, Rafael and Pires, Bernardo Avila and Tarassov, Eugene and Spangher, Lucas and Ellsworth, Will and others},
  journal={arXiv preprint arXiv:2405.19107},
  year={2024}
}

@article{widatalla2024aligning,
  title={Aligning protein generative models with experimental fitness via Direct Preference Optimization},
  author={Widatalla, Talal and Rafailov, Rafael and Hie, Brian},
  journal={bioRxiv},
  year={2024},
  doi={10.1101/2024.05.20.595026}
}

@article{turnbull2024p,
  title={p-IgGen: a paired antibody generative language model},
  author={Turnbull, Oliver M and Oglic, Dino and Croasdale-Wood, Rebecca and Deane, Charlotte M},
  journal={Bioinformatics},
  volume={40},
  number={11},
  pages={btae659},
  year={2024},
  publisher={Oxford University Press}
}
@article{shuai2023iglm,
  title={IgLM: Infilling language modeling for antibody sequence design},
  author={Shuai, Richard W and Ruffolo, Jeffrey A and Gray, Jeffrey J},
  journal={Cell systems},
  volume={14},
  number={11},
  pages={979--989},
  year={2023},
  publisher={Elsevier}
}
@article{nijkamp2023progen2,
  title={Progen2: exploring the boundaries of protein language models},
  author={Nijkamp, Erik and Ruffolo, Jeffrey A and Weinstein, Eli N and Naik, Nikhil and Madani, Ali},
  journal={Cell systems},
  volume={14},
  number={11},
  pages={968--978},
  year={2023},
  publisher={Elsevier}
}
@article{suzuki2015therapeutic,
  title={Therapeutic antibodies: their mechanisms of action and the pathological findings they induce in toxicity studies},
  author={Suzuki, Masami and Kato, Chie and Kato, Atsuhiko},
  journal={Journal of toxicologic pathology},
  volume={28},
  number={3},
  pages={133--139},
  year={2015},
  publisher={JAPANESE SOCIETY OF TOXICOLOGIC PATHOLOGY}
}
@article{lee2025reinforcement,
  title={Reinforcement Learning for Antibody Sequence Infilling},
  author={Lee, Chak Shing and Hayes, Conor F and Vashchenko, Denis and Landajuela, Mikel},
  journal={bioRxiv},
  pages={2025--08},
  year={2025},
  publisher={Cold Spring Harbor Laboratory}
}

@article{uccar2024exploring,
  title={Exploring log-likelihood scores for ranking antibody sequence designs},
  author={U{\c{c}}ar, Talip and Malherbe, Cedric and Gonzalez, Ferran},
  journal={bioRxiv},
  pages={2024--10},
  year={2024},
  publisher={Cold Spring Harbor Laboratory}
}
@article{greenig2025igcraft,
  title={IgCraft: A versatile sequence generation framework for antibody discovery and engineering},
  author={Greenig, Matthew and Zhao, Haowen and Radenkovic, Vladimir and Ramon, Aubin and Sormanni, Pietro},
  journal={arXiv preprint arXiv:2503.19821},
  year={2025}
}
@article{olsen2022observed,
  title={Observed Antibody Space: A diverse database of cleaned, annotated, and translated unpaired and paired antibody sequences},
  author={Olsen, Tobias H and Boyles, Fergus and Deane, Charlotte M},
  journal={Protein Science},
  volume={31},
  number={1},
  pages={141--146},
  year={2022},
  publisher={Wiley Online Library}
}
@article{dunbar2016anarci,
  title={ANARCI: antigen receptor numbering and receptor classification},
  author={Dunbar, James and Deane, Charlotte M},
  journal={Bioinformatics},
  volume={32},
  number={2},
  pages={298--300},
  year={2016},
  publisher={Oxford University Press}
}
@article{dunbar2014sabdab,
  title={SAbDab: the structural antibody database},
  author={Dunbar, James and Krawczyk, Konrad and Leem, Jinwoo and Baker, Terry and Fuchs, Angelika and Georges, Guy and Shi, Jiye and Deane, Charlotte M},
  journal={Nucleic Acids Research},
  volume={42},
  number={D1},
  pages={D1140--D1146},
  year={2014},
  publisher={Oxford University Press}
}
@article{deszynski2022indi,
  title={INDI—integrated nanobody database for immunoinformatics},
  author={Deszy{\'n}ski, Piotr and M{\l}okosiewicz, Jakub and Volanakis, Adam and Jaszczyszyn, Igor and Castellana, Natalie and Bonissone, Stefano and Ganesan, Rajkumar and Krawczyk, Konrad},
  journal={Nucleic Acids Research},
  volume={50},
  number={D1},
  pages={D1273--D1281},
  year={2022},
  publisher={Oxford University Press}
}
@article{passaro2025boltz,
  title={Boltz-2: Towards accurate and efficient binding affinity prediction},
  author={Passaro, Saro and Corso, Gabriele and Wohlwend, Jeremy and Reveiz, Mateo and Thaler, Stephan and Somnath, Vignesh Ram and Getz, Noah and Portnoi, Tally and Roy, Julien and Stark, Hannes and others},
  journal={BioRxiv},
  year={2025}
}
@article{touvron2023llama,
  title={{LLaMA}: Open and efficient foundation language models},
  author={Touvron, Hugo and Lavril, Thibaut and Izacard, Gautier and Martinet, Xavier and Lachaux, Marie-Anne and Lacroix, Timoth{\'e}e and Rozi{\`e}re, Baptiste and Goyal, Naman and Hambro, Eric and Azhar, Faisal and others},
  journal={arXiv preprint arXiv:2302.13971},
  year={2023}
}
@article{stark2025boltzgen,
  title={Boltzgen: Toward universal binder design},
  author={Stark, Hannes and Faltings, Felix and Choi, MinGyu and Xie, Yuxin and Hur, Eunsu and O’Donnell, Timothy and Bushuiev, Anton and U{\c{c}}ar, Talip and Passaro, Saro and Mao, Weian and others},
  journal={bioRxiv},
  pages={2025--11},
  year={2025},
  publisher={Cold Spring Harbor Laboratory}
}
@article{bennett2026atomically,
  title={Atomically accurate de novo design of antibodies with RFdiffusion},
  author={Bennett, Nathaniel R and Watson, Joseph L and Ragotte, Robert J and Borst, Andrew J and See, D{\'e}Jena{\'e} L and Weidle, Connor and Biswas, Riti and Yu, Yutong and Shrock, Ellen L and Ault, Russell and others},
  journal={Nature},
  volume={649},
  number={8095},
  pages={183--193},
  year={2026},
  publisher={Nature Publishing Group UK London}
}
@article{mille2025efficient,
  title={Efficient generation of epitope-targeted de novo antibodies with Germinal},
  author={Mille-Fragoso, Luis S and Wang, John N and Driscoll, Claudia L and Dai, Haoyu and Widatalla, Talal and Zhang, Xiaowei and Hie, Brian L and Gao, Xiaojing J},
  journal={bioRxiv},
  year={2025}
}
@article{dauparas2022robust,
  title={Robust deep learning--based protein sequence design using ProteinMPNN},
  author={Dauparas, Justas and Anishchenko, Ivan and Bennett, Nathaniel and Bai, Hua and Ragotte, Robert J and Milles, Lukas F and Wicky, Basile IM and Courbet, Alexis and de Haas, Rob J and Bethel, Neville and others},
  journal={Science},
  volume={378},
  number={6615},
  pages={49--56},
  year={2022},
  publisher={American Association for the Advancement of Science}
}
@article{li2006cd,
  title={Cd-hit: a fast program for clustering and comparing large sets of protein or nucleotide sequences},
  author={Li, Weizhong and Godzik, Adam},
  journal={Bioinformatics},
  volume={22},
  number={13},
  pages={1658--1659},
  year={2006},
  publisher={Oxford University Press}
}
@article{olsen2022ablang,
  title={{AbLang}: an antibody language model for completing antibody sequences},
  author={Olsen, Tobias H and Moal, Iain H and Deane, Charlotte M},
  journal={Bioinformatics Advances},
  volume={2},
  number={1},
  pages={vbac046},
  year={2022},
  publisher={Oxford University Press}
}
@article{prihoda2022biophi,
  title={BioPhi: A platform for antibody design, humanization, and humanness evaluation based on natural antibody repertoires and deep learning},
  author={Prihoda, David and Maamary, Jad and Waight, Andrew and Juan, Veronica and Fayadat-Dilman, Laurence and Svozil, Daniel and Bitton, Danny A},
  journal={mAbs},
  volume={14},
  number={1},
  pages={2020203},
  year={2022},
  publisher={Taylor \& Francis}
}
@article{lin2023esm2,
  title={Evolutionary-scale prediction of atomic-level protein structure with a language model},
  author={Lin, Zeming and Akin, Halil and Rao, Roshan and Hie, Brian and Zhu, Zhongkai and Lu, Wenting and Smetanin, Nikita and Verkuil, Robert and Kabeli, Ori and Shmueli, Yaniv and others},
  journal={Science},
  volume={379},
  number={6637},
  pages={1123--1130},
  year={2023},
  publisher={American Association for the Advancement of Science}
}
@inproceedings{notin2023proteingym,
  title={{ProteinGym}: Large-scale benchmarks for protein fitness prediction and design},
  author={Notin, Pascal and Kollasch, Aaron and Ritter, Daniel and van Niekerk, Lood and Paul, Steffanie and Spinner, Han and Rollins, Nathan and Shaw, Ada and Orenbuch, Rose and Weitzman, Ruben and others},
  booktitle={Advances in Neural Information Processing Systems},
  volume={36},
  year={2023}
}
@article{zhang2025nbbench,
  title={{NbBench}: A comprehensive benchmark suite for nanobody representation learning},
  author={Zhang, Yiming and Tsuda, Koji},
  journal={arXiv preprint arXiv:2505.02022},
  year={2025}
}
@book{lefranc2001immunoglobulins,
  title={The Immunoglobulin FactsBook},
  author={Lefranc, Marie-Paule and Lefranc, G{\'e}rard},
  year={2001},
  publisher={Academic Press},
  address={London}
}
@article{xu2000diversity,
  title={Diversity in the {CDR3} region of {VH} is sufficient for most antibody specificities},
  author={Xu, Jian Lian and Davis, Mark M},
  journal={Immunity},
  volume={13},
  number={1},
  pages={37--45},
  year={2000},
  publisher={Elsevier}
}
@article{abanades2023immunebuilder,
  title={{ImmuneBuilder}: Deep-learning models for predicting the structures of immune proteins},
  author={Abanades, Brennan and Wong, Wing Ki and Boyles, Fergus and Georges, Guy and Bujotzek, Alexander and Deane, Charlotte M},
  journal={Communications Biology},
  volume={6},
  number={1},
  pages={575},
  year={2023},
  publisher={Nature Publishing Group UK London}
}
@article{stocco2024guiding,
  title={Guiding Protein Language Models with Reinforcement Learning},
  author={Stocco, Filippo and Artigues-Lleix{\`a}, Maria and Hunklinger, Andrea and Widatalla, Talal and G{\"u}ell, Marc and Ferruz, Noelia},
  journal={arXiv preprint arXiv:2412.12979},
  year={2024}
}
@article{stocco2025steering,
  title={Steering Generative Models for Protein Design: Aligning and Conditioning Strategies},
  author={Stocco, Filippo and Garibbo, Michele and Ferruz, Noelia},
  journal={arXiv preprint arXiv:2511.21476},
  year={2025}
}
@article{dunbrack2025res,
  title={R{\=e}s ipSAE loquunt: What’s wrong with AlphaFold’s ipTM score and how to fix it},
  author={Dunbrack Jr, Roland L},
  journal={bioRxiv},
  year={2025}
}

@article{koenig2017mutational,
  title={Mutational landscape of antibody variable domains reveals a switch modulating the interdomain conformational dynamics and antigen binding},
  author={Koenig, Patrick and Lee, Chingwei V and Walters, Benjamin T and Janakiraman, Vasantharajan and Stinson, Jeremy and Patapoff, Thomas W and Fuh, Germaine},
  journal={Proceedings of the National Academy of Sciences},
  volume={114},
  number={4},
  pages={E486--E495},
  year={2017},
  publisher={National Academy of Sciences}
}

@article{marks2021humanization,
  title={Humanization of antibodies using a machine learning approach on large-scale repertoire data},
  author={Marks, Claire and Hummer, Alissa M and Chin, Mark and Deane, Charlotte M},
  journal={Bioinformatics},
  volume={37},
  number={22},
  pages={4041--4047},
  year={2021},
  publisher={Oxford University Press}
}

%% TODO(author): verify full author list and article/page number before submission
%% (Sci. Rep. 2024, DOI 10.1038/s41598-024-70513-4).
@article{humanizedvhh2024,
  title={Development of novel humanized VHH synthetic libraries based on physicochemical analyses},
  author={Nakakido, Makoto and Kinoshita, Seisho and Tsumoto, Kouhei},
  journal={Scientific Reports},
  volume={14},
  number={1},
  pages={19533},
  year={2024},
  publisher={Nature Publishing Group UK London}
}
@article{gordon2025therapeutic,
  title={The therapeutic nanobody profiler: characterising and predicting nanobody developability to improve therapeutic design},
  author={Gordon, Gemma L and Gervasio, Jo{\~a}o and Souders, Colby and Deane, Charlotte M},
  journal={BioRXiv},
  pages={2025--08},
  year={2025},
  publisher={Cold Spring Harbor Laboratory}
}
@article{schulman2017proximal,
  title={Proximal policy optimization algorithms},
  author={Schulman, John and Wolski, Filip and Dhariwal, Prafulla and Radford, Alec and Klimov, Oleg},
  journal={arXiv preprint arXiv:1707.06347},
  year={2017}
}
