## Supplementary material for "Generative Language Modeling for Antibody CDR Grafting and Alignment-driven De Novo Design": TeX files: reference.bib

@inproceedings{rahman2019centroidb,
  title={Centroid of Age Neighborhoods: A Generalized Approach to Estimate Biological Age},
  author={Rahman, Syed Ashiqur and Abdul, Donald},
  booktitle={2019 IEEE EMBS International Conference on Biomedical \& Health Informatics (BHI)},
  pages={1--4},
  year={2019},
  organization={IEEE}
}
@article{bahdanau2014neural,
  title={Neural machine translation by jointly learning to align and translate},
  author={Bahdanau, Dzmitry and Cho, Kyunghyun and Bengio, Yoshua},
  journal={arXiv preprint arXiv:1409.0473},
  year={2014}
}
@article{imboden2018cardiorespiratory,
  title={Cardiorespiratory fitness and mortality in healthy men and women},
  author={Imboden, Mary T and Harber, Matthew P and Whaley, Mitchell H and Finch, W Holmes and Bishop, Derron L and Kaminsky, Leonard A},
  journal={Journal of the American College of Cardiology},
  volume={72},
  number={19},
  pages={2283--2292},
  year={2018},
  publisher={Elsevier}
}
@inproceedings{motiian2017unified,
  title={Unified deep supervised domain adaptation and generalization},
  author={Motiian, Saeid and Piccirilli, Marco and Adjeroh, Donald A and Doretto, Gianfranco},
  booktitle={Proceedings of the IEEE International Conference on Computer Vision},
  pages={5715--5725},
  year={2017}
}
@book{american2013acsm,
  title={ACSM's guidelines for exercise testing and prescription},
  author={American College of Sports Medicine and others},
  year={2013},
  publisher={Lippincott Williams \& Wilkins}
}
@book{murphy2012machine,
  title={Machine learning: A probabilistic perspective},
  author={Murphy, Kevin P},
  year={2012},
  publisher={MIT press}
}
@article{ji20123d,
  title={3D convolutional neural networks for human action recognition},
  author={Ji, Shuiwang and Xu, Wei and Yang, Ming and Yu, Kai},
  journal={IEEE Transactions on Pattern Analysis and Machine Intelligence},
  volume={35},
  number={1},
  pages={221--231},
  year={2012},
  publisher={IEEE}
}
@inproceedings{krizhevsky2012imagenet,
  title={Imagenet classification with deep convolutional neural networks},
  author={Krizhevsky, Alex and Sutskever, Ilya and Hinton, Geoffrey E},
  booktitle={Advances in Neural Information Processing Systems},
  pages={1097--1105},
  year={2012}
}
@article{horvath2018dna,
  title={DNA methylation-based biomarkers and the epigenetic clock theory of ageing},
  author={Horvath, Steve and Raj, Kenneth},
  journal={Nature Reviews Genetics},
  volume={19},
  number={6},
  pages={371},
  year={2018},
  publisher={Nature Publishing Group}
}
@article{pyrkov2018quantitative,
  title={Quantitative characterization of biological age and frailty based on locomotor activity records},
  author={Pyrkov, Timothy V and Getmantsev, Evgeny and Zhurov, Boris and Avchaciov, Konstantin and Pyatnitskiy, Mikhail and Menshikov, Leonid and Khodova, Kristina and Gudkov, Andrei V and Fedichev, Peter O},
  journal={Aging (Albany NY)},
  volume={10},
  number={10},
  pages={2973},
  year={2018},
  publisher={Impact Journals, LLC}
}
@inproceedings{wang2018face,
  title={Face aging with identity-preserved conditional generative adversarial networks},
  author={Wang, Zongwei and Tang, Xu and Luo, Weixin and Gao, Shenghua},
  booktitle={Proceedings of the IEEE Conference on Computer Vision and Pattern Recognition},
  pages={7939--7947},
  year={2018}
}
@article{lecun2015deep,
  title={Deep learning},
  author={LeCun, Yann and Bengio, Yoshua and Hinton, Geoffrey},
  journal={Nature},
  volume={521},
  number={7553},
  pages={436},
  year={2015},
  publisher={Nature Publishing Group}
}
@article{zhang2018fine,
  title={Fine-Grained Age Estimation in the wild with Attention {LSTM} Networks},
  author={Zhang, Ke and Liu, Na and Yuan, Xingfang and Guo, Xinyao and Gao, Ce and Zhao, Zhenbing},
  journal={arXiv preprint arXiv:1805.10445},
  year={2018}
}
@article{ravi2016deep,
  title={Deep learning for health informatics},
  author={Rav{\`\i}, Daniele and Wong, Charence and Deligianni, Fani and Berthelot, Melissa and Andreu-Perez, Javier and Lo, Benny and Yang, Guang-Zhong},
  journal={IEEE {J}ournal of {B}iomedical and {H}ealth {I}nformatics},
  volume={21},
  number={1},
  pages={4--21},
  year={2016},
  publisher={IEEE}
}
@article{miotto2017deep,
  title={Deep learning for healthcare: review, opportunities and challenges},
  author={Miotto, Riccardo and Wang, Fei and Wang, Shuang and Jiang, Xiaoqian and Dudley, Joel T},
  journal={Briefings in {B}ioinformatics},
  volume={19},
  number={6},
  pages={1236--1246},
  year={2017},
  publisher={Oxford University Press}
}
@article{rahman2019centroida,
  title={Centroid of Age Neighborhoods: A New Approach to Estimate Biological Age},
  author={Rahman, Syed Ashiqur and Adjeroh, Donald},
  journal={IEEE {J}ournal of {B}iomedical and {H}ealth {I}nformatics},
  year={2019},
  publisher={IEEE}
}
@article{rahman2019deep,
  title={Deep Learning using Convolutional LSTM estimates Biological Age from Physical Activity},
  author={Rahman, Syed Ashiqur and Adjeroh, Donald A},
  journal={Scientific {R}eports},
  volume={9},
  number={1},
  pages={1--15},
  year={2019},
  publisher={Nature Publishing Group}
}
@inproceedings{goodfellow2014generative,
  title={Generative adversarial nets},
  author={Goodfellow, Ian and Pouget-Abadie, Jean and Mirza, Mehdi and Xu, Bing and Warde-Farley, David and Ozair, Sherjil and Courville, Aaron and Bengio, Yoshua},
  booktitle={Advances in {N}eural {I}nformation {P}rocessing {S}ystems},
  pages={2672--2680},
  year={2014}
}
@article{fu2019predicting,
  title={Predicting DNA methylation states with hybrid information based deep-learning model},
  author={Fu, Laiyi and Peng, Qinke and Chai, Ling},
  journal={IEEE/ACM Transactions on Computational Biology and Bioinformatics},
  year={2019},
  publisher={IEEE}
}
@article{liu2016pedla,
  title={PEDLA: predicting enhancers with a deep learning-based algorithmic framework},
  author={Liu, Feng and Li, Hao and Ren, Chao and Bo, Xiaochen and Shu, Wenjie},
  journal={Scientific {R}eports},
  volume={6},
  pages={28517},
  year={2016},
  publisher={Nature Publishing Group}
}
@article{cole2017predicting,
  title={Predicting brain age with deep learning from raw imaging data results in a reliable and heritable biomarker},
  author={Cole, James H and Poudel, Rudra PK and Tsagkrasoulis, Dimosthenis and Caan, Matthan WA and Steves, Claire and Spector, Tim D and Montana, Giovanni},
  journal={NeuroImage},
  volume={163},
  pages={115--124},
  year={2017},
  publisher={Elsevier}
}
@article{wang2017predicting,
  title={Predicting age by mining electronic medical records with deep learning characterizes differences between chronological and physiological age},
  author={Wang, Zichen and Li, Li and Glicksberg, Benjamin S and Israel, Ariel and Dudley, Joel T and Ma'ayan, Avi},
  journal={Journal of Biomedical Informatics},
  volume={76},
  pages={59--68},
  year={2017},
  publisher={Elsevier}
}
@article{hochschild1989improving,
  title={Improving the precision of biological age determinations. Part 1: A new approach to calculating biological age},
  author={Hochschild, Richard},
  journal={Experimental Gerontology},
  volume={24},
  number={4},
  pages={289--300},
  year={1989},
  publisher={Elsevier}
}
@article{kroll2000use,
  title={On the use of regression analysis for the estimation of human biological age},
  author={Kr{\o}ll, J and Saxtrup, O},
  journal={Biogerontology},
  volume={1},
  number={4},
  pages={363--368},
  year={2000},
  publisher={Springer}
}
@article{liu2018phenotypic,
  title={Phenotypic Age: A novel signature of mortality and morbidity risk},
  author={Liu, Zuyun and Kuo, Pei-Lun and Horvath, Steve and Crimmins, Eileen and Ferrucci, Luigi and Levine, Morgan},
  journal={BioRxiv},
  pages={363291},
  year={2018},
  publisher={Cold Spring Harbor Laboratory}
}
@article{belsky2017eleven,
  title={Eleven telomere, epigenetic clock, and biomarker-composite quantifications of biological aging: do they measure the same thing?},
  author={Belsky, Daniel W and Moffitt, Terrie E and Cohen, Alan A and Corcoran, David L and Levine, Morgan E and Prinz, Joseph A and Schaefer, Jonathan and Sugden, Karen and Williams, Benjamin and Poulton, Richie and others},
  journal={American Journal of Epidemiology},
  volume={187},
  number={6},
  pages={1220--1230},
  year={2017},
  publisher={Oxford University Press}
}
@article{mamoshina2018population,
  title={Population Specific Biomarkers of Human Aging: A Big Data Study Using {S}outh {K}orean, {C}anadian, and {E}astern {E}uropean Patient Populations},
  author={Mamoshina, Polina and Kochetov, Kirill and Putin, Evgeny and Cortese, Franco and Aliper, Alexander and Lee, Won-Suk and Ahn, Sung-Min and Uhn, Lee and Skjodt, Neil and Kovalchuk, Olga and others},
  journal={The Journals of Gerontology: Series A},
  volume={73},
  number={11},
  pages={1482--1490},
  year={2018},
  publisher={Oxford University Press US}
}
@article{bobrov2018photoageclock,
  title={Photo{A}ge{C}lock: Deep learning algorithms for development of non-invasive visual biomarkers of aging},
  author={Bobrov, Eugene and Georgievskaya, Anastasia and Kiselev, Konstantin and Sevastopolsky, Artem and Zhavoronkov, Alex and Gurov, Sergey and Rudakov, Konstantin and Tobar, Maria del Pilar Bonilla and Jaspers, S{\"o}ren and Clemann, Sven},
  journal={Aging (Albany NY)},
  volume={10},
  number={11},
  pages={3249},
  year={2018},
  publisher={Impact Journals, LLC}
}
@article{zhavoronkov2018artificial,
  title={Artificial intelligence for aging and longevity research: Recent advances and perspectives},
  author={Zhavoronkov, Alex and Mamoshina, Polina and Vanhaelen, Quentin and Scheibye-Knudsen, Morten and Moskalev, Alexey and Aliper, Alex},
  journal={Ageing Research Reviews},
  year={2018},
  publisher={Elsevier}
}
@article{srivastava2014dropout,
  title={Dropout: A simple way to prevent neural networks from overfitting},
  author={Srivastava, Nitish and Hinton, Geoffrey and Krizhevsky, Alex and Sutskever, Ilya and Salakhutdinov, Ruslan},
  journal={The Journal of Machine Learning Research},
  volume={15},
  number={1},
  pages={1929--1958},
  year={2014},
  publisher={JMLR. org}
}
@article{dozat2016incorporating,
  title={Incorporating Nesterov Momentum into Adam},
  author={Dozat, Timothy},
  year={2016},
  journal={ICLR Workshop}
}
@ONLINE{hintonRmspropOnline,
  author = {},
  title = {Neural Networks for Machine Learning. {@ONLINE}},
  month = {November},
  year = {2018},
  url = { http://www.cs.toronto.edu/~tijmen/csc321/slides/lecture_slides_lec6.pdf},
  urldate = {2018-11-15}
}
@article{morkedal2011informativeness,
  title={Informativeness of indices of blood pressure, obesity and serum lipids in relation to ischaemic heart disease mortality: {T}he {HUNT-II} study},
  author={M{\o}rkedal, Bj{\o}rn and Romundstad, P{\aa}l R and Vatten, Lars J},
  journal={European Journal of Epidemiology},
  volume={26},
  number={6},
  pages={457--461},
  year={2011},
  publisher={Springer}
}
@article{kingma2014adam,
  title={Adam: A method for stochastic optimization},
  author={Kingma, Diederik P and Ba, Jimmy},
  journal={arXiv preprint arXiv:1412.6980},
  year={2014}
}
@article{kaplan1958nonparametric,
  title={Nonparametric estimation from incomplete observations},
  author={Kaplan, Edward L and Meier, Paul},
  journal={Journal of the American Statistical Association},
  volume={53},
  number={282},
  pages={457--481},
  year={1958},
  publisher={Taylor \& Francis}
}
@article{dipietro1993descriptive,
  title={The descriptive epidemiology of selected physical activities and body weight among adults trying to lose weight: the Behavioral Risk Factor Surveillance System survey, 1989.},
  author={DiPietro, L and Williamson, DF and Caspersen, CJ and Eaker, E},
  journal={International Journal of Obesity and Related Metabolic Disorders: Journal of the International Association for the Study of Obesity},
  volume={17},
  number={2},
  pages={69--76},
  year={1993}
}

@article{stephens1994demography,
  title={The demography of physical activity.},
  author={Stephens, Thomas and Caspersen, Carl J},
  journal={Physical Activity, Fitness, and Health: International Proceedings and Consensus Statement},
  pages={204--213},
  year={1994},
  publisher={Champaign, IL, England: Human Kinetics Publishers}
}

@article{caspersen1991national,
  title={National estimates of physical activity among older adults},
  author={Caspersen, CJ and DiPietro, L},
  journal={Medicine and Science in Sports and Exercise},
  volume={23},
  number={4 supplement},
  pages={106},
  year={1991}
}

@article{dipietro2001physical,
  title={Physical activity in aging: changes in patterns and their relationship to health and function},
  author={DiPietro, Loretta},
  journal={The Journals of Gerontology Series A: Biological Sciences and Medical Sciences},
  volume={56},
  number={suppl\_2},
  pages={13--22},
  year={2001},
  publisher={Oxford University Press}
}

@ONLINE{ispah2018Online,
  author = {},
  title = {{I}nternational {S}ociety for {P}hysical {A}ctivity and {H}ealth. {@ONLINE}},
  month = {September},
  year = {2018},
  url = { http://www.ispah.org/},
  urldate = {2018-10-16}
}

@ONLINE{who2018Online,
  author = {},
  title = {{WHO}. {G}lobal {P}hysical {A}ctivity {S}urveillance. {@ONLINE}},
  month = {September},
  year = {2018},
  url = { http://www.who.int/ncds/surveillance/steps/GPAQ/en/},
  urldate = {2018-10-16}
}

@article{guthold2018worldwide,
  title={Worldwide trends in insufficient physical activity from 2001 to 2016: a pooled analysis of 358 population-based surveys with 1{\textperiodcentered} 9 million participants},
  author={Guthold, Regina and Stevens, Gretchen A and Riley, Leanne M and Bull, Fiona C},
  journal={The Lancet Global Health},
  year={2018},
  publisher={Elsevier}
}

@inproceedings{glorot2010understanding,
  title={Understanding the difficulty of training deep feedforward neural networks},
  author={Glorot, Xavier and Bengio, Yoshua},
  booktitle={Proceedings of the {T}hirteenth {I}nternational {C}onference on {A}rtificial {I}ntelligence and {S}tatistics},
  pages={249--256},
  year={2010}
}

@article{kom1997time,
  title={Time-to-event analysis of longitudinal follow-up of a survey: choice of the time-scale},
  author={Kom, Edward L and Graubard, Barry I and Midthune, Douglas},
  journal={American Journal of Epidemiology},
  volume={145},
  number={1},
  pages={72--80},
  year={1997},
  publisher={Oxford Univ Press}
}

@book{cox1984analysis,
  title={Analysis of survival data},
  author={Cox, David Roxbee and Oakes, David},
  volume={21},
  year={1984},
  publisher={CRC Press}
}

@article{cevenini2008human,
  title={Human models of aging and longevity},
  author={Cevenini, E and Invidia, L and Lescai, Francesco and Salvioli, S and Tieri, P and Castellani, G and Franceschi, C},
  journal={Expert opinion on biological therapy},
  volume={8},
  number={9},
  pages={1393--1405},
  year={2008},
  publisher={Taylor \& Francis}
}

@article{belsky2015quantification,
  title={Quantification of biological aging in young adults},
  author={Belsky, Daniel W and Caspi, Avshalom and Houts, Renate and Cohen, Harvey J and Corcoran, David L and Danese, Andrea and Harrington, HonaLee and Israel, Salomon and Levine, Morgan E and Schaefer, Jonathan D and others},
  journal={Proceedings of the National Academy of Sciences},
  volume={112},
  number={30},
  pages={E4104--E4110},
  year={2015},
  publisher={National Acad Sciences}
}
@article{mitnitski2016heterogeneity,
  title={Heterogeneity of human aging and its assessment},
  author={Mitnitski, Arnold and Howlett, Susan E and Rockwood, Kenneth},
  journal={Journals of Gerontology Series A: Biomedical Sciences and Medical Sciences},
  volume={72},
  number={7},
  pages={877--884},
  year={2016},
  publisher={Oxford University Press US}
}
@article{levine2012modeling,
  title={Modeling the rate of senescence: {C}an estimated biological age predict mortality more accurately than chronological age?},
  author={Levine, Morgan E},
  journal={Journals of Gerontology Series A: Biomedical Sciences and Medical Sciences},
  volume={68},
  number={6},
  pages={667--674},
  year={2012},
  publisher={Oxford University Press US}
}

@article{nakamura1988assessment,
  title={Assessment of biological age by principal component analysis},
  author={Nakamura, E and Miyao, K and Ozeki, T},
  journal={Mechanisms of Ageing and Development},
  volume={46},
  number={1},
  pages={1--18},
  year={1988},
  publisher={Elsevier}
}
@article{kroll2000use,
  title={On the use of regression analysis for the estimation of human biological age},
  author={Kr{\o}ll, J and Saxtrup, O},
  journal={Biogerontology},
  volume={1},
  number={4},
  pages={363--368},
  year={2000},
  publisher={Springer}
}

@article{takeda1982evaluation,
  title={Evaluation of biological age and physical age by multiple regression analysis},
  author={Takeda, Hiroshi and Inada, Hiroshi and Inoue, Michitoshi and Yoshikawa, Hiromichi and Abe, Hiroshi},
  journal={Medical Informatics},
  volume={7},
  number={3},
  pages={221--227},
  year={1982},
  publisher={Taylor \& Francis}
}
@article{xu2015novel,
  title={A novel strategy for forensic age prediction by {DNA} methylation and support vector regression model},
  author={Xu, Cheng and Qu, Hongzhu and Wang, Guangyu and Xie, Bingbing and Shi, Yi and Yang, Yaran and Zhao, Zhao and Hu, Lan and Fang, Xiangdong and Yan, Jiangwei and others},
  journal={Scientific Reports},
  volume={5},
  year={2015},
  publisher={Nature Publishing Group}
}

@inproceedings{guo2009human,
  title={Human age estimation using bio-inspired features},
  author={Guo, Guodong and Mu, Guowang and Fu, Yun and Huang, Thomas S},
  booktitle={IEEE Conference on Computer Vision and Pattern Recognition (CVPR) },
  pages={112--119},
  year={2009},
  organization={IEEE}
}

@article{fu2008human,
  title={Human age estimation with regression on discriminative aging manifold},
  author={Fu, Yun and Huang, Thomas S},
  journal={IEEE Transactions on Multimedia},
  volume={10},
  number={4},
  pages={578--584},
  year={2008},
  publisher={IEEE}
}

@article{horvath2013dna,
  title={{DNA} methylation age of human tissues and cell types},
  author={Horvath, Steve},
  journal={Genome Biology},
  volume={14},
  number={10},
  pages={3156},
  year={2013},
  publisher={BioMed Central}
}

@article{cole2017brain,
  title={Brain age predicts mortality},
  author={Cole, JH and Ritchie, SJ and Bastin, ME and Hern{\'a}ndez MC Vald{\'e}s and Maniega, S Mu{\~n}oz and Royle, N and Corley, J and Pattie, A and Harris, SE and Zhang, Q and others},
  journal={Molecular Psychiatry},
  year={2017},
  publisher={Nature Publishing Group}
}

@article{sebastiani2017biomarker,
  title={Biomarker signatures of aging},
  author={Sebastiani, Paola and Thyagarajan, Bharat and Sun, Fangui and Schupf, Nicole and Newman, Anne B and Montano, Monty and Perls, Thomas T},
  journal={Aging Cell},
  volume={16},
  number={2},
  pages={329--338},
  year={2017},
  publisher={Wiley Online Library}
}

@article{piccirilli2017framework,
  title={{A} {F}ramework for {A}nalyzing the {W}hole {B}ody {S}urface {A}rea from a {S}ingle {V}iew},
  author={Piccirilli, Marco and Doretto, Gianfranco and Adjeroh, Donald},
  journal={PLoS ONE},
  volume={12},
  number={1},
  pages={e0166749},
  year={2017},
  publisher={Public Library of Science}
}

@article{shamir2011automatic,
  title={Automatic age estimation by hand photos},
  author={Shamir, Lior},
  journal={Computer Science Letters},
  volume={3},
  number={1},
  year={2011}
}

@article{aoyagi2001comparison,
  title={Comparison of performance-based measures among native {J}apanese, {J}apanese-{A}mericans in {H}awaii and {C}aucasian women in the {U}nited {S}tates, ages 65 years and over: {A} cross-sectional study},
  author={Aoyagi, Kiyoshi and Ross, Philip D and Nevitt, Michael C and Davis, James W and Wasnich, Richard D and Hayashi, Takuo and Takemoto, Tai-ichiro},
  journal={BMC Geriatrics},
  volume={1},
  number={1},
  pages={1},
  year={2001},
  publisher={BioMed Central}
}

@article{chilima2001nutrition,
  title={Nutrition and handgrip strength of older adults in rural {M}alawi},
  author={Chilima, Dorothy M and Ismail, Suraiya J},
  journal={Public Health Nutrition},
  volume={4},
  number={01},
  pages={11--17},
  year={2001},
  publisher={Cambridge Univ Press}
}

@article{vaz2002prediction,
  title={Prediction equations for handgrip strength in healthy {I}ndian male and female subjects encompassing a wide age range},
  author={Vaz, M and Hunsberger, S and Diffey, B},
  journal={Annals of Human Biology},
  volume={29},
  number={2},
  pages={131--141},
  year={2002},
  publisher={Taylor \& Francis}
}

@article{koley2009study,
  title={A study on hand grip strength in female labourers of {J}alandhar, {P}unjab, {I}ndia},
  author={Koley, Shyamal and Kaur, Navdeep and Sandhu, JS},
  journal={J. Life Sci},
  volume={1},
  number={1},
  pages={57--62},
  year={2009}
}

@inproceedings{scholkopf1997kernel,
  title={{K}ernel {P}rincipal {C}omponent {A}nalysis},
  author={Sch{\"o}lkopf, Bernhard and Smola, Alexander and M{\"u}ller, Klaus-Robert},
  booktitle={International Conference on Artificial Neural Networks},
  pages={583--588},
  year={1997},
  organization={Springer}
}

@book{jolliffe2002principal,
  title={{P}rincipal {C}omponent {A}nalysis},
  author={Jolliffe, Ian},
  year={2002},
  publisher={Wiley Online Library}
}

@article{al2002handgrip,
  title={Handgrip strength and mortality in older {M}exican {A}mericans},
  author={Al Snih, Soham and Markides, Kyriakos S and Ray, Laura and Ostir, Glenn V and Goodwin, James S},
  journal={Journal of the American Geriatrics Society},
  volume={50},
  number={7},
  pages={1250--1256},
  year={2002},
  publisher={Wiley Online Library}
}

@article{ali2008acquired,
  title={Acquired weakness, handgrip strength, and mortality in critically ill patients},
  author={Ali, Naeem A and O'Brien Jr, James M and Hoffmann, Stephen P and Phillips, Gary and Garland, Allan and Finley, James CW and Almoosa, Khalid and Hejal, Rana and Wolf, Karen M and Lemeshow, Stanley and others},
  journal={American Journal of Respiratory and Critical Care Medicine},
  volume={178},
  number={3},
  pages={261--268},
  year={2008},
  publisher={Am Thoracic Soc}
}

@article{ling2010handgrip,
  title={Handgrip strength and mortality in the oldest old population: {T}he {L}eiden 85-plus study},
  author={Ling, Carolina HY and Taekema, Diana and de Craen, Anton JM and Gussekloo, Jacobijn and Westendorp, Rudi GJ and Maier, Andrea B},
  journal={Canadian Medical Association Journal},
  volume={182},
  number={5},
  pages={429--435},
  year={2010},
  publisher={Can Med Assoc}
}

@article{bocklandt2011epigenetic,
  title={Epigenetic predictor of age},
  author={Bocklandt, Sven and Lin, Wen and Sehl, Mary E and S{\'a}nchez, Francisco J and Sinsheimer, Janet S and Horvath, Steve and Vilain, Eric},
  journal={PLoS ONE},
  volume={6},
  number={6},
  pages={e14821},
  year={2011},
  publisher={Public Library of Science}
}

@inproceedings{ramanathan2006modeling,
  title={Modeling age progression in young faces},
  author={Ramanathan, Narayanan and Chellappa, Rama},
  booktitle={2006 IEEE Computer Society Conference on Computer Vision and Pattern Recognition (CVPR'06)},
  volume={1},
  pages={387--394},
  year={2006},
  organization={IEEE}
}

@article{campisi2003cellular,
  title={Cellular senescence and cell death},
  author={Campisi, Judith},
  journal={Physiological Basis of Aging and Geriatrics},
  pages={47--59},
  year={2003}
}

@article{campisi1997biology,
  title={The biology of replicative senescence},
  author={Campisi, J},
  journal={European Journal of Cancer},
  volume={33},
  number={5},
  pages={703--709},
  year={1997},
  publisher={Elsevier}
}

@article{richter1988normal,
  title={Normal oxidative damage to mitochondrial and nuclear {DNA} is extensive},
  author={Richter, Christoph and Park, Jeen-Woo and Ames, Bruce N},
  journal={Proceedings of the National Academy of Sciences},
  volume={85},
  number={17},
  pages={6465--6467},
  year={1988},
  publisher={National Acad Sciences}
}

@article{harman2003free,
  title={The free radical theory of aging},
  author={Harman, Denham},
  journal={Antioxidants and Redox Signaling},
  volume={5},
  number={5},
  pages={557--561},
  year={2003},
  publisher={Mary Ann Liebert, Inc.}
}

@article{tosato2007aging,
  title={The aging process and potential interventions to extend life expectancy},
  author={Tosato, Matteo and Zamboni, Valentina and Ferrini, Alessandro and Cesari, Matteo},
  journal={Clinical Interventions in Aging},
  volume={2},
  number={3},
  pages={401},
  year={2007},
  publisher={DOVE MEDICAL PRESS}
}

@article{fischer2014biomarker,
  title={Biomarker profiling by nuclear magnetic resonance spectroscopy for the prediction of all-cause mortality: {A}n observational study of 17,345 persons},
  author={Fischer, Krista and Kettunen, Johannes and W{\"u}rtz, Peter and Haller, Toomas and Havulinna, Aki S and Kangas, Antti J and Soininen, Pasi and Esko, T{\~o}nu and Tammesoo, Mari-Liis and M{\"a}gi, Reedik and others},
  journal={PLoS Med},
  volume={11},
  number={2},
  pages={e1001606},
  year={2014},
  publisher={Public Library of Science}
}

@article{ritz2000age,
  title={Age estimation: {T}he state of the art in relation to the specific demands of forensic practise},
  author={Ritz-Timme, S and Cattaneo, C and Collins, MJ and Waite, ER and Sch{\"u}tz, HW and Kaatsch, H-J and Borrman, HIM},
  journal={International Journal of Legal Medicine},
  volume={113},
  number={3},
  pages={129--136},
  year={2000},
  publisher={Springer}
}

@article{anstey1996measuring,
  title={Measuring human functional age: {A} review of empirical findings},
  author={Anstey, Kaarin J and Lord, Stephen R and Smith, Glen A},
  journal={Experimental Aging Research},
  volume={22},
  number={3},
  pages={245--266},
  year={1996},
  publisher={Taylor \& Francis}
}

@article{kroger1993development,
  title={Development of bone mass and bone density of the spine and femoral neck—a prospective study of 65 children and adolescents},
  author={Kr{\"o}ger, Heikki and Kotaniemi, Antero and Kr{\"o}ger, Liisa and Alhava, Esko},
  journal={Bone and Mineral},
  volume={23},
  number={3},
  pages={171--182},
  year={1993},
  publisher={Elsevier}
}

@article{krakauer2012new,
  title={A new body shape index predicts mortality hazard independently of {B}ody {M}ass {I}ndex},
  author={Krakauer, Nir Y and Krakauer, Jesse C},
  journal={PLoS ONE},
  volume={7},
  number={7},
  pages={e39504},
  year={2012},
  publisher={Public Library of Science}
}

@article{geng2007automatic,
  title={Automatic age estimation based on facial aging patterns},
  author={Geng, Xin and Zhou, Zhi-Hua and Smith-Miles, Kate},
  journal={IEEE Transactions on Pattern Analysis and Machine Intelligence},
  volume={29},
  number={12},
  pages={2234--2240},
  year={2007},
  publisher={IEEE}
}

@article{williams2001review,
  title={A review of the most commonly used dental age estimation techniques},
  author={Williams, Guy},
  journal={J Forensic Odontostomatol},
  volume={19},
  pages={9--17},
  year={2001}
}

@article{fu2010age,
  title={Age synthesis and estimation via faces: {A} survey},
  author={Fu, Yun and Guo, Guodong and Huang, Thomas S},
  journal={IEEE Transactions on Pattern Analysis and Machine Intelligence},
  volume={32},
  number={11},
  pages={1955--1976},
  year={2010},
  publisher={IEEE}
}

@article{han2015demographic,
  title={Demographic estimation from face images: Human vs. machine performance},
  author={Han, Hu and Otto, Charles and Liu, Xiaoming and Jain, Anil K},
   journal={IEEE Transactions on Pattern Analysis and Machine Intelligence},
  volume={37},
  number={6},
  pages={1148--1161},
  year={2015},
  publisher={IEEE}
}

@article{chua1990antecedents,
  title={The antecedents of cognitive age},
  author={Chua, Caroline and Cote, Joseph A and Leong, Siew Meng},
  journal={NA-Advances in Consumer Research Volume 17},
  year={1990}
}

@article{house1994social,
  title={The social stratification of aging and health},
  author={House, James S and Lepkowski, James M and Kinney, Ann M and Mero, Richard P and Kessler, Ronald C and Herzog, A Regula},
  journal={Journal of Health and Social Behavior},
  pages={213--234},
  year={1994},
  publisher={JSTOR}
}

@article{schmeling2008criteria,
  title={Criteria for age estimation in living individuals},
  author={Schmeling, A and Grundmann, C and Fuhrmann, A and Kaatsch, H-J and Knell, B and Ramsthaler, F and Reisinger, W and Riepert, T and Ritz-Timme, S and R{\"o}sing, FW and others},
  journal={International Journal of Legal Medicine},
  volume={122},
  number={6},
  pages={457--460},
  year={2008},
  publisher={Springer}
}

@article{maddox1973self,
  title={Self-assessment of health: {A} longitudinal study of elderly subjects},
  author={Maddox, George L and Douglass, Elizabeth B},
  journal={Journal of Health and Social Behavior},
  pages={87--93},
  year={1973},
  publisher={JSTOR}
}

@article{nurmi1987age,
  title={Age, sex, social class, and quality of family interaction as determinants of adolescents' future orientation: {A} developmental task interpretation},
  author={Nurmi, Jari-Erik},
  journal={Adolescence},
  volume={22},
  number={88},
  pages={977},
  year={1987},
  publisher={Libra Publishers.}
}

@article{rahman2015surface,
  title={{S}urface-{B}ased {B}ody {S}hape {I}ndex and {I}ts {R}elationship with {A}ll-{C}ause {M}ortality},
  author={Rahman, Syed Ashiqur and Adjeroh, Donald},
  journal={PLoS ONE},
  volume={10},
  number={12},
  pages={e0144639},
  year={2015},
  publisher={Public Library of Science}
}

@article{sanderson2014measuring,
  title={Measuring the speed of aging across population subgroups},
  author={Sanderson, Warren C and Scherbov, Sergei},
  journal={PLoS ONE},
  volume={9},
  number={5},
  pages={e96289},
  year={2014},
  publisher={Public Library of Science}
}
@article{massy2011hand,
  title={Hand Grip Strength: {A}ge and gender stratified normative data in a population-based study},
  author={Massy-Westropp, Nicola M and Gill, Tiffany K and Taylor, Anne W and Bohannon, Richard W and Hill, Catherine L},
  journal={BMC Research Notes},
  volume={4},
  number={1},
  pages={127},
  year={2011},
  publisher={BioMed Central Ltd}
}

@inproceedings{xia2014can,
  title={Can 3{D} shape of the face reveal your age?},
  author={Xia, Baiqiang and Amor, Boulbaba Ben and Daoudi, Mohamed and Drira, Hassen},
  booktitle={International Conference on Computer Vision Theory and Applications (VISAPP) },
  volume={2},
  pages={5--13},
  year={2014},
  organization={IEEE}
}

@inproceedings{guo2009study,
  title={A study on automatic age estimation using a large database.},
  author={Guo, Guodong and Mu, Guowang and Fu, Yun and Dyer, Charles R and Huang, Thomas S},
  booktitle={ICCV},
  pages={1986--1991},
  year={2009}
}

@inproceedings{egilmez2014spectral,
  title={Spectral anomaly detection using graph-based filtering for wireless sensor networks},
  author={Egilmez, Hilmi E and Ortega, Antonio},
  booktitle={IEEE International Conference on Acoustics, Speech and Signal Processing (ICASSP), 2014 },
  pages={1085--1089},
  year={2014},
  organization={IEEE}
}

@article{belkin2003laplacian,
  title={Laplacian eigenmaps for dimensionality reduction and data representation},
  author={Belkin, Mikhail and Niyogi, Partha},
  journal={Neural Computation},
  volume={15},
  number={6},
  pages={1373--1396},
  year={2003},
  publisher={MIT Press}
}

@article{klemera2006new,
  title={A new approach to the concept and computation of biological age},
  author={Klemera, Petr and Doubal, Stanislav},
  journal={Mechanisms of Ageing and Development},
  volume={127},
  number={3},
  pages={240--248},
  year={2006},
  publisher={Elsevier}
}

@article{jackson2003biological,
  title={Biological age—what is it and can it be measured?},
  author={Jackson, Stephen HD and Weale, Martin R and Weale, Robert A},
  journal={Archives of Gerontology and Geriatrics},
  volume={36},
  number={2},
  pages={103--115},
  year={2003},
  publisher={Elsevier}
}

@inproceedings{han2013age,
  title={Age estimation from face images: {H}uman vs. machine performance},
  author={Han, Hu and Otto, Christina and Jain, Anubhav K},
  booktitle={Biometrics (ICB), 2013 International Conference on},
  pages={1--8},
  year={2013},
  organization={IEEE}
}

@article{eikvil2012age,
  title={Age estimation in youths and young adults},
  author={Eikvil, Line and Kvaal, Sigrid Ingeborg and Teigland, Andr{\'e} and Haugen, Marion and Gr{\o}gaard, Jens},
  journal={Norsk Regnesentral},
  volume={12},
  year={2012},
  publisher={Citeseer}
}

@article{cho2010empirical,
  title={An empirical comparative study on biological age estimation algorithms with an application of {W}ork {A}bility {I}ndex ({WAI})},
  author={Cho, Il Haeng and Park, Kyung S and Lim, Chang Joo},
  journal={Mechanisms of Ageing and Development},
  volume={131},
  number={2},
  pages={69--78},
  year={2010},
  publisher={Elsevier}
}

@article{putin2016deep,
  title={Deep biomarkers of human aging: {A}pplication of deep neural networks to biomarker development},
  author={Putin, Evgeny and Mamoshina, Polina and Aliper, Alexander and Korzinkin, Mikhail and Moskalev, Alexey and Kolosov, Alexey and Ostrovskiy, Alexander and Cantor, Charles and Vijg, Jan and Zhavoronkov, Alex},
  journal={Aging},
  volume={8},
  number={5},
  pages={1--021},
  year={2016}
}

@article{rhodes2009age,
  title={Age estimation of faces: {A} review},
  author={Rhodes, Matthew G},
  journal={Applied Cognitive Psychology},
  volume={23},
  number={1},
  pages={1--12},
  year={2009},
  publisher={Wiley Online Library}
}

@book{alley2013social,
  title={{S}ocial and {A}pplied {A}spects of {P}erceiving {F}aces},
  author={Alley, Thomas R},
  year={2013},
  publisher={Psychology Press}
}

@ONLINE{cdc2011Online,
  author = {},
  title = {IEEE Conference on Decision and Control and European Control Conference {@ONLINE}},
  month = {December},
  year = {2011},
  url = { http://control.disp.uniroma2.it/cdcecc2011/; http://www.cdc.gov/BRFSS/},
  urldate = {2014-10-10}
}

@ONLINE{nhanes2002Online,
  author = {},
  title = { {N}ational {H}ealth and {N}utrition {E}xamination {S}urvey ({NHANES}) {A}nthropometry {P}rocedures {M}anual. {T}echnical report, {N}ational {C}enter for {H}ealth {S}tatistics {@ONLINE}},
  month = {December},
  year = {2002},
  url = {http://www.cdc.gov/nchs/data/nhanes/},
  urldate = {2014-10-10}
}

@article{mughal2014bone,
  title={Bone age assessment methods: {A} critical review},
  author={Mughal, Arsalan Manzoor and Hassan, Nuzhat and Ahmed, Anwar},
  journal={Pakistan journal of medical sciences},
  volume={30},
  number={1},
  pages={211},
  year={2014},
  publisher={Professional Medical Publications}
}

@article{zerilli1992simple,
  title={A simple technique for age estimation in adult corpses: {T}he two criteria dental method},
  author={Zerilli, A and Baccino, E and Lamendin, H and Tavernier, JC and Humbert, JF and Nossintchouk, RM},
  journal={Journal of Forensic Science},
  volume={37},
  number={5},
  pages={1373--1379},
  year={1992},
  publisher={ASTM International}
}

@article{basak2007support,
  title={{S}upport {V}ector {R}egression},
  author={Basak, Debasish and Pal, Srimanta and Patranabis, Dipak Chandra},
  journal={Neural Information Processing-Letters and Reviews},
  volume={11},
  number={10},
  pages={203--224},
  year={2007},
  publisher={Citeseer}
}

@article{liaw2002classification,
  title={Classification and {R}egression by random{F}orest},
  author={Liaw, Andy and Wiener, Matthew},
  journal={R news},
  volume={2},
  number={3},
  pages={18--22},
  year={2002}
}

@article{maber2006accuracy,
  title={Accuracy of age estimation of radiographic methods using developing teeth},
  author={Maber, M and Liversidge, HM and Hector, MP},
  journal={Forensic Science International},
  volume={159},
  pages={S68--S73},
  year={2006},
  publisher={Elsevier}
}

@article{lu2010gait,
  title={Gait-based human age estimation},
  author={Lu, Jiwen and Tan, Yap-Peng},
  journal={IEEE Transactions on Information Forensics and Security},
  volume={5},
  number={4},
  pages={761--770},
  year={2010},
  publisher={IEEE}
}

@inproceedings{wang2015deeply,
  title={Deeply-learned feature for age estimation},
  author={Wang, Xiaolong and Guo, Rui and Kambhamettu, Chandra},
  booktitle={2015 IEEE Winter Conference on Applications of Computer Vision},
  pages={534--541},
  year={2015},
  organization={IEEE}
}

@article{guo2008locally,
  title={Locally adjusted robust regression for human age estimation},
  author={Guo, Guodong and Fu, Yun and Huang, Thomas S and Dyer, Charles R},
  journal={IEEE Workshop on Applications of Computer Vision},
  volume={51},
  pages={61801},
  year={2008}
}

@article{eipel2016epigenetic,
  title={Epigenetic age predictions based on buccal swabs are more precise in combination with cell type-specific {DNA} methylation signatures},
  author={Eipel, Monika and Mayer, Felix and Arent, Tanja and Ferreira, Marcelo RP and Birkhofer, Carina and Gerstenmaier, Uwe and Costa, Ivan G and Ritz-Timme, Stefanie and Wagner, Wolfgang},
  journal={Aging},
  volume={8},
  number={5},
  year={2016}
}

@inproceedings{guo2011simultaneous,
  title={Simultaneous dimensionality reduction and human age estimation via kernel partial least squares regression},
  author={Guo, Guodong and Mu, Guowang},
  booktitle={Computer Vision and Pattern Recognition (CVPR), 2011 IEEE Conference on},
  pages={657--664},
  year={2011},
  organization={IEEE}
}

@inproceedings{ni2009web,
  title={Web image mining towards universal age estimator},
  author={Ni, Bingbing and Song, Zheng and Yan, Shuicheng},
  booktitle={Proceedings of the 17th ACM International Conference on Multimedia},
  pages={85--94},
  year={2009},
  organization={ACM}
}

@article{geng2013facial,
  title={Facial age estimation by learning from label distributions},
  author={Geng, Xin and Yin, Chao and Zhou, Zhi-Hua},
  journal={IEEE Transactions on Pattern Analysis and Machine Intelligence},
  volume={35},
  number={10},
  pages={2401--2412},
  year={2013},
  publisher={IEEE}
}

@inproceedings{robinette1999caesar,
  title={The CAESAR project: {A} 3-{D} surface anthropometry survey},
  author={Robinette, Kathleen M and Daanen, Hans and Paquet, Eric},
  booktitle={3-D Digital Imaging and Modeling, 1999. Proceedings. Second International Conference on},
  pages={380--386},
  year={1999},
  organization={IEEE}
}

% http://cyberware.com/products/scanners/
@ONLINE{cyberware:Online,
  author = {},
  title = {Cyberware scanner {@ONLINE}},
  month = {},
  year = {},
  url = {http://cyberware.com/products/scanners/},
  urldate = {2014-10-10}
}

@inproceedings{adjeroh2010predictability,
  title={Predictability and correlation in human metrology},
  author={Adjeroh, Donald and Cao, Deng and Piccirilli, Marco and Ross, Arun},
  booktitle={Information Forensics and Security (WIFS), 2010 IEEE International Workshop on},
  pages={1--6},
  year={2010},
  organization={IEEE}
}

@inproceedings{cao2012predicting,
  title={Predicting gender and weight from human metrology using a copula model},
  author={Cao, Deng and Chen, Cunjian and Adjeroh, Donald and Ross, Arun},
  booktitle={Biometrics: Theory, Applications and Systems (BTAS), 2012 IEEE Fifth International Conference on},
  pages={162--169},
  year={2012},
  organization={IEEE}
}

@article{box1964analysis,
  title={An analysis of transformations},
  author={Box, George EP and Cox, David R},
  journal={Journal of the Royal Statistical Society. Series B (Methodological)},
  pages={211--252},
  year={1964},
  publisher={JSTOR}
}

@inproceedings{xingjian2015convolutional,
  title={Convolutional LSTM network: A machine learning approach for precipitation nowcasting},
  author={Xingjian, SHI and Chen, Zhourong and Wang, Hao and Yeung, Dit-Yan and Wong, Wai-Kin and Woo, Wang-chun},
  booktitle={Advances in Neural Information Processing Systems},
  pages={802--810},
  year={2015}
}

@article{hochreiter1997long,
  title={Long short-term memory},
  author={Hochreiter, Sepp and Schmidhuber, J{\"u}rgen},
  journal={Neural Computation},
  volume={9},
  number={8},
  pages={1735--1780},
  year={1997},
  publisher={MIT Press}
}

@article{pyrkov2018extracting,
  title={Extracting biological age from biomedical data via deep learning: too much of a good thing?},
  author={Pyrkov, Timothy V and Slipensky, Konstantin and Barg, Mikhail and Kondrashin, Alexey and Zhurov, Boris and Zenin, Alexander and Pyatnitskiy, Mikhail and Menshikov, Leonid and Markov, Sergei and Fedichev, Peter O},
  journal={Scientific Reports},
  volume={8},
  number={1},
  pages={5210},
  year={2018},
  publisher={Nature Publishing Group}
}

@ONLINE{CDC2017adult_vaccination,
  author = {},
  title = {US Centers for Disease Control (CDC), Recommended Immunization Schedule for Adults Aged 19 Years or Older, by Vaccine and Age Group, United States {@ONLINE}},
  month = {December},
  year = {2017},
  url = { https://www.cdc.gov/vaccines/schedules/hcp/imz/adult.html},
  urldate = {2017-12-29}
}

@article{altmann2010permutation,
  title={Permutation importance: {A} corrected feature importance measure},
  author={Altmann, Andr{\'e} and Tolo{\c{s}}i, Laura and Sander, Oliver and Lengauer, Thomas},
  journal={Bioinformatics},
  volume={26},
  number={10},
  pages={1340--1347},
  year={2010},
  publisher={Oxford University Press}
}

@article{hamano2017forensic,
  title={Forensic age prediction for saliva samples using methylation-sensitive high resolution melting: exploratory application for cigarette butts},
  author={Hamano, Yuya and Manabe, Sho and Morimoto, Chie and Fujimoto, Shuntaro and Tamaki, Keiji},
  journal={Scientific {R}eports},
  volume={7},
  number={1},
  pages={10444},
  year={2017},
  publisher={Nature Publishing Group}
}

@article{xu2015novel,
  title={A novel strategy for forensic age prediction by {DNA} methylation and support vector regression model},
  author={Xu, Cheng and Qu, Hongzhu and Wang, Guangyu and Xie, Bingbing and Shi, Yi and Yang, Yaran and Zhao, Zhao and Hu, Lan and Fang, Xiangdong and Yan, Jiangwei and others},
  journal={Scientific {R}eports},
  volume={5},
  year={2015},
  publisher={Nature Publishing Group}
}


@article{piccirilli2017framework,
  title={{A} {F}ramework for {A}nalyzing the {W}hole {B}ody {S}urface {A}rea from a {S}ingle {V}iew},
  author={Piccirilli, Marco and Doretto, Gianfranco and Adjeroh, Donald},
  journal={PLoS ONE},
  volume={12},
  number={1},
  pages={e0166749},
  year={2017},
  publisher={Public Library of Science}
}

@article{shamir2011automatic,
  title={Automatic age estimation by hand photos},
  author={Shamir, Lior},
  journal={Computer Science Letters},
  volume={3},
  number={1},
  year={2011}
}

@article{aoyagi2001comparison,
  title={Comparison of performance-based measures among native {J}apanese, {J}apanese-{A}mericans in {H}awaii and {C}aucasian women in the {U}nited {S}tates, ages 65 years and over: {A} cross-sectional study},
  author={Aoyagi, Kiyoshi and Ross, Philip D and Nevitt, Michael C and Davis, James W and Wasnich, Richard D and Hayashi, Takuo and Takemoto, Tai-ichiro},
  journal={BMC Geriatrics},
  volume={1},
  number={1},
  pages={1},
  year={2001},
  publisher={BioMed Central}
}

@article{chilima2001nutrition,
  title={Nutrition and handgrip strength of older adults in rural {M}alawi},
  author={Chilima, Dorothy M and Ismail, Suraiya J},
  journal={Public Health Nutrition},
  volume={4},
  number={01},
  pages={11--17},
  year={2001},
  publisher={Cambridge Univ Press}
}

@article{vaz2002prediction,
  title={Prediction equations for handgrip strength in healthy {I}ndian male and female subjects encompassing a wide age range},
  author={Vaz, M and Hunsberger, S and Diffey, B},
  journal={Annals of Human Biology},
  volume={29},
  number={2},
  pages={131--141},
  year={2002},
  publisher={Taylor \& Francis}
}

@article{koley2009study,
  title={A study on hand grip strength in female labourers of {J}alandhar, {P}unjab, {I}ndia},
  author={Koley, Shyamal and Kaur, Navdeep and Sandhu, JS},
  journal={J. Life Sci},
  volume={1},
  number={1},
  pages={57--62},
  year={2009}
}

@inproceedings{scholkopf1997kernel,
  title={{K}ernel {P}rincipal {C}omponent {A}nalysis},
  author={Sch{\"o}lkopf, Bernhard and Smola, Alexander and M{\"u}ller, Klaus-Robert},
  booktitle={International Conference on Artificial Neural Networks},
  pages={583--588},
  year={1997},
  organization={Springer}
}

@book{jolliffe2002principal,
  title={{P}rincipal {C}omponent {A}nalysis},
  author={Jolliffe, Ian},
  year={2002},
  publisher={Wiley Online Library}
}

@article{al2002handgrip,
  title={Handgrip strength and mortality in older {M}exican {A}mericans},
  author={Al Snih, Soham and Markides, Kyriakos S and Ray, Laura and Ostir, Glenn V and Goodwin, James S},
  journal={Journal of the American Geriatrics Society},
  volume={50},
  number={7},
  pages={1250--1256},
  year={2002},
  publisher={Wiley Online Library}
}

@article{ali2008acquired,
  title={Acquired weakness, handgrip strength, and mortality in critically ill patients},
  author={Ali, Naeem A and O'Brien Jr, James M and Hoffmann, Stephen P and Phillips, Gary and Garland, Allan and Finley, James CW and Almoosa, Khalid and Hejal, Rana and Wolf, Karen M and Lemeshow, Stanley and others},
  journal={American Journal of Respiratory and Critical Care Medicine},
  volume={178},
  number={3},
  pages={261--268},
  year={2008},
  publisher={Am Thoracic Soc}
}

@article{ling2010handgrip,
  title={Handgrip strength and mortality in the oldest old population: {T}he {L}eiden 85-plus study},
  author={Ling, Carolina HY and Taekema, Diana and de Craen, Anton JM and Gussekloo, Jacobijn and Westendorp, Rudi GJ and Maier, Andrea B},
  journal={Canadian Medical Association Journal},
  volume={182},
  number={5},
  pages={429--435},
  year={2010},
  publisher={Can Med Assoc}
}

@article{bocklandt2011epigenetic,
  title={Epigenetic predictor of age},
  author={Bocklandt, Sven and Lin, Wen and Sehl, Mary E and S{\'a}nchez, Francisco J and Sinsheimer, Janet S and Horvath, Steve and Vilain, Eric},
  journal={PLoS ONE},
  volume={6},
  number={6},
  pages={e14821},
  year={2011},
  publisher={Public Library of Science}
}

@inproceedings{ramanathan2006modeling,
  title={Modeling age progression in young faces},
  author={Ramanathan, Narayanan and Chellappa, Rama},
  booktitle={2006 IEEE Computer Society Conference on Computer Vision and Pattern Recognition (CVPR'06)},
  volume={1},
  pages={387--394},
  year={2006},
  organization={IEEE}
}

@article{campisi2003cellular,
  title={Cellular senescence and cell death},
  author={Campisi, Judith},
  journal={Physiological Basis of Aging and Geriatrics},
  pages={47--59},
  year={2003}
}

@article{campisi1997biology,
  title={The biology of replicative senescence},
  author={Campisi, J},
  journal={European Journal of Cancer},
  volume={33},
  number={5},
  pages={703--709},
  year={1997},
  publisher={Elsevier}
}

@article{richter1988normal,
  title={Normal oxidative damage to mitochondrial and nuclear {DNA} is extensive},
  author={Richter, Christoph and Park, Jeen-Woo and Ames, Bruce N},
  journal={Proceedings of the National Academy of Sciences},
  volume={85},
  number={17},
  pages={6465--6467},
  year={1988},
  publisher={National Acad Sciences}
}

@article{harman2003free,
  title={The free radical theory of aging},
  author={Harman, Denham},
  journal={Antioxidants and Redox Signaling},
  volume={5},
  number={5},
  pages={557--561},
  year={2003},
  publisher={Mary Ann Liebert, Inc.}
}

@article{tosato2007aging,
  title={The aging process and potential interventions to extend life expectancy},
  author={Tosato, Matteo and Zamboni, Valentina and Ferrini, Alessandro and Cesari, Matteo},
  journal={Clinical Interventions in Aging},
  volume={2},
  number={3},
  pages={401},
  year={2007},
  publisher={DOVE MEDICAL PRESS}
}

@article{ritz2000age,
  title={Age estimation: {T}he state of the art in relation to the specific demands of forensic practise},
  author={Ritz-Timme, S and Cattaneo, C and Collins, MJ and Waite, ER and Sch{\"u}tz, HW and Kaatsch, H-J and Borrman, HIM},
  journal={International Journal of Legal Medicine},
  volume={113},
  number={3},
  pages={129--136},
  year={2000},
  publisher={Springer}
}

@article{kroger1993development,
  title={Development of bone mass and bone density of the spine and femoral neck—a prospective study of 65 children and adolescents},
  author={Kr{\"o}ger, Heikki and Kotaniemi, Antero and Kr{\"o}ger, Liisa and Alhava, Esko},
  journal={Bone and Mineral},
  volume={23},
  number={3},
  pages={171--182},
  year={1993},
  publisher={Elsevier}
}

@article{geng2007automatic,
  title={Automatic age estimation based on facial aging patterns},
  author={Geng, Xin and Zhou, Zhi-Hua and Smith-Miles, Kate},
  journal={IEEE Transactions on Pattern Analysis and Machine Intelligence},
  volume={29},
  number={12},
  pages={2234--2240},
  year={2007},
  publisher={IEEE}
}

@article{williams2001review,
  title={A review of the most commonly used dental age estimation techniques},
  author={Williams, Guy},
  journal={J Forensic Odontostomatol},
  volume={19},
  pages={9--17},
  year={2001}
}

@article{fu2010age,
  title={Age synthesis and estimation via faces: {A} survey},
  author={Fu, Yun and Guo, Guodong and Huang, Thomas S},
  journal={IEEE Transactions on Pattern Analysis and Machine Intelligence},
  volume={32},
  number={11},
  pages={1955--1976},
  year={2010},
  publisher={IEEE}
}


@article{chua1990antecedents,
  title={The antecedents of cognitive age},
  author={Chua, Caroline and Cote, Joseph A and Leong, Siew Meng},
  journal={NA-Advances in Consumer Research Volume 17},
  year={1990}
}

@article{house1994social,
  title={The social stratification of aging and health},
  author={House, James S and Lepkowski, James M and Kinney, Ann M and Mero, Richard P and Kessler, Ronald C and Herzog, A Regula},
  journal={Journal of Health and Social Behavior},
  pages={213--234},
  year={1994},
  publisher={JSTOR}
}

@article{schmeling2008criteria,
  title={Criteria for age estimation in living individuals},
  author={Schmeling, A and Grundmann, C and Fuhrmann, A and Kaatsch, H-J and Knell, B and Ramsthaler, F and Reisinger, W and Riepert, T and Ritz-Timme, S and R{\"o}sing, FW and others},
  journal={International Journal of Legal Medicine},
  volume={122},
  number={6},
  pages={457--460},
  year={2008},
  publisher={Springer}
}

@article{maddox1973self,
  title={Self-assessment of health: {A} longitudinal study of elderly subjects},
  author={Maddox, George L and Douglass, Elizabeth B},
  journal={Journal of Health and Social Behavior},
  pages={87--93},
  year={1973},
  publisher={JSTOR}
}

@article{nurmi1987age,
  title={Age, sex, social class, and quality of family interaction as determinants of adolescents' future orientation: {A} developmental task interpretation},
  author={Nurmi, Jari-Erik},
  journal={Adolescence},
  volume={22},
  number={88},
  pages={977},
  year={1987},
  publisher={Libra Publishers.}
}

@article{sanderson2014measuring,
  title={Measuring the speed of aging across population subgroups},
  author={Sanderson, Warren C and Scherbov, Sergei},
  journal={PLoS ONE},
  volume={9},
  number={5},
  pages={e96289},
  year={2014},
  publisher={Public Library of Science}
}
@article{massy2011hand,
  title={Hand Grip Strength: {A}ge and gender stratified normative data in a population-based study},
  author={Massy-Westropp, Nicola M and Gill, Tiffany K and Taylor, Anne W and Bohannon, Richard W and Hill, Catherine L},
  journal={BMC Research Notes},
  volume={4},
  number={1},
  pages={127},
  year={2011},
  publisher={BioMed Central Ltd}
}

@inproceedings{xia2014can,
  title={Can 3{D} shape of the face reveal your age?},
  author={Xia, Baiqiang and Amor, Boulbaba Ben and Daoudi, Mohamed and Drira, Hassen},
  booktitle={International Conference on Computer Vision Theory and Applications (VISAPP) },
  volume={2},
  pages={5--13},
  year={2014},
  organization={IEEE}
}

@inproceedings{guo2009study,
  title={A study on automatic age estimation using a large database.},
  author={Guo, Guodong and Mu, Guowang and Fu, Yun and Dyer, Charles R and Huang, Thomas S},
  booktitle={ICCV},
  pages={1986--1991},
  year={2009}
}

@inproceedings{egilmez2014spectral,
  title={Spectral anomaly detection using graph-based filtering for wireless sensor networks},
  author={Egilmez, Hilmi E and Ortega, Antonio},
  booktitle={IEEE International Conference on Acoustics, Speech and Signal Processing (ICASSP), 2014 },
  pages={1085--1089},
  year={2014},
  organization={IEEE}
}

@article{belkin2003laplacian,
  title={Laplacian eigenmaps for dimensionality reduction and data representation},
  author={Belkin, Mikhail and Niyogi, Partha},
  journal={Neural Computation},
  volume={15},
  number={6},
  pages={1373--1396},
  year={2003},
  publisher={MIT Press}
}

@inproceedings{han2013age,
  title={Age estimation from face images: {H}uman vs. machine performance},
  author={Han, Hu and Otto, Christina and Jain, Anubhav K},
  booktitle={Biometrics (ICB), 2013 International Conference on},
  pages={1--8},
  year={2013},
  organization={IEEE}
}

@article{eikvil2012age,
  title={Age estimation in youths and young adults},
  author={Eikvil, Line and Kvaal, Sigrid Ingeborg and Teigland, Andr{\'e} and Haugen, Marion and Gr{\o}gaard, Jens},
  journal={Norsk Regnesentral},
  volume={12},
  year={2012},
  publisher={Citeseer}
}

@article{rhodes2009age,
  title={Age estimation of faces: {A} review},
  author={Rhodes, Matthew G},
  journal={Applied Cognitive Psychology},
  volume={23},
  number={1},
  pages={1--12},
  year={2009},
  publisher={Wiley Online Library}
}

@book{alley2013social,
  title={{S}ocial and {A}pplied {A}spects of {P}erceiving {F}aces},
  author={Alley, Thomas R},
  year={2013},
  publisher={Psychology Press}
}

@ONLINE{cdc2011Online,
  author = {},
  title = {IEEE Conference on Decision and Control and European Control Conference {@ONLINE}},
  month = {December},
  year = {2011},
  url = { http://control.disp.uniroma2.it/cdcecc2011/; http://www.cdc.gov/BRFSS/},
  urldate = {2014-10-10}
}

@ONLINE{nhanes2002Online,
  author = {},
  title = { {N}ational {H}ealth and {N}utrition {E}xamination {S}urvey ({NHANES}) {A}nthropometry {P}rocedures {M}anual. {T}echnical report, {N}ational {C}enter for {H}ealth {S}tatistics {@ONLINE}},
  month = {December},
  year = {2002},
  url = {http://www.cdc.gov/nchs/data/nhanes/},
  urldate = {2014-10-10}
}

@article{mughal2014bone,
  title={Bone age assessment methods: {A} critical review},
  author={Mughal, Arsalan Manzoor and Hassan, Nuzhat and Ahmed, Anwar},
  journal={Pakistan journal of medical sciences},
  volume={30},
  number={1},
  pages={211},
  year={2014},
  publisher={Professional Medical Publications}
}

@article{zerilli1992simple,
  title={A simple technique for age estimation in adult corpses: {T}he two criteria dental method},
  author={Zerilli, A and Baccino, E and Lamendin, H and Tavernier, JC and Humbert, JF and Nossintchouk, RM},
  journal={Journal of Forensic Science},
  volume={37},
  number={5},
  pages={1373--1379},
  year={1992},
  publisher={ASTM International}
}

@article{basak2007support,
  title={{S}upport {V}ector {R}egression},
  author={Basak, Debasish and Pal, Srimanta and Patranabis, Dipak Chandra},
  journal={Neural Information Processing-Letters and Reviews},
  volume={11},
  number={10},
  pages={203--224},
  year={2007},
  publisher={Citeseer}
}

@article{liaw2002classification,
  title={Classification and {R}egression by random{F}orest},
  author={Liaw, Andy and Wiener, Matthew},
  journal={R news},
  volume={2},
  number={3},
  pages={18--22},
  year={2002}
}

@article{maber2006accuracy,
  title={Accuracy of age estimation of radiographic methods using developing teeth},
  author={Maber, M and Liversidge, HM and Hector, MP},
  journal={Forensic Science International},
  volume={159},
  pages={S68--S73},
  year={2006},
  publisher={Elsevier}
}

@article{lu2010gait,
  title={Gait-based human age estimation},
  author={Lu, Jiwen and Tan, Yap-Peng},
  journal={IEEE Transactions on Information Forensics and Security},
  volume={5},
  number={4},
  pages={761--770},
  year={2010},
  publisher={IEEE}
}

@inproceedings{wang2015deeply,
  title={Deeply-learned feature for age estimation},
  author={Wang, Xiaolong and Guo, Rui and Kambhamettu, Chandra},
  booktitle={2015 IEEE Winter Conference on Applications of Computer Vision},
  pages={534--541},
  year={2015},
  organization={IEEE}
}

@article{guo2008locally,
  title={Locally adjusted robust regression for human age estimation},
  author={Guo, Guodong and Fu, Yun and Huang, Thomas S and Dyer, Charles R},
  journal={IEEE Workshop on Applications of Computer Vision},
  volume={51},
  pages={61801},
  year={2008}
}

@article{eipel2016epigenetic,
  title={Epigenetic age predictions based on buccal swabs are more precise in combination with cell type-specific {DNA} methylation signatures},
  author={Eipel, Monika and Mayer, Felix and Arent, Tanja and Ferreira, Marcelo RP and Birkhofer, Carina and Gerstenmaier, Uwe and Costa, Ivan G and Ritz-Timme, Stefanie and Wagner, Wolfgang},
  journal={Aging},
  volume={8},
  number={5},
  year={2016}
}

@inproceedings{guo2011simultaneous,
  title={Simultaneous dimensionality reduction and human age estimation via kernel partial least squares regression},
  author={Guo, Guodong and Mu, Guowang},
  booktitle={Computer Vision and Pattern Recognition (CVPR), 2011 IEEE Conference on},
  pages={657--664},
  year={2011},
  organization={IEEE}
}

@inproceedings{ni2009web,
  title={Web image mining towards universal age estimator},
  author={Ni, Bingbing and Song, Zheng and Yan, Shuicheng},
  booktitle={Proceedings of the 17th ACM International Conference on Multimedia},
  pages={85--94},
  year={2009},
  organization={ACM}
}

@article{geng2013facial,
  title={Facial age estimation by learning from label distributions},
  author={Geng, Xin and Yin, Chao and Zhou, Zhi-Hua},
  journal={IEEE Transactions on Pattern Analysis and Machine Intelligence},
  volume={35},
  number={10},
  pages={2401--2412},
  year={2013},
  publisher={IEEE}
}

@inproceedings{robinette1999caesar,
  title={The CAESAR project: {A} 3-{D} surface anthropometry survey},
  author={Robinette, Kathleen M and Daanen, Hans and Paquet, Eric},
  booktitle={3-D Digital Imaging and Modeling, 1999. Proceedings. Second International Conference on},
  pages={380--386},
  year={1999},
  organization={IEEE}
}

% http://cyberware.com/products/scanners/
@ONLINE{cyberware:Online,
  author = {},
  title = {Cyberware scanner {@ONLINE}},
  month = {},
  year = {},
  url = {http://cyberware.com/products/scanners/},
  urldate = {2014-10-10}
}

@inproceedings{adjeroh2010predictability,
  title={Predictability and correlation in human metrology},
  author={Adjeroh, Donald and Cao, Deng and Piccirilli, Marco and Ross, Arun},
  booktitle={Information Forensics and Security (WIFS), 2010 IEEE International Workshop on},
  pages={1--6},
  year={2010},
  organization={IEEE}
}

@inproceedings{cao2012predicting,
  title={Predicting gender and weight from human metrology using a copula model},
  author={Cao, Deng and Chen, Cunjian and Adjeroh, Donald and Ross, Arun},
  booktitle={Biometrics: Theory, Applications and Systems (BTAS), 2012 IEEE Fifth International Conference on},
  pages={162--169},
  year={2012},
  organization={IEEE}
}
@misc{IBM_SPSS,
  author = {{IBM Corp.}},
  title = {{IBM SPSS Statistics for Windows}},
  year = {2023},
  version = {29.0},
  howpublished = {{Computer software}},
  url = {https://www.ibm.com/products/spss-statistics},
}

@misc{Pew_News,
  author = {{Pew Research Center}},
  title = {{The State of the News Media 2023}},
  year = {2023},
  note = {{Data set}},
  organization = {{American Trends Panel Survey}},
  url = {https://www.pewresearch.org/journalism/dataset/},
}
