## Supplementary material for "Generative Language Modeling for Antibody CDR Grafting and Alignment-driven De Novo Design": TeX files: zHenriquesLab-Mendeley.bib

@article{barreiro2009high,
    title = {{High-throughput SNP genotyping: combining tag SNPs and molecular beacons.}},
    year = {2009},
    journal = {Methods in molecular biology (Clifton, N.J.)},
    author = {Barreiro, Luis B and Henriques, Ricardo and Mhlanga, Musa M},
    pages = {255--76},
    volume = {578},
    publisher = {Humana Press},
    url = {http://www.ncbi.nlm.nih.gov/pubmed/19768600},
    isbn = {1940-6029 (Electronic){\textbackslash}r1064-3745 (Linking)},
    doi = {10.1007/978-1-60327-411-1{\_}17},
    issn = {1940-6029},
    pmid = {19768600},
    keywords = {C-Type, C-Type: genetics, Cell Adhesion Molecules, Cell Adhesion Molecules: genetics, Cell Surface, Cell Surface: genetics, DNA Primers, DNA Primers: metabolism, Fluorescent Dyes, Fluorescent Dyes: metabolism, Genetic Loci, Genetic Loci: genetics, Genotype, High-Throughput Screening Assays, High-Throughput Screening Assays: methods, Humans, Lectins, Molecular Probes, Molecular Probes: metabolism, Polymorphism, Receptors, Reverse Transcriptase Polymerase Chain Reaction, Single Nucleotide, Single Nucleotide: genetics}
}

@article{Almada2015,
    title = {{PALM and STORM: Into large fields and high-throughput microscopy with sCMOS detectors.}},
    year = {2015},
    journal = {Methods (San Diego, Calif.)},
    author = {Almada, Pedro and Culley, Siân and Henriques, Ricardo},
    month = {10},
    pages = {109--21},
    volume = {88},
    url = {http://www.ncbi.nlm.nih.gov/pubmed/26079924},
    doi = {10.1016/j.ymeth.2015.06.004},
    issn = {1095-9130},
    pmid = {26079924},
    keywords = {Hardware, Homogenization, Single-molecule localization, sCMOS}
}

@article{culley2016infection,
    title = {{Infection Counter: Automated Quantification of in Vitro Virus Replication by Fluorescence Microscopy.}},
    year = {2016},
    journal = {Viruses},
    author = {Culley, Siân and Towers, Greg J. and Selwood, David L. and Henriques, Ricardo and Grove, Joe},
    number = {7},
    month = {7},
    pages = {201},
    volume = {8},
    publisher = {Multidisciplinary Digital Publishing Institute},
    url = {http://www.ncbi.nlm.nih.gov/pubmed/27455304 http://www.pubmedcentral.nih.gov/articlerender.fcgi?artid=PMC4974536},
    doi = {10.3390/v8070201},
    issn = {1999-4915},
    pmid = {27455304},
    keywords = {ImageJ, focus-forming assay, hepatitis C virus, image quantification, plaque assay}
}

@article{Soares2013,
    title = {{Regulated vesicle fusion generates signaling nanoterritories that control T cell activation at the immunological synapse}},
    year = {2013},
    journal = {Journal of Experimental Medicine},
    author = {Soares, H. and Henriques, R. and Sachse, M. and Ventimiglia, L. and Alonso, M. A. and Zimmer, C. and Thoulouze, M.-I. and Alcover, A.},
    number = {11},
    pages = {2415--2433},
    volume = {210},
    url = {http://www.jem.org/cgi/doi/10.1084/jem.20130150},
    doi = {10.1084/jem.20130150},
    issn = {0022-1007}
}

@article{Gustafsson2016,
    title = {{Fast live-cell conventional fluorophore nanoscopy with ImageJ through super-resolution radial fluctuations.}},
    year = {2016},
    journal = {Nature communications},
    author = {Gustafsson, Nils and Culley, Siân and Ashdown, George and Owen, Dylan M and Pereira, Pedro Matos and Henriques, Ricardo},
    number = {12471},
    month = {8},
    pages = {12471},
    volume = {7},
    url = {http://www.ncbi.nlm.nih.gov/pubmed/27514992 http://www.pubmedcentral.nih.gov/articlerender.fcgi?artid=PMC4990649},
    isbn = {2041-1723 (Electronic){\textbackslash}r2041-1723 (Linking)},
    doi = {10.1038/ncomms12471},
    issn = {2041-1723},
    pmid = {27514992}
}

@article{sirianni2016mitochondria,
    title = {{Mitochondria mediate septin cage assembly to promote autophagy of Shigella.}},
    year = {2016},
    journal = {EMBO reports},
    author = {Sirianni, Andrea and Krokowski, Sina and Lobato-M{\'{a}}rquez, Damián and Buranyi, Stephen and Pfanzelter, Julia and Galea, Dieter and Willis, Alexandra and Culley, Siân and Henriques, Ricardo and Larrouy-Maumus, Gerald and Hollinshead, Michael and Sancho-Shimizu, Vanessa and Way, Michael and Mostowy, Serge},
    number = {7},
    month = {7},
    pages = {1029--43},
    volume = {17},
    publisher = {EMBO Press},
    url = {http://www.ncbi.nlm.nih.gov/pubmed/27259462 http://www.pubmedcentral.nih.gov/articlerender.fcgi?artid=PMC4931556},
    isbn = {1469-221x},
    doi = {10.15252/embr.201541832},
    issn = {1469-3178},
    pmid = {27259462},
    keywords = {Shigella, autophagy, cytoskeleton, mitochondria, septin}
}

@article{ehsani2012hierarchies,
    title = {{Hierarchies of host factor dynamics at the entry site of Shigella flexneri during host cell invasion.}},
    year = {2012},
    journal = {Infection and immunity},
    author = {Ehsani, Soudeh and Santos, José Carlos and Rodrigues, Cristina D and Henriques, Ricardo and Audry, Laurent and Zimmer, Christophe and Sansonetti, Philippe and Tran Van Nhieu, Guy and Enninga, Jost},
    number = {7},
    month = {7},
    pages = {2548--57},
    volume = {80},
    publisher = {American Society for Microbiology},
    url = {http://www.pubmedcentral.nih.gov/articlerender.fcgi?artid=3416480&tool=pmcentrez&rendertype=abstract http://www.ncbi.nlm.nih.gov/pubmed/22526677 http://www.pubmedcentral.nih.gov/articlerender.fcgi?artid=PMC3416480},
    doi = {10.1128/IAI.06391-11},
    issn = {1098-5522},
    pmid = {22526677},
    keywords = {Cytoplasm, Cytoplasm: microbiology, Cytoskeleton, Cytoskeleton: metabolism, Epithelial Cells, Epithelial Cells: microbiology, Fluorescence, HeLa Cells, Host-Pathogen Interactions, Humans, Microscopy, Shigella flexneri, Shigella flexneri: pathogenicity, Time Factors, Time-Lapse Imaging, Vacuoles, Vacuoles: microbiology}
}

@article{clough2016k63,
    title = {{K63-Linked Ubiquitination Targets Toxoplasma gondii for Endo-lysosomal Destruction in IFN{$\gamma$}-Stimulated Human Cells}},
    year = {2016},
    journal = {PLOS Pathogens},
    author = {Clough, Barbara and Wright, Joseph D and Pereira, Pedro M and Hirst, Elizabeth M and Johnston, Ashleigh C and Henriques, Ricardo and Frickel, Eva-Maria},
    editor = {Denkers, Eric Y},
    number = {11},
    month = {11},
    pages = {e1006027},
    volume = {12},
    publisher = {Public Library of Science},
    url = {http://dx.plos.org/10.1371/journal.ppat.1006027},
    doi = {10.1371/journal.ppat.1006027},
    issn = {1553-7374}
}

@article{gomes2012highly,
    title = {{Highly dynamic host actin reorganization around developing Plasmodium inside hepatocytes.}},
    year = {2012},
    journal = {PloS one},
    author = {Gomes-Santos, Carina S S and Itoe, Maurice A and Afonso, Cristina and Henriques, Ricardo and Gardner, Rui and Sep{\'{u}}lveda, Nuno and Simões, Pedro D and Raquel, Helena and Almeida, António Paulo and Moita, Luis F and Frischknecht, Friedrich and Mota, Maria M and {others}},
    number = {1},
    month = {1},
    pages = {e29408},
    volume = {7},
    publisher = {Public Library of Science},
    url = {http://www.pubmedcentral.nih.gov/articlerender.fcgi?artid=3253080&tool=pmcentrez&rendertype=abstract},
    doi = {10.1371/journal.pone.0029408},
    issn = {1932-6203},
    pmid = {22238609},
    keywords = {Actin Cytoskeleton, Actin Cytoskeleton: metabolism, Actins, Actins: metabolism, Animals, Cells, Cultured, Gelsolin, Gelsolin: metabolism, Gelsolin: physiology, Genetically Modified, Green Fluorescent Proteins, Green Fluorescent Proteins: genetics, Green Fluorescent Proteins: metabolism, Hepatocytes, Hepatocytes: metabolism, Hepatocytes: parasitology, Host-Parasite Interactions, Host-Parasite Interactions: genetics, Host-Parasite Interactions: physiology, Humans, Inbred BALB C, Kinetics, Liver, Liver: metabolism, Liver: parasitology, Mice, Organisms, Plasmodium, Plasmodium: genetics, Plasmodium: metabolism, Plasmodium: physiology, Protein Multimerization, Protein Multimerization: physiology, Tubulin, Tubulin: metabolism}
}

@article{Lelek2012,
    title = {{Superresolution imaging of HIV in infected cells with FlAsH-PALM.}},
    year = {2012},
    journal = {Proceedings of the National Academy of Sciences of the United States of America},
    author = {Lelek, Mickaël and Di Nunzio, Francesca and Henriques, Ricardo and Charneau, Pierre and Arhel, Nathalie and Zimmer, Christophe},
    number = {22},
    month = {5},
    pages = {8564--9},
    volume = {109},
    publisher = {National Academy of Sciences},
    url = {http://www.ncbi.nlm.nih.gov/pubmed/22586087 http://www.pubmedcentral.nih.gov/articlerender.fcgi?artid=PMC3365178 http://www.pubmedcentral.nih.gov/articlerender.fcgi?artid=3365178&tool=pmcentrez&rendertype=abstract},
    isbn = {1013267109},
    doi = {10.1073/pnas.1013267109},
    issn = {1091-6490},
    pmid = {22586087},
    keywords = {Animals, Capsid, Capsid: chemistry, Capsid: metabolism, Cell Nucleus, Cell Nucleus: virology, Cell Shape, Cell Shape: physiology, Cysteine, Cysteine: chemistry, Cysteine: metabolism, Cytoplasm, Cytoplasm: virology, Fluorescence, Fluorescence: methods, Fluorescent Dyes, Fluorescent Dyes: chemistry, Fluorescent Dyes: metabolism, HIV Integrase, HIV Integrase: chemistry, HIV Integrase: metabolism, HIV-1, HIV-1: enzymology, HIV-1: physiology, HeLa Cells, Host-Pathogen Interactions, Humans, Mice, Microscopy, NIH 3T3 Cells, Peptides, Peptides: chemistry, Peptides: metabolism, Reproducibility of Results, Virion, Virion: chemistry, Virion: metabolism}
}

@article{herbert2012enhanced,
    title = {{Enhanced epifluorescence microscopy by uniform and intensity optimized illumination.}},
    year = {2012},
    journal = {Cytometry. Part A : the journal of the International Society for Analytical Cytology},
    author = {Herbert, Sébastien and Henriques, Ricardo},
    number = {4},
    month = {4},
    pages = {278--80},
    volume = {81},
    publisher = {Wiley Subscription Services, Inc., A Wiley Company},
    url = {http://www.ncbi.nlm.nih.gov/pubmed/22392634},
    doi = {10.1002/cyto.a.21177},
    issn = {1552-4930},
    pmid = {22392634}
}

@article{Scholefield2016,
    title = {{Super-resolution microscopy reveals a preformed NEMO lattice structure that is collapsed in incontinentia pigmenti.}},
    year = {2016},
    journal = {Nature communications},
    author = {Scholefield, Janine and Henriques, Ricardo and Savulescu, Anca F. and Fontan, Elisabeth and Boucharlat, Alix and Laplantine, Emmanuel and Smahi, Asma and Isra{\"{e}}l, Alain and Agou, Fabrice and Mhlanga, Musa M.},
    pages = {12629},
    volume = {7},
    url = {http://www.nature.com/doifinder/10.1038/ncomms12629%5Cnhttp://www.ncbi.nlm.nih.gov/pubmed/27586688%5Cnhttp://www.pubmedcentral.nih.gov/articlerender.fcgi?artid=PMC5025789},
    doi = {10.1038/ncomms12629},
    issn = {2041-1723},
    pmid = {27586688}
}

@article{gray2016virusmapper,
    title = {{VirusMapper: open-source nanoscale mapping of viral architecture through super-resolution microscopy.}},
    year = {2016},
    journal = {Scientific reports},
    author = {Gray, Robert D. M. and Beerli, Corina and Pereira, Pedro Matos and Scherer, Kathrin Maria and Samolej, Jerzy and Bleck, Christopher Karl Ernst and Mercer, Jason and Henriques, Ricardo},
    month = {7},
    pages = {29132},
    volume = {6},
    publisher = {Nature Publishing Group},
    url = {http://www.nature.com/articles/srep29132 http://www.ncbi.nlm.nih.gov/pubmed/27374400 http://www.pubmedcentral.nih.gov/articlerender.fcgi?artid=PMC4931586},
    doi = {10.1038/srep29132},
    issn = {2045-2322},
    pmid = {27374400}
}

@article{Pereira2015,
    title = {{High-content 3D multicolor super-resolution localization microscopy}},
    year = {2015},
    journal = {Methods in Cell Biology},
    author = {Pereira, Pedro M. and Almada, Pedro and Henriques, Ricardo},
    pages = {95--117},
    volume = {125},
    isbn = {9780128011034},
    doi = {10.1016/bs.mcb.2014.10.004},
    issn = {0091679X},
    pmid = {25640426},
    keywords = {3D, DSTORM, High-content screening, Immunofluorescence, Localization microscopy, Multicolor, Super-resolution}
}

@article{silva2016hiv,
    title = {{HIV-1 Nef Impairs the Formation of Calcium Membrane Territories Controlling the Signaling Nanoarchitecture at the Immunological Synapse}},
    year = {2016},
    journal = {The Journal of Immunology},
    author = {Silva, Joana G. and Martins, Nuno P. and Henriques, Ricardo and Soares, Helena},
    number = {10},
    month = {11},
    pages = {4042--4052},
    volume = {197},
    publisher = {American Association of Immunologists},
    url = {http://www.jimmunol.org/cgi/doi/10.4049/jimmunol.1601132 http://www.jimmunol.org/lookup/doi/10.4049/jimmunol.1601132},
    doi = {10.4049/jimmunol.1601132},
    issn = {0022-1767}
}

@article{Tarantino2014,
    title = {{TNF and IL-1 exhibit distinct ubiquitin requirements for inducing NEMO-IKK supramolecular structures.}},
    year = {2014},
    journal = {The Journal of cell biology},
    author = {Tarantino, Nadine and Tinevez, Jean-Yves and Crowell, Elizabeth Faris and Boisson, Bertrand and Henriques, Ricardo and Mhlanga, Musa and Agou, Fabrice and Isra{\"{e}}l, Alain and Laplantine, Emmanuel},
    number = {2},
    month = {1},
    pages = {231--45},
    volume = {204},
    url = {http://www.ncbi.nlm.nih.gov/pubmed/24446482 http://www.pubmedcentral.nih.gov/articlerender.fcgi?artid=PMC3897181},
    doi = {10.1083/jcb.201307172},
    issn = {1540-8140},
    pmid = {24446482},
    keywords = {Cell Line, HeLa Cells, Humans, I-kappa B Kinase, I-kappa B Kinase: analysis, I-kappa B Kinase: metabolism, Interleukin-1, Interleukin-1 Receptor-Associated Kinases, Interleukin-1 Receptor-Associated Kinases: analysi, Interleukin-1 Receptor-Associated Kinases: metabol, Interleukin-1: analysis, Interleukin-1: metabolism, Interleukin-1: physiology, NF-kappa B, NF-kappa B: analysis, NF-kappa B: metabolism, Receptors, Signal Transduction, Tumor, Tumor Necrosis Factor, Tumor Necrosis Factor-alpha, Tumor Necrosis Factor-alpha: analysis, Tumor Necrosis Factor-alpha: metabolism, Tumor Necrosis Factor-alpha: physiology, Tumor Necrosis Factor: analysis, Tumor Necrosis Factor: metabolism, Ubiquitin, Ubiquitin: genetics, Ubiquitin: metabolism, Ubiquitin: physiology, Ubiquitination}
}

@article{caetano2008degradation,
    title = {{Chronic arthritis directly induces quantitative and qualitative bone disturbances leading to compromised biomechanical properties.}},
    year = {2008},
    journal = {Clinical and experimental rheumatology},
    author = {Caetano-Lopes, J and Nery, A M and Henriques, R and Canh{\~{a}}o, H and Duarte, J and Amaral, P M and Vale, M and Moura, R A and Pereira, P a and Weinmann, P and Abdulghani, S and Souto-Carneiro, M and Rego, P and Monteiro, J and Sakagushi, S and Queiroz, M V and Konttinen, Y T and Gra{\c{c}}a, L and Vaz, M F and Fonseca, J E},
    number = {3},
    pages = {475--82},
    volume = {27},
    publisher = {Clinical and Experimental Rheumatology},
    url = {http://www.ncbi.nlm.nih.gov/pubmed/19604441},
    institution = {SPRINGER 233 SPRING ST, NEW YORK, NY 10013 USA},
    issn = {0392-856X},
    pmid = {19604441},
    keywords = {Animal, Animals, Arthritis, Arthritis: metabolism, Arthritis: pathology, Arthritis: physiopathology, Biomechanics, Bone Density, Bone Density: physiology, Chronic Disease, Collagen, Collagen: metabolism, Collagen: ultrastructure, Disease Models, Electron, Female, Femur, Femur: metabolism, Femur: pathology, Femur: physiopathology, Inbred BALB C, Lumbar Vertebrae, Lumbar Vertebrae: metabolism, Lumbar Vertebrae: pathology, Lumbar Vertebrae: physiopathology, Mice, Microscopy, Mutant Strains, Scanning, Spectrometry, X-Ray Emission}
}

@article{Henriques2010QuickPALM:ImageJ,
    title = {{QuickPALM: 3D real-time photoactivation nanoscopy image processing in ImageJ}},
    year = {2010},
    journal = {Nature Methods},
    author = {Henriques, Ricardo and Lelek, Mickael and Fornasiero, Eugenio F and Valtorta, Flavia and Zimmer, Christophe and Mhlanga, Musa M},
    number = {5},
    month = {5},
    pages = {339--340},
    volume = {7},
    url = {http://dx.doi.org/10.1038/nmeth0510-339 http://www.nature.com/doifinder/10.1038/nmeth0510-339},
    isbn = {1548-7091},
    doi = {10.1038/nmeth0510-339},
    issn = {1548-7091},
    pmid = {20431545}
}
