## Supplementary figures and images for "Generative Language Modeling for Antibody CDR Grafting and Alignment-driven De Novo Design"

### alignment_naturalness_monitoring.pdf

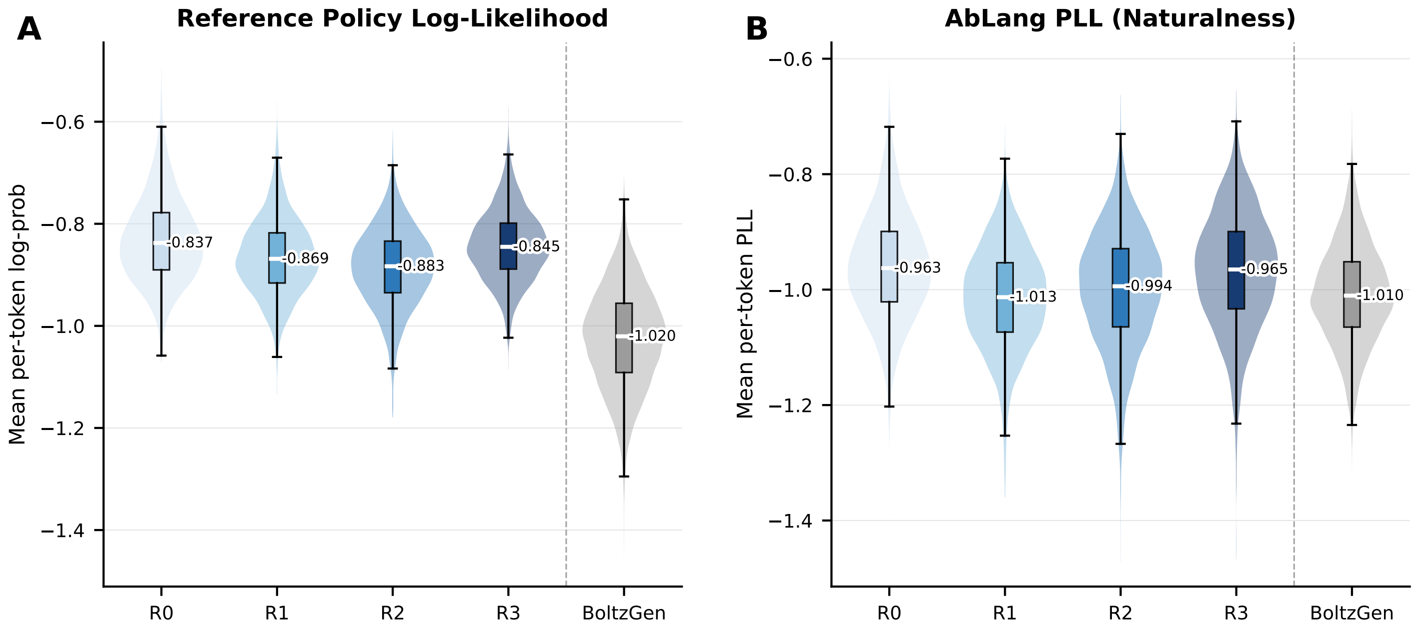

### alignment_naturalness_monitoring.png

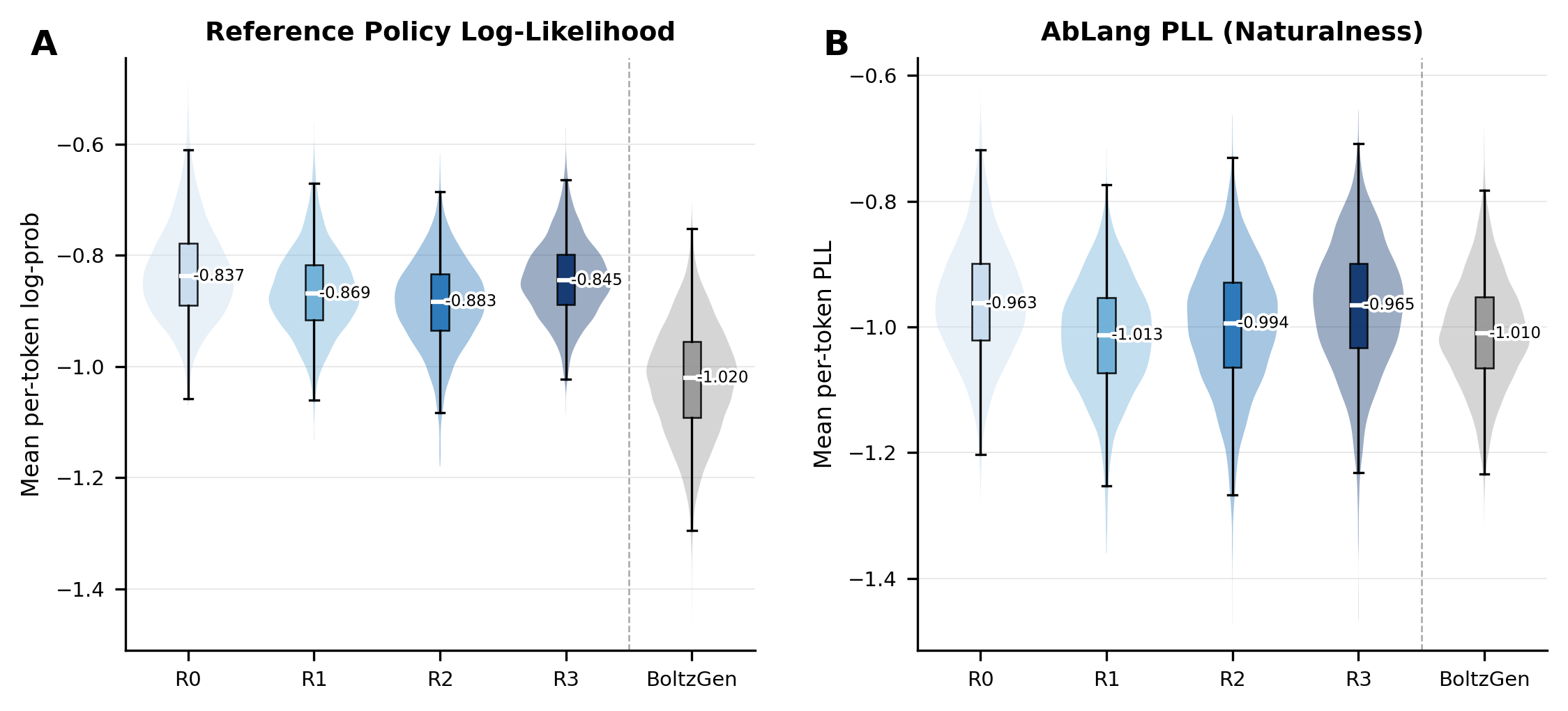

### alignment_pass_rate.pdf

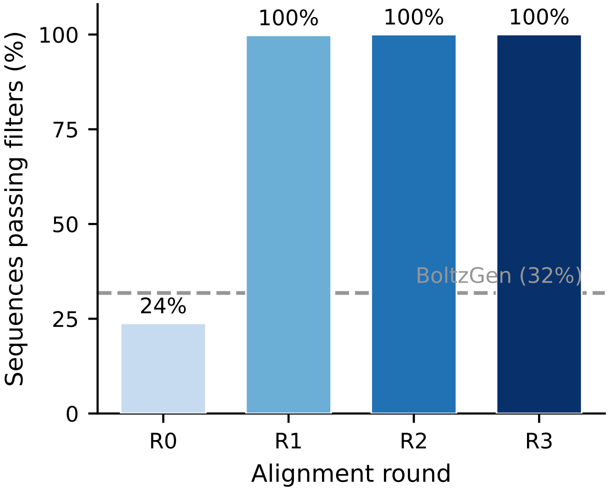

### alignment_progression.pdf

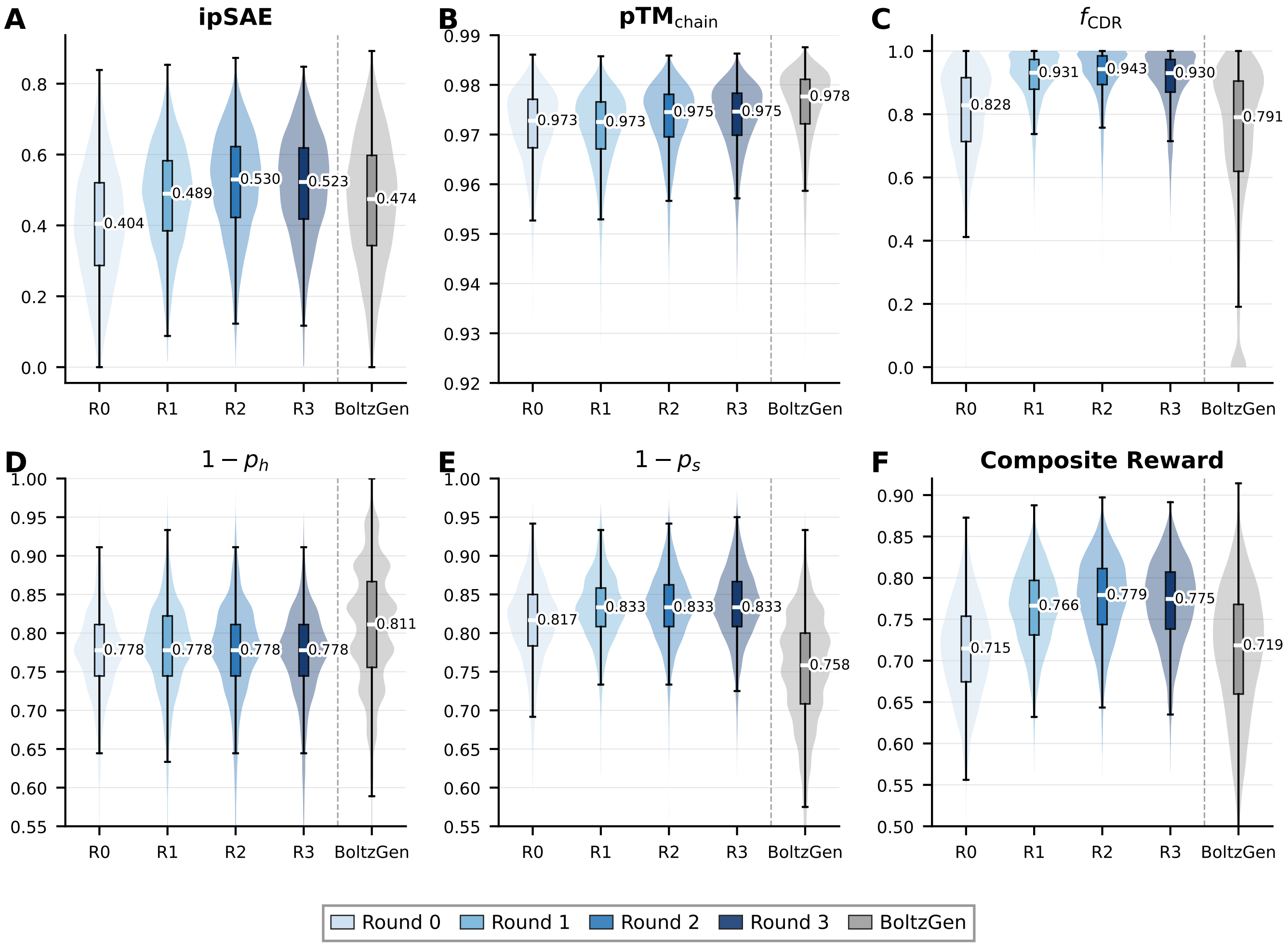

### alignment_progression.png

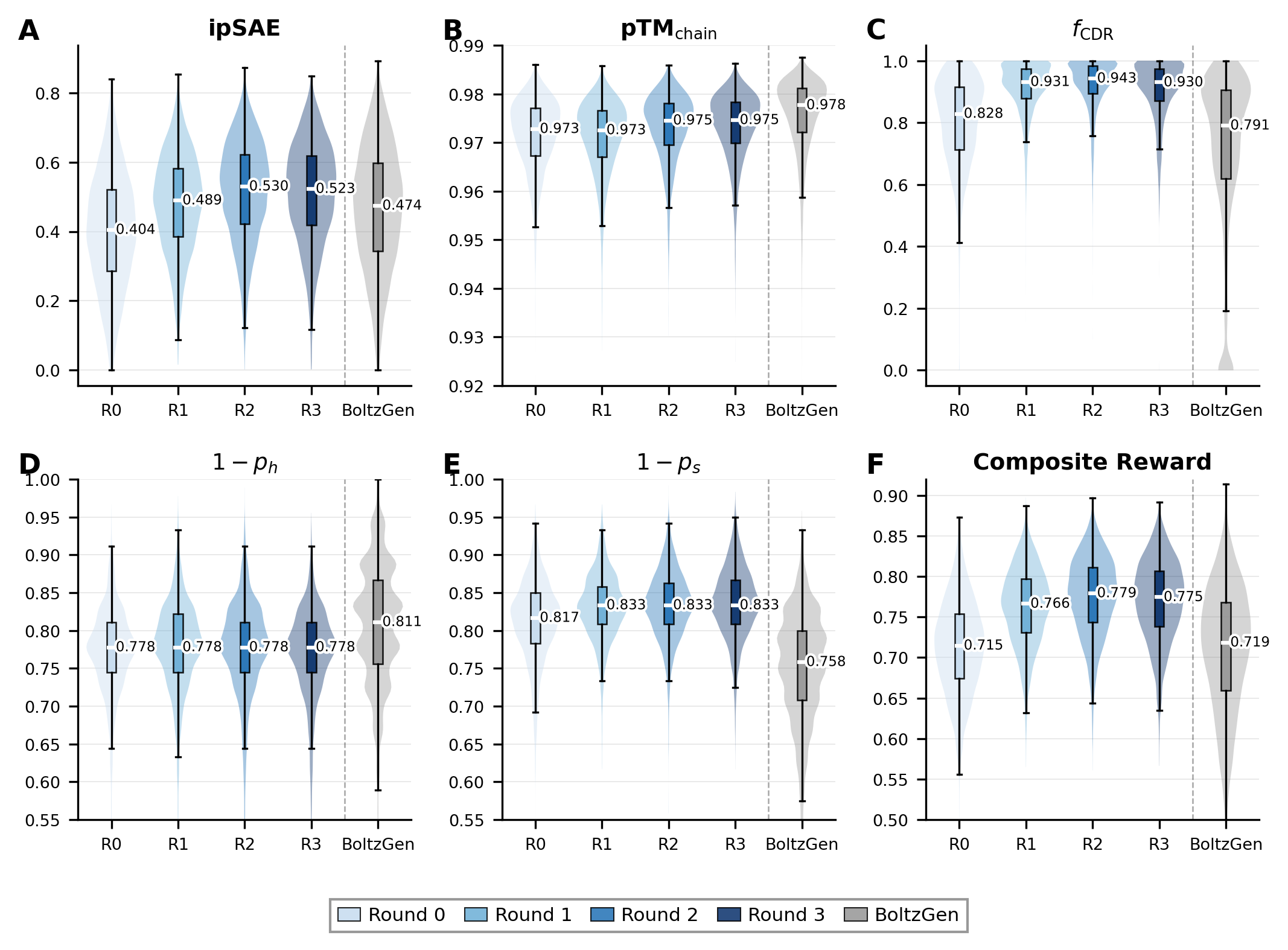

### alignment_reward_per_framework.pdf

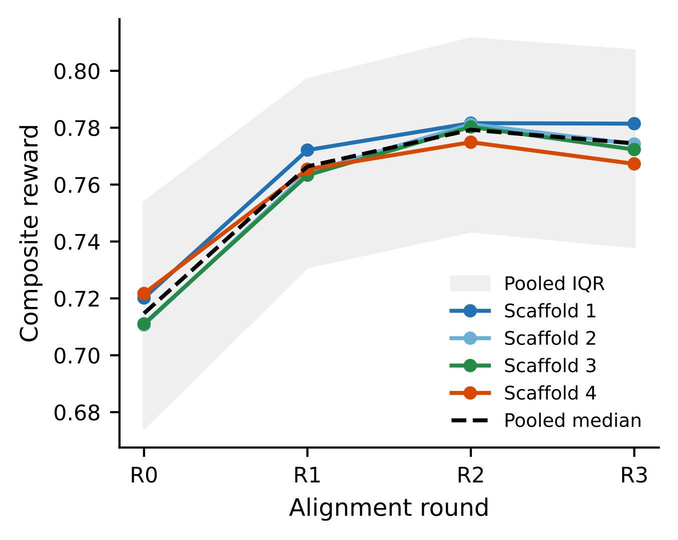

### cdr3_lenC_boxplot.pdf

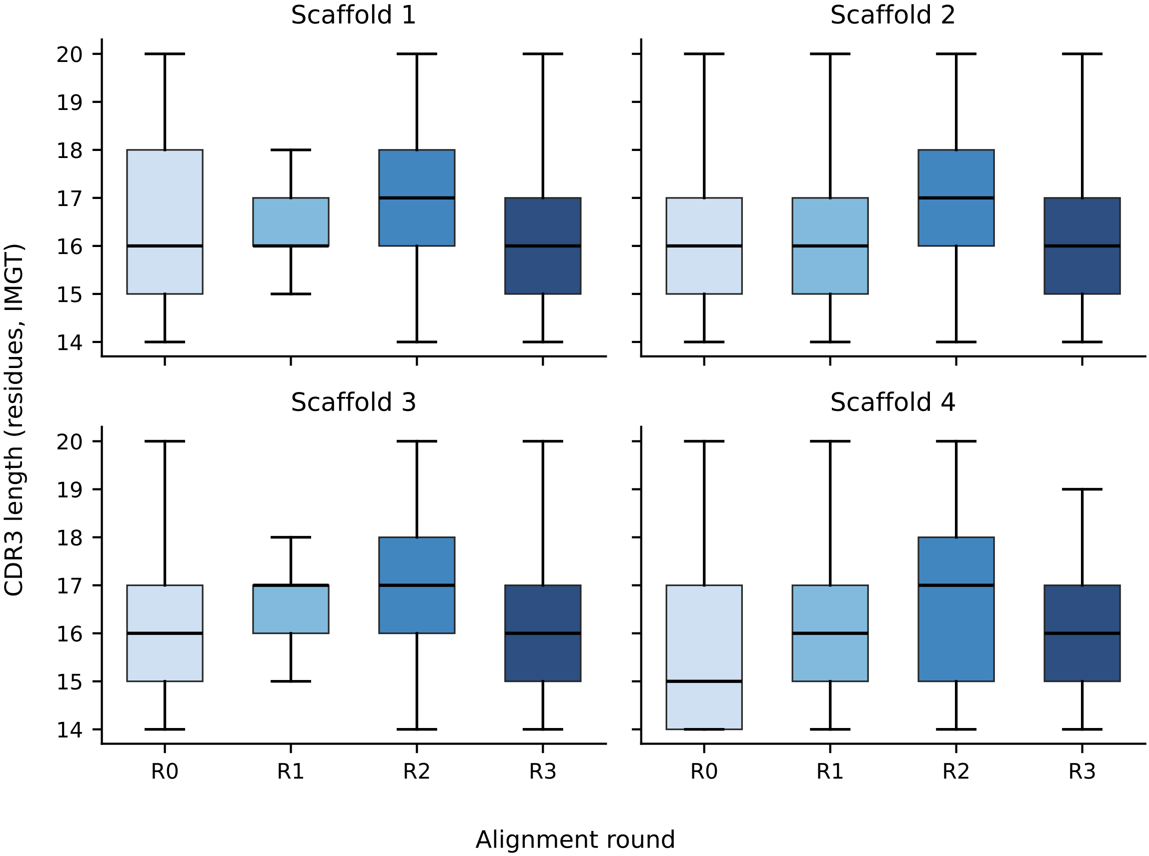

### cdr_confidence_histogram.png

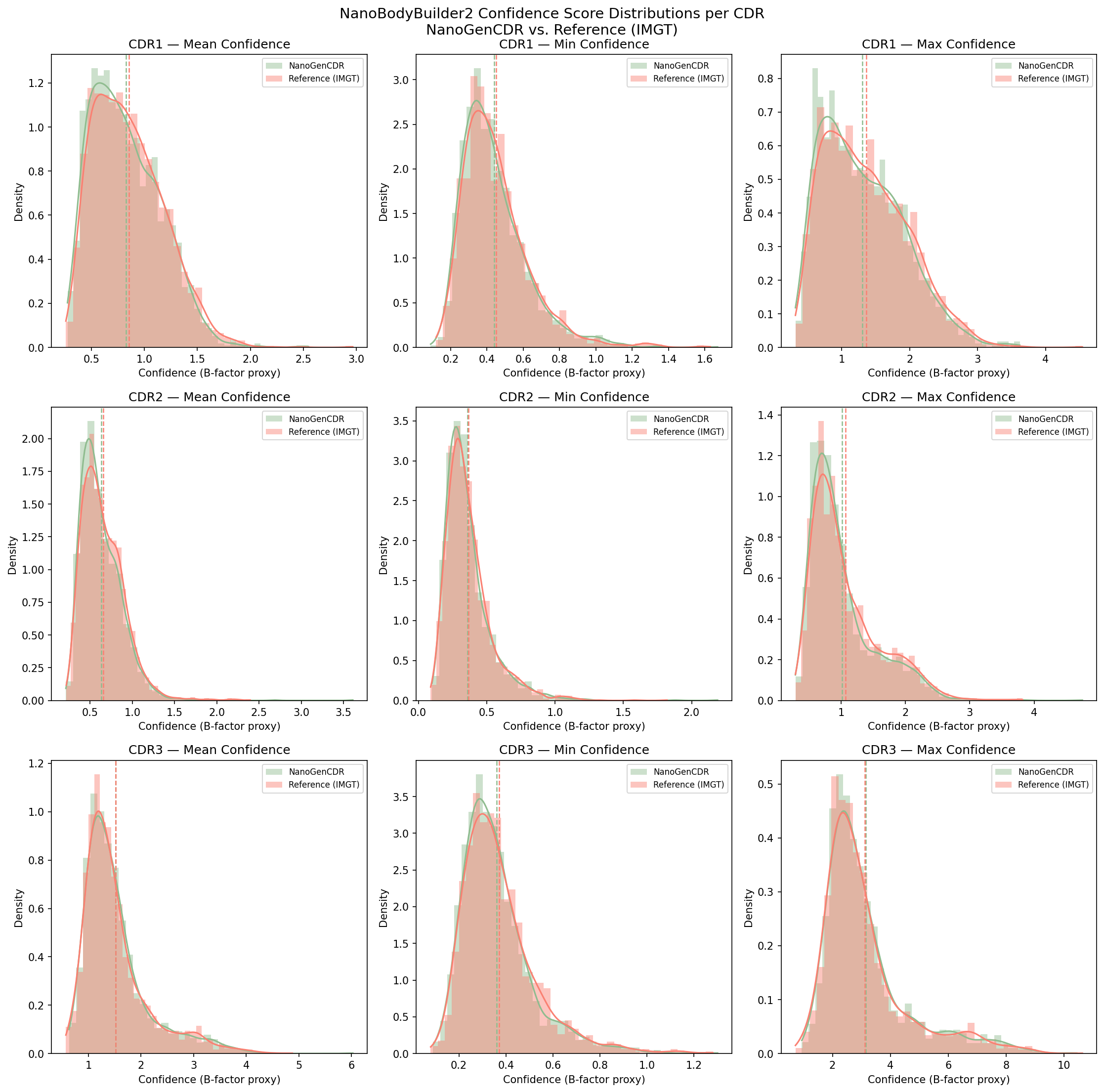

### cdr_length_comparison.png

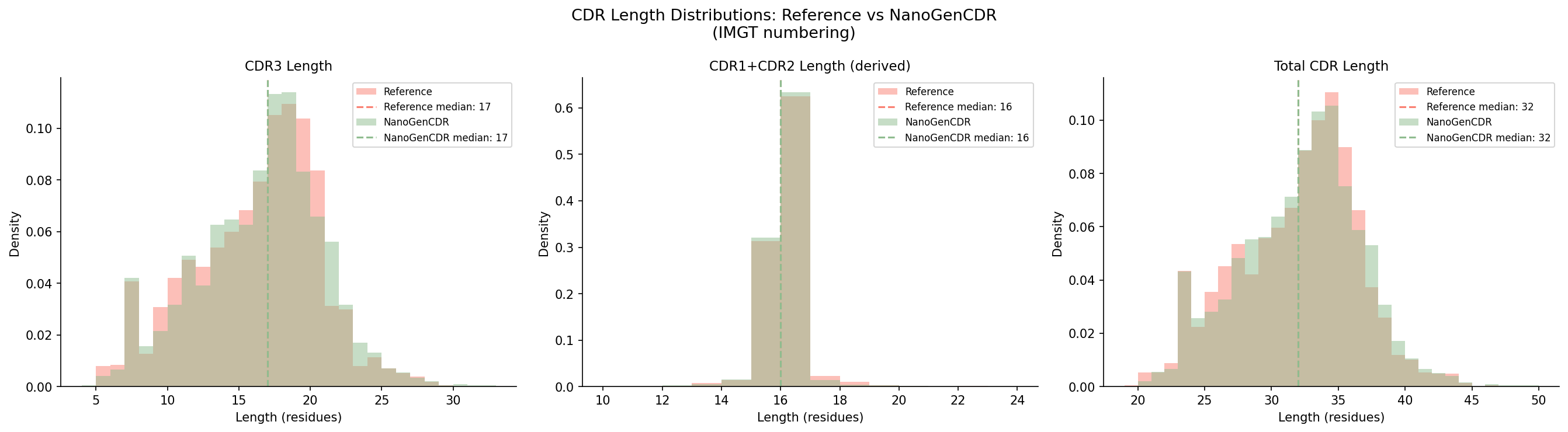

### cdr_length_distributions.pdf

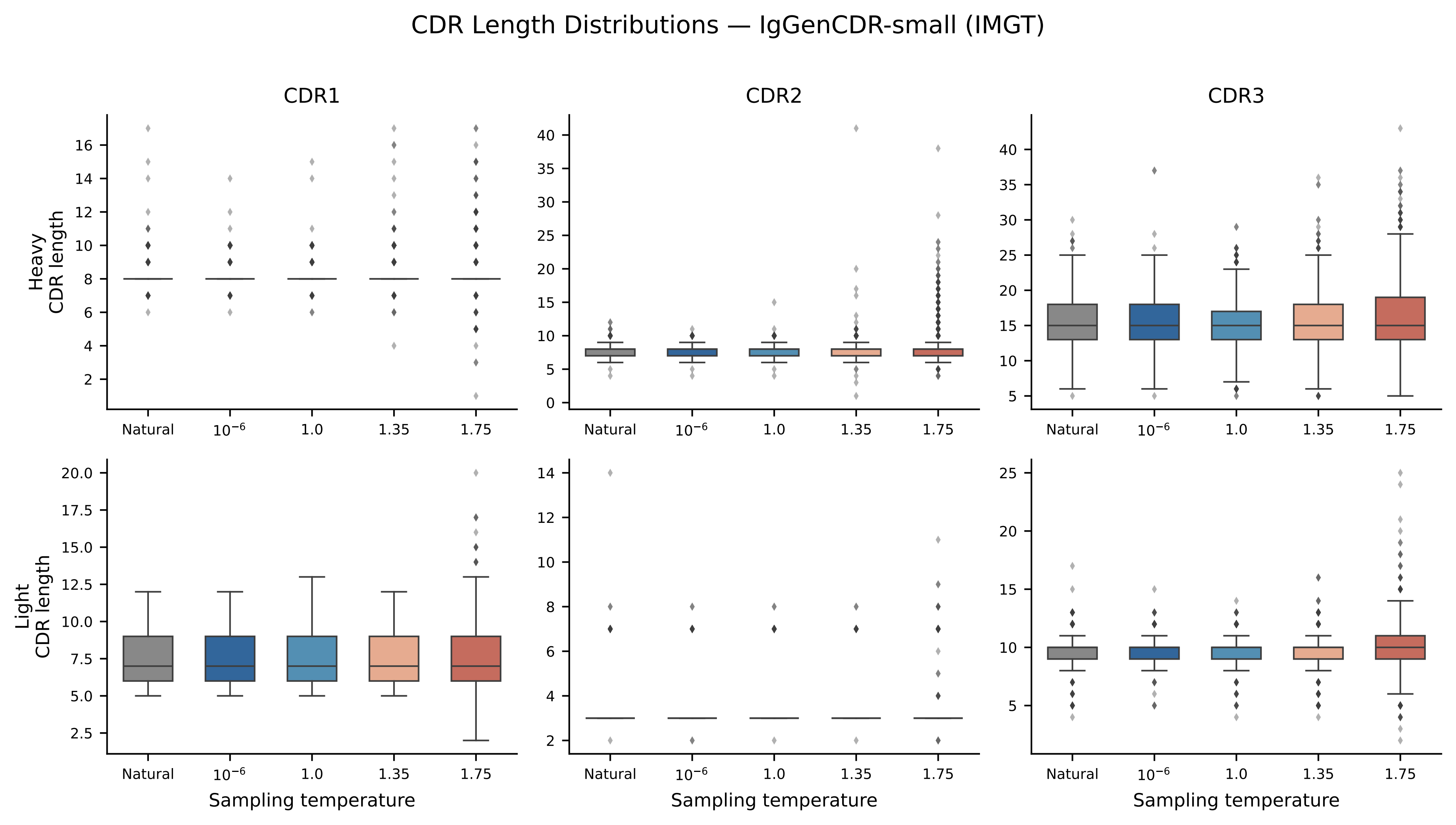

### cdr_lengths_20k_boltzgen.png

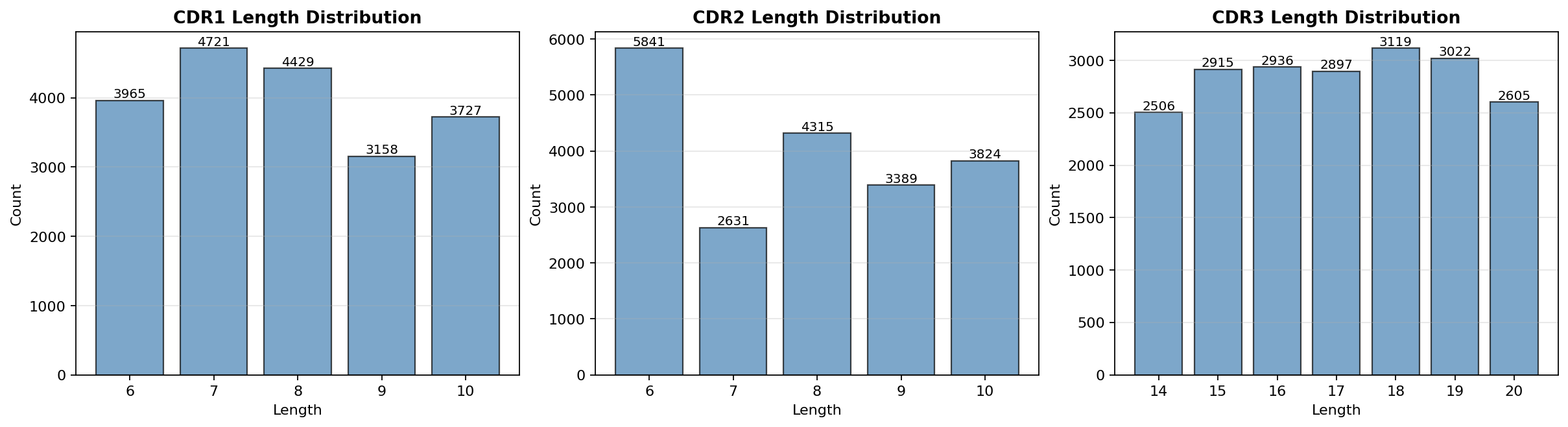

### cdr_lengths_histogram.pdf

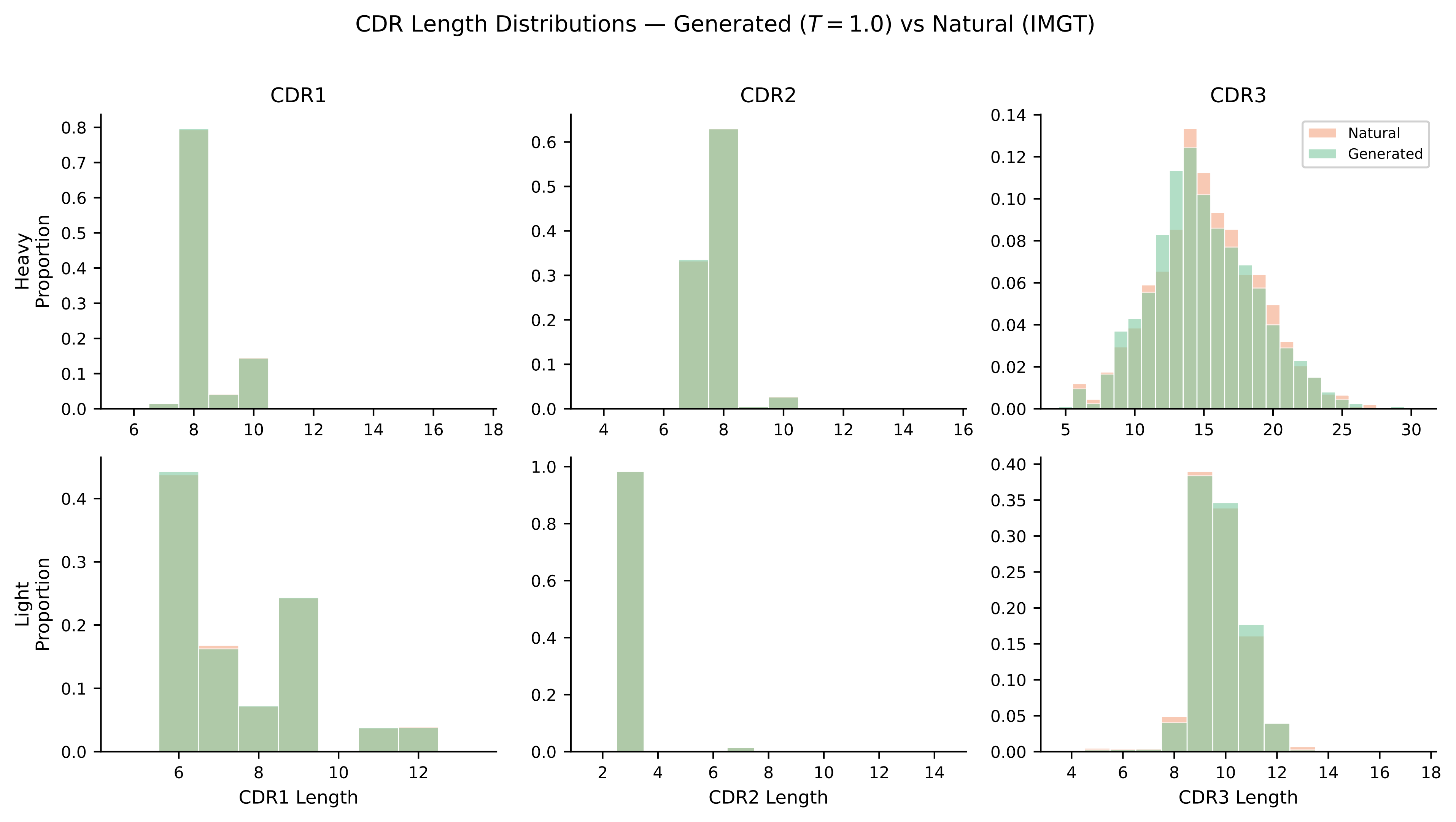

### cdr_lengths_histogram_nanobody_oasTrue_long.pdf

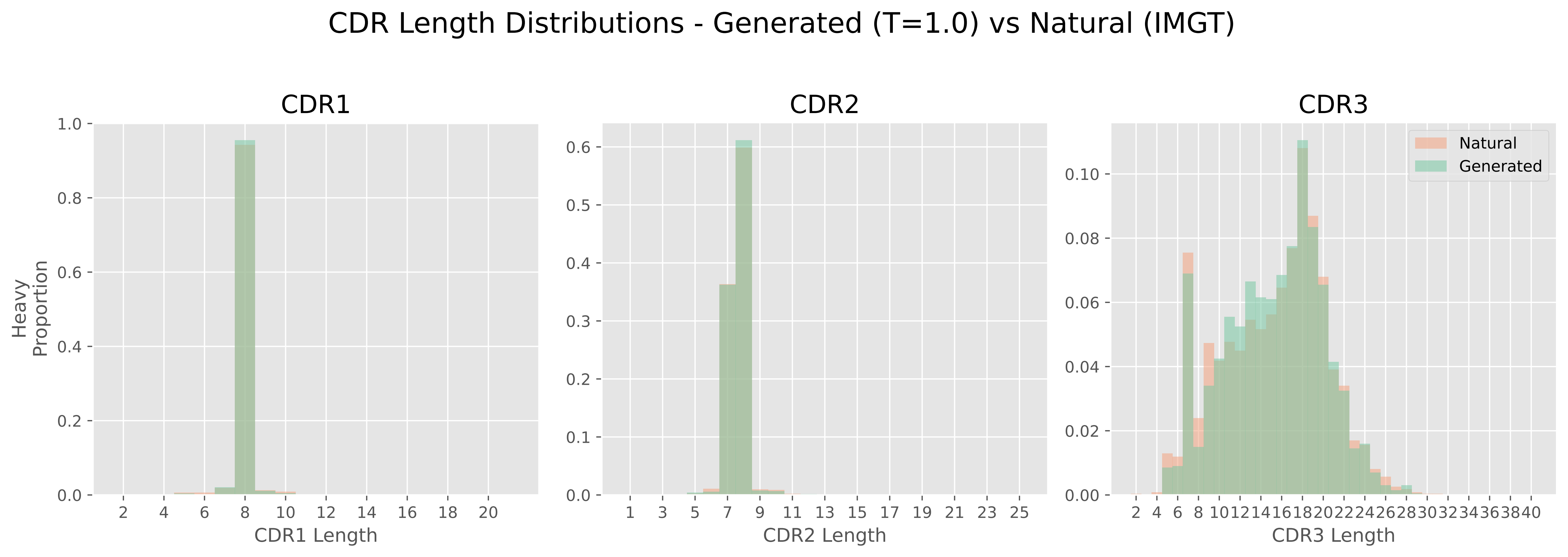

### esm2_histogram.pdf

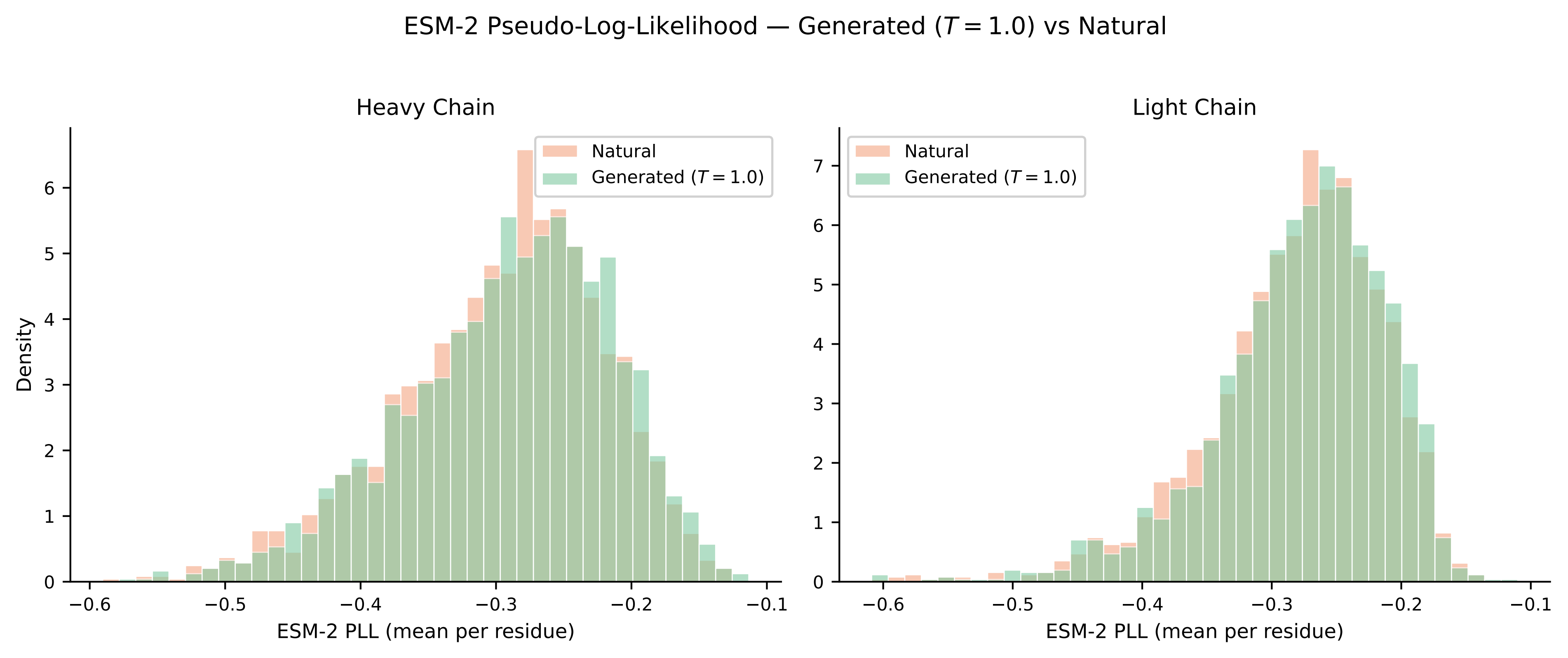

### esm2_pll.pdf

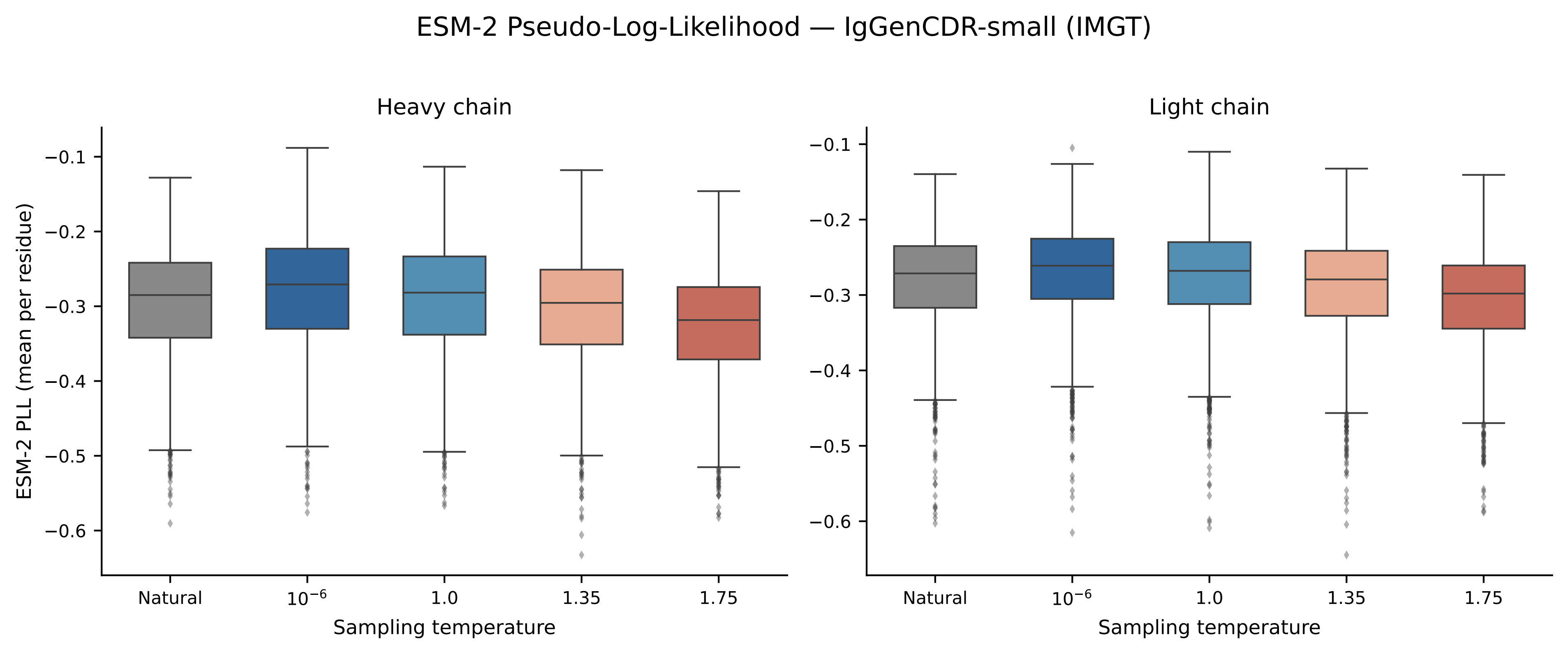

### flag_comparison.png

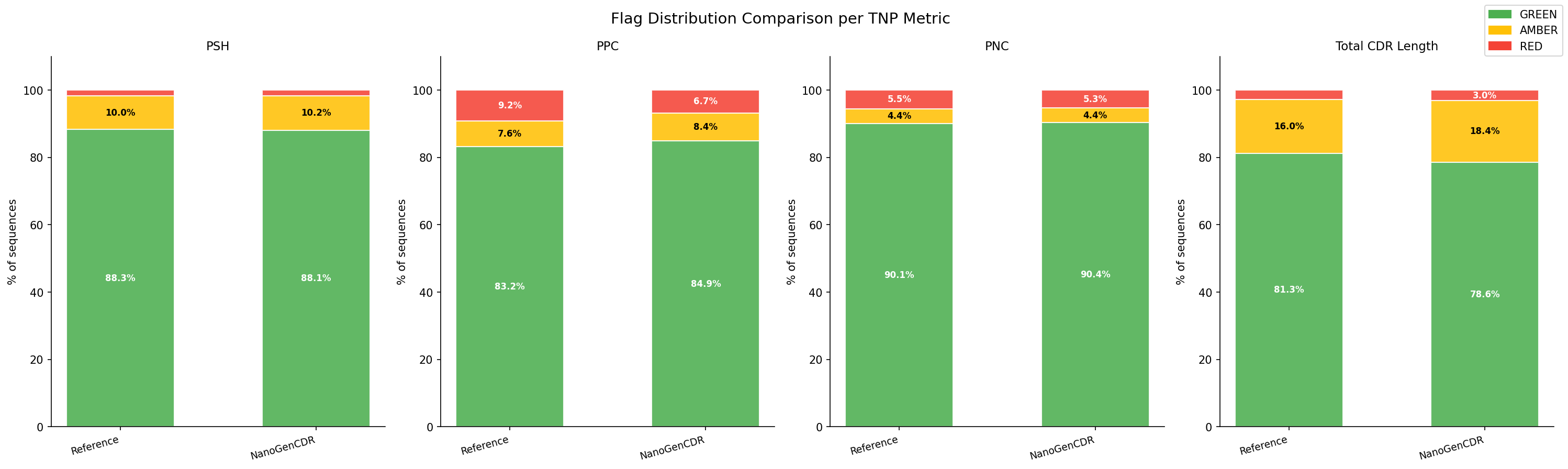

### GenCDR_main.pdf

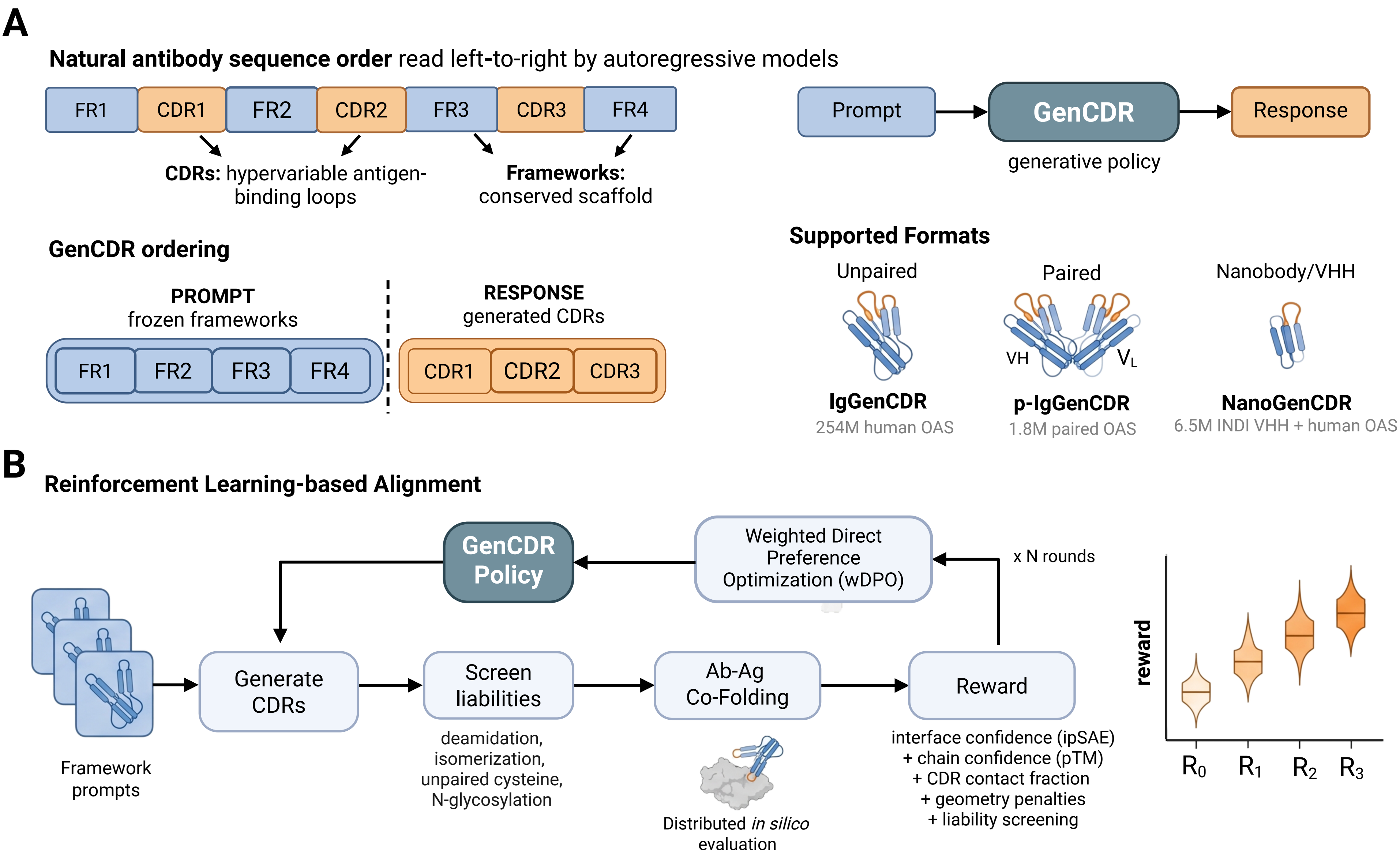

### iggencdr_cdr_lengths.pdf

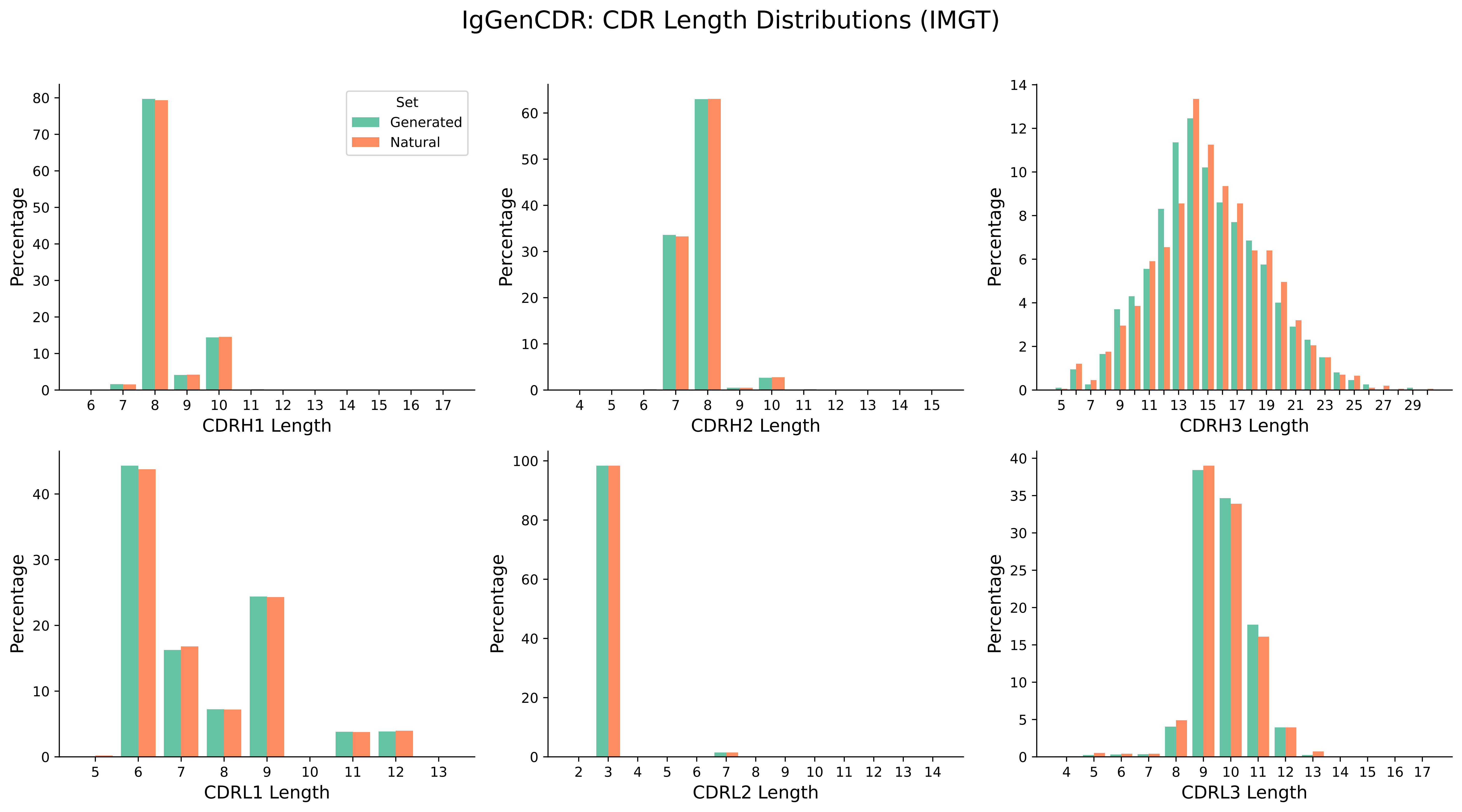

### nanogencdr_cdr_lengths.pdf

NanoGenCDR: CDR Length Distributions (IMGT)

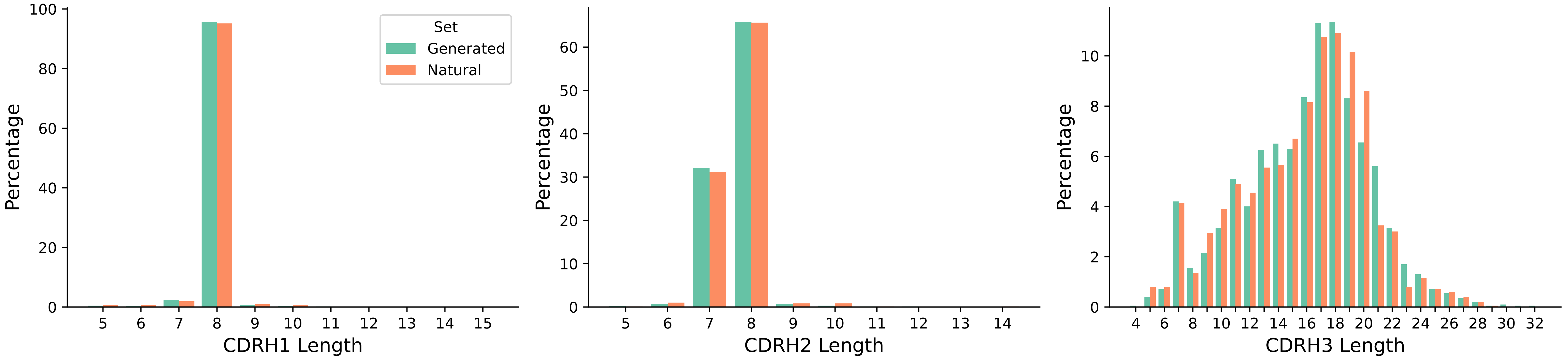

### oasis_humanness.pdf

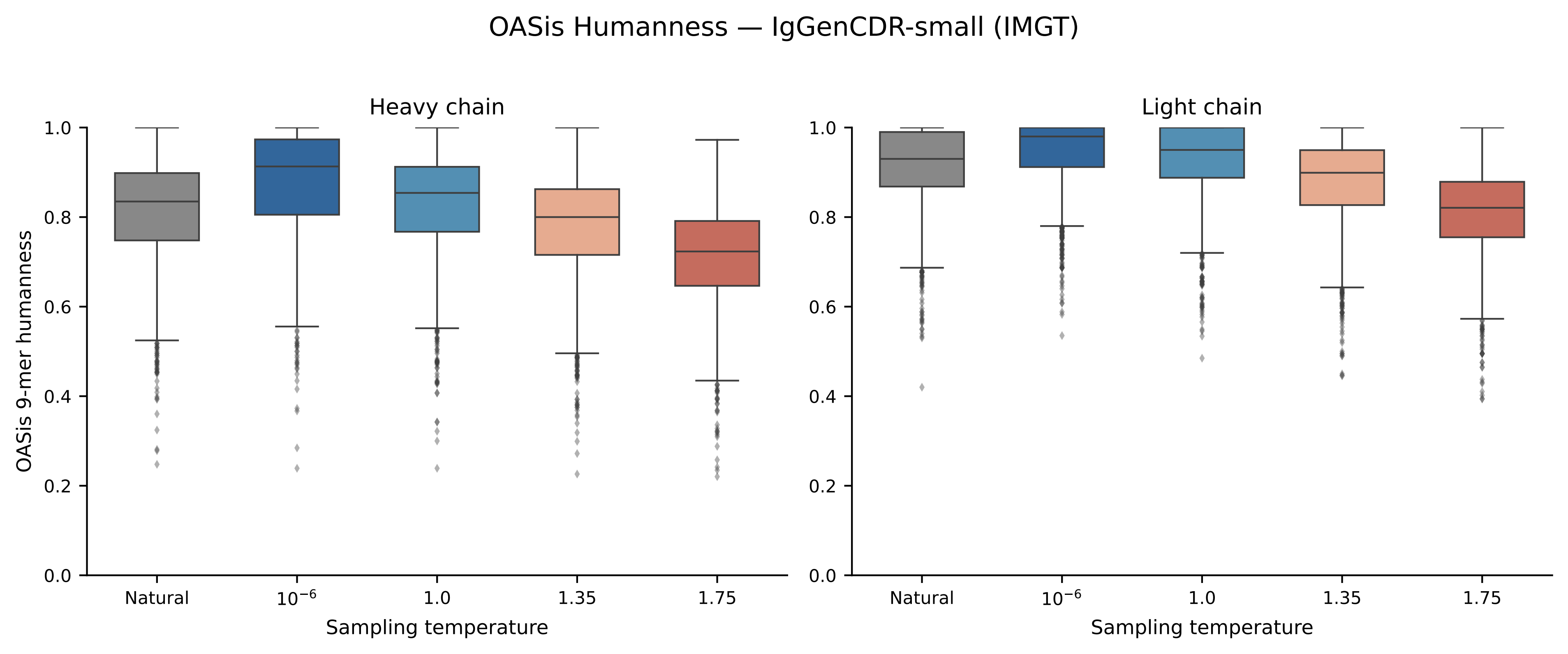

### oasis_humanness_per_framework.pdf

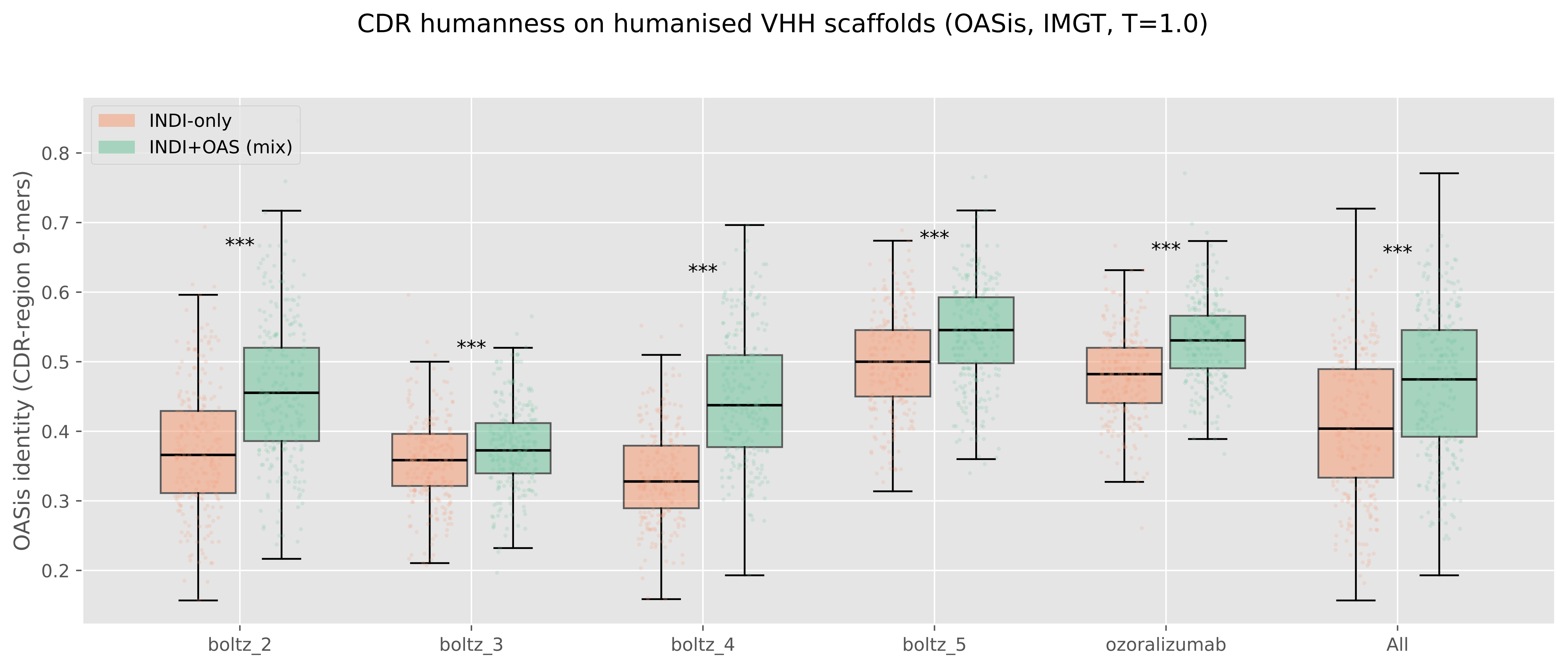

### oasis_humanness_per_framework.png

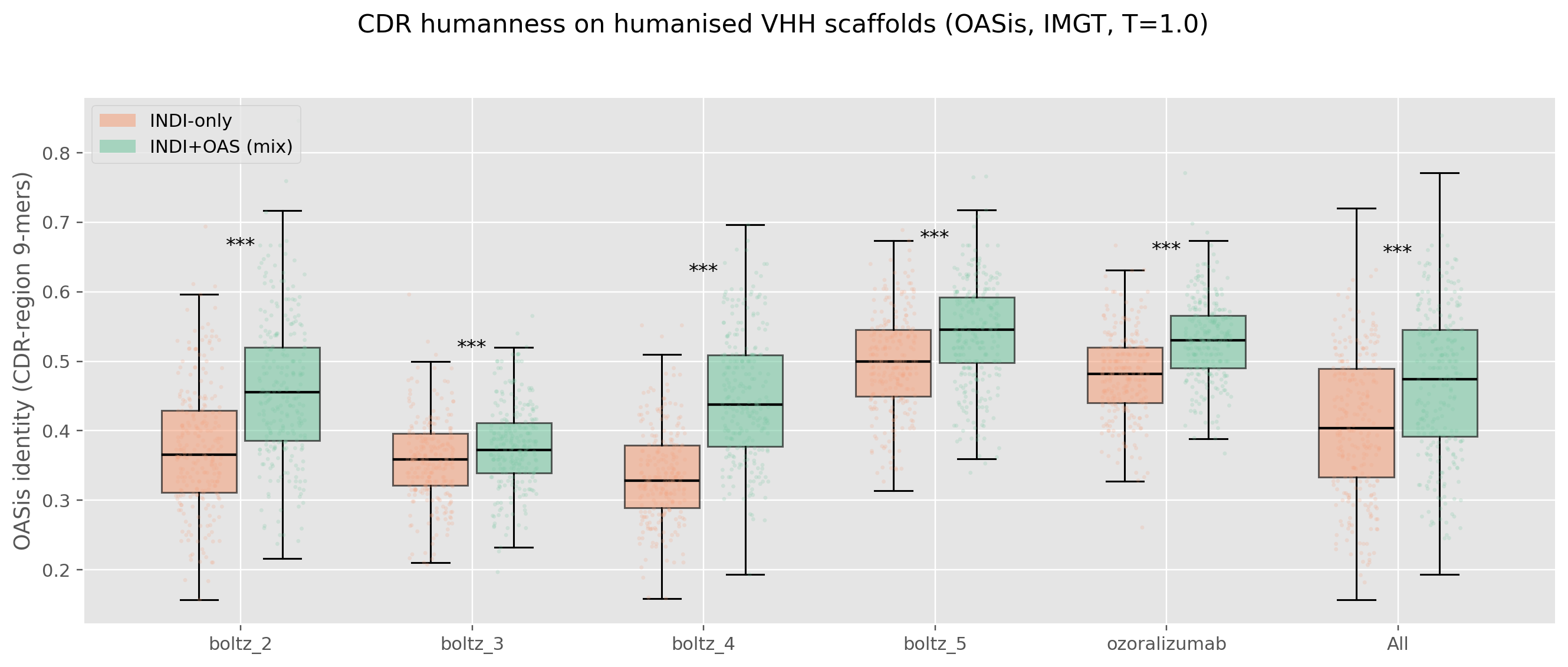

### p_iggencdr_cdr_lengths.pdf

#### p-IgGenCDR: CDR Length Distributions (IMGT)

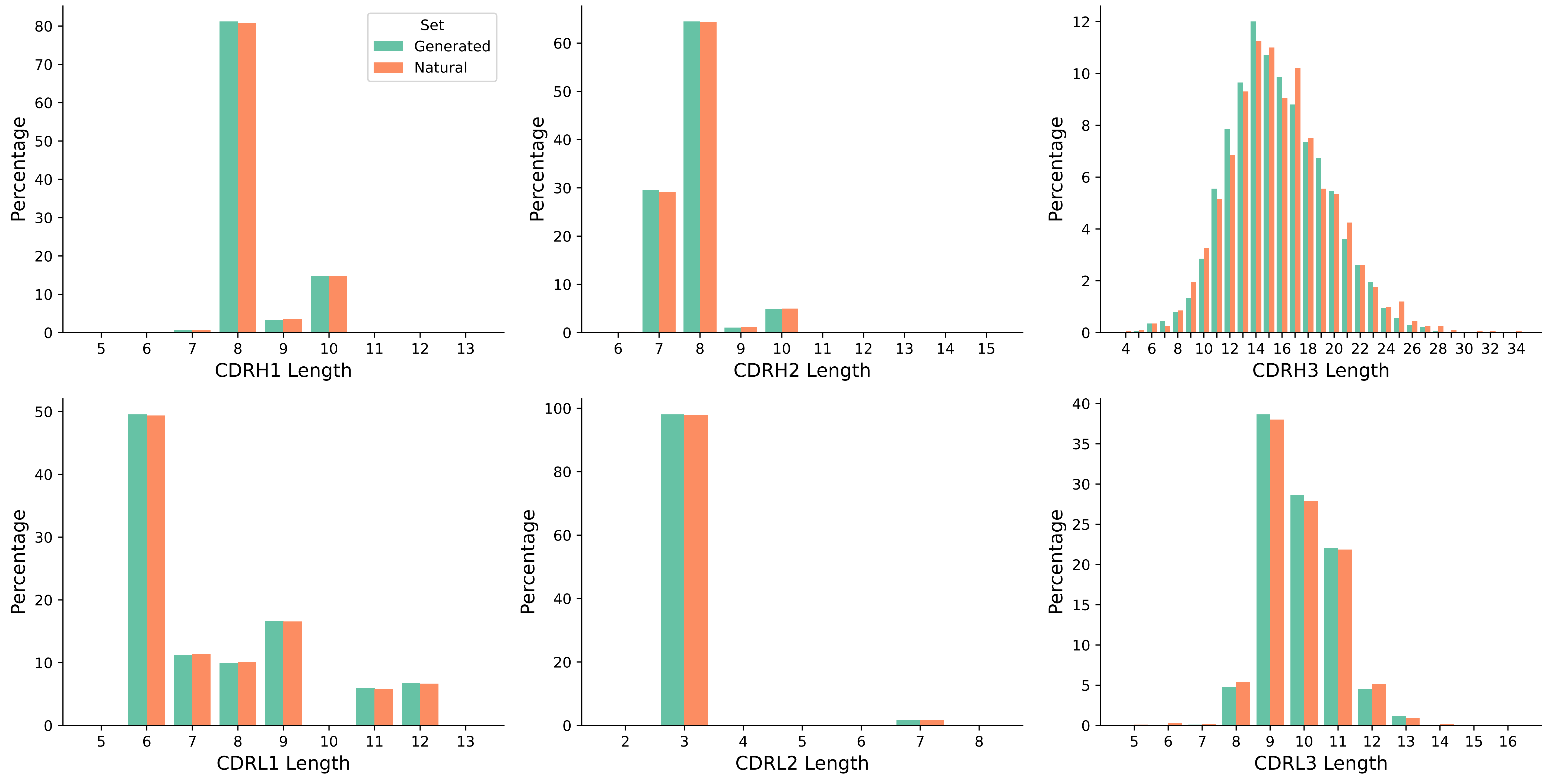

### sabdab_fcdr_reference.pdf

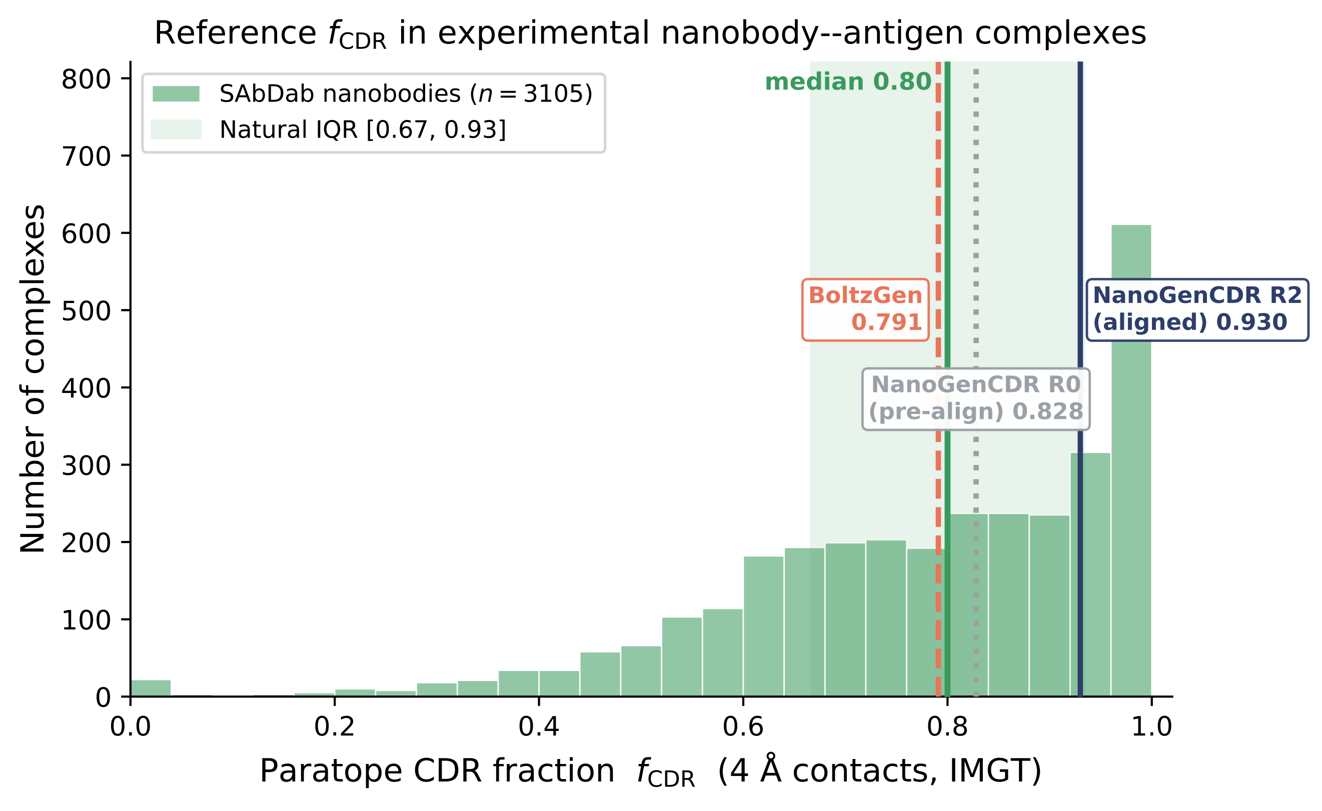

### sabdab_fcdr_reference.png

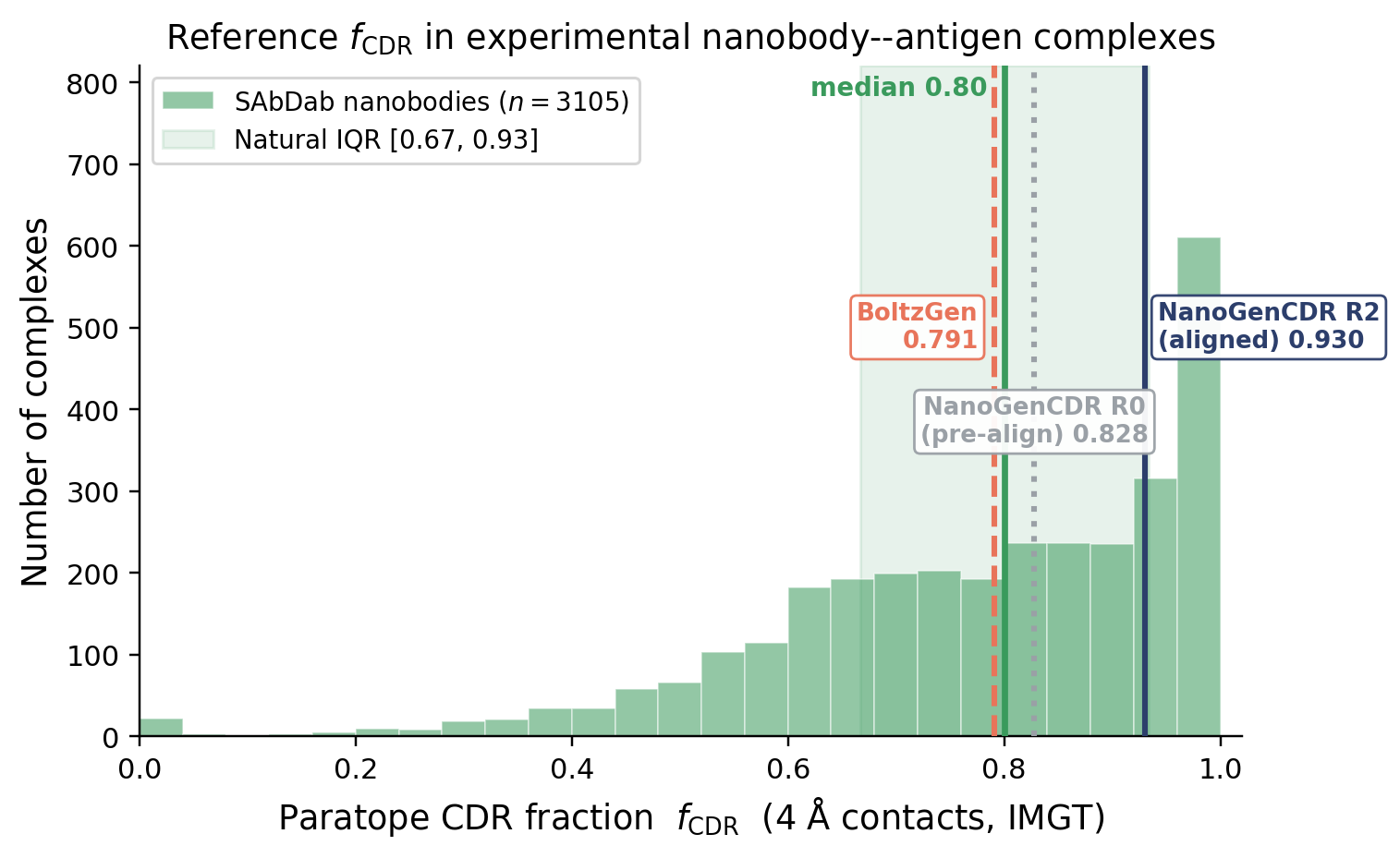

### sabdab_fcdr_therapeutic.pdf

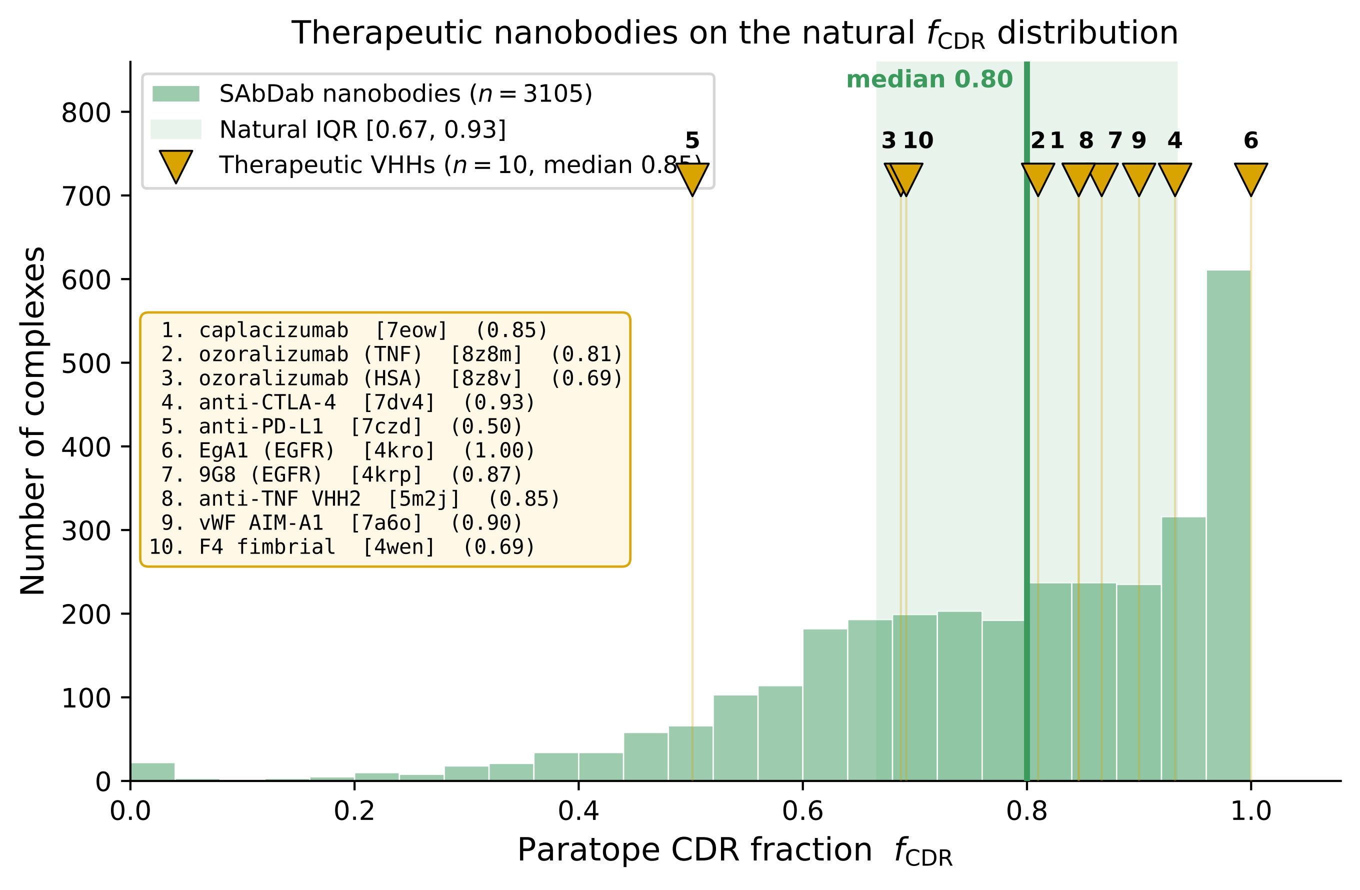

### sabdab_fcdr_therapeutic.png

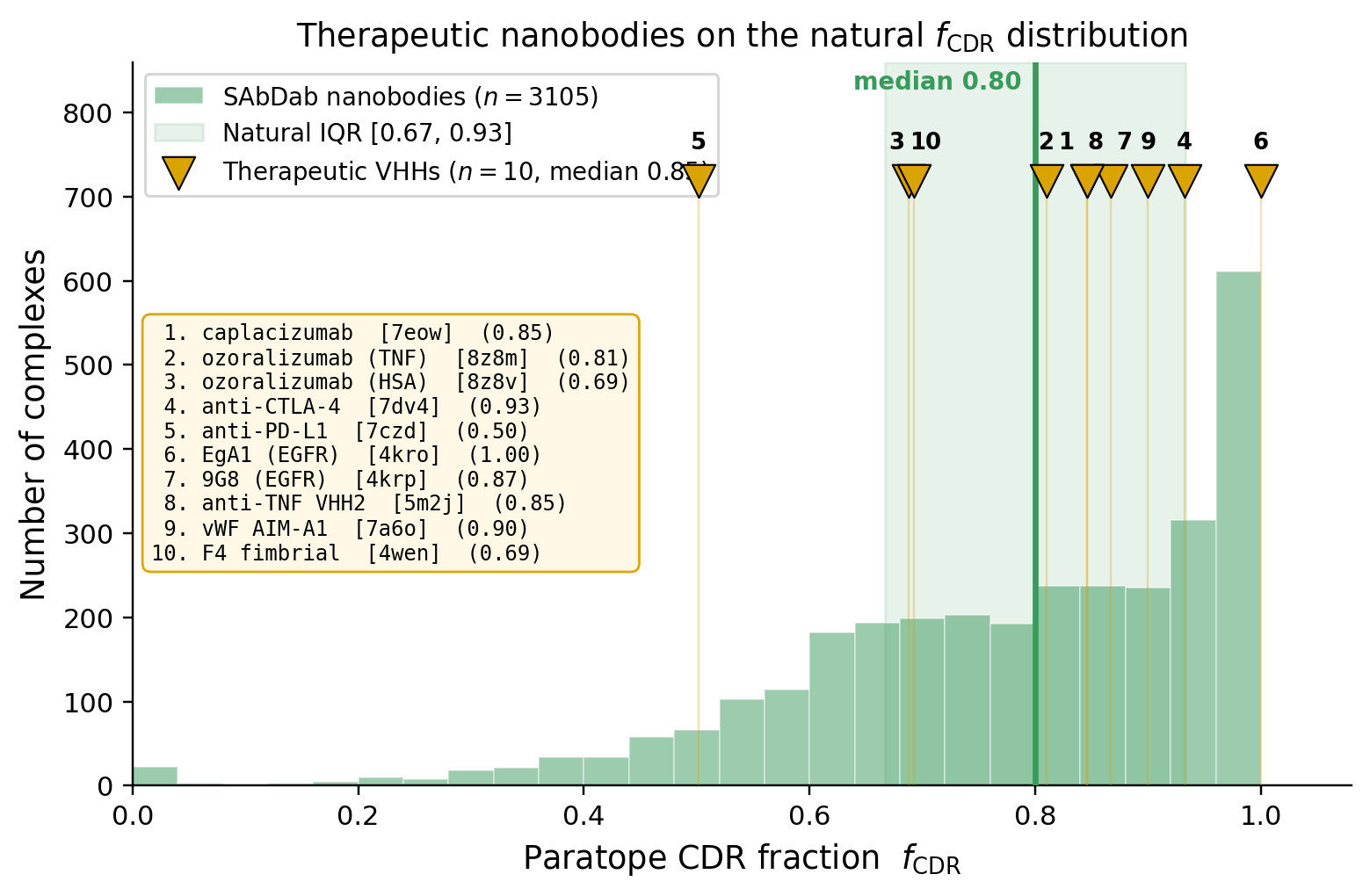

### sabdab_fcdr_therapeutic_cb8.pdf

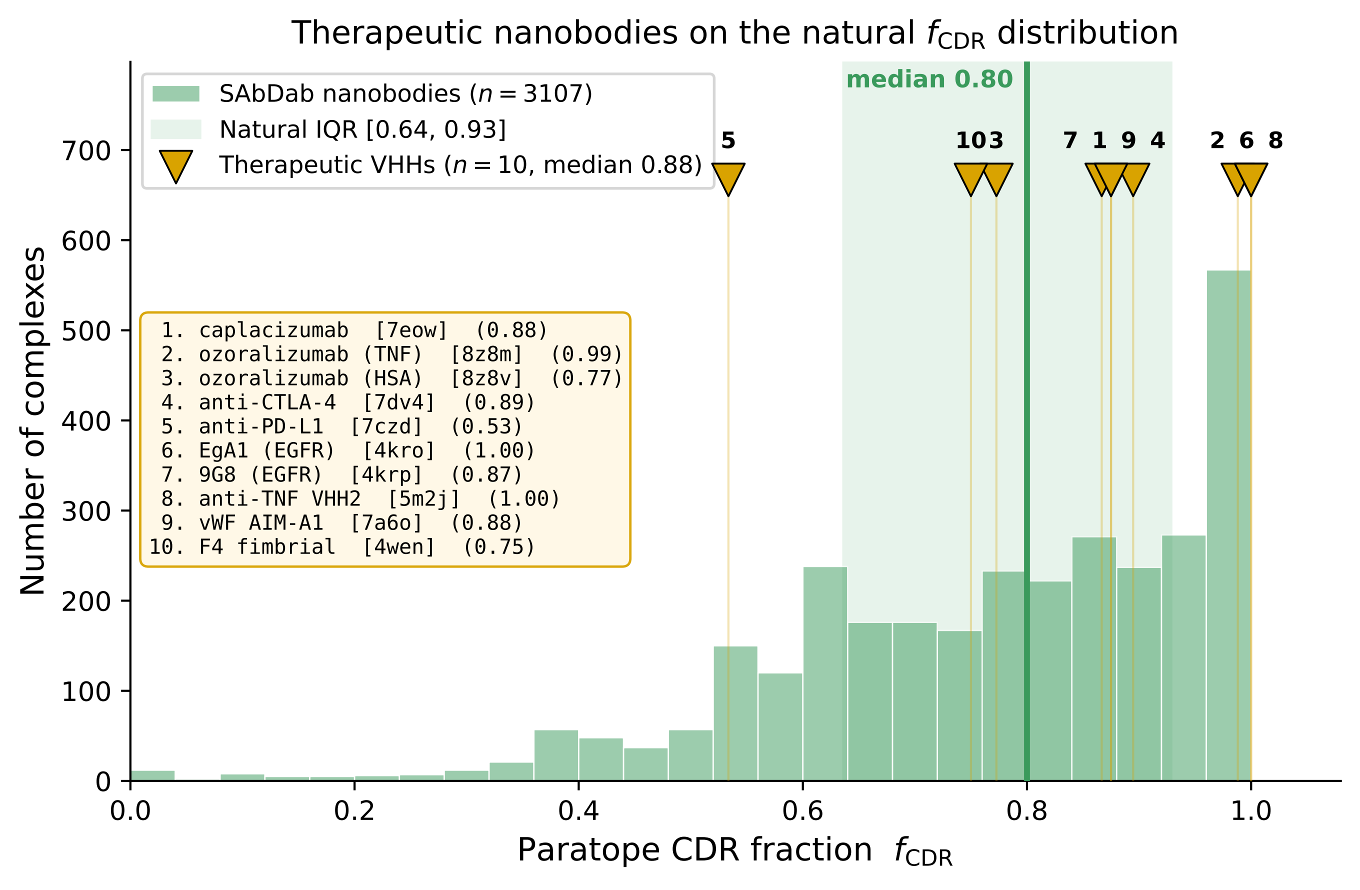

### sabdab_fcdr_therapeutic_cb8.png

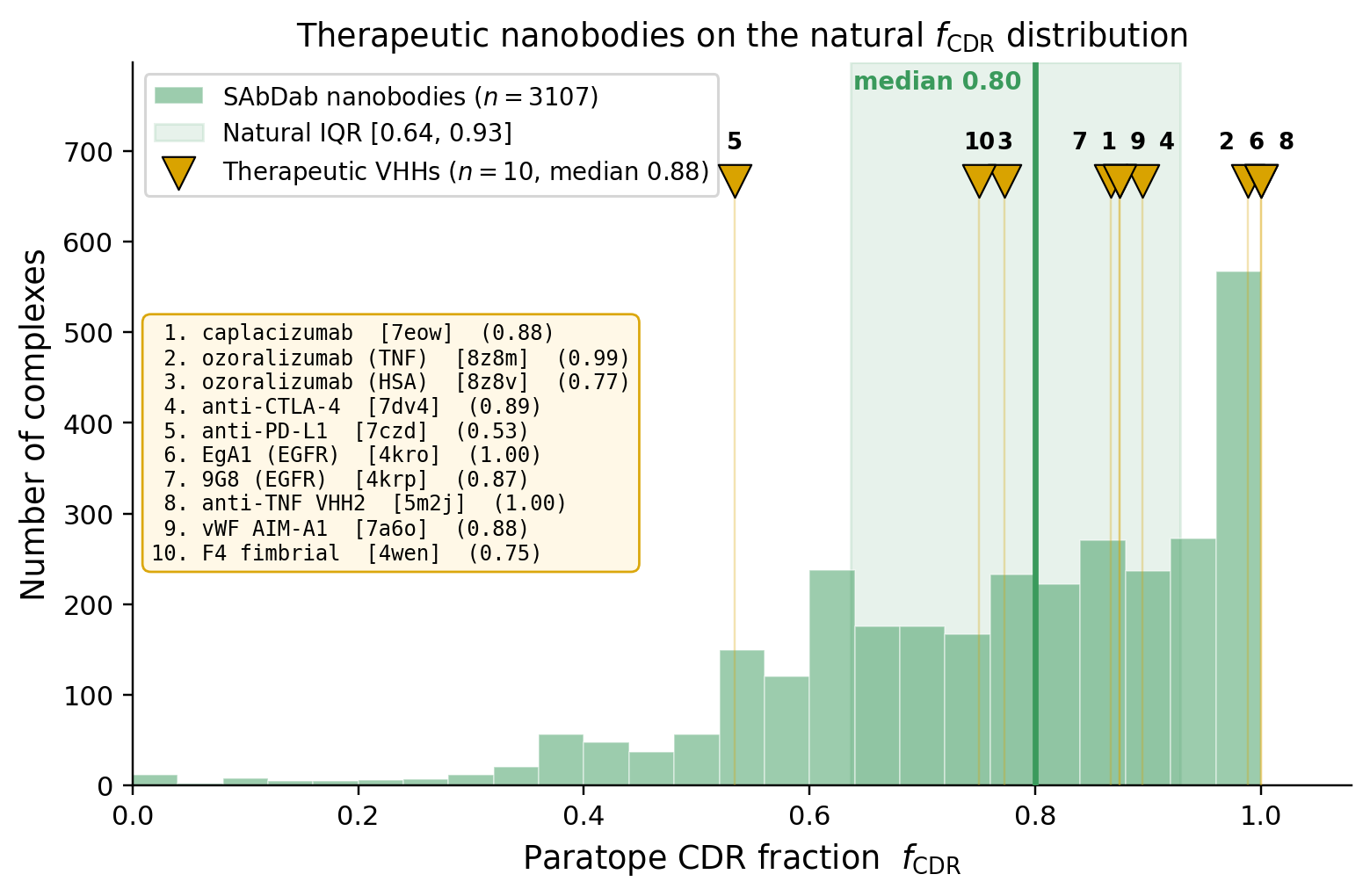

### sabdab_fcdr_therapeutic_cb.pdf

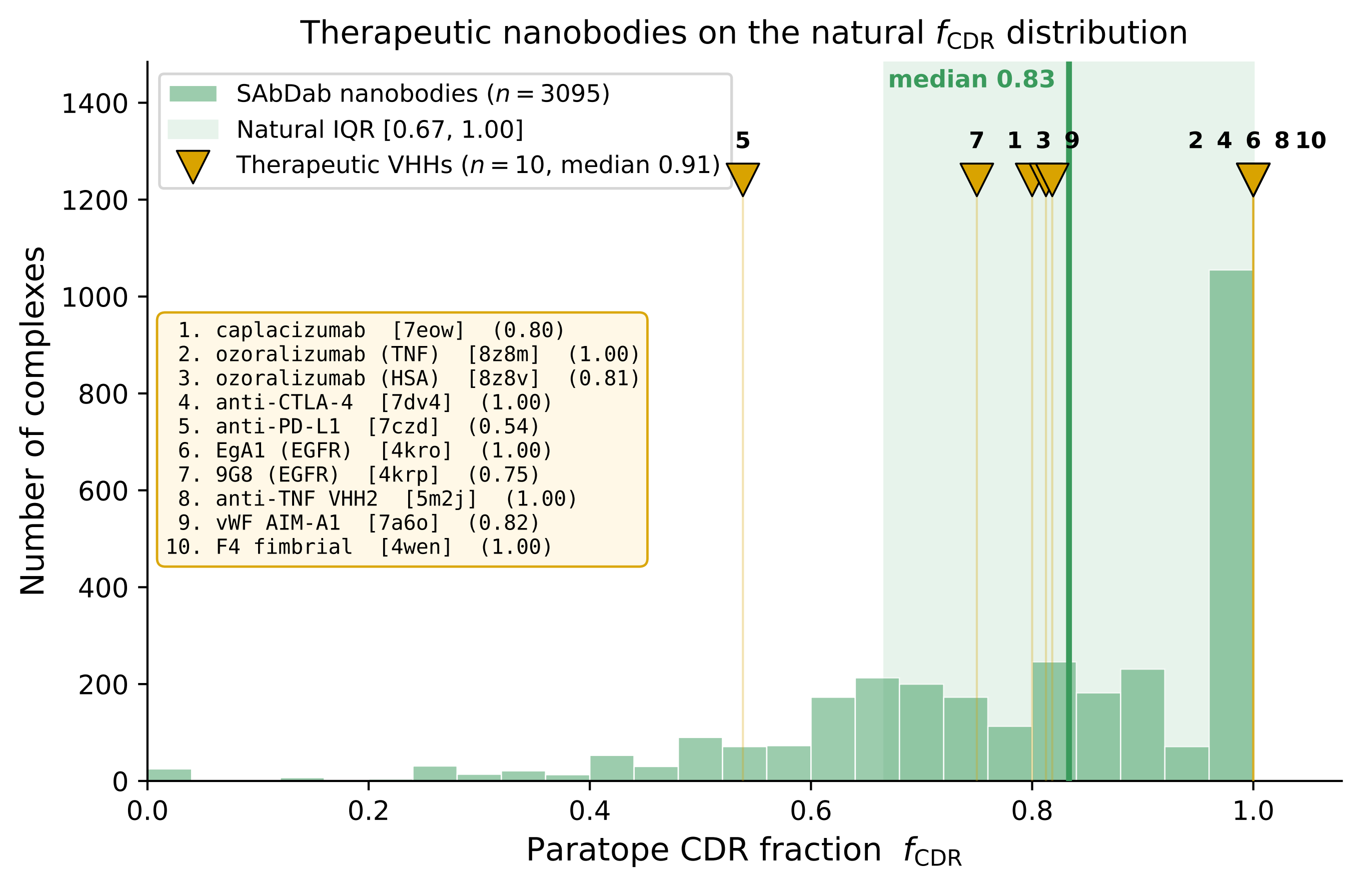

### sabdab_fcdr_therapeutic_cb.png

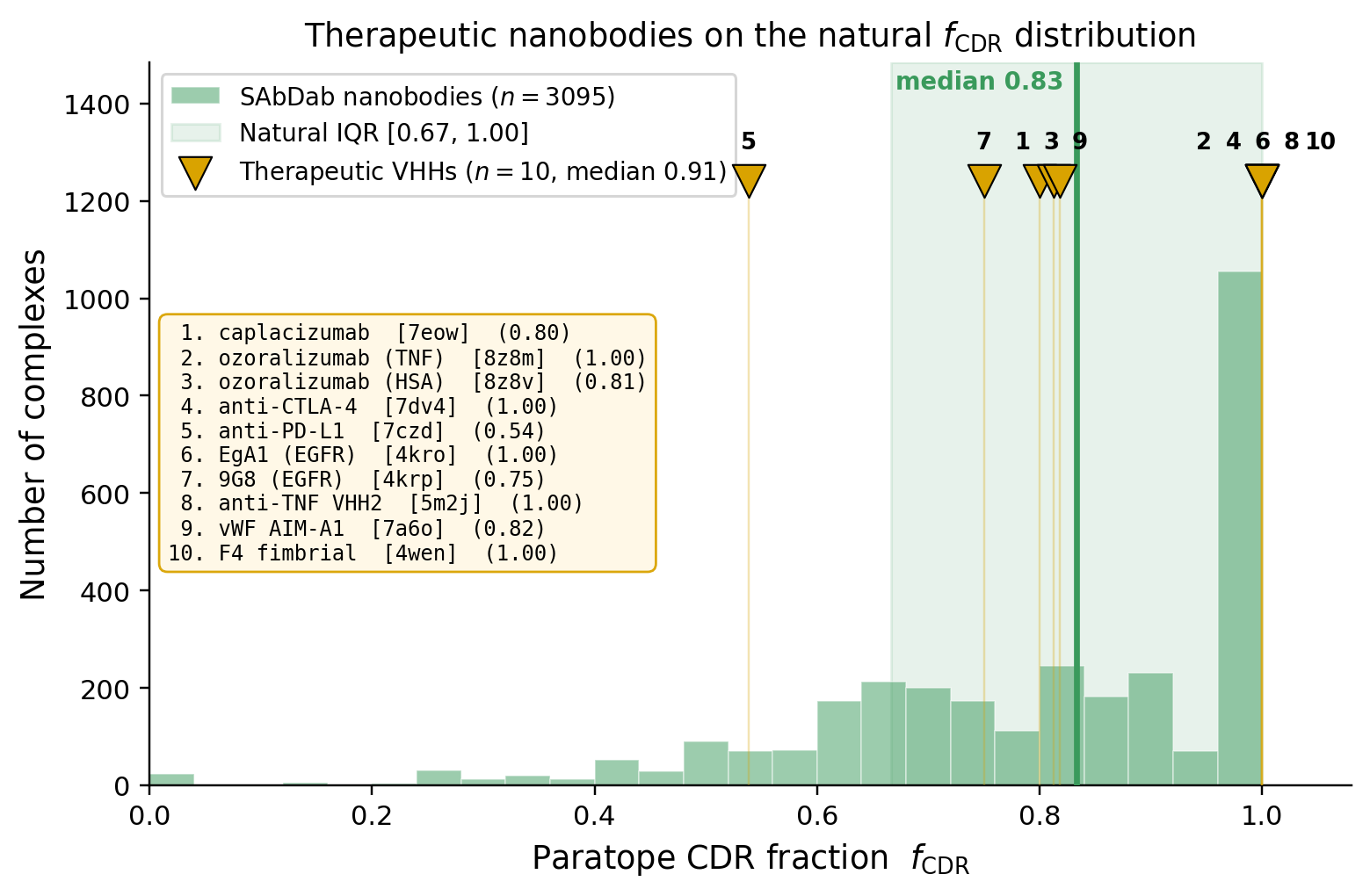

### sabdab_fcdr_therapeutic_cb_models.pdf

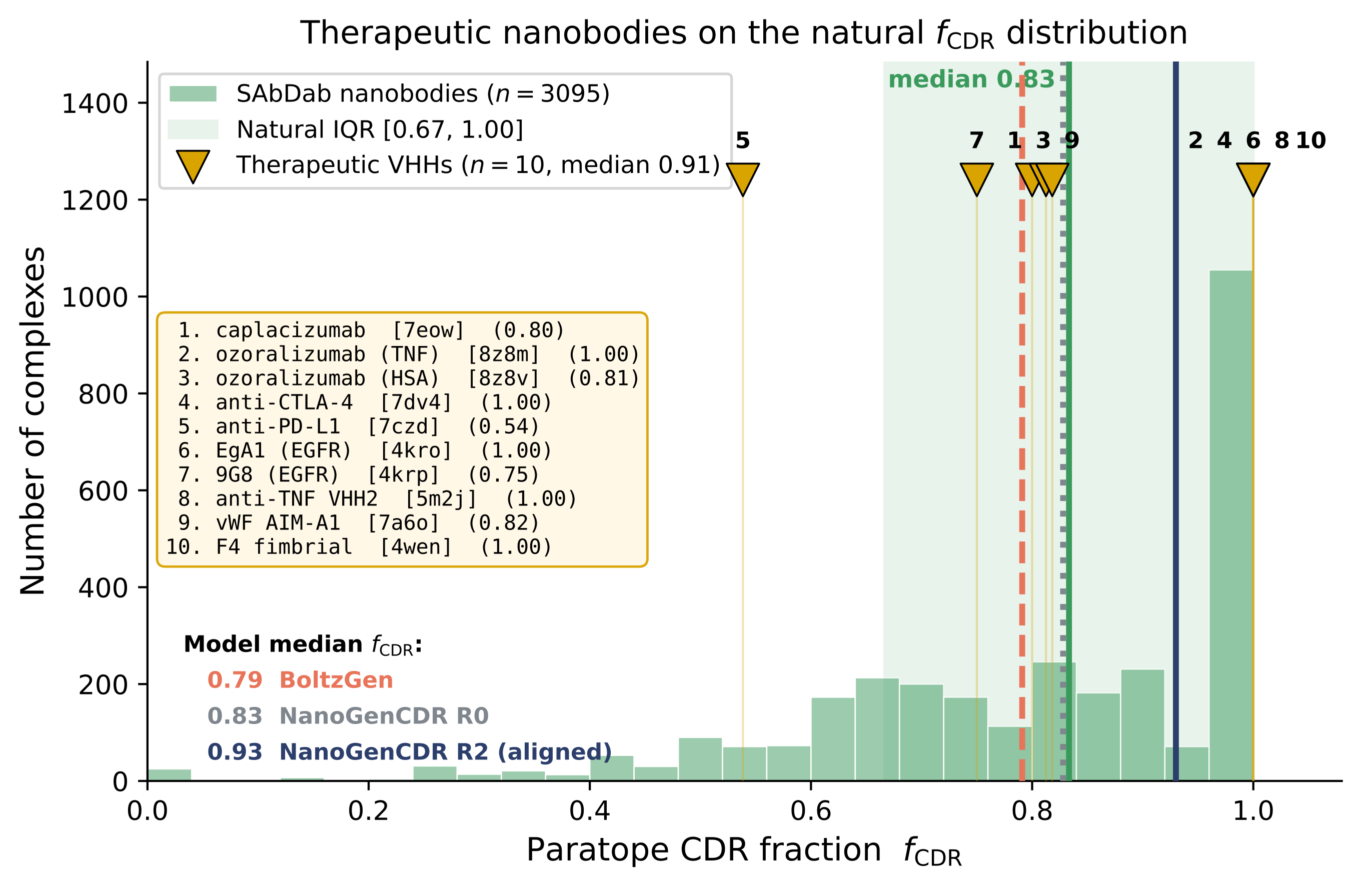

### sabdab_fcdr_therapeutic_cb_models.png

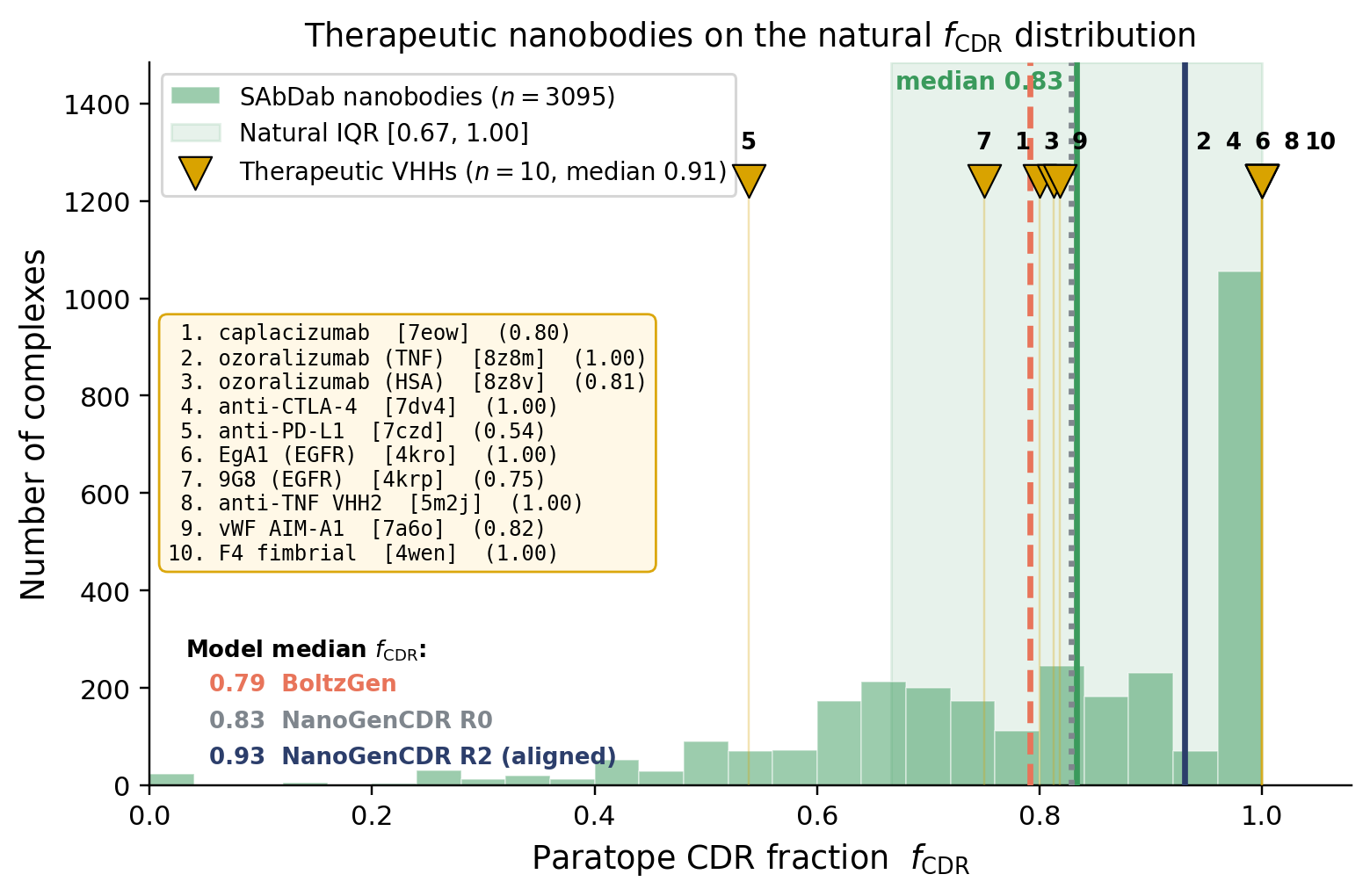

### sabdab_fcdr_therapeutic_models.pdf

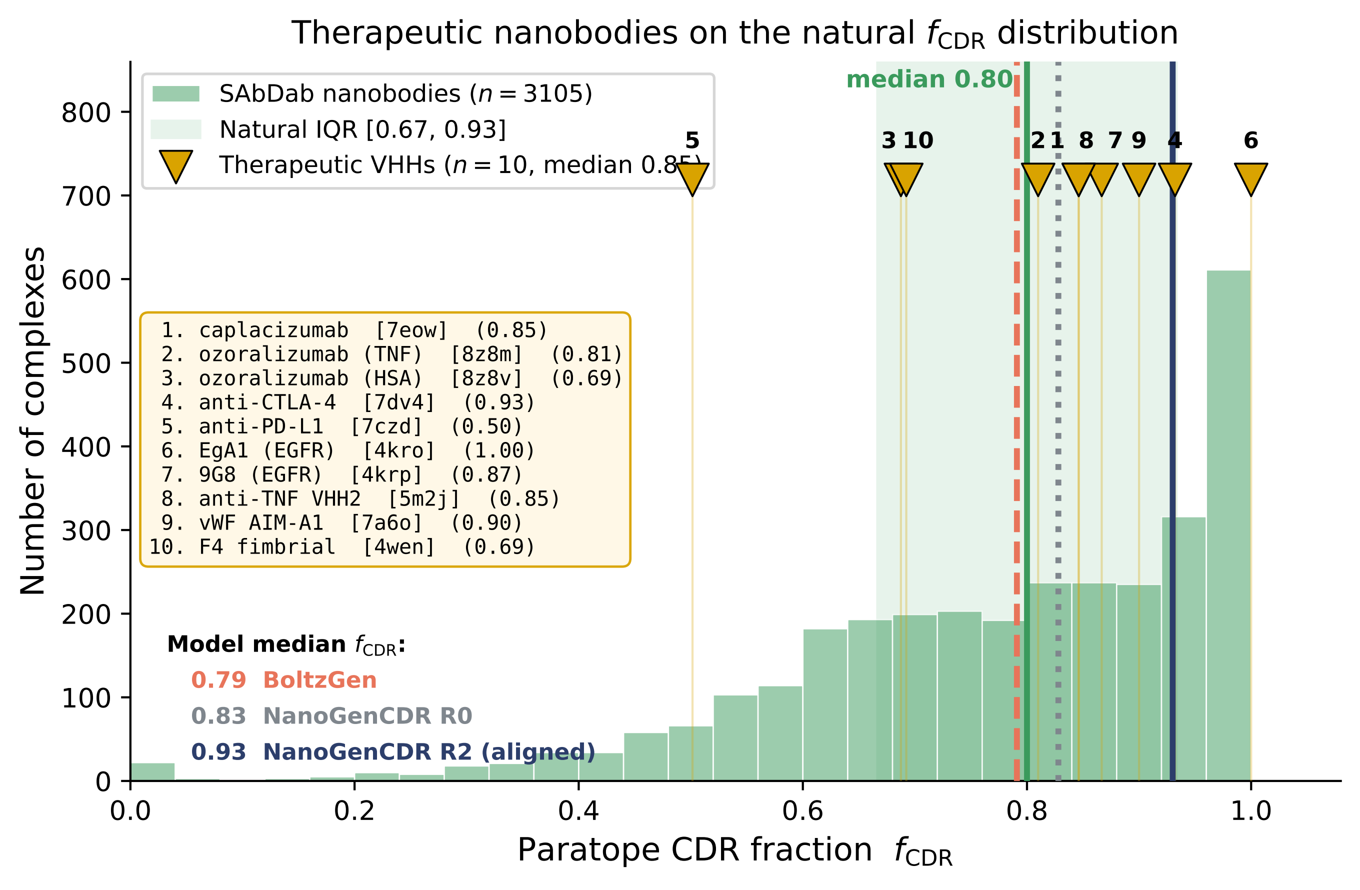

### sabdab_fcdr_therapeutic_models.png

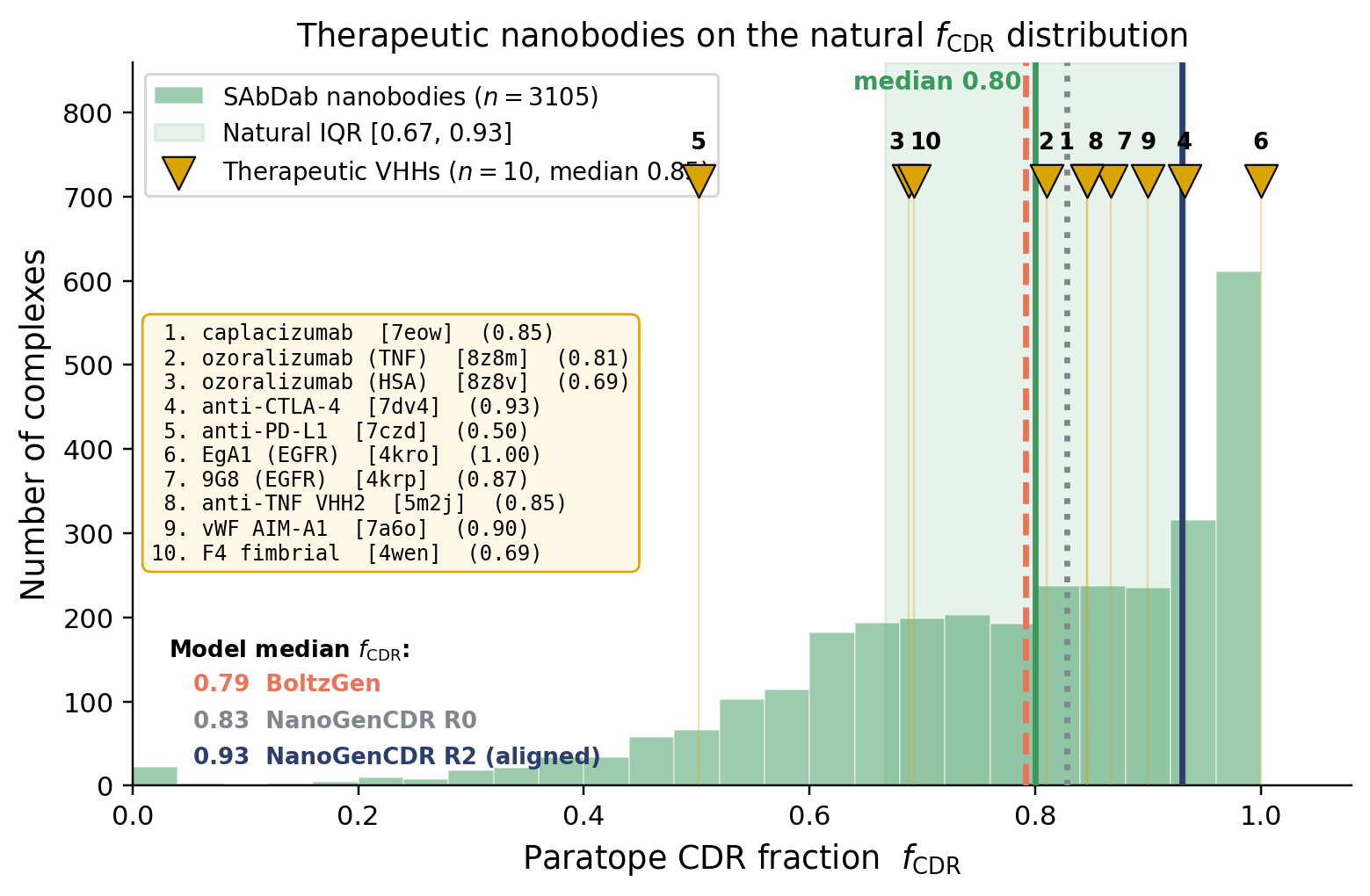

### temp_sweep_esm2.pdf

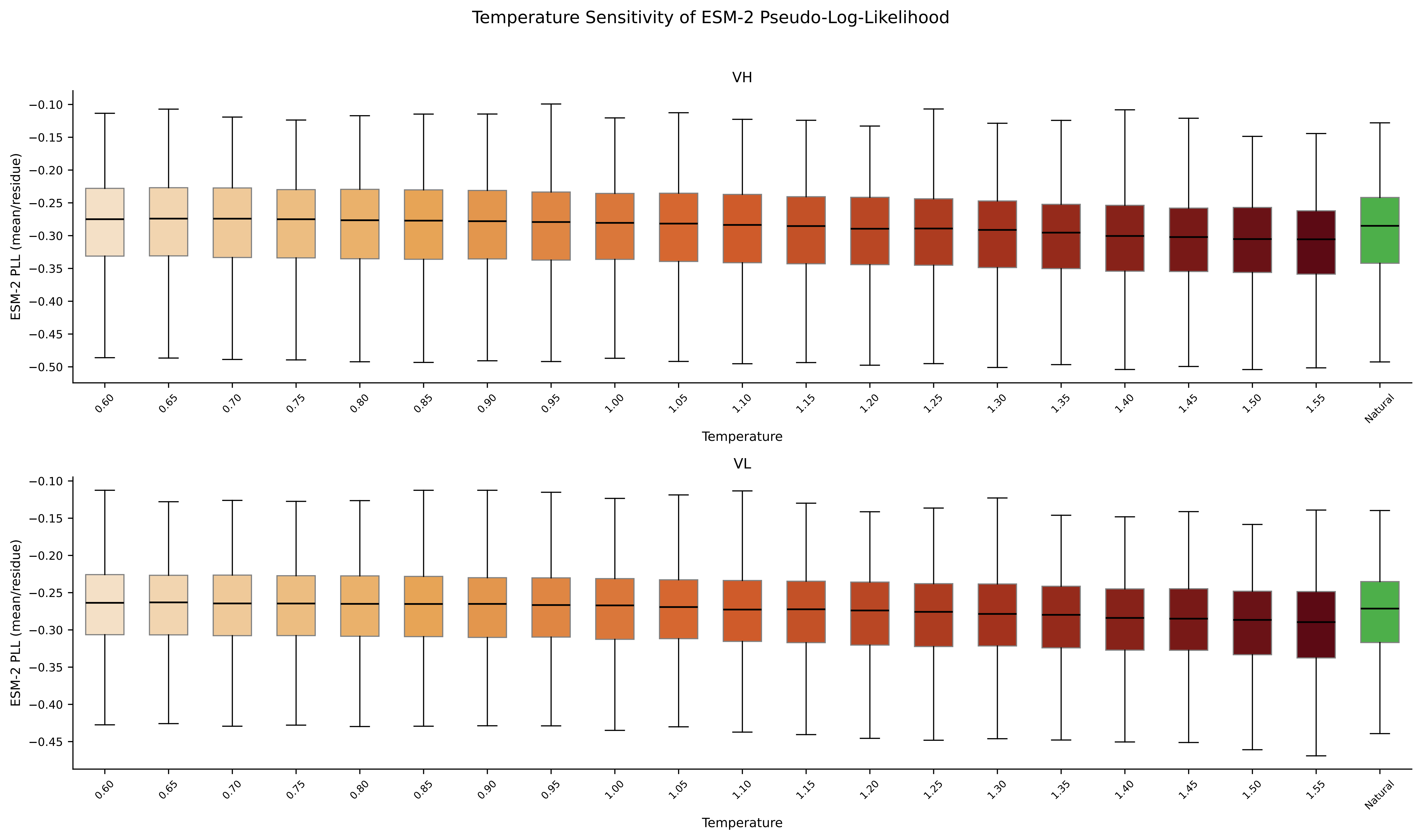

### temp_sweep_novelty.pdf

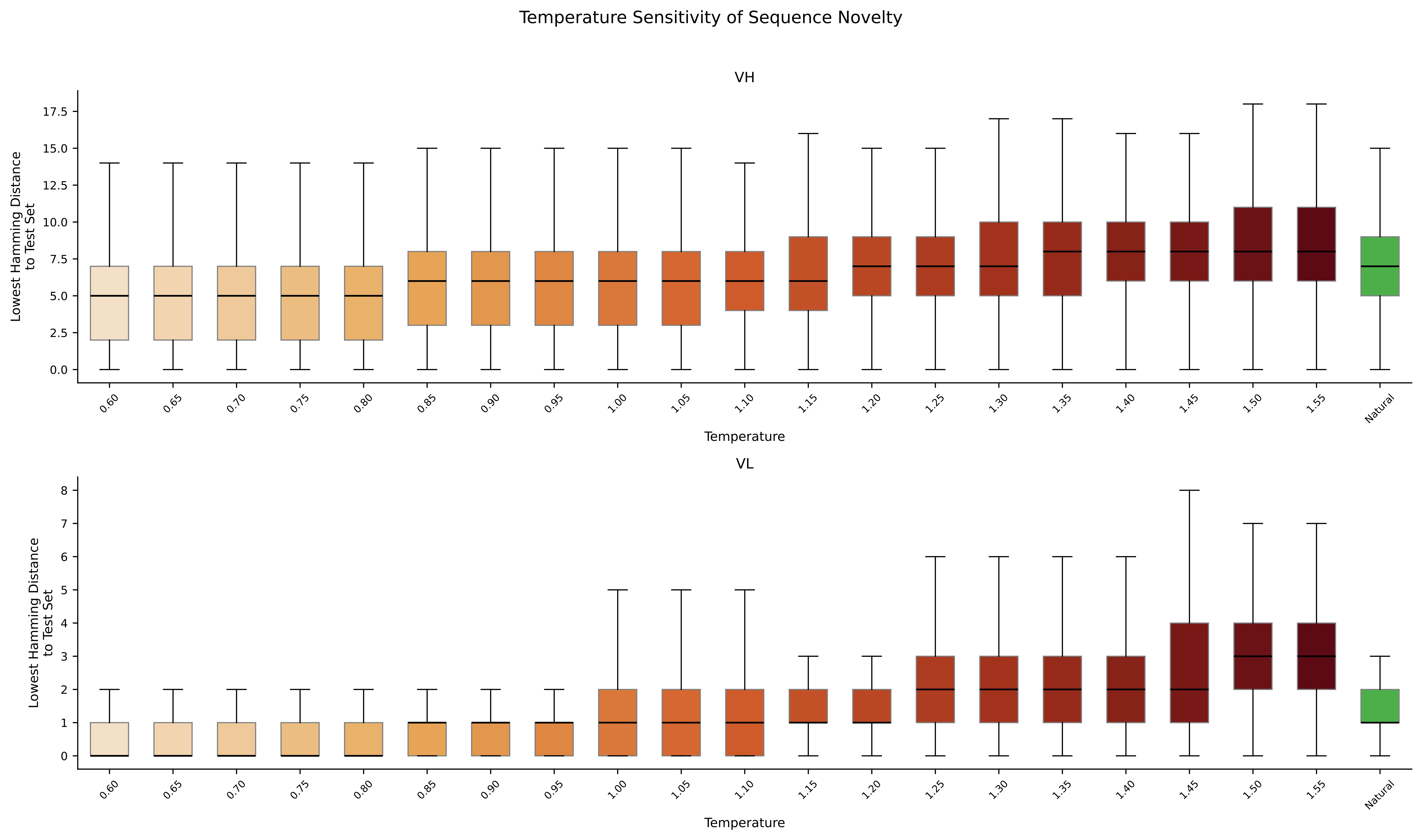

### temp_sweep_oasis.pdf

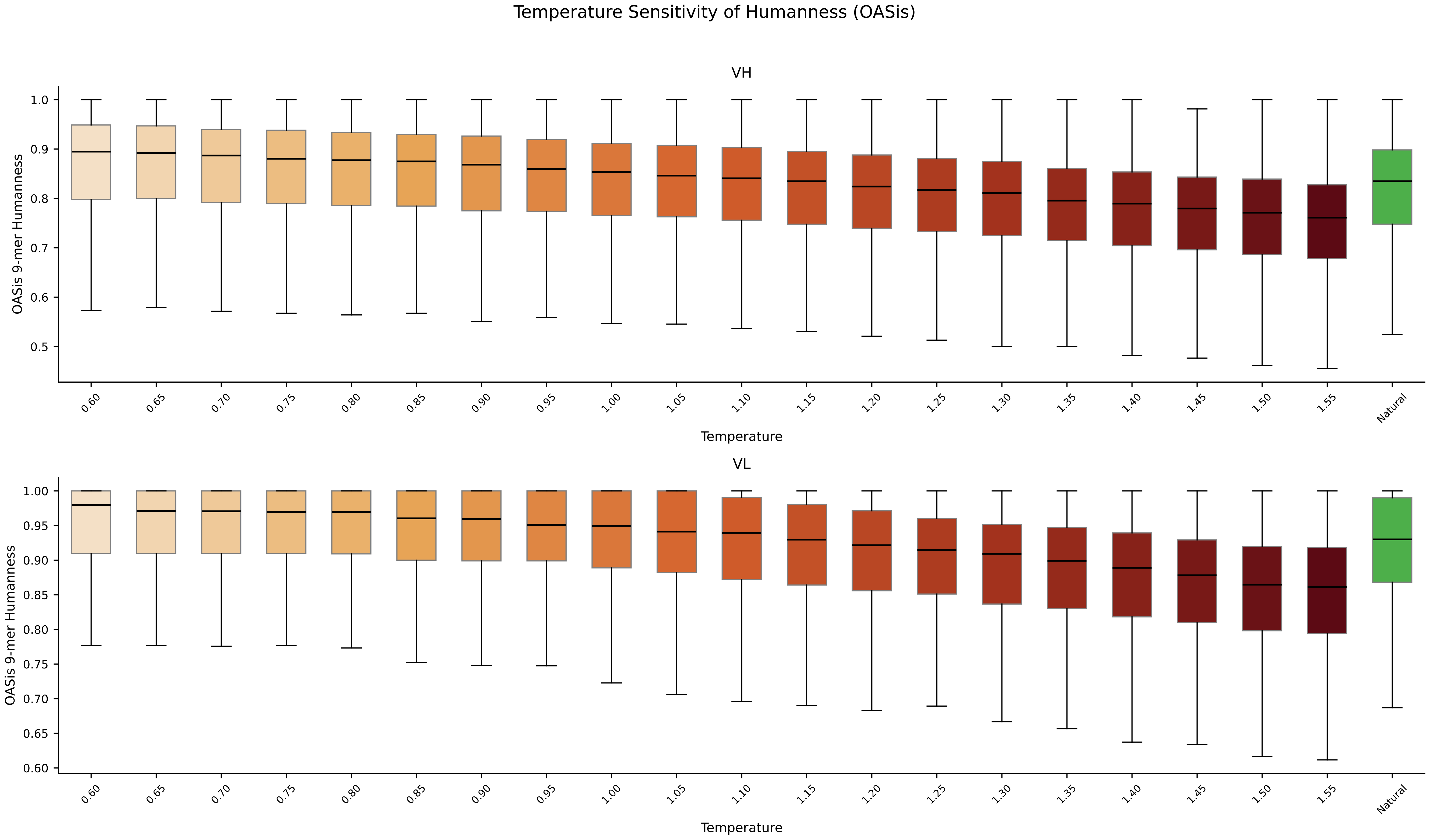
