## Supplementary Split for "Generative Language Modeling for Antibody CDR Grafting and Alignment-driven De Novo Design"

### Supplementary Note S3: Architecture and training hyperparameters

Table S1 lists the full architecture and training hyperparameters, complementing the summary in the main text. All variants share the same architecture, differing only in the fine-tuning corpus. AdamW used $\beta_1$=0.9, $\beta_2$=0.999 and a weight decay of 0.1, with gradient norms clipped to 1.0. The effective batch size of 1,024 was reached on 8$\times$A100 (40 GB) GPUs under Distributed Data Parallel with a per-GPU batch size of 32 and 4 gradient accumulation steps.

[3]https://naturalantibody.com/indi2/

DRAFT

| Hyperparameter | Value |
|---|---|
| Layers | 4 |
| Hidden dimension | 768 |
| Feed-forward size | 2,048 |
| Attention heads | 12 (4 key-value heads) |
| Context length | 1,024 tokens |
| Vocabulary | 48 (26 amino acids, 12 special, 10 reserved) |
| Parameters | ∼25M |
| Optimiser | AdamW ($\beta_1$=0.9, $\beta_2$=0.999) |
| Weight decay | 0.1 |
| Peak learning rate | $2\times10^{-4}$ |
| LR schedule | Cosine decay to 0 |
| Warm-up steps | 3,000 (linear) |
| Gradient norm clipping | 1.0 |
| Precision | BF16 (automatic mixed precision) |
| Hardware | 8×A100 (40 GB), DDP |
| Per-GPU batch size | 32 |
| Gradient accumulation steps | 4 |
| Effective batch size | 1,024 |
| Unpaired pre-training epochs | 10 |
| Paired fine-tuning epochs | 5 |
| Nanobody fine-tuning epochs | 5 |

**Table S1.** Architecture and training hyperparameters for all IgGenCDR variants. All variants share the LLaMA-style architecture above. Both p-IgGenCDR and NanoGenCDR were initialised from the IgGenCDR (unpaired) checkpoint and fine-tuned for 5 epochs on paired VH–VL sequences and the camelid–human mixture respectively.

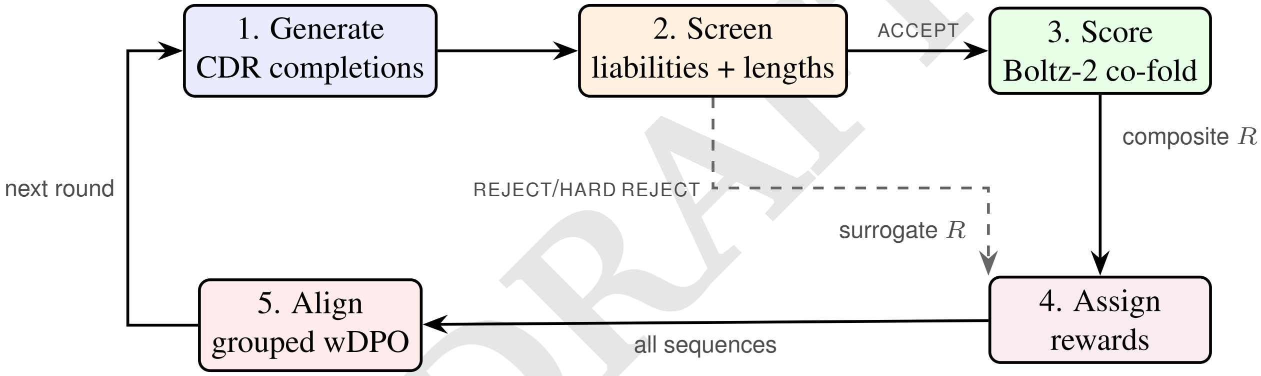


**Fig. S1.** Iterative alignment loop. ACCEPT sequences are co-folded and receive the composite reward; REJECT/HARD REJECT sequences bypass co-folding and receive surrogate rewards from calibrated bands. The updated policy generates CDR sequences for the next round.

DRAFT

### Supplementary Note S4: Antigen-driven alignment: methods and details

Full methodological detail for the antigen-driven alignment experiment (Section D).

**A. Pre-screening and reward calibration.** Figure S1 illustrates the complete iterative alignment loop.
Before any structural evaluation, generated CDR sequences are gated to scaffold-compatible length ranges (CDR1: 6–10, CDR2: 6–10, CDR3: 14–20 residues, IMGT) and screened against six sequence-level developability liability rules, adapted from the motif set of the Blaze antibody sequence-analysis platform and consistent with established developability rules (32) (Table S2), both shared with the BoltzGen reference (Section E). Sequences passing all checks are labelled ACCEPT and proceed to Boltz-2 co-folding, the only sequences to incur that cost. Those failing one or more rules are labelled REJECT or HARD REJECT according to the number of violations.
Rejected sequences are still included in the DPO signal so the model learns that liability-prone regions yield low reward, but they bypass co-folding and instead receive surrogate rewards sampled uniformly from bands placed below the ACCEPT distribution. The REJECT band spans the two robust standard deviations immediately below the 1st percentile of the initial ACCEPT rewards, and the HARD REJECT band the two below that, where the robust standard deviation is 1.4826 times the median absolute deviation of the ACCEPT rewards. These bounds are calibrated once on the first round and fixed thereafter, keeping the surrogate rewards close enough to provide a gradient signal yet separated enough not to confound the reward landscape.

| Rule | Scope | Motif / condition |
|---|---|---|
| `unpaired_cys_chain` | Full chain | Odd number of cysteines |
| `glycosylation_cdrs` | CDRs only | N-X-[ST] sequon (X $\neq$ P) |
| `invalid_characters_chain` | Full chain | Any non-standard amino acid character |
| `deamidation_high_ng` | Full chain | NG dipeptide (high deamidation risk) |
| `deamidation_medium_nsn` | Full chain | NS or NN dipeptide (medium deamidation risk) |
| `asp_isomerization_dsg` | Full chain | DS or DG dipeptide (aspartate isomerisation risk) |

**Table S2.** Liability rules applied during pre-screening, adapted from the Blaze platform and consistent with established developability motifs (32). A sequence failing any single rule is rejected.

**B. Composite reward: metric definitions.** Formal definitions of the five terms in Eq. 2, all evaluated on Boltz-2 (6) co-folded nanobody–antigen predictions.

**Interface confidence** (ipSAE): the interaction prediction Score from Aligned Errors (20), computed from the PAE matrix restricted to inter-chain residue pairs. Higher values indicate greater confidence in the predicted binding interface.

**Chain TM-score** ($\mathrm{pTM}_{\mathrm{chain}}$): predicted TM-score restricted to the nanobody chain, measuring internal fold quality independently of the interface.

**Paratope CDR fraction** ($f_{\mathrm{CDR}}$): with paratope residues defined as nanobody residues having any heavy atom within 4 Å of any antigen heavy atom and classified as CDR or framework by IMGT boundaries, $f_{\mathrm{CDR}} = C/(C+F)$ where $C$ and $F$ are CDR and framework contact counts.

**CDR geometry penalties** ($p_h$, $p_s$): adapted from the $\alpha$-helical and $\beta$-strand distogram losses in Germinal (10), converted to post-hoc metrics on predicted structures. The $\alpha$-helix propensity $p_h$ is the fraction of CDR-internal $(i, i{+}3)$ C$\alpha$ pairs within the helix window:

$$\begin{aligned} p_h &= \frac{1}{|\mathcal{P}_h|} \sum_{(i,i+3)\in\mathcal{P}_h} \mathbb{1}[2.0 \le d_i \le 6.2], \\ \mathcal{P}_h &= \{(i,i{+}3) \mid m_i^{\mathrm{CDR}}{=}1,\ m_{i+3}^{\mathrm{CDR}}{=}1\}. \end{aligned} \tag{S1}$$

*$\beta$-strand propensity* ($p_s$): the same calculation with a one-residue-expanded CDR mask (to capture CDR–framework junctions) and the strand window:

$$\begin{aligned} p_s &= \frac{1}{|\mathcal{P}_s|} \sum_{(i,i+3)\in\mathcal{P}_s} \mathbb{1}[9.75 \le d_i \le 11.5], \\ \mathcal{P}_s &= \{(i,i{+}3) \mid \tilde{m}_i^{\mathrm{CDR}}{=}1,\ \tilde{m}_{i+3}^{\mathrm{CDR}}{=}1\}, \end{aligned} \tag{S2}$$

where $d_i = \|x_{i+3} - x_i\|_2$ and $\tilde{m}_i^{\mathrm{CDR}} = \max(m_{i-1}^{\mathrm{CDR}}, m_i^{\mathrm{CDR}}, m_{i+1}^{\mathrm{CDR}})$. The distance windows and mask expansion follow Mille-Fragoso et al. (10).

**C. Scaffold-grouped weighted DPO: derivation and implementation details. Single-trajectory alignment.** Standard DPO (13) operates on preference pairs $(x, y_w, y_l)$. When continuous scalar rewards are available, constructing preference pairs introduces information loss. The single-trajectory setting (33) instead works with triplets $(x, y, R)$ and derives the optimal KL-regularised policy directly. Weighted DPO (11, 19) implements this by replacing the pairwise sigmoid with a listwise cross-entropy over softmax-normalised rewards (Eq. 1), avoiding the need for an auxiliary value function. Alternative single-trajectory formulations exist (e.g. DRO (33)) and a comparison is left to future work.

**Completion-only log-ratio.** The sequence-level log-ratio in Eq. 1 is computed as a mean over CDR completion tokens only, those after the `<HPRED>` (or `<LPRED>`) boundary through `<EOS>`, including inter-CDR `<SEP>` tokens. Framework tokens are excluded since they form the fixed prompt:

$$\log \frac{\pi_\theta(y_i \mid x_i)}{\pi_{\mathrm{ref}}(y_i \mid x_i)} = \frac{1}{|C_i|} \sum_{t\in C_i} \left[\log \pi_\theta(y_{i,t} \mid x_i, y_{i,<t}) - \log \pi_{\mathrm{ref}}(y_{i,t} \mid x_i, y_{i,<t})\right] \tag{S3}$$

where $C_i$ is the set of CDR completion token positions for sequence $i$. This decomposition follows directly from the framework-first input ordering.

DRAFT

**Scaffold grouping.** Each batch contains CDR completions from all four framework scaffolds. The listwise softmax of Eq. 1 is meaningful only among completions of a shared prompt: the $\beta$-scaled log-ratios $s_k$ are comparable across responses to the same scaffold, but not across different scaffolds, whose frameworks impose different baseline likelihoods. We therefore compute the softmax targets $w_k$ and logits $s_k$ within each scaffold group, never across the full batch. This grouping is what makes Eq. 1 well defined, rather than a post-hoc variance-reduction step.
The per-scaffold losses are then combined by group fraction $|g|/B$, each group $g$ collecting the $|g|$ responses to one scaffold prompt:

$$\mathcal{L} = \sum_{g \in \mathcal{G}} \frac{|g|}{B}\, \mathcal{L}^{(g)}_{\text{wDPO}}, \tag{S4}$$

with $B = \sum_g |g|$ the batch size. Because each round generates comparable numbers of accepted completions per scaffold, group sizes are near-balanced ($|g| \approx B/|\mathcal{G}|$), so the group-fraction weighting reduces to a uniform average over scaffolds ($|g|/B \approx 1/|\mathcal{G}|$) and no scaffold is systematically up-weighted. We adopt $|g|/B$ as it requires no assumption of exact balance while reducing to the uniform weighting whenever balance holds. Figure S3 confirms that all four scaffolds improve in lockstep, with none dominating the update. A systematic comparison of the two aggregations under strong scaffold imbalance is left to future work.

**Choice of $\beta$.** We set $\beta = 0.15$, more conservative than the 0.01–0.1 used for single-property protein alignment (11), for three reasons: (i) the multi-CDR action space amplifies the risk of degenerate drift, (ii) the composite reward is derived from stochastic Boltz-2 predictions and is therefore noisy, (iii) we aim to preserve pre-trained CDR length and composition distributions while steering only interface-relevant properties.

**Optimisation.** Each round performs 10 epochs of wDPO on that round's generated set with AdamW (learning rate $5 \times 10^{-5}$, weight decay 0.01), a batch size of 512, gradient-norm clipping at 1.0, and 16-bit mixed precision, on a single A100 GPU. A stratified 20% of each round's data, by reward, is held out for validation. The reference policy $\pi_{\text{ref}}$ is the frozen pre-aligned NanoGenCDR checkpoint and is kept fixed across all four rounds.

**D. Boltz-2 co-folding inference settings.** All structural evaluations in this work (both alignment rounds and BoltzGen reference designs) use the same Boltz-2 (6) co-folding configuration to ensure directly comparable scores (Table S3). No template structure is provided for the antigen, only an MSA is supplied for the antigen sequence (empty for nanobody inputs).

| Parameter | Value |
|---|---|
| Recycling steps | 3 |
| Diffusion samples | 5 |
| Sampling steps | 200 |
| Step scale | 1.638 |
| Antigen template | None (MSA only) |
| Nanobody MSA | Empty |

**Table S3.** Boltz-2 co-folding inference settings used for all structural evaluations. Scores are aggregated by taking the average across the 5 diffusion samples.

DRAFT

**E. BoltzGen reference design methodology.** BoltzGen designs were generated against PD-L1 using the four nanobody scaffolds of Section D, following the protocol of Stark et al. (9): for each scaffold the framework structure and sequence are fixed while the three CDR regions are replaced with loops of random lengths within the shared constraints (Section A). We ran BoltzGen with `num_design = 20,000`, `diffusion_batch_size = 10`, `protocol = nanobody_anything`, `step_scale = 1.8`, and `noise_scale = 0.9`, inverse-folding each generated backbone into 4 sequences with the BoltzIF inverse-folding model (9), a soluble-protein inverse folder analogous to SolubleMPNN. Of the 80,000 candidates, 31.8% passed the shared liability and length filters from which we sampled 20,000 ACCEPT sequences for co-folding. These were co-folded with Boltz-2 under the shared settings (Table S3, ≈48 h on 8×A100), with structural and interface metrics computed identically to the alignment pipeline.

**F. Alignment naturalness monitoring.** Table S4 reports the full per-round distributions (median and IQR) of every reward component, the composite reward, and the reference policy log-likelihood summarised in the main text (Section D). To monitor policy drift during alignment, we track the reference policy log-likelihood (mean per-token log-probability under the frozen NanoGenCDR checkpoint) and AbLang pseudo-log-likelihood (30) across rounds (Figure S2).
The round-by-round trajectory resolves the drift underlying the reward plateau. The reference log-likelihood decreases modestly from round 0 (−0.837) to round 2 (−0.883) as the policy diverges to push reward, then recovers in round 3 (−0.845), tracking the reward plateau (Table S4). The conservative $\beta$=0.15 therefore bounds drift, with the policy exploring early and being

| Source | ipSAE | $\mathrm{pTM_{chain}}$ | $f_{\mathrm{CDR}}$ | $1-p_h$ | $1-p_s$ | **Reward** | Ref LL |
|---|---|---|---|---|---|---|---|
| Round 0 ($n$=5K) | 0.404 [0.287, 0.520] | 0.973 [0.967, 0.977] | 0.828 [0.713, 0.916] | 0.778 [0.744, 0.811] | 0.817 [0.783, 0.850] | 0.715 [0.674, 0.754] | **−0.837** [−0.890, −0.778] |
| Round 1 ($n$=5K) | 0.489 [0.385, 0.583] | 0.973 [0.967, 0.977] | 0.931 [0.879, 0.973] | 0.778 [0.744, 0.822] | 0.833 [0.808, 0.858] | 0.766 [0.731, 0.797] | −0.869 [−0.916, −0.818] |
| Round 2 ($n$=5K) | **0.530** [0.422, 0.623] | 0.975 [0.970, 0.978] | **0.943** [0.894, 0.985] | 0.778 [0.744, 0.811] | 0.833 [0.808, 0.863] | **0.779** [0.744, 0.811] | −0.883 [−0.935, −0.834] |
| Round 3 ($n$=5K) | 0.523 [0.418, 0.619] | 0.975 [0.970, 0.978] | 0.930 [0.870, 0.973] | 0.778 [0.744, 0.811] | 0.833 [0.808, 0.867] | 0.775 [0.738, 0.807] | −0.845 [−0.889, −0.799] |
| BoltzGen ($n$=20K) | 0.474 [0.343, 0.598] | **0.978** [0.972, 0.981] | 0.791 [0.619, 0.905] | **0.811** [0.756, 0.867] | 0.758 [0.708, 0.800] | 0.719 [0.660, 0.768] | −1.020 [−1.091, −0.956] |

**Table S4.** Median [IQR] of reward components, composite reward (Eq. 2), and reference policy log-likelihood across alignment rounds (ACCEPT sequences only). Both pipelines use identical Boltz-2 co-folding settings and scoring (Table S3). $f_{\mathrm{CDR}}$: paratope CDR fraction; $1-p_h/1-p_s$: CDR geometry penalties (higher=less regular secondary structure). Ref LL: mean per-token log-probability under the frozen NanoGenCDR reference policy (not a reward component; monitors policy drift). **Bold**: best value in each column.

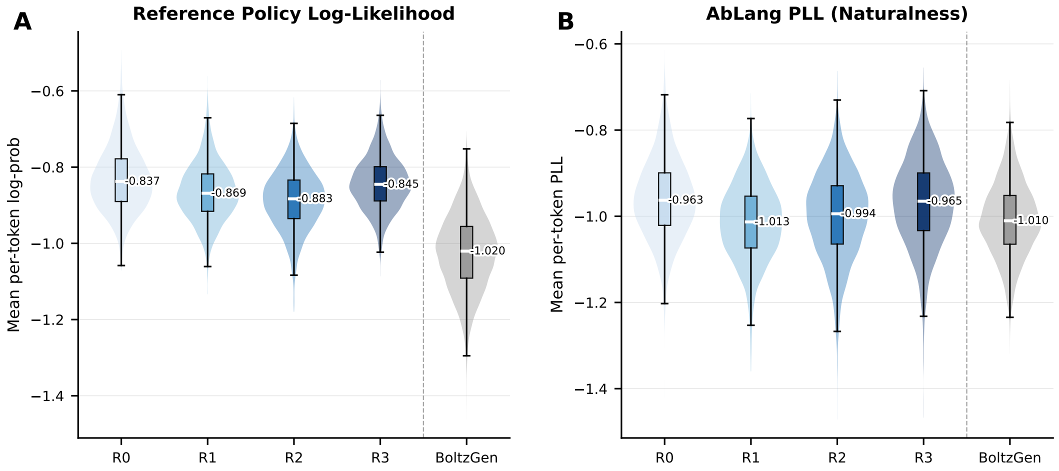


**Fig. S2.** Naturalness monitoring across alignment rounds. **(A)** Reference policy (NanoGenCDR) log-likelihood of generated CDR sequences. **(B)** AbLang pseudo-log-likelihood (30), an independent antibody-specific naturalness metric. Violins show full distributions; boxes indicate IQR with median (white line). Annotated values are medians. Dashed vertical line separates alignment rounds from BoltzGen.

pulled back toward the reference as the KL penalty grows. For reference, BoltzGen designs score $-1.020$ (NanoGenCDR log-likelihood) and $-1.010$ (AbLang PLL), below every alignment round.

DRAFT

**G. Tail analysis: top-5% designs.** This section details the tail analysis summarised in the main text (Section D). To mirror how a campaign triages designs, we rank all ACCEPT designs of each pipeline by the composite reward, take the top $5\%$, and report the mean of every metric over that single selected cohort (Table S5). Ranking by a single criterion, rather than per metric, means the reported values are jointly realised by the same set of designs. The round-2 threshold defining this cohort is a composite reward of $0.847$, which only $0.58\%$ of round-0 designs meet.

| Source | ipSAE | $\mathrm{pTM_{chain}}$ | $f_{\mathrm{CDR}}$ | $1-p_h$ | $1-p_s$ | Reward |
|---|---|---|---|---|---|---|
| Round 0 ($n$=252) | 0.683 | 0.979 | 0.915 | 0.788 | 0.831 | 0.826 |
| Round 1 ($n$=251) | 0.716 | 0.979 | 0.960 | 0.797 | 0.839 | 0.848 |
| Round 2 ($n$=250) | **0.745** | 0.980 | **0.966** | 0.796 | 0.842 | **0.859** |
| Round 3 ($n$=250) | 0.741 | 0.981 | 0.961 | 0.794 | **0.849** | 0.857 |
| BoltzGen ($n$=1000) | 0.741 | **0.981** | 0.929 | **0.827** | 0.775 | 0.845 |

**Table S5.** Tail analysis: designs ranked by composite reward, top $5\%$ selected, and the mean of every metric reported over that single selected cohort (ACCEPT sequences only), complementing the median comparison of Table S4. Alignment improves the selected cohort across rounds, and at equal $5\%$ selection the aligned round-2 cohort matches BoltzGen on ipSAE and leads on composite reward and $f_{\mathrm{CDR}}$. **Bold**: best value in each column.

**H. Per-scaffold reward and length steering.** The main-text progression (Section D, Figure 4) pools the four scaffolds. Figure S3 disaggregates the composite reward by scaffold, with all four rising from a median of $\sim$0.71 at round 0 to $\sim$0.78 by round 2 and then plateauing.

Figure S4 disaggregates the CDR3 length steering described in the main text (Section D) by scaffold, showing the broad round 0 distributions concentrating toward the longer loops at round 2 and tightening again at round 3 as the reward plateaus, consistently across all four scaffolds.

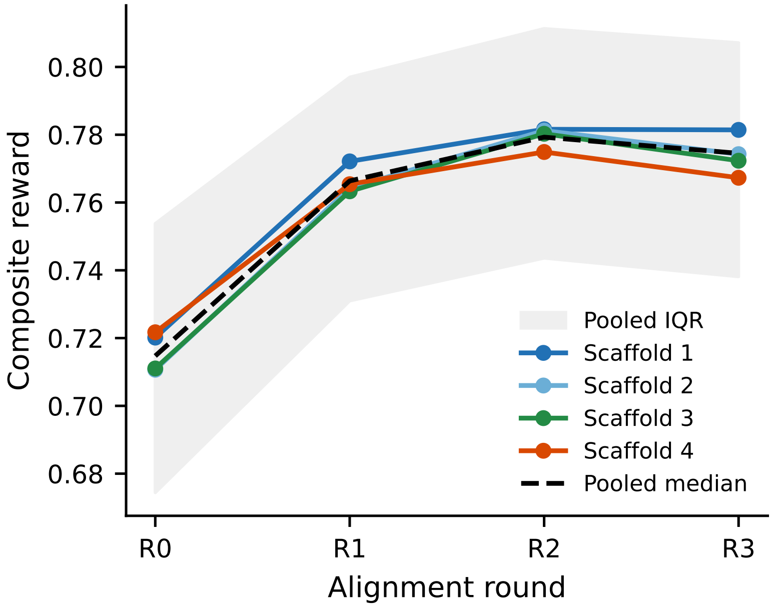


**Fig. S3.** Composite reward per scaffold across alignment rounds (median over accepted designs, shading shows the pooled interquartile range). All four scaffolds improve together and plateau after round 2, mirroring the pooled trend (dashed).

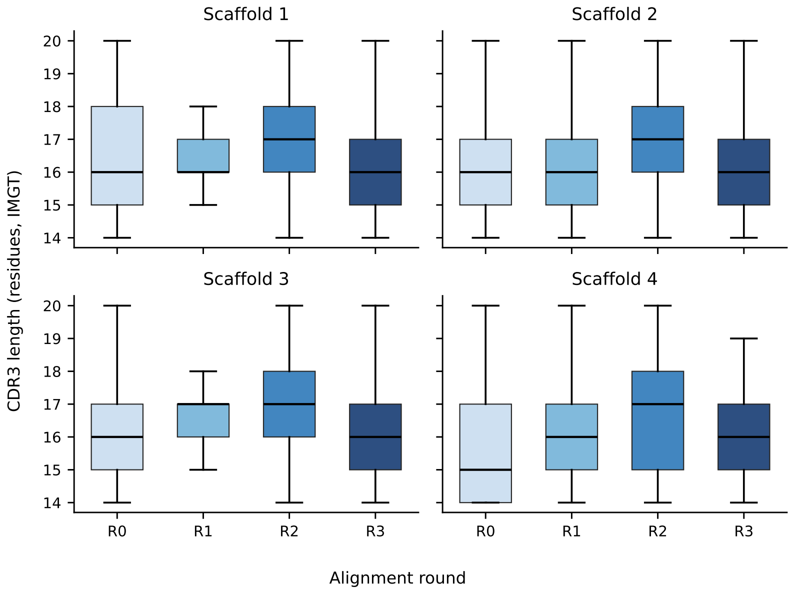


**Fig. S4.** CDR3 length distribution per scaffold across alignment rounds (boxes show IQR with median; whiskers to $1.5\times$IQR). Distributions shift toward longer loops at round 2, coincident with the reward peak, then partly relax at round 3.

### Supplementary Note S5: CDR amino acid recovery under alternative numbering schemes

Tables S6 and S7 reproduce the main-text CDR-disjoint recovery comparison (Table 1) under Kabat and Chothia CDR boundaries. The model ordering is preserved across schemes.

### Supplementary Note S6: CDR recovery: detailed baseline analysis

**Input-framing ablation.** With an identical 25 M-parameter architecture and training set, switching from left-to-right to framework-first ordering improves recovery on five of the six CDR loops. CDR1 benefits the most ($+2.91\%$ heavy, $+3.44\%$ light) and CDR3 improves moderately ($+0.99\%$ heavy, $+0.44\%$ light), whereas CDR2 barely moves ($-0.50\%$ heavy, $+0.10\%$ light). CDR1 appears before FR2–FR4 in the canonical left-to-right order and thus never sees the downstream frameworks that constrain it. The framework-first layout resolves this by conditioning CDR1 on all four frameworks. CDR2, by contrast, already receives most of its relevant context from FR1 and FR2 even under left-to-right decoding.

| Model | Heavy chain (%) CDR1 | Heavy chain (%) CDR2 | Heavy chain (%) CDR3 | Light chain (%) CDR1 | Light chain (%) CDR2 | Light chain (%) CDR3 |
|---|---|---|---|---|---|---|
| *Input-framing ablation* | | | | | | |
| IgGenCDR (framework-first) | **88.50 ± .04**$^{***}$ | 91.27 ± .03 | **44.81 ± .04**$^{***}$ | **88.75 ± .07**$^{***}$ | 87.58 ± .08 | **58.75 ± .08**$^{***}$ |
| IgGenCDR-LTR (left-to-right) | 85.56 ± .04 | **91.87 ± .03**$^{***}$ | 43.66 ± .05 | 86.34 ± .07 | **87.60 ± .09** | 58.10 ± .08 |
| *Autoregressive baselines* | | | | | | |
| p-IgGen (left-to-right) (4) | 81.16 ± .05 | 88.08 ± .03 | 34.24 ± .03 | 82.72 ± .08 | 82.02 ± .09 | 50.38 ± .08 |
| p-IgGen (right-to-left) (4) | 84.97 ± .05 | 85.93 ± .03 | 37.90 ± .04 | 84.72 ± .08 | 81.40 ± .09 | 42.73 ± .08 |
| p-IgGen (developable, left-to-right) (4) | 81.35 ± .05 | 87.77 ± .03 | 33.94 ± .03 | 82.47 ± .08 | 81.74 ± .09 | 50.14 ± .08 |
| p-IgGen (developable, right-to-left) (4) | 84.69 ± .05 | 85.70 ± .03 | 37.17 ± .04 | 84.61 ± .08 | 81.29 ± .09 | 42.52 ± .08 |
| ProGen2-OAS | 80.84 ± .05 | 88.34 ± .03 | 35.53 ± .04 | 60.29 ± .19 | 52.22 ± .19 | 39.38 ± .12 |
| ProGen2-small | 51.46 ± .06 | 61.86 ± .04 | 14.03 ± .02 | 50.90 ± .11 | 50.71 ± .11 | 24.56 ± .08 |
| ProGen2-base | 73.28 ± .06 | 84.01 ± .04 | 26.64 ± .03 | 70.63 ± .12 | 70.64 ± .12 | 40.66 ± .09 |
| ProGen2-large | 74.48 ± .06 | 83.89 ± .04 | 26.40 ± .03 | 75.79 ± .12 | 73.34 ± .12 | 40.12 ± .10 |
| IgLM (CDR infill) | 85.53 ± .05$^{\dagger}$ | 88.49 ± .03$^{\dagger}$ | 33.53 ± .03$^{\dagger}$ | 85.33 ± .08$^{\dagger}$ | 83.13 ± .09$^{\dagger}$ | 53.17 ± .08$^{\dagger}$ |
| IgLM (full LTR) | 81.81 ± .05 | 88.35 ± .03 | 23.24 ± .03 | 84.57 ± .08 | 83.25 ± .09 | 41.08 ± .08 |
| IgLM-S (CDR infill) | 83.51 ± .05$^{\dagger}$ | 87.69 ± .03$^{\dagger}$ | 33.83 ± .04$^{\dagger}$ | 83.76 ± .08$^{\dagger}$ | 81.59 ± .10$^{\dagger}$ | 49.34 ± .08$^{\dagger}$ |
| IgLM-S (full LTR) | 81.19 ± .05 | 87.91 ± .03 | 25.28 ± .04 | 84.09 ± .08 | 82.46 ± .09 | 40.86 ± .09 |

**Table S6.** CDR amino acid recovery (NTP accuracy, %, Kabat numbering) on the CDR-disjoint unpaired test set, following the protocol of the main-text Table 1. Subscripts are one bootstrap standard error (2000 sequence-level resamples). $^{\dagger}$Each CDR scored independently with full bidirectional context via IgLM's single-span masked format. Stars mark the best-vs-second-best gap per column (paired Wilcoxon, Holm-corrected across loops): $^{***}p < 0.001$, $^{**}p < 0.01$, $^{*}p < 0.05$.

| Model | Heavy chain (%) CDR1 | Heavy chain (%) CDR2 | Heavy chain (%) CDR3 | Light chain (%) CDR1 | Light chain (%) CDR2 | Light chain (%) CDR3 |
|---|---|---|---|---|---|---|
| *Input-framing ablation* | | | | | | |
| IgGenCDR (framework-first) | **92.01 ± .03**$^{***}$ | 89.32 ± .04 | **44.93 ± .05**$^{***}$ | **88.75 ± .07**$^{***}$ | 87.57 ± .08 | **58.75 ± .08**$^{***}$ |
| IgGenCDR-LTR (left-to-right) | 89.87 ± .03 | **89.41 ± .04**$^{***}$ | 43.67 ± .05 | 86.34 ± .07 | **87.59 ± .09** | 58.10 ± .08 |
| *Autoregressive baselines* | | | | | | |
| p-IgGen (left-to-right) (4) | 87.47 ± .04 | 84.04 ± .05 | 34.24 ± .03 | 82.72 ± .08 | 82.01 ± .09 | 50.38 ± .08 |
| p-IgGen (right-to-left) (4) | 88.43 ± .04 | 81.77 ± .05 | 37.90 ± .04 | 84.72 ± .08 | 81.40 ± .10 | 42.73 ± .08 |
| p-IgGen (developable, left-to-right) (4) | 87.63 ± .04 | 83.22 ± .05 | 33.94 ± .03 | 82.46 ± .08 | 81.74 ± .09 | 50.14 ± .08 |
| p-IgGen (developable, right-to-left) (4) | 88.29 ± .04 | 81.43 ± .05 | 37.17 ± .04 | 84.61 ± .08 | 81.28 ± .10 | 42.52 ± .08 |
| ProGen2-OAS | 87.56 ± .04 | 84.26 ± .05 | 35.53 ± .04 | 60.29 ± .18 | 52.22 ± .19 | 39.38 ± .12 |
| ProGen2-small | 72.78 ± .05 | 44.31 ± .06 | 14.03 ± .02 | 50.89 ± .11 | 50.71 ± .11 | 24.55 ± .08 |
| ProGen2-base | 83.37 ± .04 | 78.58 ± .06 | 26.64 ± .03 | 70.63 ± .12 | 70.64 ± .12 | 40.66 ± .09 |
| ProGen2-large | 84.09 ± .04 | 78.01 ± .06 | 26.40 ± .03 | 75.79 ± .12 | 73.34 ± .12 | 40.11 ± .09 |
| IgLM (CDR infill) | 88.66 ± .04$^{\dagger}$ | 84.77 ± .05$^{\dagger}$ | 33.53 ± .03$^{\dagger}$ | 85.33 ± .08$^{\dagger}$ | 83.13 ± .09$^{\dagger}$ | 53.17 ± .08$^{\dagger}$ |
| IgLM (full LTR) | 87.78 ± .04 | 84.31 ± .05 | 23.24 ± .03 | 84.57 ± .08 | 83.25 ± .09 | 41.08 ± .08 |
| IgLM-S (CDR infill) | 88.36 ± .04$^{\dagger}$ | 83.30 ± .05$^{\dagger}$ | 33.83 ± .04$^{\dagger}$ | 83.76 ± .08$^{\dagger}$ | 81.58 ± .10$^{\dagger}$ | 49.34 ± .08$^{\dagger}$ |
| IgLM-S (full LTR) | 87.57 ± .04 | 83.74 ± .05 | 25.28 ± .03 | 84.09 ± .08 | 82.46 ± .09 | 40.85 ± .09 |

DRAFT

**Table S7.** CDR amino acid recovery (NTP accuracy, %, Chothia numbering) on the CDR-disjoint unpaired test set, following the protocol of the main-text Table 1. Subscripts are one bootstrap standard error (2000 sequence-level resamples). $^{\dagger}$Each CDR scored independently with full bidirectional context via IgLM's single-span masked format. Stars mark the best-vs-second-best gap per column (paired Wilcoxon, Holm-corrected across loops): $^{***}p < 0.001$, $^{**}p < 0.01$, $^{*}p < 0.05$.

**Published baselines.** p-IgGen (17 M parameters) was fine-tuned for paired heavy–light generation on ~1.8 M paired sequences (4). ProGen2-OAS (151 M parameters) achieves competitive CDR1 and CDR2 accuracy on the heavy chain but collapses on light-chain CDR2 (43.88%), likely because its OAS training mixture is skewed towards heavy chains and lacks chain-type conditioning (3). IgLM (13 M parameters) was trained on 558 M multi-species sequences from OAS with a GPT-2 backbone and species-conditioning tokens (2).

**Bidirectional CDR context.** IgLM's CDR-infill mode provides bidirectional context for a single loop at a time. In this mode, heavy-chain CDR3 accuracy rises from 30.65% to 39.09% and light-chain CDR3 from 41.08% to 53.17%, confirming the value of surrounding-sequence conditioning. However, IgLM's masked-span format cannot provide simultaneous context for all three CDRs.

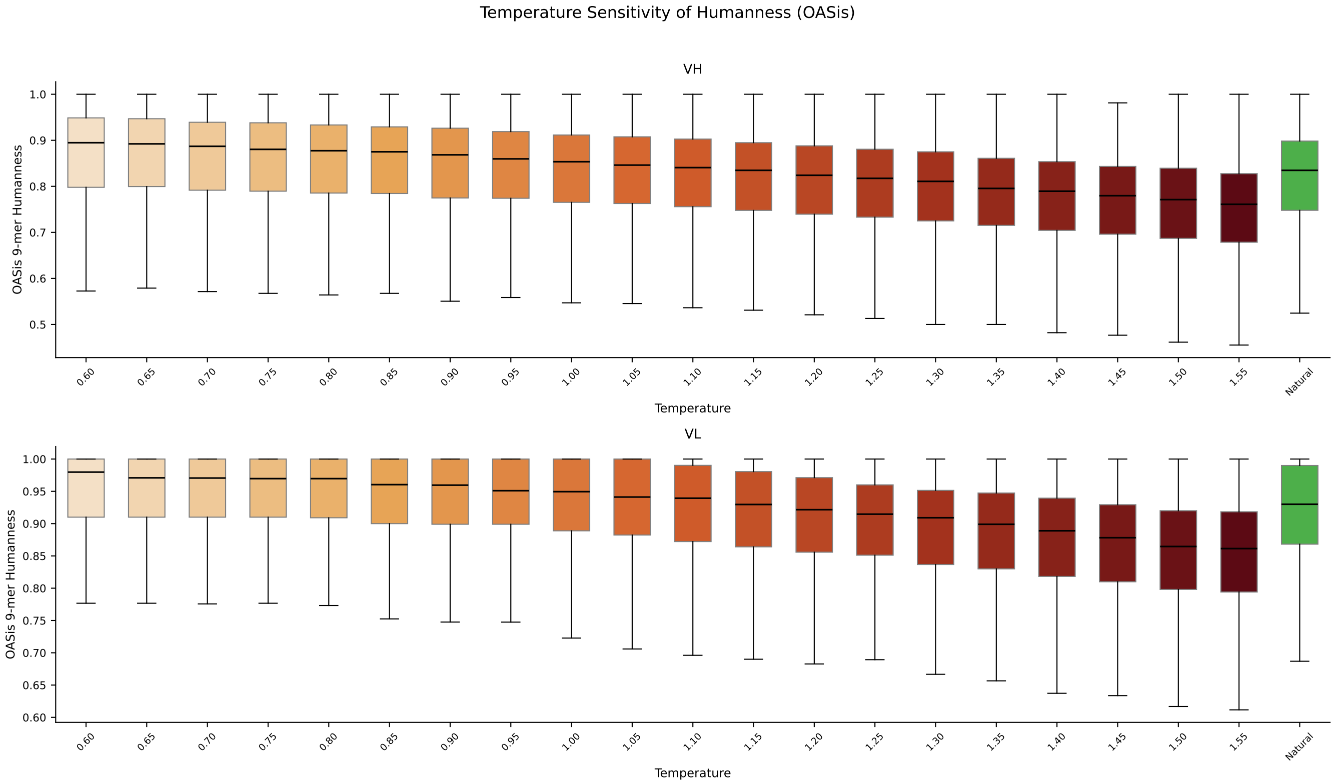


**Fig. S5.** OASis 9-mer humanness across the temperature sweep ($T = 0.60$–$1.55$) for heavy (top) and light (bottom) chains. The green box (far right) shows the natural test-set distribution. At $T \leq 1.0$ the generated distribution exceeds the natural reference; humanness degrades smoothly at higher temperatures.

### Supplementary Note S7: Generated CDR quality: full temperature sweep

This section collects the full temperature-sweep quality figures summarised in the IgGenCDR generation-quality results (main text). Across the sweep ($T = 0.60$–$1.55$), OASis 9-mer humanness (Figure S5) and ESM-2 pseudo-log-likelihood (Figure S7) degrade smoothly while sequence novelty (Figure S8) rises monotonically, and at $T = 1.0$ the generated ESM-2 distribution overlaps closely with the natural test set (Figure S6). Together these confirm that $T \in [0.90, 1.15]$ offers the best trade-off between naturalness and diversity.

DRAFT

### Supplementary Note S8: Zero-shot fitness correlation

We evaluate zero-shot fitness correlation on two assays from the FLAb benchmark collection (24, 25). Marks et al. (24) ($n$=217) measures the percentage of anti-drug antibody (%ADA) response of clinical-stage therapeutics to humanisation mutations, providing a proxy for immunogenicity, lower %ADA is preferable. Koenig et al. (25) ($n$=4,276) reports the expression ratio (ER) of single-point mutants of the G6.31 anti-VEGF antibody, where higher expression is preferable. Both datasets contain paired heavy and light chain sequences with a scalar fitness label.

Each sequence is scored by its mean per-token log-likelihood under the model:

$$\text{score}(x) = \frac{1}{|\mathcal{S}|} \sum_{i \in \mathcal{S}} \log p_\theta(x_i \mid x_{<i}), \tag{S5}$$

where $\mathcal{S}$ indexes either all variable-domain tokens (full-sequence scoring), the framework tokens only, or the CDR tokens only, giving the Full, Framework, and CDR scores reported throughout. For the framework-first models the CDR score uses the framework-conditioned factorisation $\log p_\theta(x_i^{\text{CDR}} \mid \text{FR}, x_{<i}^{\text{CDR}})$, the quantity directly optimised during training. Log-likelihoods are computed separately for the heavy and light chains, and the pair score is their arithmetic mean. CDR boundaries follow the indicated numbering scheme, and for each dataset the sign is adjusted so that a positive Spearman $\rho$ indicates correct model behaviour: for Marks et al. (24) immunogenicity, lower is better and for Koenig et al. (25) expression, higher is better. Reported values are averaged across IMGT, Kabat, and Chothia to marginalise over the arbitrary choice of scheme, with per-scheme results in Table S8.

IgGenCDR uses the framework-first factorisation: framework tokens condition CDR log-likelihoods, matching the training objective. IgGenCDR-LTR is the left-to-right ablation trained on identical data. Its full-sequence log-likelihood is scheme-

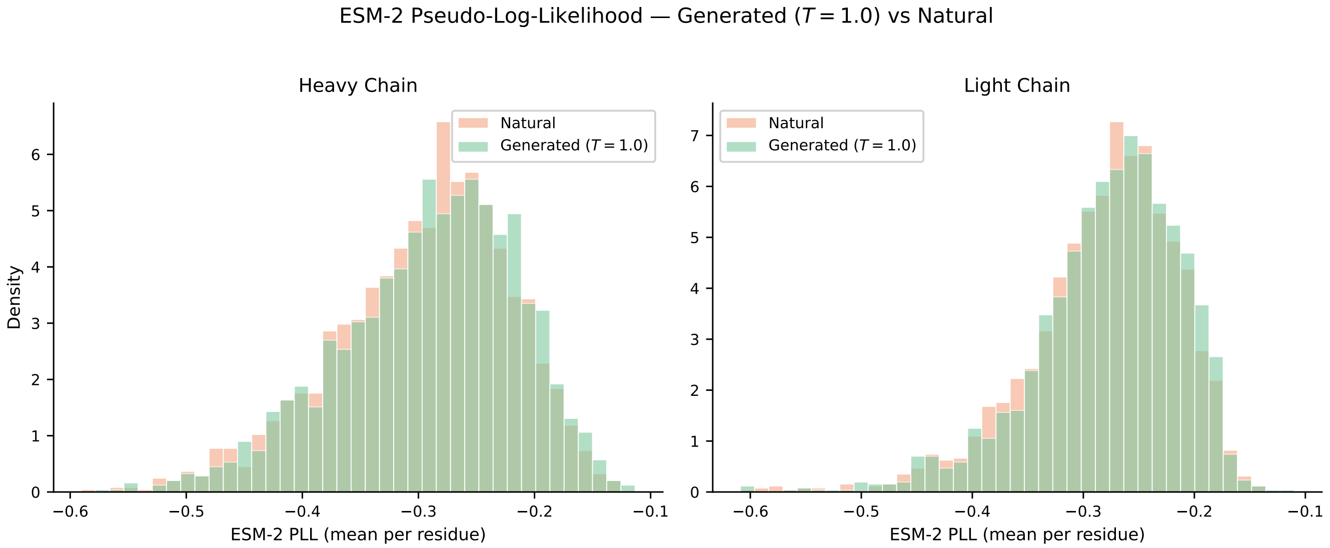


**Fig. S6.** ESM-2 pseudo-log-likelihood histograms for IgGenCDR-generated sequences at $T = 1.0$ (green) overlaid on natural OAS test-set sequences (coral). Heavy chain (left) and light chain (right). The two distributions overlap extensively, indicating comparable naturalness.

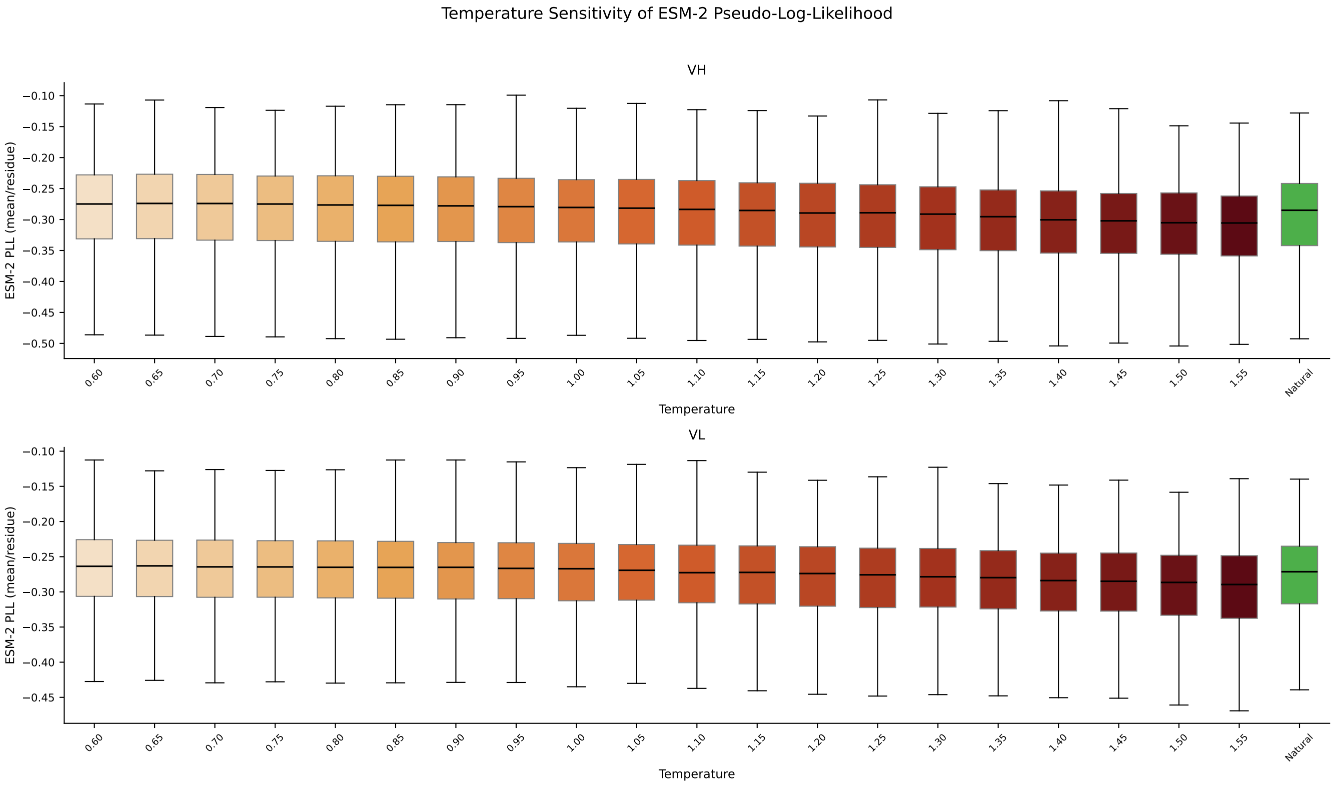


**Fig. S7.** ESM-2 pseudo-log-likelihood (PLL, mean per residue) across the temperature sweep for heavy (top) and light (bottom) chains. Natural test-set reference in green. PLL is stable up to $T \approx 1.0$ and declines smoothly at higher temperatures.

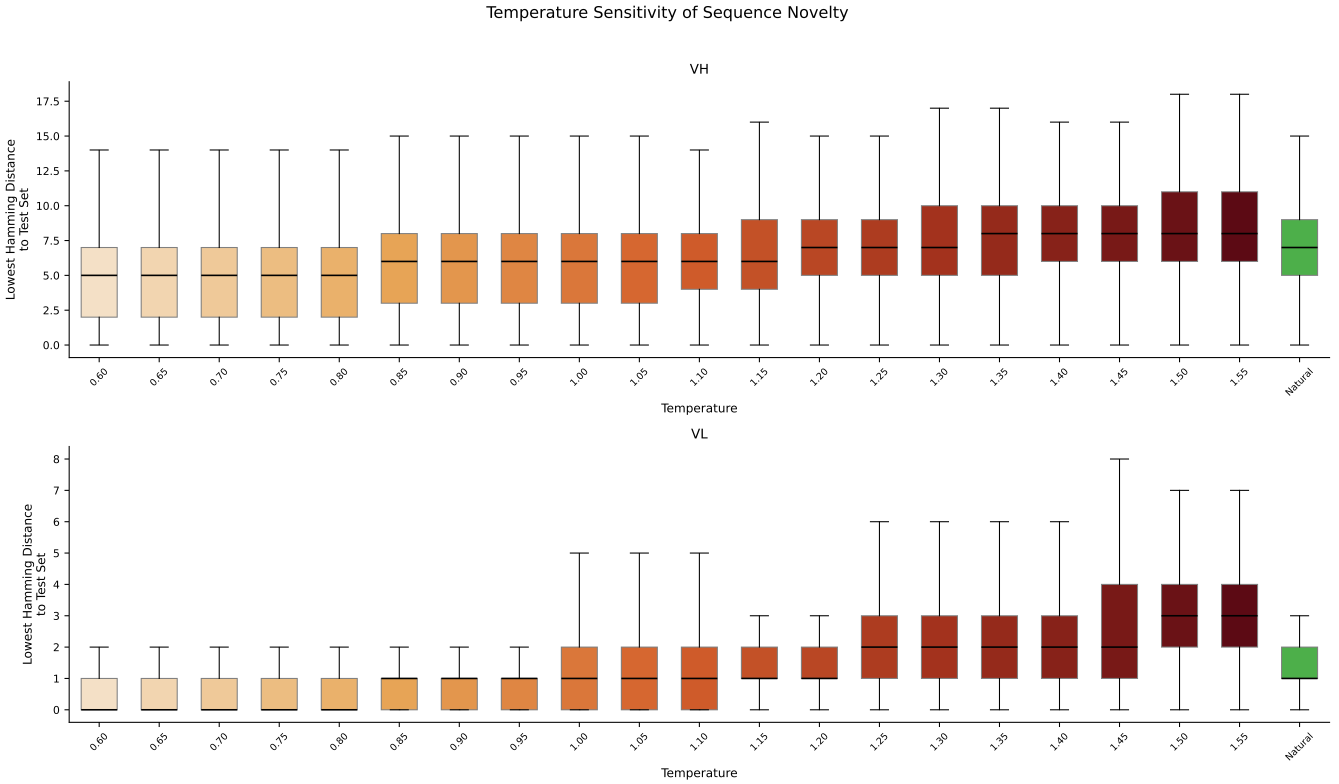


**Fig. S8.** Sequence novelty across the temperature sweep, measured as the minimum Hamming distance to the nearest training-set sequence. Heavy chain (top) and light chain (bottom). Natural test-set reference in green. Novelty increases smoothly with temperature.

independent (same token order regardless of numbering), but framework and CDR decompositions differ across schemes as FR/CDR boundaries change. IgLM (2) is a GPT-2 language model conditioned on chain type and species. It is the most architecturally similar infilling baseline and is evaluated in its standard full-sequence left-to-right mode (HEAVY/LIGHT + HUMAN prefix).

The scheme-averaged correlations are reported in the main text (Table 2), and the per-scheme breakdown is given in Table S8.

DRAFT

| | | Marks2021 (24) ($n$=217) | | | Koenig2017 (25) ($n$=4276) | | |
|---|---|---|---|---|---|---|---|
| Scheme | Model | Full | Framework | CDR | Full | Framework | CDR |
| IMGT | IgGenCDR | **0.353** | **0.329** | **0.240** | 0.209 | 0.199 | 0.062 |
| | IgGenCDR-LTR | 0.310 | 0.307 | 0.224 | **0.237** | **0.212** | 0.059 |
| | IgLM | 0.244 | 0.294 | 0.162 | 0.211 | 0.195 | **0.064** |
| KABAT | IgGenCDR | **0.338** | **0.286** | 0.207 | 0.235 | 0.183 | **0.105** |
| | IgGenCDR-LTR | 0.310 | 0.278 | **0.219** | **0.237** | **0.220** | 0.037 |
| | IgLM | 0.244 | 0.283 | 0.161 | 0.211 | 0.201 | 0.052 |
| CHOTHIA | IgGenCDR | **0.351** | **0.329** | **0.235** | **0.251** | 0.197 | **0.126** |
| | IgGenCDR-LTR | 0.310 | 0.304 | 0.231 | 0.237 | **0.205** | 0.084 |
| | IgLM | 0.244 | 0.304 | 0.166 | 0.211 | 0.191 | 0.078 |

**Table S8.** Per-scheme log-likelihood–fitness correlation (Spearman $\rho$) for paired heavy+light chain sequences. Region boundaries (FR/CDR) are defined by each numbering scheme. **Bold**: highest $\rho$ in each column within each numbering scheme.

### Supplementary Note S9: Paired IgGenCDR (p-IgGenCDR)

This section provides additional detail and per-scheme breakdowns for the paired model introduced in Section B. p-IgGenCDR is fine-tuned from the IgGenCDR checkpoint on ~1.8M natively paired VH–VL sequences from paired OAS only. Both chains are rendered as a single paired sequence in the framework-first layout (Figure S9), presenting all framework regions

`<BOS>` `<H> FR1 <SEP> ··· <SEP> FR4` `<L> FR1 <SEP> ··· <SEP> FR4` `<HPRED> CDR1 <SEP> CDR2 <SEP> CDR3` `<LPRED> CDR1 <SEP> CDR2 <SEP> CDR3` `<EOS>`

heavy framework — light framework — heavy CDR targets — light CDR targets

**Fig. S9.** Paired framework-first input representation (heavy-first shown). Both chains' framework regions form a single joint conditioning prompt, followed by the heavy and light CDR blocks, each preceded by its own generation-boundary token (<HPRED>, <LPRED>). The chain order (heavy-first or light-first) is sampled uniformly per training example. This extends the single-chain layout of the main text to two chains without architectural change.

of both chains first as a joint conditioning prompt, followed by the two CDR blocks, each introduced by its own generation-boundary token (<HPRED>, <LPRED>). This lets the model learn inter-chain co-evolutionary dependencies inaccessible to models trained on unpaired sequences. We also fine-tune a left-to-right ablation, p-IgGenCDR-LTR, from the IgGenCDR-LTR checkpoint under the same data and schedule, and include the unpaired IgGenCDR (base) checkpoint as an ablation. The full per-scheme recovery breakdown for all models follows in Table S9.

**A. CDR recovery: per-scheme breakdown.** Table S9 reports CDR recovery for all models under each numbering scheme (IMGT, Kabat, Chothia).

| | | Heavy chain (%) | | | Light chain (%) | | |
|---|---|---|---|---|---|---|---|
| **Scheme** | **Model** | CDR1 | CDR2 | CDR3 | CDR1 | CDR2 | CDR3 |
| IMGT | p-IgGenCDR | 90.46±.06 | **88.04±.07**$^{***}$ | **49.71±.06**$^{***}$ | **91.65±.05**$^{***}$ | **91.57±.07**$^{***}$ | **84.36±.05**$^{***}$ |
| | p-IgGenCDR-LTR | 90.27±.06 | 87.69±.07 | 48.58±.05 | 91.48±.06 | 91.30±.07 | 83.97±.05 |
| | IgGenCDR (base) | 90.02±.06 | 87.49±.08 | 47.91±.06 | 91.31±.06 | 90.89±.07 | 81.16±.05 |
| | p-IgGen (l-to-r) (4) | 88.11±.06 | 87.82±.07 | 46.89±.05 | 90.47±.06 | 91.33±.07 | 82.09±.05 |
| | p-IgGen (r-to-l) (4) | **90.60±.06**$^{***}$ | 85.58±.08 | 48.09±.06 | 91.58±.06 | 91.02±.07 | 73.52±.06 |
| | p-IgGen (dev., l-to-r) (4) | 88.02±.06 | 87.50±.07 | 46.69±.05 | 90.41±.06 | 91.23±.07 | 82.02±.05 |
| | p-IgGen (dev., r-to-l) (4) | 90.57±.06 | 85.34±.08 | 47.98±.06 | 91.56±.06 | 90.87±.07 | 73.30±.06 |
| Kabat | p-IgGenCDR | 87.84±.07 | **91.15±.05**$^{***}$ | 43.95±.06 | 93.82±.04 | **93.73±.04**$^{***}$ | **84.37±.05**$^{***}$ |
| | p-IgGenCDR-LTR | 87.47±.07 | 90.94±.05 | 42.66±.06 | 93.69±.04 | 93.56±.04 | 84.00±.05 |
| | IgGenCDR (base) | 87.18±.07 | 90.77±.05 | 41.92±.06 | 93.54±.04 | 93.30±.05 | 80.98±.05 |
| | p-IgGen (l-to-r) (4) | 84.06±.07 | 90.91±.05 | 40.73±.06 | 92.95±.04 | 93.58±.04 | 82.09±.05 |
| | p-IgGen (r-to-l) (4) | **88.19±.07**$^{***}$ | 89.32±.05 | **45.44±.06**$^{***}$ | **93.92±.04**$^{***}$ | 93.46±.04 | 73.52±.06 |
| | p-IgGen (dev., l-to-r) (4) | 83.94±.08 | 90.76±.05 | 40.50±.06 | 92.91±.04 | 93.53±.04 | 82.02±.05 |
| | p-IgGen (dev., r-to-l) (4) | 88.14±.07 | 89.18±.05 | 45.19±.06 | 93.90±.04 | 93.39±.04 | 73.30±.06 |
| Chothia | p-IgGenCDR | **90.85±.06**$^{***}$ | **86.91±.08**$^{***}$ | 43.95±.06 | 93.82±.04 | **93.73±.04**$^{***}$ | **84.38±.05**$^{***}$ |
| | p-IgGenCDR-LTR | 90.69±.06 | 86.55±.08 | 42.66±.06 | 93.68±.04 | 93.56±.04 | 84.01±.05 |
| | IgGenCDR (base) | 90.55±.06 | 86.28±.09 | 41.94±.06 | 93.54±.04 | 93.31±.05 | 80.97±.05 |
| | p-IgGen (l-to-r) (4) | 89.55±.06 | 86.36±.08 | 40.73±.06 | 92.95±.04 | 93.58±.04 | 82.09±.05 |
| | p-IgGen (r-to-l) (4) | 90.69±.06 | 84.30±.09 | **45.44±.06**$^{***}$ | **93.92±.04**$^{***}$ | 93.46±.04 | 73.52±.06 |
| | p-IgGen (dev., l-to-r) (4) | 89.46±.06 | 85.94±.08 | 40.50±.06 | 92.91±.04 | 93.53±.04 | 82.02±.05 |
| | p-IgGen (dev., r-to-l) (4) | 90.65±.06 | 84.01±.09 | 45.19±.06 | 93.90±.04 | 93.39±.04 | 73.30±.06 |

DRAFT

**Table S9.** Per-scheme CDR amino acid recovery (NTP accuracy, %) on the paired OAS test set ($\sim$69.8K pairs), for all models under each numbering scheme. p-IgGen and p-IgGen (developable) are the public baselines (4), evaluated on the same pairs in both decoding directions (l-to-r: `1+VH+VL+2`, r-to-l: `2+rev+1`). Subscripts are one bootstrap standard error (2000 sequence-level resamples). Stars mark the best-vs-second-best gap per column (paired Wilcoxon, Holm-corrected across the six loops) and sit on the column-best (bold) cell. Framework-first also exceeds its LTR ablation on all six loops under every scheme (paired Wilcoxon, Holm-corrected). $^{***}p < 0.001$, $^{**}p < 0.01$, $^{*}p < 0.05$.

**B. Generated CDR length distributions.** We assess *de novo* generation quality for p-IgGenCDR by prompting the model with 2,000 held-out paired framework scaffolds from the paired OAS test set and generating all six CDR loops at $T = 1.0$ (top-$p = 0.95$, IMGT numbering). Figure S10 compares the resulting CDR length distributions against the natural paired test-set sequences. As for the unpaired model (main-text Figure 2), the generated and natural distributions overlap closely across all six loops of both chains, confirming that paired fine-tuning preserves the length-distribution fidelity of the base model.

**C. Zero-shot fitness correlation on paired FLAb.** The scheme-averaged paired FLAb correlations are reported in the main text (Table 2), where they appear alongside the independent-chain (pair-mean) scores of the unpaired models. For p-IgGenCDR and p-IgGenCDR-LTR, the H+L pair is rendered as a single paired sequence (`render_paired`) and per-region log-likelihoods are computed jointly: *Full* sums over all amino-acid tokens of both chains, *Framework* over the H- and L-framework blocks, and *CDR* over the post-<HPRED> and post-<LPRED> blocks. The unpaired IgGenCDR baseline cannot be evaluated under this rendering because it has not been trained on the paired layout, so its independent-chain pair-mean values

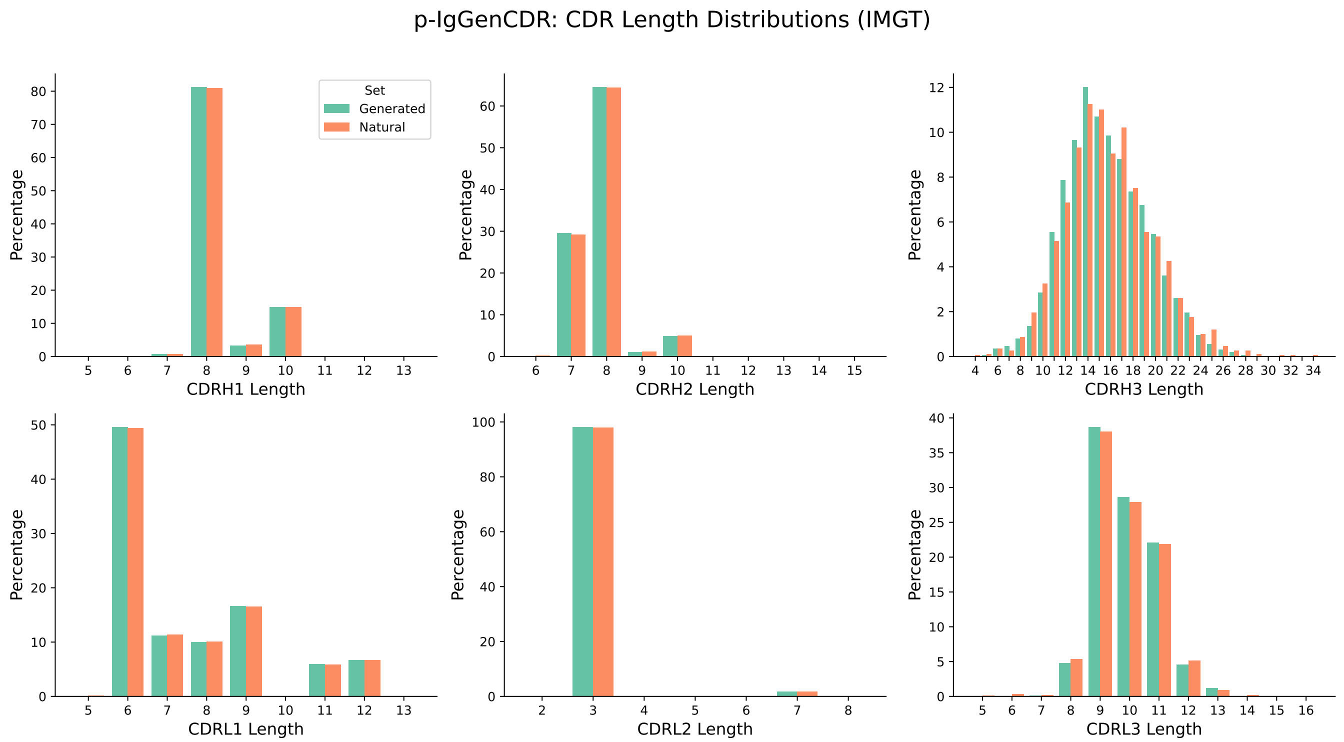


**Fig. S10.** CDR length distributions (IMGT) for p-IgGenCDR-generated sequences at $T = 1.0$ (green) versus natural paired OAS test-set sequences (coral). Heavy chain (top) and light chain (bottom); CDR1, CDR2, CDR3 from left to right. Bars show the per-length proportion within each set.

(from Table S8) are reproduced here for reference. Sign convention follows Section S8: positive Spearman $\rho$ indicates correct model behaviour. Table S10 gives the per-scheme breakdown.

DRAFT

| | | Marks2021 (24) ($n$=217) | | | Koenig2017 (25) ($n$=4276) | | |
|---|---|---|---|---|---|---|---|
| Scheme | Model | Full | Framework | CDR | Full | Framework | CDR |
| IMGT | IgGenCDR (base, pair-mean) | 0.353 | **0.329** | **0.240** | 0.209 | 0.199 | 0.062 |
| | p-IgGenCDR | **0.371** | 0.344 | 0.231 | 0.259 | 0.271 | −0.044 |
| | p-IgGenCDR-LTR | 0.367 | 0.338 | 0.270 | **0.349** | **0.309** | **0.050** |
| Kabat | IgGenCDR (base, pair-mean) | 0.338 | 0.286 | 0.207 | 0.235 | 0.183 | **0.105** |
| | p-IgGenCDR | 0.374 | **0.300** | 0.217 | **0.311** | 0.264 | −0.015 |
| | p-IgGenCDR-LTR | **0.382** | 0.289 | **0.270** | 0.307 | **0.271** | 0.002 |
| Chothia | IgGenCDR (base, pair-mean) | 0.351 | **0.329** | 0.235 | 0.251 | 0.197 | **0.126** |
| | p-IgGenCDR | **0.377** | 0.341 | 0.235 | 0.320 | 0.259 | 0.090 |
| | p-IgGenCDR-LTR | 0.369 | 0.324 | **0.285** | **0.324** | **0.268** | 0.071 |

**Table S10.** Per-scheme log-likelihood–fitness correlation (Spearman $\rho$) on FLAb. p-IgGenCDR and p-IgGenCDR-LTR use joint paired scoring, whereas the IgGenCDR (base) row reproduces the per-scheme pair-mean values from Table S8. Sign convention: positive $\rho$ indicates correct model behaviour. **Bold**: highest $\rho$ in each column within each numbering scheme.

### Supplementary Note S10: NanoGenCDR additional results

Table S11 reports the scheme-averaged CDR recovery on the INDI test set summarised in the main text (Section C), and Table S12 gives the per-scheme breakdown.

**A. Effect of the OAS mixture on CDR humanness.** This section expands the humanness comparison summarised in the main text (Section C, Figure 3, Table S13). NanoGenCDR is fine-tuned on a 1:1 mixture of camelid (INDI) and human OAS

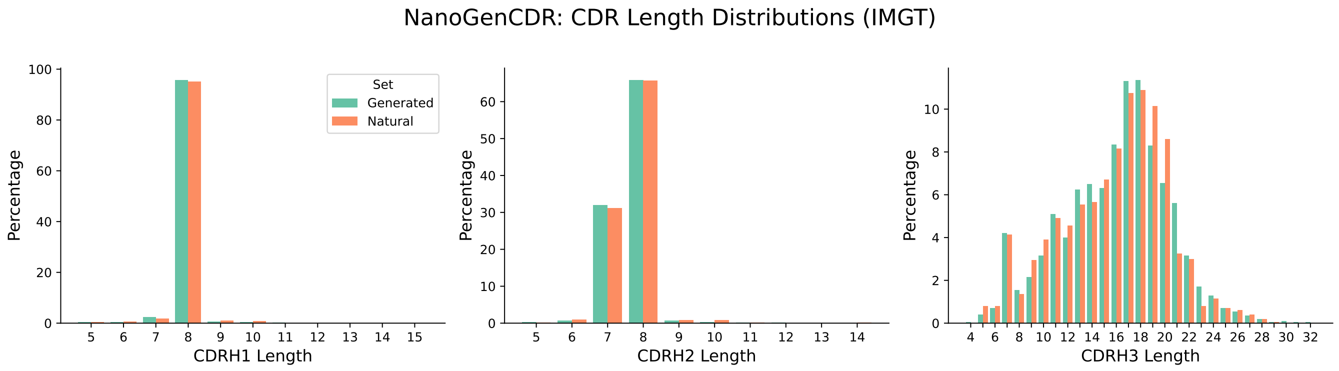


**Fig. S11.** CDR length distributions (IMGT) for NanoGenCDR generations from 2,000 INDI framework prompts at $T = 1.0$ (green) versus natural INDI test-set heavy-chain sequences (coral). CDR1, CDR2, CDR3 from left to right; bars show the per-length proportion within each set.

DRAFT

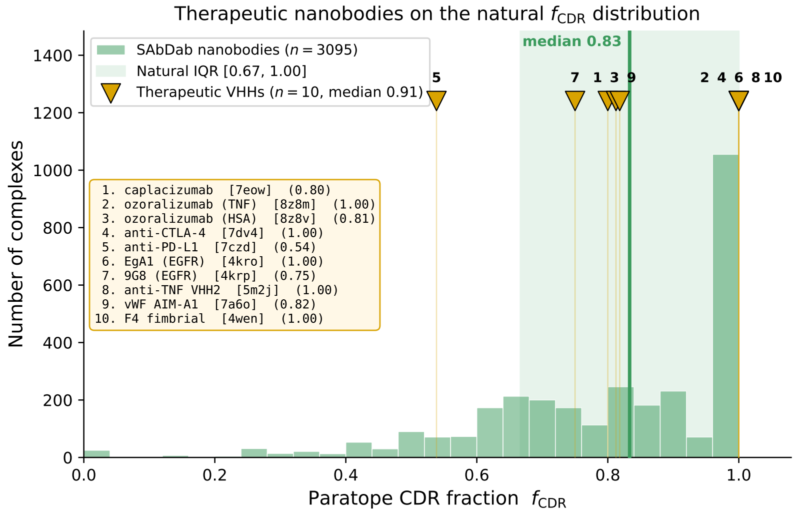


**Fig. S12.** Reference paratope CDR fraction ($f_{\mathrm{CDR}}$) over $3{,}095$ experimental nanobody–antigen complexes from SAbDab (29) (C$\beta$ contacts within 6 Å, IMGT). Gold triangles mark $10$ clinical/approved therapeutic VHHs (key inset, with PDB identifiers).

| Model | CDR1 (%) | CDR2 (%) | CDR3 (%) |
|---|---|---|---|
| NanoGenCDR | **79.03** $\pm$ **.03**$^{***}$ | **78.90** $\pm$ **.03**$^{***}$ | **56.96** $\pm$ **.04**$^{***}$ |
| NanoGenCDR-LTR | 74.95 $\pm$ .03 | 78.85 $\pm$ .03 | 55.58 $\pm$ .04 |
| IgGenCDR (base) | 39.68 $\pm$ .03 | 43.03 $\pm$ .02 | 13.34 $\pm$ .01 |
| IgLM$^{\dagger}$ (CDR3 infill, camelid prompt) | 44.64 $\pm$ .05$^{\ddagger}$ | 45.48 $\pm$ .04$^{\ddagger}$ | 20.46 $\pm$ .02$^{\ddagger}$ |
| IgLM-S$^{\dagger}$ (CDR3 infill, camelid prompt) | 44.70 $\pm$ .05$^{\ddagger}$ | 44.09 $\pm$ .04$^{\ddagger}$ | 20.44 $\pm$ .02$^{\ddagger}$ |

**Table S11.** CDR amino acid recovery (NTP accuracy, %) on the INDI nanobody test set, averaged across numbering schemes (IMGT, Kabat, Chothia). Subscripts are one bootstrap standard error (2000 sequence-level resamples, resampling unit is the test sequence aligned across schemes). Stars mark the best-vs-second-best gap per column on the scheme-averaged per-sequence recovery (paired Wilcoxon, Holm-corrected across the three loops) and sit on the column-best (bold) cell: $^{***}p < 0.001$, $^{**}p < 0.01$, $^{*}p < 0.05$. $^{\dagger}$IgLM is prompted with its camelid species token. $^{\ddagger}$IgLM evaluated under IMGT only with CDR3 masked, leveraging bidirectional context.

| Scheme | Model | CDR1 (%) | CDR2 (%) | CDR3 (%) |
|---|---|---|---|---|
| IMGT | NanoGenCDR | **78.96** $\pm$ **.05**$^{***}$ | **78.61** $\pm$ **.05**$^{***}$ | **58.49** $\pm$ **.06**$^{***}$ |
| | NanoGenCDR-LTR | 73.29 $\pm$ .05 | 78.16 $\pm$ .05 | 58.23 $\pm$ .06 |
| | IgGenCDR (base) | 44.58 $\pm$ .05 | 41.06 $\pm$ .04 | 15.12 $\pm$ .02 |
| Kabat | NanoGenCDR | **78.19** $\pm$ **.05** | 82.57 $\pm$ .03 | **56.18** $\pm$ **.07**$^{***}$ |
| | NanoGenCDR-LTR | 78.17 $\pm$ .05 | **83.46** $\pm$ **.03**$^{***}$ | 55.75 $\pm$ .07 |
| | IgGenCDR (base) | 26.30 $\pm$ .04 | 55.63 $\pm$ .03 | 12.47 $\pm$ .02 |
| Chothia | NanoGenCDR | **79.94** $\pm$ **.05**$^{***}$ | **75.51** $\pm$ **.06**$^{***}$ | **56.20** $\pm$ **.07**$^{***}$ |
| | NanoGenCDR-LTR | 73.38 $\pm$ .05 | 74.94 $\pm$ .06 | 55.75 $\pm$ .07 |
| | IgGenCDR (base) | 48.17 $\pm$ .05 | 32.41 $\pm$ .04 | 12.44 $\pm$ .02 |

**Table S12.** Per-scheme CDR amino acid recovery (NTP accuracy, %) on the INDI nanobody test set. IgLM and IgLM-S are evaluated under IMGT numbering only (CDR3-masked, bidirectional context) and are therefore reported only in the scheme-averaged Table S11. Subscripts are one bootstrap standard error (2000 sequence-level resamples). Stars mark the best-vs-second-best gap per column (paired Wilcoxon, Holm-corrected across the three loops) and sit on the column-best (bold) cell: $^{***}p < 0.001$, $^{**}p < 0.01$, $^{*}p < 0.05$.

DRAFT

sequences (Section S1). To test whether the human OAS fraction improves the humanness of generated CDRs, we compare it against an otherwise-identical checkpoint trained on camelid sequences only (*INDI-only*), sharing the same initialisation, schedule and seed and differing solely in the OAS mixture. Both models complete CDRs ($T$=1.0, IMGT) on five humanised VHH scaffolds: four humanised camelid frameworks from Stark et al. (9) and the framework of ozoralizumab, an approved anti-TNF$\alpha$ humanised VHH therapeutic. These scaffolds are hybrids that combine a human VH3-derived framework with retained camelid hallmark and loop-supporting residues, and are the practical substrate for therapeutic nanobody design. The ozoralizumab framework was taken from its sequence (deposited in SAbDab (29)) and is representative of the humanised synthetic VHH frameworks now used to build therapeutic libraries (34), and we segment it into FR1–FR4 with ANARCI (17) under IMGT. We generate 500 sequences per framework per model on identical prompts, and score humanness with OASis (22), restricted to the 9-mers overlapping the generated CDRs so that the shared framework does not dilute the signal. Per-framework means are given in Table S13.

| Framework | NanoGenCDR (mix) | INDI-only | Sig. |
|---|---|---|---|
| boltz_2 | **0.459** $\pm$ 0.102 | 0.376 $\pm$ 0.093 | *** |
| boltz_3 | **0.377** $\pm$ 0.062 | 0.360 $\pm$ 0.060 | *** |
| boltz_4 | **0.446** $\pm$ 0.094 | 0.338 $\pm$ 0.072 | *** |
| boltz_5 | **0.541** $\pm$ 0.076 | 0.498 $\pm$ 0.074 | *** |
| ozoralizumab | **0.531** $\pm$ 0.060 | 0.480 $\pm$ 0.063 | *** |
| All | **0.471** $\pm$ 0.101 | 0.410 $\pm$ 0.098 | *** |

**Table S13.** CDR-region OASis humanness (mean $\pm$ s.d., $n$=500 per framework per model) of CDRs generated on humanised VHH scaffolds. The OAS mixture yields significantly more human CDRs on every framework. Significance from a two-sided Mann–Whitney $U$ test: $^{*}p$<0.05, $^{**}p$<0.01, $^{***}p$<0.001. **Bold**: higher mean.

The per-framework means (Table S13) show the OAS-mixture gain holds on every individual humanised scaffold, and on camelid INDI frameworks the two models are indistinguishable (median OASis 0.31 for both), as expected when the scaffold itself is non-human.

The effect is even more pronounced on a fully humanised acceptor framework. Repeating the experiment on the human IGHV3-23 (DP-47) germline framework, the canonical human acceptor toward which camelid VHHs are humanised but which retains no camelid hallmark residues and is therefore not included in the humanised-VHH average above, the OAS mixture reaches an OASis of $0.864 \pm 0.061$ versus $0.691 \pm 0.071$ for the INDI-only model ($n$=500 each, $p<10^{-100}$, two-sided Mann–Whitney $U$). As the scaffold approaches a fully human framework, the human-distribution signal retained by the mixture becomes increasingly decisive.

**B. Effect of the OAS mixture on recovery: a favourable trade-off.** Table S14 details the recovery trade-off summarised in the main text (Section C), comparing NanoGenCDR (mix) and the INDI-only checkpoint on next-token CDR recovery (IMGT, heavy) on the camelid INDI and unpaired human OAS test sets. The collapse on human OAS is sharpest on the discriminative CDR3 loop, where the INDI-only model falls to 24.34% against the mixture's 60.53%, indicating that camelid-only fine-tuning largely forgets the human CDR3 distribution while the mixture preserves it.

| Test set | Model | CDR1 | CDR2 | CDR3 | Mean |
|---|---|---|---|---|---|
| INDI (camelid) | NanoGenCDR (mix) | 78.96 $\pm$ .05 | 78.60 $\pm$ .05 | 58.49 $\pm$ .06 | 72.02 $\pm$ .04 |
| | INDI-only | **79.88 $\pm$ .05**$^{***}$ | **79.75 $\pm$ .05**$^{***}$ | **60.47 $\pm$ .06**$^{***}$ | **73.37 $\pm$ .04**$^{***}$ |
| OAS (human) | NanoGenCDR (mix) | **90.94 $\pm$ .02**$^{***}$ | **89.47 $\pm$ .02**$^{***}$ | **60.53 $\pm$ .03**$^{***}$ | **80.31 $\pm$ .02**$^{***}$ |
| | INDI-only | 62.90 $\pm$ .03 | 45.33 $\pm$ .03 | 24.34 $\pm$ .01 | 44.19 $\pm$ .02 |

**Table S14.** CDR amino acid recovery (NTP accuracy, %, IMGT, heavy) for the OAS-mixed vs INDI-only NanoGenCDR checkpoints on camelid (INDI) and human (OAS) test sets. The mixture costs $1.35$ mean points on camelid but retains $+36$ mean points on human relative to the camelid-only model. Subscripts are one bootstrap standard error (2000 sequence-level resamples, resampling unit is the test sequence). Stars mark the mix vs INDI-only gap per loop (paired Wilcoxon on per-sequence recovery, Holm-corrected across the three loops and the mean) and sit on the better model: $^{***}p < 0.001$, $^{**}p < 0.01$, $^{*}p < 0.05$.

**C. Zero-shot fitness correlation on NbBench.** We evaluate zero-shot fitness correlation on two nanobody thermostability assays from the NbBench collection (28): thermo-seq ($n$=764), measuring sequencing-based enrichment labels, and thermo-tm ($n$=567), measuring melting temperature ($T_m$), with higher values preferable for both. All datasets contain single heavy-chain (VHH) sequences with a scalar fitness label.

Sequences are scored by mean per-token log-likelihood (Eq. S5) computed over the single heavy chain. Region-specific scores (Full, Framework, CDR) use the same token partitioning as in Section S8, with CDR boundaries defined by the indicated numbering scheme. For framework-first models (NanoGenCDR, IgGenCDR), the CDR log-likelihood is conditioned on all framework tokens, matching the training objective. IgLM (2) is evaluated in full-sequence left-to-right mode with its camelid species token.

| | thermo-seq ($n$=764) | | | thermo-tm ($n$=567) | | |
|---|---|---|---|---|---|---|
| Model | Full | Framework | CDR | Full | Framework | CDR |
| NanoGenCDR | **0.255** | **0.331** | **0.105** | **0.245** | **0.325** | 0.100 |
| IgGenCDR (base) | 0.159 | 0.264 | −0.004 | 0.176 | 0.280 | 0.028 |
| IgLM | 0.160 | 0.211 | 0.150 | 0.191 | 0.224 | **0.215** |

**Table S15.** Log-likelihood–fitness correlation (Spearman $\rho$) on NbBench nanobody benchmarks, averaged across numbering schemes (IMGT, Kabat, Chothia). **Bold**: highest $\rho$ in each column.

DRAFT

| Scheme | Model | thermo-seq ($n$=764) | | | thermo-tm ($n$=567) | | |
|---|---|---|---|---|---|---|---|
| | | Full | Framework | CDR | Full | Framework | CDR |
| IMGT | NanoGenCDR | **0.241** | **0.316** | 0.101 | **0.234** | **0.313** | 0.094 |
| | IgGenCDR (base) | 0.178 | 0.261 | 0.071 | 0.197 | 0.282 | 0.088 |
| | IgLM | 0.160 | 0.184 | **0.232** | 0.191 | 0.194 | **0.300** |
| Kabat | NanoGenCDR | **0.271** | **0.312** | 0.133 | **0.261** | **0.312** | 0.124 |
| | IgGenCDR (base) | 0.130 | 0.224 | −0.007 | 0.154 | 0.233 | 0.039 |
| | IgLM | 0.160 | 0.196 | **0.157** | 0.191 | 0.217 | **0.175** |
| Chothia | NanoGenCDR | **0.247** | **0.355** | 0.074 | **0.237** | **0.342** | 0.078 |
| | IgGenCDR (base) | 0.163 | 0.294 | −0.074 | 0.169 | 0.305 | −0.051 |
| | IgLM | 0.160 | 0.246 | **0.051** | 0.191 | 0.245 | **0.149** |

**Table S16.** Per-scheme Spearman $\rho$ between log-likelihood and label on NbBench thermostability benchmarks. **Bold**: highest $\rho$ in each column within each numbering scheme.

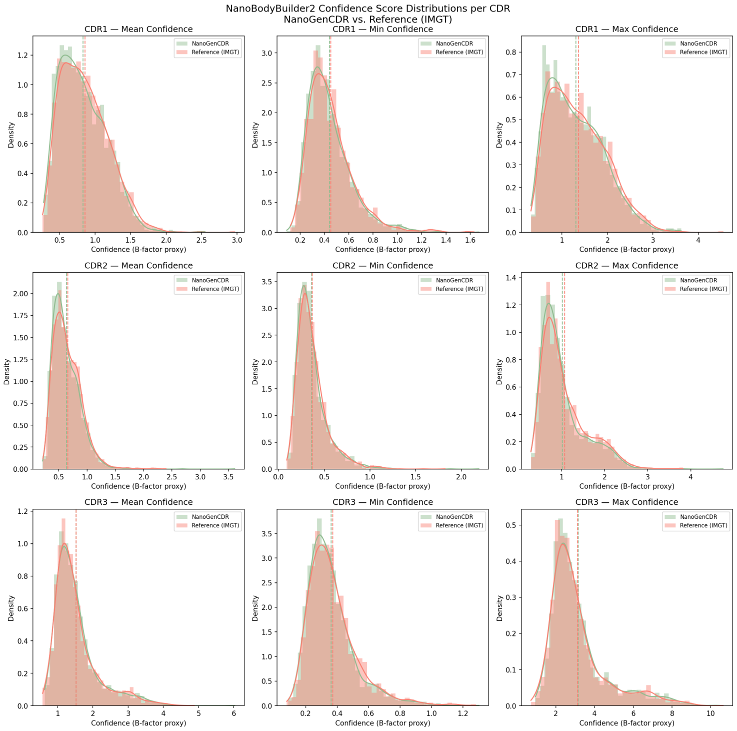


**Fig. S13.** NanoBodyBuilder2 (26) mean confidence score per each CDR (IMGT), with minimum and maximum confidence values, based on b-factor proxy. NanoGenCDR results are overlaid against Reference set.

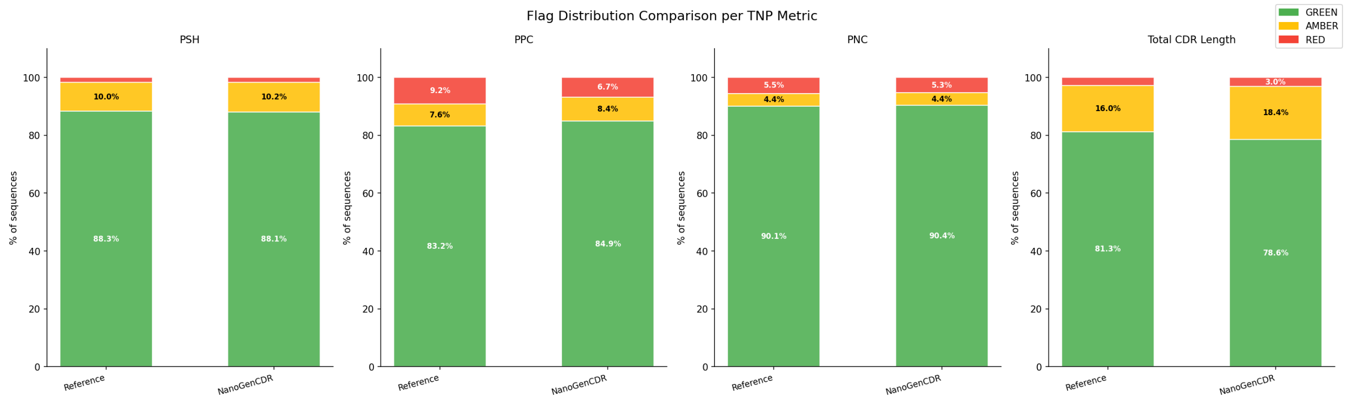


**Fig. S14.** Developability flag distributions for NanoGenCDR versus the reference set, computed with the Therapeutic Nanobody Profiler (TNP) (27). PSH (Patch CDR Surface Hydrophobicity) flags hydrophobic surface patches on the CDRs that predispose to aggregation. PPC and PNC (Patch Positive and Negative Charge) flag charged CDR patches associated with poor specificity and clearance. Total IMGT CDR length flags unusually long combined loops, which tend to be harder to express and develop. Each metric is binned into TNP green/amber/red risk bands, and the plot compares the proportion of designs in each band against the reference cohort.
